# Brain charts for the human lifespan

**DOI:** 10.1101/2021.06.08.447489

**Authors:** R.A.I. Bethlehem, J. Seidlitz, S.R. White, J.W. Vogel, K.M. Anderson, C. Adamson, S. Adler, G.S. Alexopoulos, E. Anagnostou, A. Areces-Gonzalez, D.E. Astle, B. Auyeung, M. Ayub, G. Ball, S. Baron-Cohen, R. Beare, S.A. Bedford, V. Benegal, F. Beyer, J. Bin Bae, J. Blangero, M. Blesa Cábez, J.P. Boardman, M. Borzage, J.F. Bosch-Bayard, N. Bourke, V.D. Calhoun, M.M. Chakravarty, C. Chen, C. Chertavian, G. Chetelat, Y.S. Chong, J.H. Cole, A. Corvin, M. Costantino, E. Courchesne, F. Crivello, V.L. Cropley, J. Crosbie, N. Crossley, M. Delarue, R. Delorme, S. Desrivieres, G. Devenyi, M.A. Di Biase, R. Dolan, K.A. Donald, G. Donohoe, K. Dunlop, A.D. Edwards, J.T. Elison, C.T. Ellis, J.A. Elman, L. Eyler, D.A. Fair, E. Feczko, P.C. Fletcher, P. Fonagy, C.E. Franz, L. Galan-Garcia, A. Gholipour, J. Giedd, J.H. Gilmore, D.C. Glahn, I. Goodyer, P.E. Grant, N.A. Groenewold, F.M. Gunning, R.E. Gur, R.C. Gur, C.F. Hammill, O. Hansson, T. Hedden, A. Heinz, R.N. Henson, K. Heuer, J. Hoare, B. Holla, A.J. Holmes, R. Holt, H. Huang, K. Im, J. Ipser, C.R. Jack, A.P. Jackowski, T. Jia, K.A. Johnson, P.B. Jones, D.T. Jones, R. Kahn, H. Karlsson, L. Karlsson, R. Kawashima, E.A. Kelley, S. Kern, K. Kim, M.G. Kitzbichler, W.S. Kremen, F. Lalonde, B. Landeau, S. Lee, J. Lerch, J.D. Lewis, J. Li, W. Liao, C. Liston, M.V. Lombardo, J. Lv, C. Lynch, T.T. Mallard, M. Marcelis, R.D. Markello, S.R. Mathias, B. Mazoyer, P. McGuire, M.J. Meaney, A. Mechelli, N. Medic, B. Misic, S.E. Morgan, D. Mothersill, J. Nigg, M.Q.W. Ong, C. Ortinau, R. Ossenkoppele, M. Ouyang, L. Palaniyappan, L. Paly, P.M. Pan, C. Pantelis, M.M. Park, T. Paus, Z. Pausova, D. Paz-Linares, A. Pichet Binette, K. Pierce, X. Qian, J. Qiu, A. Qiu, A. Raznahan, T. Rittman, A. Rodrigue, C.K. Rollins, R. Romero-Garcia, L. Ronan, M.D. Rosenberg, D.H. Rowitch, G.A. Salum, T.D. Satterthwaite, H.L. Schaare, R.J. Schachar, A.P. Schultz, G. Schumann, M. Schöll, D. Sharp, R.T. Shinohara, I. Skoog, C.D. Smyser, R.A. Sperling, D.J. Stein, A. Stolicyn, J. Suckling, G. Sullivan, Y. Taki, B. Thyreau, R. Toro, N. Traut, K.A. Tsvetanov, N.B. Turk-Browne, J.J. Tuulari, C. Tzourio, É. Vachon-Presseau, M.J. Valdes-Sosa, P.A. Valdes-Sosa, S.L. Valk, T. van Amelsvoort, S.N. Vandekar, L. Vasung, L.W. Victoria, S. Villeneuve, A. Villringer, P.E. Vértes, K. Wagstyl, Y.S. Wang, S.K. Warfield, V. Warrier, E. Westman, M.L. Westwater, H.C. Whalley, A.V. Witte, N. Yang, B. Yeo, H. Yun, A. Zalesky, H.J. Zar, A. Zettergren, J.H. Zhou, H. Ziauddeen, A. Zugman, X.N. Zuo, 3R-BRAIN, AIBL, Alzheimer’s Disease Neuroimaging Initiative, Alzheimer’s Disease Repository Without Borders Investigators, UMN BCP, CALM Team, Cam-CAN, CCNP, COBRE, Developing Human Connectome Project, ENIGMA Developmental Brain Age working group, FinnBrain, Harvard Aging Brain Study, IMAGEN, KNE96, The Mayo Clinic Study of Aging, NSPN, POND, The PREVENT-AD Research Group, VETSA, E.T. Bullmore, A.F. Alexander-Bloch

**Affiliations:** Autism Research Centre, Department of Psychiatry, University of Cambridge, Cambridge, CB2 0SZ, UK; Brain Mapping Unit, Department of Psychiatry, University of Cambridge, Cambridge, CB2 0SZ, UK; Department of Psychiatry, University of Pennsylvania, Philadelphia, PA 19104; Department of Child and Adolescent Psychiatry and Behavioral Science, The Children’s Hospital of Philadelphia, Philadelphia, PA 19104; Department of Psychiatry, University of Cambridge, Cambridge, CB2 0SZ, UK; MRC Biostatistics Unit, University of Cambridge, Cambridge, England; Lifespan Informatics & Neuroimaging Center, University of Pennsylvania, Philadelphia, PA 19104; Department of Psychology, Yale University, New Haven, CT, USA; Developmental Imaging, Murdoch Children’s Research Institute, Melbourne, Victoria, Australia; Department of Medicine, Monash University, Melbourne, Victoria, Australia; UCL Great Ormond Street Institute for Child Health, 30 Guilford St, Holborn, London WC1N 1EH; Weill Cornell Institute of Geriatric Psychiatry, Department of Psychiatry, Weill Cornell Medicine; Department of Pediatrics University of Toronto; Holland Bloorview Kids Rehabilitation Hospital, Toronto, Canada; The Clinical Hospital of Chengdu Brain Science Institute, MOE Key Lab for NeuroInformation, University of Electronic Science and Technology of China, No. 2006, Xiyuan Ave., West Hi-Tech Zone, Chengdu, 611731, China; University of Pinar del Río “Hermanos Saiz Montes de Oca”, Cuba; MRC Cognition and Brain Sciences Unit, University of Cambridge, Cambridge UK; Department of Psychology, School of Philosophy, Psychology and Language Sciences, University of Edinburgh, Edinburgh, United Kingdom; University College London, Mental Health Neuroscience Research Department, Division of Psychiatry, London UK; Department of Paediatrics, University of Melbourne, Melbourne, Victoria, Australia; Cambridge Lifetime Asperger Syndrome Service (CLASS), Cambridgeshire and Peterborough NHS Foundation Trust, Cambridge, United Kingdom; Centre for Addiction Medicine, National Institute of Mental Health and Neurosciences, Bengaluru, India 560029; Department of Neurology, Max Planck Institute for Human Cognitive and Brain Sciences, Leipzig, 04103, Germany; Department of Neuropsychiatry, Seoul National University Bundang Hospital, Seongnam, Korea; Department of Human Genetics, South Texas Diabetes and Obesity Institute, University of Texas Rio Grande Valley; MRC Centre for Reproductive Health, University of Edinburgh, UK; Fetal and Neonatal Institute, Division of Neonatology, Children’s Hospital Los Angeles, Department of Pediatrics, Keck School of Medicine, University of Southern California, Los Angeles, California USA; McGill Centre for Integrative Neuroscience, Ludmer Centre for Neuroinformatics and Mental Health, Montreal Neurological Institute; McGill University; Department of Brain Sciences, Imperial College London, London UK & Care Research & Technology Centre, UK Dementia Research Institute; Tri-institutional Center for Translational Research in Neuroimaging and Data Science, Georgia State University, Georgia Institute of Technology, and Emory University, Atlanta, GA, USA; Computational Brain Anatomy (CoBrA) Laboratory, Cerebral Imaging Centre, Douglas Mental Health University Institute; Penn Statistics in Imaging and Visualization Center, Department of Biostatistics, Epidemiology, and Informatics, Perelman School of Medicine, University of Pennsylvania, Philadelphia, PA, USA; Normandie Univ, UNICAEN, INSERM, U1237, PhIND “Physiopathology and Imaging of Neurological Disorders”, Institut Blood and Brain @ Caen-Normandie, Cyceron, 14000 Caen, France; Singapore Institute for Clinical Sciences, Agency for Science, Technology and Research, Singapore; Department of Obstetrics and Gynaecology, Yong Loo Lin School of Medicine, National University of Singapore, Singapore; Centre for Medical Image Computing (CMIC), University College London; Dementia Research Centre (DRC), University College London; Department of Psychiatry, Trinity College, Dublin, Ireland; Cerebral Imaging Centre, Douglas Mental Health University Institute, Verdun, Canada; Undergraduate program in Neuroscience, McGill University, Montreal, Canada; Department of Neuroscience, University of California, San Diego, San Diego, CA 92093, USA; Autism Center of Excellence, University of California, San Diego, San Diego, CA 92037, USA; Institute of Neurodegenerative Disorders, CNRS UMR5293, University of Bordeaux; Melbourne Neuropsychiatry Centre, University of Melbourne, Melbourne, Australia; The Hospital for Sick Children, Toronto, Canada; Department of Psychiatry, School of Medicine, Pontificia Universidad Católica de Chile, Diagonal Paraguay 362, Santiago 8330077, Chile; Department of Psychosis Studies, Institute of Psychiatry, Psychology and Neuroscience, King’s College London, De Crespigny Park, London SE5 8AF, UK; Child and Adolescent Psychiatry Department, Robert Debré University Hospital, AP-HP, F-75019, Paris France; Human Genetics and Cognitive Functions, Institut Pasteur, F-75015, Paris France; Social, Genetic and Developmental Psychiatry Centre, Institute of Psychiatry, Psychology & Neuroscience, King’s College London, London, United Kingdom; Cerebral Imaging Centre, Douglas Mental Health University Institute, Montreal, QC, Canada; Department of Psychiatry, McGill University, Montreal, QC, Canada; Department of Psychiatry, Brigham and Women’s Hospital, Harvard Medical School, Boston, Massachusetts, United States; Max Planck UCL Centre for Computational Psychiatry and Ageing Research, University College London, London, UK; Wellcome Centre for Human Neuroimaging, University College London, London, UK; University of Cape Town, South Africa, Cape Town, South Africa; Center for Neuroimaging, Cognition & Genomics (NICOG), School of Psychology, National University of Ireland Galway, Galway, Ireland; Weil Family Brain and Mind Research Institute, Department of Psychiatry, Weill Cornell Medicine; Centre for the Developing Brain, King’s College London, London, UK; Evelina London Children’s Hospital; Institute of Child Development, Department of Pediatrics, Masonic Institute for the Developing Brain, University of Minnesota, Minneapolis, MN, United States; Department of Psychiatry, Center for Behavior Genetics of Aging, University of California, San Diego, La Jolla, CA; Desert-Pacific Mental Illness Research Education and Clinical Center, VA San Diego Healthcare, San Diego, CA, USA; Department of Psychiatry, University of California San Diego, Los Angeles, CA, USA; Department of Psychiatry, University of Cambridge, and Wellcome Trust MRC Institute of Metabolic Science, Cambridge Biomedical Campus, Cambridge, United Kingdom; Cambridgeshire and Peterborough NHS Foundation Trust; Department of Clinical, Educational and Health Psychology, University College London, London, UK; Department of Psychiatry, Center for Behavior Genetics of Aging, University of California, San Diego, La Jolla, CA 92093; Cuban Center for Neuroscience, La Habana, Cuba; Computational Radiology Laboratory, Boston Children’s Hospital, Boston, MA 02115; Department of Child and Adolescent Psychiatry, University of California, San Diego, San Diego, CA 92093, USA; Department of Psychiatry, University of California San Diego, San Diego, CA, USA; Department of Psychiatry, University of North Carolina, Chapel Hill, NC, USA; Department of Psychiatry, Boston Children’s Hospital and Harvard Medical School, Boston, MA 02115; Harvard Medical School, Boston, MA 02115; Division of Newborn Medicine and Neuroradiology, Fetal Neonatal Neuroimaging and Developmental Science Center, Boston Children’s Hospital, Harvard Medical School, Boston, MA 02115, USA; Department of Paediatrics and Child Health, Red Cross War Memorial Children’s Hospital, SA-MRC Unit on Child & Adolescent Health, University of Cape Town, South Africa; Neuroscience Institute, University of Cape Town, Cape Town, South Africa; Lifespan Brain Institute, The Children’s Hospital of Philadelphia, Philadelphia, PA 19104; Lifespan Brain Institute, The Children’s Hospital of Philadelphia, Philadelphia, PA 19105; Mouse Imaging Centre, Toronto, Canada; Clinical Memory Research Unit, Department of Clinical Sciences Malmö, Lund University, Malmö, Sweden; Memory Clinic, Skåne University Hospital, Malmö, Sweden; Department of Neurology, Icahn School of Medicine at Mount Sinai, New York, NY 10029, USA; Athinoula A. Martinos Center for Biomedical Imaging, Department of Radiology, Massachusetts General Hospital, Harvard Medical School, Boston, MA 02129, USA; Department of Psychiatry and Psychotherapy, Charite University Hospital Berlin, Berlin, Germany; Department of Psychiatry, University of Cambridge, Cambridge, UK; Department of Neuropsychology, Max Planck Institute for Human Cognitive and Brain Sciences, Leipzig, Germany; Université de Paris, Paris, France; Department of Psychiatry, University of Cape Town, Cape Town, South Africa; Departments of Psychiatry and Integrative Medicine, NIMHANS, Bengaluru, India; Departments of Psychology and Psychiatry, Yale University, New Haven, CT, USA; Radiology Research, Children’s Hospital of Philadelphia, Philadelphia, United States; The Department of Radiology, Perelman School of Medicine, University of Pennsylvania, Philadelphia, United States; Division of Newborn Medicine, Fetal Neonatal Neuroimaging and Developmental Science Center, Boston Children’s Hospital, Harvard Medical School, Boston, MA 02115, USA; Boston Children’s Hospital, Boston, MA 02115; Department of Psychiatry and Mental Health, Clinical Neuroscience Institute, University of Cape Town; Department of Radiology, Mayo Clinic, Rochester, MN 55905, USA; Department of Psychiatry, Universidade Federal de São Paulo; National Institute of Developmental Psychiatry, CNPq; Institute of Science and Technology for Brain-Inspired Intelligence, Fudan University, Shanghai, 200433, China; Key Laboratory of Computational Neuroscience and BrainInspired Intelligence (Fudan University), Ministry of Education, Shanghai, China; Centre for Population Neuroscience and Precision Medicine (PONS), Institute of Psychiatry, Psychology and Neuroscience, SGDP Centre, King’s College London, London SE5 8AF, UK; Harvard Aging Brain Study, Department of Neurology, Massachusetts General Hospital, Boston, MA 02114; Center for Alzheimer Research and Treatment, Department of Neurology, Brigham and Women’s Hospital, Boston, MA 02115; Department of Radiology, Massachusetts General Hospital, Boston, MA; Cambridgeshire and Peterborough NHS Foundation Trust, Huntingdon, United Kingdom; Department of Neurology, Mayo Clinic, Rochester, MN, USA; Department of Radiology, Mayo Clinic, Rochester, MN, USA; Department of Psychiatry Division of Neurosciences, University Medical Center, Utrecht, The Netherlands; Department of Psychiatry, Icahn School of Medicine, Mount Sinai, New York, USA; Department of Clinical Medicine, Department of Psychiatry and Turku Brain and Mind Center, FinnBrain Birth Cohort Study, University of Turku and Turku University Hospital, Turku, Finland; Centre for Population Health Research, Turku University Hospital and University of Turku, Turku, Finland; Institute of Development, Aging and Cancer, Tohoku University, Seiryocho, Aobaku, Sendai 980-8575, Japan; Queen’s University, Departments of Psychology and Psychiatry, Centre for Neuroscience Studies, Kingston, Ontario, Canada; Institute of Neuroscience and Physiology, University of Gothenburg, Gothenburg, Sweden; Department of Brain and Cognitive Sciences, Seoul National University College of Natural Sciences, Seoul, Republic of Korea; Department of Neuropsychiatry, Seoul National University Bundang Hospital, Seongnam, Republic of Korea; Department of Psychiatry, Seoul National University College of Medicine, Seoul, Republic of Korea; Department of Brain and Cognitive Science, Seoul National University College of Natural Sciences; Section on Developmental Neurogenomics, Human Genetics Branch, National Institute of Mental Health, Bethesda, MD, USA; Department of Brain & Cognitive Sciences, Seoul National University College of Natural Sciences, Seoul, Korea; Department of Medical Biophysics, University of Toronto, Toronto, ON, Canada; Mouse Imaging Centre, The Hospital for Sick Children, Toronto, ON, Canada; Wellcome Centre for Integrative Neuroimaging, FMRIB, Nuffield Department of Clinical Neuroscience, University of Oxford, Oxford, UK; Montreal Neurological Institute, McGill University, Montreal, Canada; The Clinical Hospital of Chengdu Brain Science Institute, University of Electronic Science and Technology of China, Chengdu 611731, China; Department of Psychiatry and Brain and Mind Research Institute, Weill Cornell Medicine; Laboratory for Autism and Neurodevelopmental Disorders, Center for Neuroscience and Cognitive Systems @UniTn, Istituto Italiano di Tecnologia, Rovereto, Italy; School of Biomedical Engineering & Brain and Mind Centre, The University of Sydney, Sydney, NSW, Australia; Department of Psychology, University of Texas, Austin, Texas 78712, USA; Department of Psychiatry and Neuropsychology, School of Mental Health and Neuroscience, EURON, Maastricht University Medical Centre, PO Box 616, 6200 MD, Maastricht, the Netherlands; Institute for Mental Health Care Eindhoven (GGzE), Eindhoven, the Netherlands; McConnell Brain Imaging Centre, Montreal Neurological Institute, McGill University, Montreal, QC H3A 2B4, Canada; Bordeaux University Hospital; Professor, Department of Psychosis Studies, Institute of Psychiatry, Psychology and Neuroscience, King’s College London, UK; Ludmer Centre for Neuroinformatics and Mental Health, Douglas Mental Health University Institute, McGill University, Montreal, Quebec, Canada; Singapore Institute for Clinical Sciences, Singapore; Department of Computer Science and Technology, University of Cambridge, Cambridge CB3 0FD, United Kingdom; Department of Psychiatry, University of Cambridge, Cambridge CB2 0SZ, United Kingdom; The Alan Turing Institute, London, NW1 2DB; Department of Psychology, School of Business, National College of Ireland, Dublin, Ireland; School of Psychology & Center for Neuroimaging and Cognitive Genomics, National University of Ireland Galway, Galway, Ireland; Department of Psychiatry, Trinity College Dublin, Dublin, Ireland; Department of Psychiatry, School of Medicine, Oregon Health and Science University, Portland, United States; Center for Sleep and Cognition, Yong Loo Lin School of Medicine, National University of Singapore, Singapore; Department of Pediatrics, Washington University in St. Louis, St. Louis, Missouri, United States; Alzheimer Center Amsterdam, Department of Neurology, Amsterdam Neuroscience, Vrije Universiteit Amsterdam, Amsterdam UMC, Amsterdam, The Netherlands; Lund University, Clinical Memory Research Unit, Lund, Sweden; Robarts Research Institute & The Brain and Mind Institute, University of Western Ontario,London,Ontario,Canada; Department of Psychiatry, Federal University of Sao Poalo (UNIFESP); National Institute of Developmental Psychiatry for Children and Adolescents (INPD), Brazil; Melbourne Neuropsychiatry Centre, Department of Psychiatry, The University of Melbourne and Melbourne Health, Carlton South, Victoria, Australia; Melbourne School of Engineering, The University of Melbourne, Parkville, Victoria, Australia; Florey Institute of Neuroscience and Mental Health, Parkville, VIC, Australia; Department of Psychiatry, Schulich School of Medicine and Dentistry, Western University, London, ON, Canada; Department of Psychiatry, Faculty of Medicine and Centre Hospitalier Universitaire Sainte-Justine, University of Montreal, Montreal, Quebec, Canada; Departments of Psychiatry and Psychology, University of Toronto, Toronto, ON, Canada; Departments of Physiology and Nutritional Sciences, University of Toronto, Toronto, Canada; Cuban Neuroscience Center, Havana, Cuba; Department of Psychiatry, Faculty of Medicine, McGill University, Montreal, Qc, H3A 1Y2, Canada; Douglas Mental Health University Institute, Montreal, Qc, H4H 1R3, Canada; Autism Center of Excellence, Department of Neurosciences, University of California, San Diego La Jolla, CA, USA; School of Psychology, Southwest University, Chongqing 400715, P.R. China; Department of Biomedical Engineering, The N.1 Institute for Health, National University of Singapore; Department of Clinical Neurosciences, University of Cambridge, Cambridge UK; Department of Neurology, Harvard Medical School; Department of Neurology, Boston Children’s Hospital, Boston, MA 02115; Instituto de Biomedicina de Sevilla (IBiS) HUVR/CSIC/Universidad de Sevilla, Dpto. de Fisiología Médica y Biofísica, Spain; Department of Psychology, Neuroscience Institute, University of Chicago; Department of Paediatrics and Wellcome-MRC Cambridge Stem Cell Institute, University of Cambridge, Hills Road, Cambridge, UK; Department of Psychiatry, Universidade Federal do Rio Grande do Sul (UFRGS); National Institute of Developmental Psychiatry (INPD); Otto Hahn Group Cognitive Neurogenetics, Max Planck Institute for Human Cognitive and Brain Sciences, Leipzig, Germany; Institute of Neuroscience and Medicine (INM-7: Brain and Behaviour), Research Centre Juelich, Juelich, Germany; Athinoula A. Martinos Center for Biomedical Imaging, Department of Radiology, Massachusetts General Hospital, Charlestown, MA 02129, USA; Centre for Population Neuroscience and Stratified Medicine (PONS), SGDP Centre, IoPPN, KCL, UK; PONS-Centre, Dept of Psychiatry and Psychotherapy, Campus Charite Mitte, Humboldt University, Berlin, Germany; PONS Centre, Institute for Science and Technology of Brain-inspired Intelligence (ISTBI), Fudan University, Shanghai, China; Wallenberg Centre for Molecular and Translational Medicine, University of Gothenburg, Gothenburg, Sweden; Department of Psychiatry and Neurochemistry, University of Gothenburg, Sweden; Dementia Research Centre, Queen’s Square Institute of Neurology, University College London, UK; Center for Biomedical Image Computing and Analytics, Department of Radiology, Perelman School of Medicine, University of Pennsylvania, Philadelphia, PA, USA; Departments of Neurology, Pediatrics, and Radiology, Washington University School of Medicine, St. Louis, United States; SA MRC Unit on Risk & Resilience in Mental Disorders, Dept of Psychiatry and Neuroscience Institute, University of Cape Town, Cape Town, South Africa; Division of Psychiatry, Centre for Clinical Brain Sciences, University of Edinburgh, UK; Department of Neuroscience, Institut Pasteur, Paris, France; Center for Research and Interdisciplinarity (CRI), Université Paris Descartes, Paris, France; Department of Psychology, University of Cambridge, Cambridge, UK; Wu Tsai Institute, Yale University, New Haven, CT, USA; Department of Clinical Medicine, Department of Psychiatry, FinnBrain Birth Cohort Study, University of Turku, Turku, Finland; Department of Clinical Medicine, University of Turku, Turku Finland; Turku Collegium for Science, Medicine and Technology, University of Turku, Turku, Finland; Univ. Bordeaux, Inserm, Bordeaux Population Health Research Center, U1219, CHU Bordeaux, F-33000 Bordeaux, France; Department of Anesthesia, Faculty of Medicine, McGill University, Montreal, Qc, H3A 1G1, Canada; Faculty of Dentistry, McGill University, Montreal, Qc, H3A 1G1, Canada; Alan Edwards Centre for Research on Pain (AECRP), McGill University, Montreal, Qc, H3A 1G1, Canada; Joint China-Cuba Lab,University of Electronic Science and Technology, Chengdu China/Cuban Center for Neuroscience, La Habana, Cuba; University of Electronic Science and Technology of China/Cuban Center for Neuroscience; Institute for Neuroscience and Medicine 7, Forschungszentrum Juelich; Max Planck Institute for Human Cognitive and Brain Sciences; Department of Psychiatry & Neurosychology, Maastricht University, Maastricht, The Netherlands; Department of Biostatistics, Vanderbilt University, Nashville, Tennessee, USA; Department of Biostatistics, Vanderbilt University Medical Center, Nashville, Tennessee, USA; Division of Newborn Medicine, Fetal Neonatal Neuroimaging and Developmental Science Center, Department of Pediatrics, Boston Children’s Hospital, Boston, MA 02115; McConnell Brain Imaging Center, Montreal Neurological Institute, McGill University, Montreal, Quebec, Canada; Clinic for Cognitive Neurology, University of Leipzig Medical Center, Leipzig, 04103, Germany; The Alan Turing Institute, London NW1 2DB, UK; Wellcome Centre for Human Neuroimaging, Institute of Neurology, University College London, WC1N 3AR; State Key Laboratory of Cognitive Neuroscience and Learning, Beijing Normal University, Beijing 100875, China; Developmental Population Neuroscience Research Center, IDG/McGovern Institute for Brain Research, Beijing Normal University, Beijing 100875, China; National Basic Science Data Center, Beijing 100190, China; Research Center for Lifespan Development of Brain and Mind, Institute of Psychology, Chinese Academy of Sciences, Beijing 100101, China; Division of Clinical Geriatrics, Center for Alzheimer Research, Department of Neurobiology, Care Sciences and Society, Karolinska Institutet, Stockholm, Sweden; Faculty of Medicine, CRC 1052 ‘Obesity Mechanisms’, University of Leipzig, Leipzig, 04103, Germany; Department of Electrical and Computer Engineering, National University of Singapore, Singapore; Centre for Sleep & Cognition and Centre for Translational MR Research, Yong Loo Lin School of Medicine, National University of Singapore, Singapore; N.1 Institute for Health & Institute for Digital Medicine, National University of Singapore, Singapore; Integrative Sciences and Engineering Programme (ISEP), National University of Singapore, Singapore; Fetal Neonatal Neuroimaging and Developmental Science Center, Division of Newborn Medicine, Boston Children’s Hospital, Harvard Medical School, Boston, MA 02115, USA; Melbourne Neuropsychiatry Centre, University of Melbourne, Melbourne, Australia; Department of Biomedical Engineering, University of Melbourne, Melbourne, Australia; SAMRC Unit on Child & Adolescent Health, University of Cape Town, South Africa; Center for Translational Magnetic Resonance Research, Yong Loo Lin School of Medicine, National University of Singapore, Singapore; Wellcome Trust-MRC Institute of Metabolic Science, University of Cambridge, Cambridge, CB2 0SZ; Cambridgeshire and Peterborough Foundation Trust, Cambridge, CB21 5EF; National Institute of Mental Health (NIMH), National Institutes of Health (NIH), Bethesda, Maryland, USA; Department of Psychiatry, Escola Paulista de Medicina, São Paulo, Brazil; Key Laboratory of Brain and Education, School of Education Science, Nanning Normal University, Nanning 530001, China

## Abstract

Over the past few decades, neuroimaging has become a ubiquitous tool in basic research and clinical studies of the human brain. However, no reference standards currently exist to quantify individual differences in neuroimaging metrics over time, in contrast to growth charts for anthropometric traits such as height and weight^1^. Here, we built an interactive resource to benchmark brain morphology, www.brainchart.io, derived from any current or future sample of magnetic resonance imaging (MRI) data. With the goal of basing these reference charts on the largest and most inclusive dataset available, we aggregated 123,984 MRI scans from 101,457 participants aged from 115 days post-conception through 100 postnatal years, across more than 100 primary research studies. Cerebrum tissue volumes and other global or regional MRI metrics were quantified by centile scores, relative to non-linear trajectories^2^ of brain structural changes, and rates of change, over the lifespan. Brain charts identified previously unreported neurodevelopmental milestones^3^; showed high stability of individual centile scores over longitudinal assessments; and demonstrated robustness to technical and methodological differences between primary studies. Centile scores showed increased heritability compared to non-centiled MRI phenotypes, and provided a standardised measure of atypical brain structure that revealed patterns of neuroanatomical variation across neurological and psychiatric disorders. In sum, brain charts are an essential first step towards robust quantification of individual deviations from normative trajectories in multiple, commonly-used neuroimaging phenotypes. Our collaborative study proves the principle that brain charts are achievable on a global scale over the entire lifespan, and applicable to analysis of diverse developmental and clinical effects on human brain structure. Furthermore, we provide open resources to support future advances towards adoption of brain charts as standards for quantitative benchmarking of typical or atypical brain MRI scans.

## Main

The simple framework of growth charts to quantify age-related change was first published in the late 18th century^1^ and remains a cornerstone of paediatric healthcare – an enduring example of the utility of standardised norms to benchmark individual trajectories of development. However, growth charts are currently available only for a small set of anthropometric variables, e.g., height, weight and head circumference, and only for the first decade of life. There are no analogous charts available for quantification of age-related changes in the human brain, although it is known to go through a prolonged and complex maturational program from pregnancy to the third decade^4^, followed by progressive senescence from the sixth decade^5^, approximately. The lack of tools for standardised assessment of brain development and aging is particularly relevant to research studies of psychiatric disorders, which are increasingly recognised as a consequence of atypical brain development^6^, and neurodegenerative diseases that cause pathological brain changes in the context of normative senescence^7^. Preterm birth and neurogenetic disorders are also associated with marked abnormalities of brain structure^8,9^ that persist into adult life^9,10^ and are associated with learning disabilities and mental health disorders. Mental illness and dementia collectively represent the single biggest global health burden^11^, highlighting the urgent need for normative brain charts as an anchorpoint for standardised quantification of brain structure over the lifespan^12^.

Such standards for human brain measurement have not yet materialised from decades of neuroimaging research, likely due to the challenges of integrating magnetic resonance imaging (MRI) data across multiple, methodologically diverse studies targeting distinct developmental epochs and clinical conditions^13^. For example, the perinatal period is rarely incorporated in analysis of age-related brain changes, despite evidence that early biophysical and molecular processes powerfully influence life-long neurodevelopmental trajectories^14,15^ and vulnerability to psychiatric disorders^3^. Primary case-control studies are usually focused on a single disorder despite evidence of trans-diagnostically shared risk factors and pathogenic mechanisms, especially in psychiatry^16,17^. Harmonization of MRI data across primary studies to address these and other deficiencies in the extant literature is challenged by methodological and technical heterogeneity. Compared to relatively simple anthropometric measurements, like height or weight, brain morphometrics are known to be highly sensitive to variation in scanner platforms and sequences, data quality control, pre-processing and statistical analysis^18^, thus severely limiting the generalisability of trajectories estimated from any individual study^19^. Collaborative initiatives spurring collection of large-scale datasets^20,21^, recent advances in neuroimaging data processing^22,23^, and proven statistical frameworks for modelling biological growth curves^2,24,25^ provide the building blocks for a more comprehensive and generalisable approach to age-normed quantification of MRI phenotypes over the entire lifespan (see **SI 1** for details and consideration of prior work focused on the related but distinct objective of inferring brain age from MRI data). Here, we demonstrate that these convergent advances now enable the generation of brain charts that i) robustly define normative processes of sex-stratified, age-related change in multiple MRI-derived phenotypes; ii) identify previously unreported brain growth milestones; iii) increase sensitivity to detect genetic and early life environmental effects on brain structure; and iv) provide standardised effect sizes to quantify neuroanatomical atypicality of brain scans collected across multiple clinical disorders. We do not claim to have yet reached the ultimate goal of quantitatively precise diagnosis of MRI scans from individual patients in clinical practice. However, the present work proves the principle that building normative charts to benchmark individual differences in brain structure is already achievable at global scale and over the entire life-course; and provides a suite of open science resources for the neuroimaging research community to accelerate further progress in the direction of standardised quantitative assessment of MRI data.

### Mapping normative brain growth

We created brain charts for the human lifespan using generalised additive models for location, scale and shape (GAMLSS)^2,24^, a robust and flexible framework for modelling non-linear growth trajectories recommended by the World Health Organization^24^. GAMLSS and related statistical frameworks have previously been applied to developmental modelling of brain structural and functional MRI phenotypes in open datasets^19,26–31^. Our approach to GAMLSS modelling leveraged the greater scale of data available to optimise model selection empirically, to estimate non-linear age-related trends (in median and variance) across the lifespan stratified by sex over the entire lifespan, and to account for site- or study-specific “batch effects” on MRI phenotypes in terms of multiple random effect parameters. Specifically, GAMLSS models were fitted to structural MRI data from control subjects for the four main tissue volumes of the cerebrum (total cortical grey matter volume [GMV] and total white matter volume [WMV], total subcortical grey matter volume [sGMV], and total ventricular cerebrospinal fluid volume [Ventricles or CSF]). See **Online Methods, Supplementary Table** [**ST**] **1.1-1.8** for details on acquisition, processing and demographics of the dataset. See **Supplementary Information** [**SI**] for details regarding GAMLSS model specification and estimation (**SI1**); image quality control, which utilized a combination of expert visual curation and automated metrics of image quality (**SI2**); model stability and robustness (**SI3-4**); phenotypic validation against non-imaging metrics (**SI3** and **SI5.2**); inter-study harmonisation (**SI5**); and assessment of cohort effects (**SI6**). See **SI19** for details on all primary studies contributing to the reference dataset, including multiple publicly available open MRI datasets^32–42^.

Lifespan curves (**Fig.1**; **ST2.1**) showed an initial strong increase in GMV from mid-gestation onwards, peaking at 5.9_CI-Bootstrap:5.8-6.1_ years, followed by a near-linear decrease. This peak was observed 2-3 years later than prior reports relying on smaller, more age-restricted samples^43,44^. WMV also increased rapidly from mid-gestation through early childhood peaking at 28.7_CI-Bootstrap:28.1-29.2_ years, with subsequent accelerated decline in WMV after 50 years. Subcortical GMV showed an intermediate growth pattern compared to GMV and WMV, peaking in adolescence at 14.4_CI-Bootstrap:14.0-14.7_ years. Both the WMV and sGMV peaks are consistent with prior neuroimaging and postmortem reports^45,46^. In contrast, CSF showed an increase until age 2, followed by a plateau until age 30, and then a slow linear increase that exponentiated in the sixth decade of life. Age-related variance (**Fig.1D**), explicitly estimated by GAMLSS, formally quantifies developmental changes in between-subject variability. There was an early developmental increase in GMV variability that peaked at 4 years, whereas subcortical volume variability peaked in late adolescence. WMV variability peaked during the fourth decade of life, and CSF was maximally variable at the end of the human lifespan.

**Fig. 1.**
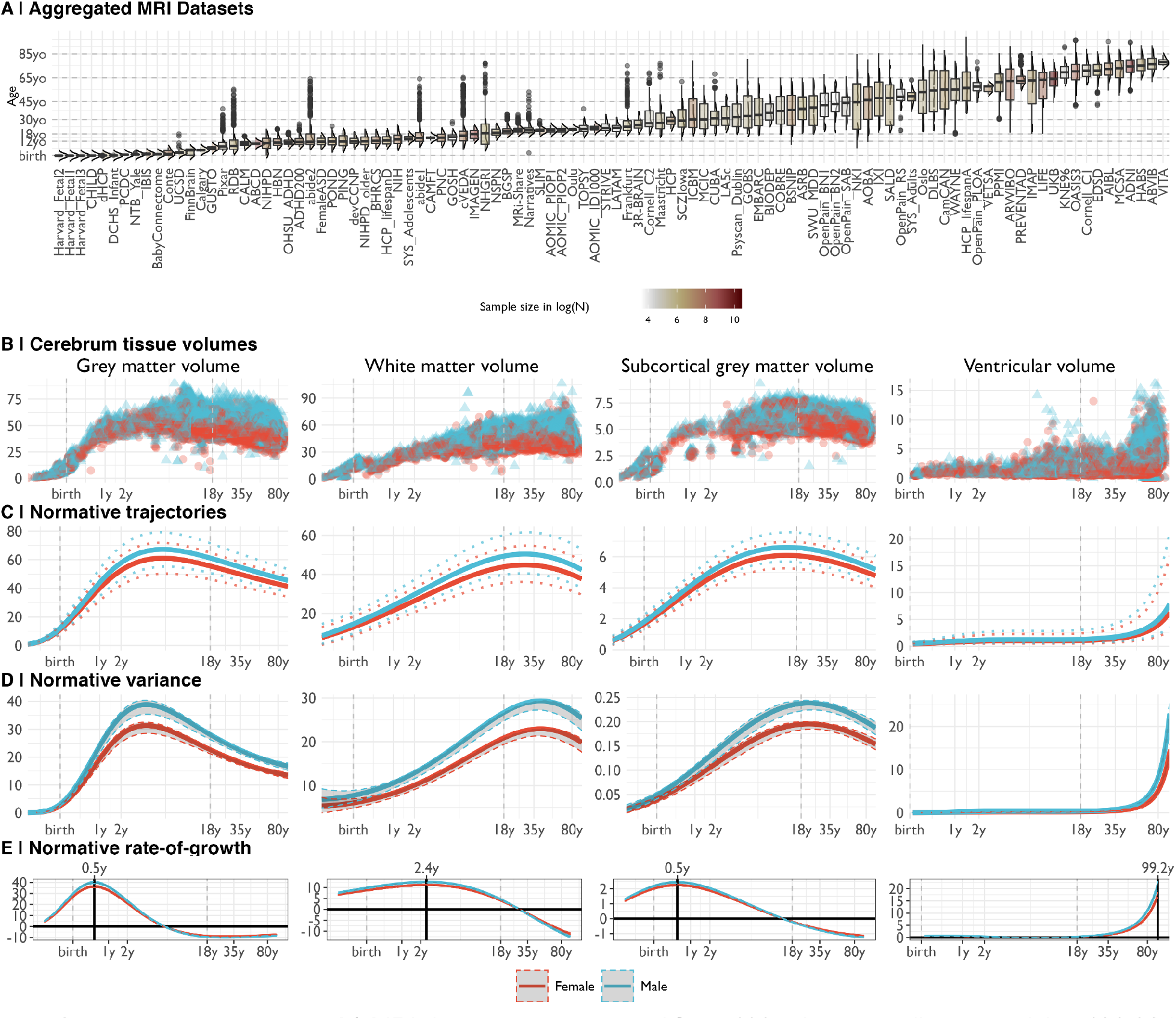
Human brain charts. A | MRI data were aggregated from 100 primary studies comprising 123,984 scans that collectively spanned the age range from mid-gestation to 100 postnatal years. Box-violin plots show the age distribution (log-scaled) for each study coloured by its relative sample-size (log-scaled). B | Non-centiled, “raw” bilateral cerebrum tissue volumes (from left to right: grey matter, white matter, subcortical grey matter and ventricles) are plotted for each cross-sectional control scan, point-coloured by sex, as a function of age (log-scaled). C | Normative brain trajectories were estimated by generalised additive modelling for location, scale and shape (GAMLSS), accounting for site- and study-specific batch effects, and stratified by sex (female/male curves coloured red/blue). All four cerebrum tissue volumes demonstrated distinct, non-linear trajectories of their medians (with 2.5% and 97.5% centiles denoted as dotted lines) as a function of age over the lifespan. Demographics for each cross-sectional sample of healthy controls included in the reference dataset for normative GAMLSS modelling of each MRI phenotype are detailed in **ST1.2-1.8**. D | Trajectories of median between-subject variability and 95% confidence intervals for four cerebrum issue volumes were estimated by sex-stratified bootstrapping (see **SI3** for details). E | Rates of volumetric change across the lifespan for each tissue volume, stratified by sex, were estimated by the first derivatives of the median volumetric trajectories. For solid (parenchymal) tissue volumes, the horizontal line (y=0) indicates when the volume of each tissue stops growing and starts shrinking; the solid vertical line indicates the age of maximum growth of each tissue. See **ST2.1** for all neurodevelopmental milestones and their confidence intervals. Note that y-axes in panels B-E are scaled in units of 10,000 mm^3^ (10ml).

### Extended neuroimaging phenotypes

To extend the scope of brain charts beyond the four cerebrum tissue volumes, we generalised the same GAMLSS modelling approach to estimate normative trajectories for additional MRI phenotypes including other geometric properties at a similar scale (mean cortical thickness and total surface area) and regional volume at each of 34 cortical areas^47^ (**Fig.2, SI7-9, ST1-2**). We found, as expected, that total surface area closely tracked the development of total cerebrum volume (TCV) across the lifespan (**Fig.2A**), with both metrics peaking at ∼11-12 years (SA 10.97_CI-Bootstrap:10.42-11.51_; TCV 12.5_CI-Bootstrap:12.14-12.89_). In contrast, cortical thickness peaked distinctively early at 1.7_CI-Bootstrap:1.3-2.1_ years, which reconciles prior observations that cortical thickness increases during the perinatal period^48^ and declines during later development^49^ (**SI7**).

**Fig. 2.**
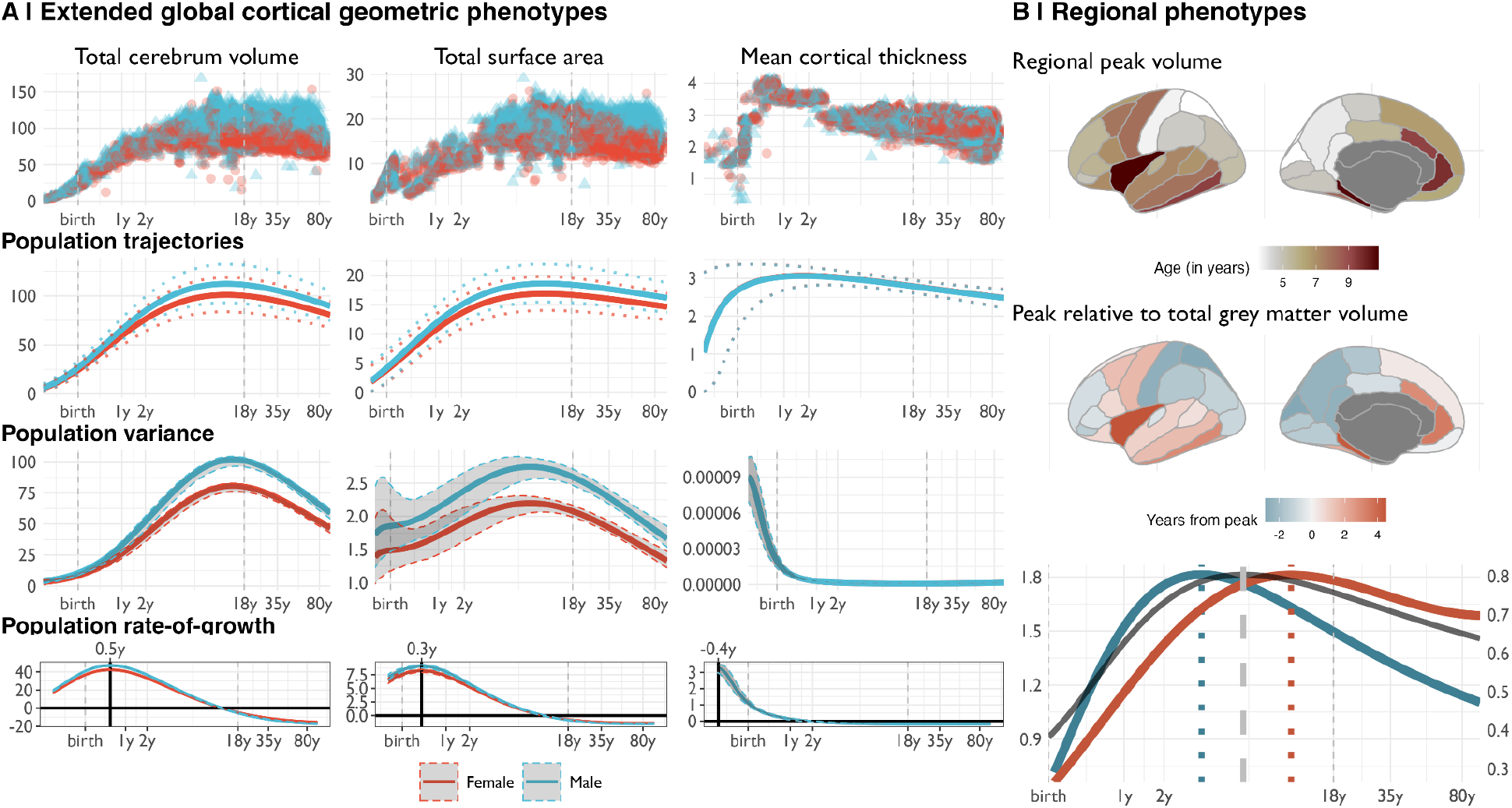
Extended global and regional cortical geometric phenotypes. A | Trajectories for total cerebrum volume (TCV; left column), total surface area (SA; middle column), and mean cortical thickness (CT; right column). For each global cortical geometric MRI phenotype, the following sex-stratified results are shown as a function of age over the lifespan, from top to bottom rows: raw, non-centiled data; population trajectories of the median (with 2.5% and 97.5% centiles; dotted lines); between-subject variance (and 95% confidence intervals); and rate-of-growth (the first derivatives of the median trajectory and 95% confidence intervals). All trajectories are plotted as a function of log-scaled age (x-axis) and y-axes are scaled in units of the corresponding MRI metrics (10,000 mm^3^ for TCV, 10,000 mm^2^ for SA and mm for CT). B | Regional variability of cortical volume trajectories for 34 bilateral brain regions, as defined by the Desikan-Killiany parcellation^47^, averaged across sex (see also **SI7-8** for details). Since models were generated from bilateral averages of each cortical region, the cortical maps are plotted on the left hemisphere purely for visualisation purposes. In the top panel, a cortical map of age at peak regional volume (range 2-10 years); in the middle panel, a cortical map of age at peak regional volume relative to age at peak GMV (5.9 years), highlighting regions that peak earlier (blue) or later (red) than GMV; in the bottom panel, illustrative trajectories for the earliest peaking region (superior parietal lobe, blue line) and the latest peaking region (insula, red line), showing the range of regional variability relative to the GMV trajectory (grey line). Regional volume peaks are denoted as dotted vertical lines either side of the global peak, denoted as a dashed vertical line, in the bottom panel. The left hand y-axis on the bottom panel refers to the earliest peak (blue line), the right hand y-axis refers to the latest peak (red line), and both are in units of 10,000 mm^3^ (10ml).

We also found evidence for regional variability in volumetric neurodevelopmental trajectories. Compared to peak GMV at 5.9 years, the age of peak regional grey matter volume varied considerably – from approximately 2 to 10 years – across 34 cortical areas. Primary sensory regions reached peak volume earliest, and fronto-temporal association cortical areas peaked later (**Fig.2B; SI8**). In general, earlier maturing ventral-caudal regions also showed faster post-peak declines in cortical volume, and later maturing dorsal-rostral regions showed slower post-peak declines (**Fig.2B; SI8.2**). Notably, this spatial pattern recapitulated a gradient from sensory-to-association cortex that has been previously associated with multiple aspects of brain structure and function^50^.

### Developmental milestones

Neuroimaging milestones are defined by inflection points of the tissue-specific volumetric trajectories (**Fig.3; Online Methods**). Among the total tissue volumes, only GMV peaked before the typical age at onset of puberty^51^, with sGMV peaking mid-puberty and WMV peaking in young adulthood (**Fig.3**). The rate-of-growth (velocity) peaked for GMV (5.08_CI-Bootstrap:4.85-5.22_ months), sGMV (5.65_CI-Bootstrap:5,75-5.83_ months) and WMV (2.4_CI-Bootstrap:2.2-2.6_ years) in infancy and early childhood. TCV velocity peaked between the maximum velocity for GMV and WMV at ∼7 months. Two major milestones of TCV and sGMV (peak velocity and size; **Fig.3**) coincided with the early neonatal and adolescent peaks of height and weight velocity^52,53^. The velocity of mean cortical thickness peaked even earlier, in the prenatal period at −0.38_CI-Bootstrap:-0.4 to −0.34_ years (relative to birth), corresponding approximately to mid-gestation. This early peak in cortical thickness velocity has not been reported previously, in part due to obstacles in acquiring adequate and consistent signal from typical MRI sequences in the perinatal period^54^. Similarly, normative trajectories revealed an early period of GMV:WMV differentiation, beginning in the first month after birth with the switch from WMV to GMV as the proportionally dominant tissue compartment, and ending when the absolute difference of GMV and WMV peaked around 3 years (**SI9**). This epoch of GMV:WMV differentiation, which may reflect underlying changes in myelination and synaptic proliferation^4,55–58^, has not been demarcated by prior studies^45,59^. It was likely identified in this study due to the substantial amount of early developmental MRI data available for analysis in the aggregated dataset (in total across all primary studies, N=2,571 and N=1,484 participants aged <2 years were available for analysis of cerebrum tissue volumes and extended global MRI phenotypes, respectively). The period of GMV:WMV differentiation encompasses dynamic changes in brain metabolites^60^ (0-3 months), resting metabolic rate (RMR; minimum=7 months, maximum=4.2 years)^61^, the typical period of acquisition of motor capabilities and other early paediatric milestones^62^, and the most rapid change in TCV (**Fig.3**).

**Fig. 3.**
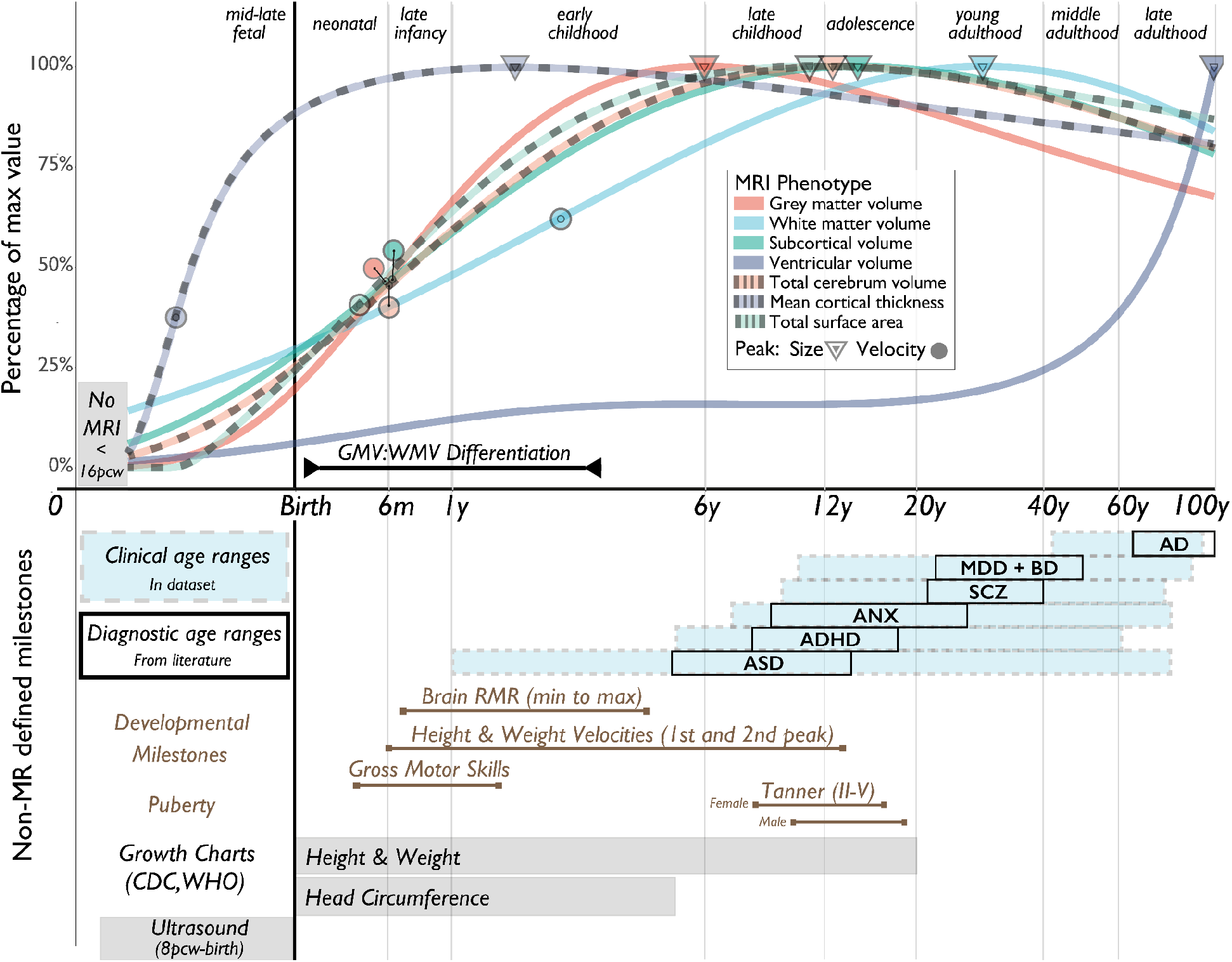
Neurodevelopmental milestones. Top panel: A graphical summary of the normative trajectories of the median (or 50th centile) for each global MRI phenotype, and key developmental milestones, as a function of age (log-scaled). Circles depict the peak rate-of-growth milestones for each phenotype (defined by the maxima of the first derivatives of the median trajectories; see **Fig.1E**). Triangles depict the peak volume of each phenotype (defined by the maxima of the median trajectories); the definition of GMV:WMV differentiation is detailed in **SI9.1**. Bottom panel: A graphical summary of additional MRI and non-MRI developmental stages and milestones. From top to bottom: blue shaded boxes denote the age-range of incidence for each of the major clinical disorders represented in the MRI dataset; black boxes denote the age at which these conditions are generally diagnosed as derived from literature^63^ (**Online Methods**); brown lines represent the normative intervals for developmental milestones derived from non-MRI data, based on previous literature and averaged across males and females (**Online Methods**); grey bars depict age ranges for existing (WHO and Centres for Disease Control and Prevention [CDC]) growth charts of anthropometric and ultrasonographic variables. Across both panels, light grey vertical lines delimit lifespan epochs (labelled above the top panel) previously defined by neurobiological criteria^64^. Abbreviations: resting metabolic rate (RMR), Alzheimer’s disease (AD), attention deficit hyperactivity disorder (ADHD), anxiety or phobic disorders (ANX), autism spectrum disorder (ASD, including high-risk individuals with confirmed diagnosis at a later age), major depressive disorder (MDD), bipolar disorder (BD), and schizophrenia (SCZ).

### Individualised centile scores in clinical samples

We computed individualised centile scores that benchmarked each individual scan in the context of normative age-related trends (**SI1-6**). This approach is conceptually similar to quantile rank mapping, as previously reported^26,28,29^, where the (a)typicality of each phenotype in each scan is quantified by its score on the distribution of phenotypic parameters in the normative or reference sample of scans, with more atypical phenotypes having more extreme centile (or quantile) scores. The clinical diversity of the aggregated dataset allowed us to comprehensively investigate case-control differences in individually-specific centile scores across multiple conditions. Relative to the control group (CN), there were highly significant differences in centile scores across large (N>500) diagnostic groups of multiple disorders (**Fig.4A; SI10)**, with effect-sizes ranging from medium (Cohen’s *d* ranging from 0.2 to 0.8) to large (Cohen’s *d* > 0.8) (see **ST3-4** for all false discovery rate (FDR)-corrected *P*-values and effect-sizes). The pattern of these group differences in centile scores varied across tissue types and disorders. Clinical differences in cortical thickness and surface area generally followed the same trend as volume differences (**SI10**). Alzheimer’s disease (AD) showed the greatest overall difference, with a maximum difference localised to gray matter in biologically female patients (median centile score = 14%, 36% points difference from CN median, corresponding to Cohen’s *d*=0.88; **Fig.4A**). In addition, we generated a cumulative deviation metric, the centile Mahalanobis distance (CMD), to summarise a comparative assessment of brain morphology across all global MRI phenotypes relative to the CN group (**Fig.4B**; **SI1.6**). Notably, schizophrenia (SCZ) ranked third overall behind AD and mild cognitive impairment (MCI), based on CMD (**Fig.4C**). Assessment across diagnostic groups, based on profiles of the multiple centile scores for each MRI phenotype and for CMD, highlighted shared and distinct patterns across clinical conditions (**SI10-11**). However, when examining cross-disorder similarity of multivariate centile scores, hierarchical clustering yielded three clusters broadly comprising neurodegenerative, mood/anxiety, and neurodevelopmental disorders (**SI11**). Overall, these analyses highlight some complementary use-cases for examining both absolute and relative differences in centile scores within and across conventional diagnostic categories.

**Fig. 4.**
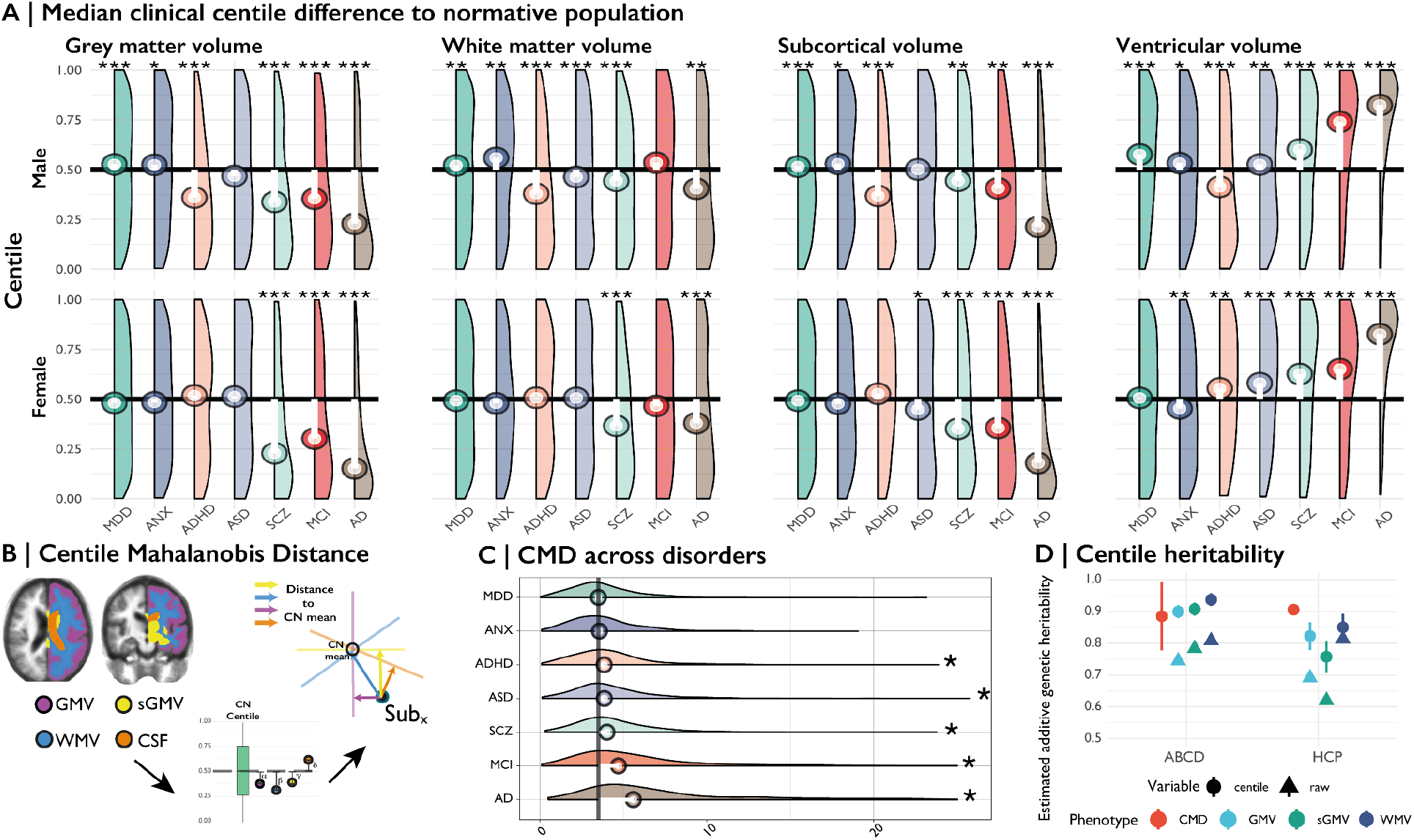
Case-control differences and heritability of centile scores. A | Centile score distributions for each diagnostic category of clinical cases relative to the control group median (depicted as a horizontal black line). The median deviation of centile scores in each diagnostic category is overlaid as a lollipop plot (white line with circle corresponding to the median centile score for each group of cases). Pairwise tests for significance were based on Monte-Carlo resampling (10,000 permutations) and P-values were adjusted for multiple comparisons using the Benjamini-Hochberg (FDR) correction across all possible case-control differences. Only significant deviations from the control group median (with corrected P<0.001) are highlighted with an asterisk. For a complete overview of all pairwise comparisons, see **SI10** and **ST3**. Groups are ordered by their multivariate distance from the control group (see panel C and **SI10.3**). B | The centile Mahalanobis distance (CMD) is a summary metric of multivariate deviation that quantifies the aggregate atypicality of an individual scan in terms of all global MRI phenotypes. The schematic shows segmentation of four cerebrum tissue volumes, followed by estimation of univariate centile scores, leading to the orthogonal projection of a single participant’s scan (Sub_x_) onto the four principal components of the control group (CN; coloured axes and arrows): the CMD for Sub_x_ is then the sum of its distances from the CN group mean on all 4 dimensions of the multivariate space. C | Probability density plots of CMD across disorders. Vertical black line depicts the median CMD of the control group. Asterisks indicate an FDR-corrected significant difference from the CN group (P<0.001). D | Heritability of raw volumetric phenotypes and their centile scores across two twin studies (ABCD and HCP; see **SI 19**). All dots have error bars for the standard error, but in some cases these are too narrow to be observed. Abbreviations: control (CN), Alzheimer’s disease (AD), attention deficit hyperactivity disorder (ADHD), anxiety or phobia (ANX), autism spectrum disorder (ASD, including high-risk individuals with confirmed diagnosis at a later age), mild cognitive impairment (MCI), major depressive disorder (MDD), schizophrenia (SCZ); grey matter volume (GMV), subcortical grey matter volume (sGMV), white matter volume (WMV), centile Mahalanobis distance (CMD). Asterisks indicate level of FDR-corrected significance: P<0.05, P<0.01 or P<0.001 for *, ** and *** respectively.

Between-subject variation in centile scores also showed strong associations with development, early-life events, and shared genetic architecture. Across all major epochs of the lifespan, CMD was consistently greater in cases relative to controls, irrespective of diagnostic category, with the largest difference found in adolescence and late adulthood across epochs^64^ (**SI10.3**). Adolescence also showed the greatest overlap between diagnostic categories in our dataset, and is well-recognised as a period of increased incidence of mental health disorders (**Fig.4; SI10-11**). Across 5 primary studies covering the lifespan, average centile scores were related to two metrics of premature birth (gestational age at birth: *t*=13.164, *P*<2e-16; birth weight: *t*=36.395, *P*<2e-16; SI12). Centile scores also showed increased twin-based heritability in two independent studies (total N=913 twin-pairs) compared to non-centiled phenotypes (average increase of 11.8% points in h^2^ across phenotypes; **Fig.4D, SI13**). In summary, centile normalisation of brain metrics reproducibly detected case-control differences and genetic effects on brain structure, as well as long-term sequelae of adverse birth outcomes even in the adult brain^10^.

### Longitudinal centile changes

Due to the relative paucity of longitudinal imaging data (∼10% of the reference dataset), normative models were estimated from cross-sectional data collected at a single time point. However, the generalisability of cross-sectional models to longitudinal assessment is important for future research. Within-subject variability of centile scores derived from longitudinally repeated scans, measured with the interquartile range (IQR; **SI1.7**), was low across both clinical and control groups (all median IQR < 0.05 centile points), indicating that centile scoring of brain structure was generally stable over time, although there was also some evidence of between-study and cross-disorder differences in within-subject variability (**SI14**). Notably, individuals who changed diagnostic categories, e.g., progressed from MCI to AD, over the course of repeated scanning showed small but significant increases in within-subject variability of centile scores (**SI14**; **ST5-6**). Within-subject variability was also slightly higher in younger samples (**SI14**), which could reflect increased noise due to the technical or data quality difficulties associated with scanning younger individuals, but is also consistent with the evidence of increased variability in earlier development observed across other anthropometric traits^65^.

### Out-of-sample centile scoring of “new” MRI data

A key challenge for brain charts is the accurate centile scoring of out-of-sample MRI data, not represented in the reference dataset used to estimate normative trajectories. We therefore carefully evaluated the reliability and validity of brain charts for centile scoring of such “new” scans. For each new MRI study, we used maximum likelihood to estimate study-specific statistical offsets from the age-appropriate epoch of the normative trajectory; then we estimated centile scores for each individual in the new study benchmarked against the offset trajectory (**Fig.5; SI1.8**). Extensive jack-knife and leave-one-study-out (LOSO) analyses indicated that a study size of N>100 scans was sufficient for stable and unbiased estimation of out-of-sample centile scores (**SI4**). This study size limit is in line with the size of many contemporary brain MRI research studies. However, these results do not immediately support the use of brain charts to generate centile scores from smaller scale research studies, or from an individual patient’s scan in clinical practice – this remains a goal for future work. Out-of-sample centile scores proved highly reliable in multiple test-retest datasets and were robust to variations in image processing pipelines (**SI4**).

**Fig. 5.**
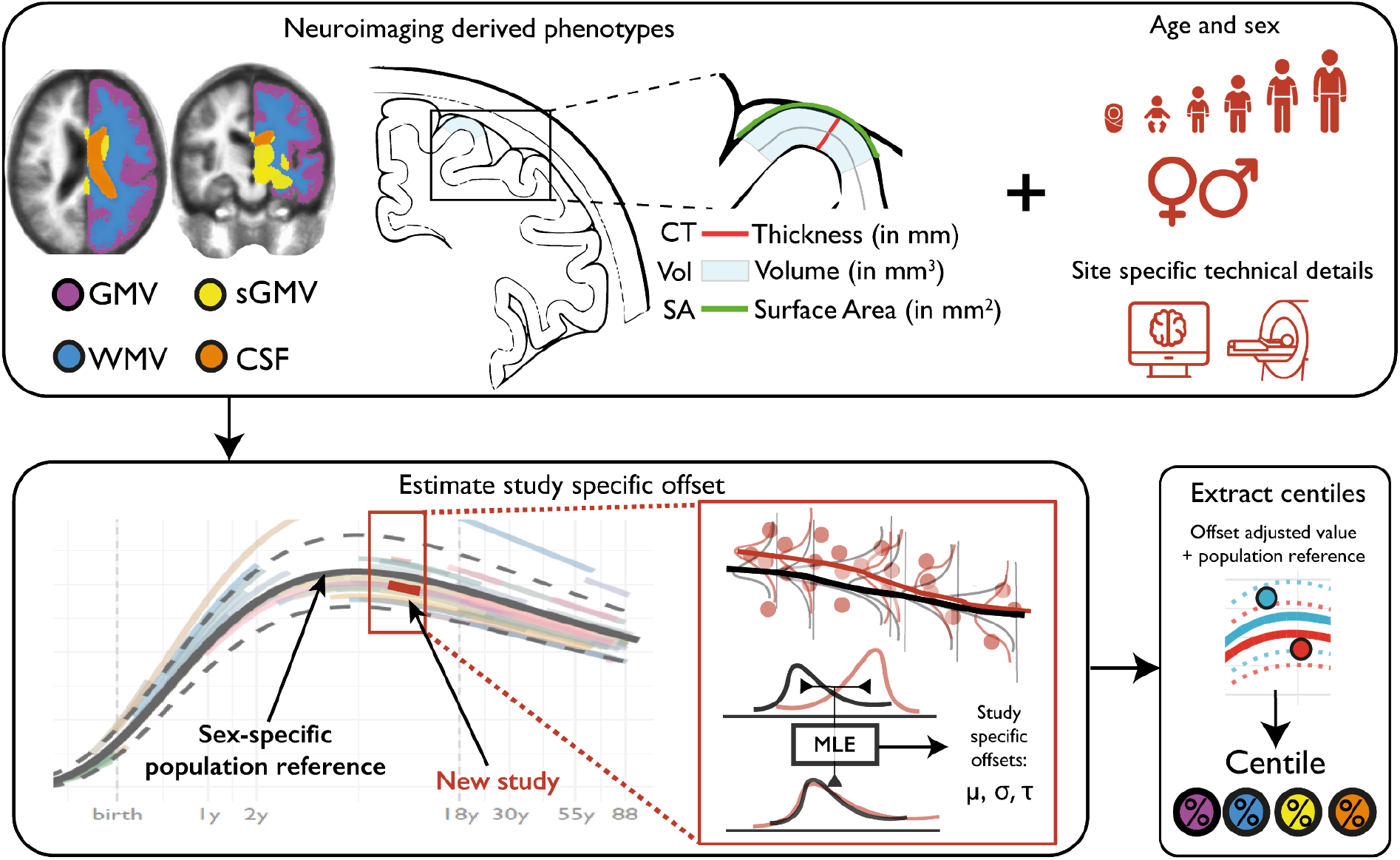
Schematic overview of brain charts, highlighting methods for out-of-sample centile scoring. Top panel: Brain phenotypes were measured in a reference dataset of MRI scans. GAMLSS modelling was used to estimate the relationship between (global) MRI phenotypes and age, stratified by sex, and controlling for technical and other sources of variation between scanning sites and primary studies. Bottom panel: The normative trajectory of the median and confidence interval for each phenotype was plotted as a population reference curve. Out-of-sample data from a new MRI study were aligned to the corresponding epoch of the normative trajectory, using maximum likelihood to estimate the study specific offsets (random effects) for three moments of the underlying statistical distributions: mean (μ), variance (σ), and skewness (**ν**) in an age- and sex-specific manner. Centile scores could then be estimated for each scan in the new study, on the same scale as the reference population curve, while accounting for study-specific “batch effects” on technical or other sources of variation (see **SI1.8** for details).

## Discussion

We have aggregated the largest neuroimaging dataset to date to modernise the concept of growth charts for mapping typical and atypical human brain development and ageing. The ∼100-year age range enabled the delineation of milestones and critical periods in maturation of the human brain, revealing an early growth epoch across its constituent tissue classes -- starting before 17 post-conception weeks, when the brain is at ∼10% of its maximum size and ending at ∼80% maximum size by age 3. Individual centile scores benchmarked by normative neurodevelopmental trajectories were significantly associated with neuropsychiatric disorders as well as with dimensional phenotypes (**SI5.2** and **SI12**). Furthermore, imaging-genetics studies^66^ may benefit from the increased heritability of centile scores compared to raw volumetric data (**SI13**). Perhaps most importantly, GAMLSS modelling enabled harmonisation across technically diverse studies (**SI5**), and thus unlocked the potential value of combining primary MRI studies at scale to generate normative, sex-stratified brain growth charts, and individual centile scores of (a)typicality.

The analogy to paediatric growth charts is not meant to imply that brain charts are immediately suitable for benchmarking or quantitative diagnosis of individual patients in clinical practice. Even for traditional anthropometric growth charts (height, weight and BMI) there are still significant caveats and nuances concerning their diagnostic interpretation in individual children^67^; and, likewise, it is expected that considerable further research will be required to validate the clinical diagnostic utility of brain charts. However, the current results bode well for future progress towards digital diagnosis of atypical brain structure and development^68^. By providing an age- and sex-normalised metric, centile scores enable trans-diagnostic comparisons between disorders that emerge at different stages of the lifespan (**SI10-11**). The generally high stability of centile scores across longitudinal measurements also enabled assessment of documented changes in diagnosis such as the transition from MCI to AD (**SI14**), which provides one example of how centile scoring could be clinically useful in quantitatively predicting or diagnosing progressive neurodegenerative disorders in the future. Our provision of appropriate normative growth charts and on-line tools also creates an immediate opportunity to quantify atypical brain structure in clinical research samples, to leverage available legacy neuroimaging datasets, and to enhance ongoing studies.

Several important caveats are worth highlighting. The use of brain charts does not circumvent the fundamental requirement for quality control of MRI data. We have shown that GAMLSS modelling of global structural MRI phenotypes is in fact remarkably robust to inclusion of poor quality scans (**SI2**), but it should not be assumed that this level of robustness will apply to future brain charts of regional MRI or functional MRI phenotypes; therefore the importance of quality control remains paramount. It will also be important in future to represent ethnic diversity appropriately in normative brain charts^69,70^. Even this large MRI dataset was heavily biased towards European and North American populations, as is also common in anthropometric growth charts and existing genetic datasets. Further increasing ethnic and demographic diversity in MRI research will enable more population-representative normative trajectories^69,70^ that can be expected to improve the accuracy and strengthen the interpretation of centile scores in relation to appropriate norms^26^. The available reference data were also not equally representative of all ages, e.g., foetal, neonatal and mid-adulthood (30-40y) epochs were under-represented (**SI17-19**). While our statistical modelling approach was designed to mitigate study- or site-specific effects on centile scores, it cannot entirely correct for limitations of primary study design, such as ascertainment bias or variability in diagnostic criteria. Our decision to stratify the lifespan models by sex followed the analogous logic of sex-stratified anthropometric growth charts. Males have larger brain tissue volumes than females in absolute terms (**SI16**), but this is not indicative of any difference in clinical or cognitive outcomes. Future work would also benefit from more detailed and dimensional self-report variables relating to sex and gender^71^.

We have focused primarily on global brain phenotypes, which were measurable in the largest possible aggregated sample over the widest age range, with the fewest methodological, theoretical and data sharing constraints. However, we have also provided proof-of-concept brain charts for regional grey matter volumetrics, demonstrating plausible heterochronicity of cortical patterning, and illustrating the potential generalisability of this approach to a diverse range of fine-grained MRI phenotypes (**Fig.2**; **SI8**). As ongoing and future efforts provide increasing amounts of high-quality MRI data, we predict an iterative process of improved brain charts for an increasing number of multimodal^72^ neuroimaging phenotypes. Such diversification will require the development, implementation, and standardisation of additional data quality control procedures^27^ to underpin robust brain chart modelling. To facilitate further research using our reference charts, we have provided interactive tools to explore these statistical models and to derive normalised centile scores for new datasets across the lifespan at www.brainchart.io.

## Acknowledgements

RAIB was supported by a British Academy Postdoctoral fellowship and by the Autism Research Trust. JS was supported by NIMH T32MH019112-29 and K08MH120564. SRW was funded by UKRI Medical Research Council MC_UU_00002/2 and was supported by the NIHR Cambridge Biomedical Research Centre (BRC-1215-20014). ETB was supported by an NIHR Senior Investigator award and the Wellcome Trust collaborative award for the Neuroscience in Psychiatry Network. AFA-B was supported by NIMH K08MH120564. Data were curated and analysed using a computational facility funded by an MRC research infrastructure award (<u>MR/M009041/1</u>) to the School of Clinical Medicine, University of Cambridge and supported by the mental health theme of the NIHR Cambridge Biomedical Research Centre. The views expressed are those of the authors and not necessarily those of the NIH, NHS, the NIHR or the Department of Health and Social Care. See supplementary information (**SI 23**) for a comprehensive list of all contributing authors. We would particularly like to acknowledge the invaluable contribution to this effort made by several openly-shared MRI datasets, specifically:s OpenNeuro (https://openneuro.org/), the Healthy Brain Network (https://healthybrainnetwork.org/), UK BioBank (https://www.ukbiobank.ac.uk/), ABCD (https://abcdstudy.org/), the Laboratory of NeuroImaging (https://loni.usc.edu/), data made available through the Open Science Framework (https://osf.io/), COllaborative Informatics and Neuroimaging Suite Data Exchange tool (COINS; http://coins.mrn.org/dx), the Developing Human Connectome Project (http://www.developingconnectome.org/), the Human Connectome Project (http://www.humanconnectomeproject.org/), the OpenPain project (https://www.openpain.org), the International Neuroimaging Datasharing Initiative (INDI; http://fcon_1000.projects.nitrc.org/), and the NIMH Data Archive (https://nda.nih.gov/); see **SI21** for details on open human MRI science..

## Data and code availability

All code is made available on https://github.com/ucam-department-of-psychiatry/Lifespan and summary statistics are provided in the **Supplementary Tables (ST1-8)** and through www.brainchart.io. Sharing or re-sharing of MRI scans aggregated here is through application procedures managed at the discretion of each contributing study individually.

## Contributions

RAIB, JS, SRW, ETB and AFA-B designed the study, conducted analyses, wrote and edited the manuscript. JV and KA helped to design the study and contributed to data analysis. All other authors made a significant contribution in one or more of the following areas: design of the study, acquisition or analysis of data, provision of software, or drafting or substantive review of the paper.

## Online methods: Brain charts for the human lifespan

To accurately and comprehensively establish standardised brain reference charts across the lifespan, it is crucial to leverage multiple independent and diverse datasets, especially those spanning prenatal and early postnatal life. Here we sought to chart normative brain development and ageing across the largest age-span and largest aggregated neuroimaging dataset to date using a robust and scalable methodological framework^1,2^. We leveraged these normative reference charts in clinical cohorts to generate individualised assessments of age-relative centiles. These centiles were then leveraged to investigate cross-diagnostic and longitudinal atypicalities of brain morphology across the lifespan. We used generalised additive models for location, scale and shape (GAMLSS)^1^ to estimate cross-sectional normative age-related trends from 100 studies (see supplementary tables [**ST**] **1.1-1.7** for full demographic information and supplementary information [**SI**] **19** for dataset descriptions). The GAMLSS approach allows not only modelling of age-related changes in brain phenotypes but also age related-changes in the variability, in the form of both linear and nonlinear changes over time, thereby overcoming potential limitations of conventional additive models that only allow additive means to be modelled^1^. In addition, site-specific offsets (mean and variance) for each brain phenotype are also modelled. These modelling criteria are particularly important in the context of establishing growth references as recommended by the World Health Organisation^2^, as it is reasonable to assume the distribution of higher order moments (e.g., variance) changes with age, sex, site/study and pre-processing pipeline—especially given the impossibility of fully comprehensive longitudinal data for individuals spanning the ∼100 year age range. Furthermore, recent studies suggest that changes in across-individual variability might intersect with vulnerability for developing a mental health condition^3^. The use of data spanning the entire age range is also critical, as estimation from partial age-windows can lead to biased estimations when extrapolated to the whole lifespan. In summary, using a sex-stratified approach^2^, age, preprocessing pipeline and study were each included in the GAMLSS model estimation of first order (*μ*) and second order (***σ***) distribution parameters of a generalised gamma distribution using fractional polynomials to model nonlinear trends. See **Supplementary Information** [**SI**] for more details regarding GAMLSS model specification and estimation (**SI1**), image quality control (**SI2**), model stability and robustness (**SI3-4**), phenotypic validation against non-imaging metrics (**SI3 & SI5.2**), inter-study harmonisation (**SI5**) and assessment of cohort effects (**SI6**).

In general, the GAMLSS framework can be specified in the following way:

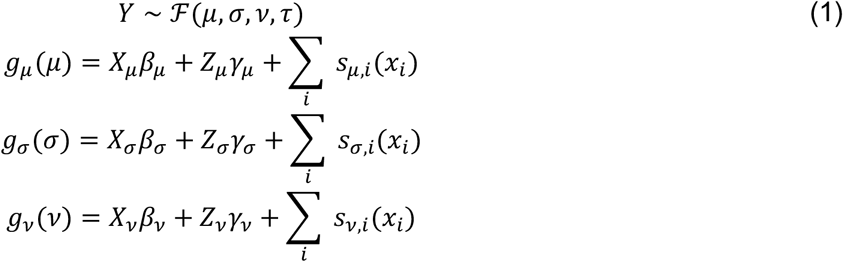

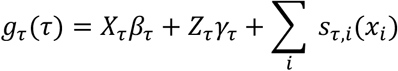

Here, the outcome vector, *Y*, follows a probability distribution *F* parameterised by up to four parameters, (*μ, σ, ν, τ*). The four parameters, depending on the parameterisation of the probability density function, may correspond to the mean, variance, skewness, and kurtosis (i.e., the first four moments); however, for many distributions there is not a direct one-to-one correspondence. Each component is linked to a linear equation through a link-function, *g*_·_(), and each component equation may include three types of terms: fixed effects, *β*_·_(with design matrix, *X*_·_); random-effects, *γ*_·_ (with design matrix, *Z*_·_); and non-parametric smoothing functions, *s*_*·,i*_ applied to the *i*^th^ covariate. The nature of the outcome distribution determines the appropriate link-functions and which components are used. In principle any outcome distribution can be used, from well-behaved continuous and discrete outcomes, through to mixtures and truncations.

Within this paper we consider fractional polynomials as a flexible, yet limited in complexity, approach to modelling age-related changes. Although non-parametric smoothers are more flexible, they can become unstable and infeasible, especially in the presence of random-effects. Hence, the fractional polynomials enter the model within the *X*_·_terms, with associated coefficients in *β*_·_. The GAMLSS framework includes the ability to estimate the most appropriate powers within the iterative fitting algorithm, searching across the standard set of powers, *p* ∈ {−2, −1, −0.5,0,0.5,1,2,3}, where the design matrix includes the covariate (in our setting, age) raised to the power, namely, *x*^*p*^. Fractional polynomials naturally extend to higher-orders, for example a second-order fractional polynomial of the form, 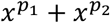.

There are several options for including random-effects within the GAMLSS framework depending on the desired covariance structures. We consider the simplest case, including a factor-level (or group-level) random-intercept, where the observations are grouped by the study covariate. The random-effects are drawn from a normal distribution with zero mean and variance to be estimated, *γ*_·_ ∼ *N*(0, *δ*_·_^2^). The ability to include random-effects is fundamental to accounting for co-dependence between observations. It is therefore possible to take advantage of the flexibility of “standard” GAMLSS, as typically used to develop growth charts^2,4,5^, while accounting for co-dependence between observations using random-effects. The typical applications of GAMLSS assume independent and identically distributed outcomes; however within our context it is essential to account for within-study covariance implying the observations are no longer independent.

The resulting models were evaluated using several sensitivity analyses and validation approaches. See **Supplementary Information** [**SI**] for further details regarding GAMLSS model specification and estimation (**SI1**), image quality control (**SI2**), model stability and robustness (**SI3-4**), phenotypic validation against non-imaging metrics (**SI3 & SI5.2**), between-study harmonisation (**SI5**) and assessment of cohort effects (**SI6**). While the models of whole brain and regional morphometric development were robust to variations in image quality, and cross-validated by non-imaging metrics, we expect that several sources of variance, including but not limited to MRI data quality and variability of acquisition protocols, may become increasingly important as brain charting methods are applied to more innovative and/or anatomically fine-grained MRI phenotypes. It will be important for future work to remain vigilant about the potential impact of data quality and other sources of noise on robustness and generalisability of both normative trajectories and the centile scores derived from them.

Based on the model selection criteria, outlined in the Supplementary Information (**SI 1**) in detail, the final models for normative trajectories of all MRI phenotypes were specified as illustrated below for GMV:

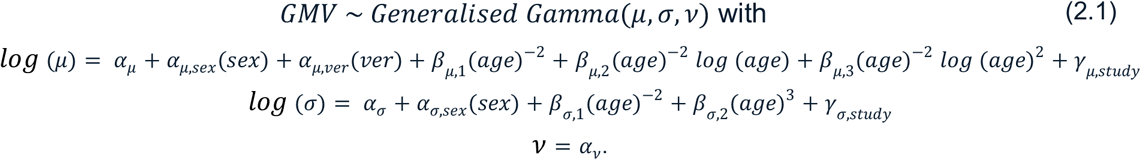

For each component of the generalised gamma distribution, *α* terms correspond to fixed effects of the intercept, sex (female/male), and software version (five categories); *β* terms correspond to the fixed effects of age, modeled as fractional polynomial functions with the number of terms reflecting the order of the fractional polynomials; and *γ* terms correspond to the study-level random-effects. Note that we have explicitly included the link-functions for each component of the generalised gamma, namely the natural logarithm for *μ* and *σ* (since these parameters must be positive) and the identity for *ν*.

Similarly for the other phenotypes:

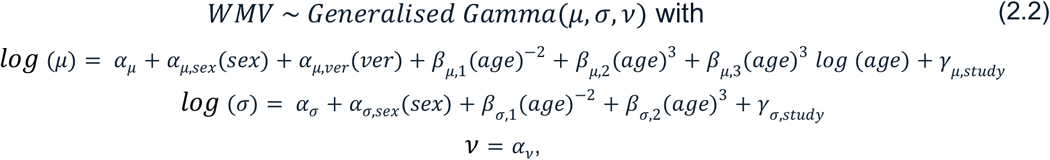

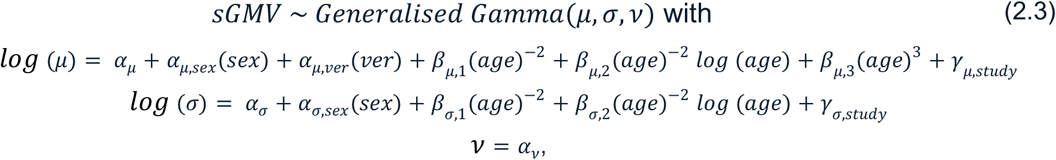

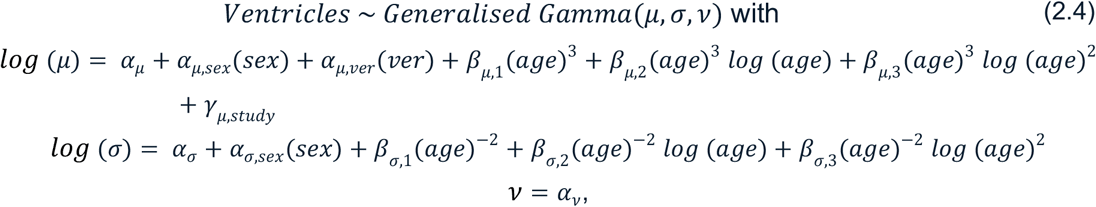

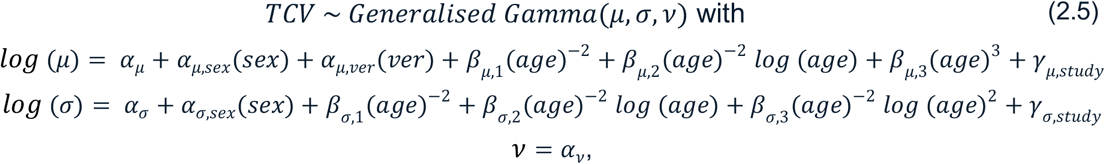

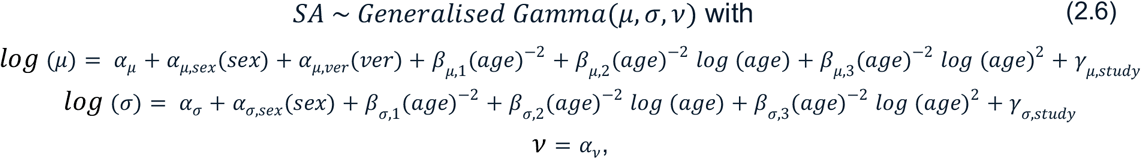

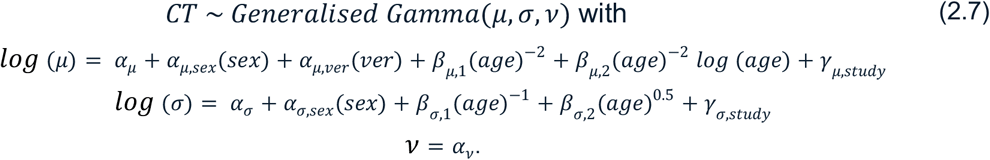

No smoothing terms were used in any GAMLSS models implemented in this study, although the fractional polynomials can be regarded as effectively a parametric form of smoothing. Reliably estimating higher order moments requires increasing amounts of data, hence none of our models specified any fixed-nor random-effects in the *ν* term. However, *α*_*ν*_ was found to be important in terms of model fit and hence we have used a generalised gamma distribution.

This approach also allowed us to leverage the aggregated life-spanning neuroimaging dataset to derive developmental milestones (i.e., peaks of trajectories) and compare them to existing literature. These sex-stratified models incorporated variation in study and processing pipeline to allow computation of standardised reference charts. The cerebrum tissue classes from 100 studies (**Fig. 1A, ST1.1-1.7, SI18**) showed clear, predominantly age-related trends, even prior to any modelling (**Fig. 1B**). Yet, marked heterogeneity of growth curves for individual studies (www.brainchart.io) reinforces the importance of using the full aggregated dataset to achieve representative norms not biased by individual studies. The validity of the models is supported by high stability under cross-validation and bootstrap resampling (**SI3**). Comparing these models to multiple non-MRI metrics of brain size demonstrated high correspondence across the lifespan (**SI3**). Peaks were determined based on the GAMLSS model output (50^th^ centile) for each of the tissue classes and TCV, for both total tissue volumes (**Fig. 1B**) and rates of change or growth (“velocity”; **Fig. 1E**). Diagnostic age ranges from previous literature^6,7^ were plotted (blue boxes in **Fig. 3**) to compare with empirical age ranges of patients with a given diagnosis across the aggregated neuroimaging dataset (black boxes in **Fig. 3**). Note that age of diagnosis is significantly later than age of symptom onset for many disorders^6^. Developmental milestones were re-plotted based on published work for brain resting metabolic rate (RMR)^8^, from its minimum in infancy to its maximum in early childhood; anthropometric variables (height and weight), which reach a first peak in velocity during infancy and a second peak in velocity in adolescence^9^; and typical acquisition of the six gross motor capabilities^4^. Pubertal age ranges were taken from reported typical age ranges^10,11^.

Furthermore, these neuroimaging-derived brain reference charts also enabled each individual to be quantified relative to a statistical distribution defined at the reference level for any point during the lifespan^12,13^. Individual centile scores were obtained relative to the reference curves, conceptually similar to traditional anthropometric growth charts. These normative scores represent a novel set of population and age standardised clinical phenotypes, providing the capacity for cross-phenotype, cross-study and cross-disorder comparison. A single summary deviation metric for each individual was also generated. Main group effects were analysed with a bootstrapped (500 bootstraps) non-parametric generalisation of Welch’s one-way ANOVA. Pairwise, sex stratified, post-hoc comparisons were conducted using non-parametric Monte-Carlo permutation tests (10,000 permutations) and thresholded at a Benjamini-Hochberg false discovery rate (FDR) of q < 0.05.

To utilise the centiles in a diagnostically meaningful or predictive way, they need to be stable across multiple measuring points. To assess this intra-individual stability we calculated the subject specific interquartile range (IQR) of centiles across timepoints for the datasets that included longitudinal scans (n = 9,306, 41 unique studies). Exploratory longitudinal clinical analyses were restricted to clinical groups that had at least 50 subjects with longitudinal data to allow for robust group-wise estimates of longitudinal variability. In addition, there was a small subset of individuals with documented pathological progression across longitudinal scans, for instance from high-risk status to formal diagnosis. Here, we would expect an associated change in centile measurement. To test this hypothesis we assessed whether these individuals showed differences in centile variability (as assessed with IQR), and their approximate direction of change.

Finally, we provide an interactive tool (www.brainchart.io) and have made our code and models openly available (https://github.com/ucam-department-of-psychiatry/Lifespan). The tool not only allows the user to visualise the underlying datasets’ demographics and reported reference charts in a much more detailed fashion than static images allow, it also provides the opportunity for interactive exploration of differences in centile scores across many clinical groups that is beyond the present manuscript. Perhaps most significantly, it includes an out-of-sample estimator of model parameters for novel data that enables the user to compute percentile scores for their own datasets without the computational or data-sharing hurdles involved in adding that data to the reference chart. All modelling included extensive validation, sensitivity analyses and multi-modal validation against existing growth chart references.

Though already based on the largest and most comprehensive neuroimaging dataset to date and supporting analyses of out-of-sample data, the underlying reference charts will also be updated as additional data is made available.

