## Supplemental information for "Brain charts for the human lifespan"

Bethlehem, R.A.I.[1,2]#, Seidlitz J.[3,4]#, White, S.R.[5,6]#, Vogel, J.W.[7,3], Anderson, K.M.[8], Adamson C.[9,10], Adler S.[11], Alexopoulos, G.S.[12], Anagnostou E.[13,14], Areces-Gonzalez A.[15,16], Astle, D.E.[17], Auyeung B.[18,1], Ayub M.[19], Ball G.[9,20], Baron-Cohen S.[1,21], Beare R.[9,10], Bedford, S.A.[1], Benegal V.[22], Beyer F.[23], Bin Bae J.[24], Blangero J.[25], Blesa Cábez M.[26], Boardman, J.P.[26], Borzage M.[27], Bosch-Bayard, J.F.[28,29], Bourke N.[30], Calhoun, V.D.[31], Chakravarty, M.M.[32,29], Chen C.[33], Chertavian C.[3], Chetelat G.[34], Chong, Y.S.[35,36], Cole, J.H.[37,38], Corvin A.[39], Costantino M.[40,41], Courchesne E.[42,43], Crivello F.[44], Cropley, V.L.[45], Crosbie J.[46], Crossley N.[47,48], Delarue M.[34], Delorme R.[49,50], Desrivieres S.[51], Devenyi G.[52,53], Di Biase, M.A.[45,54], Dolan R.[55,56], Donald, K.A.[57], Donohoe G.[58], Dunlop K.[59], Edwards, A.D.[60,61], Elison, J.T.[62], Ellis, C.T.[8], Elman, J.A.[63], Eyler L.[64,65], Fair, D.A.[62], Feczko E.[62], Fletcher, P.C.[66,67], Fonagy P.[68], Franz, C.E.[69], Galan-Garcia L.[70], Gholipour A.[71], Giedd J.[72,73], Gilmore, J.H.[74], Glahn, D.C.[75,76], Goodyer I.[5], Grant, P.E.[77], Groenewold, N.A.[78,79], Gunning, F.M.[12], Gur, R.E.[3,80], Gur, R.C.[3,81], Hammill, C.F.[46,82], Hansson O.[83,84], Hedden T.[85,86], Heinz A.[87], Henson, R.N.[17,88], Heuer K.[89,90], Hoare J.[91], Holla B.[92], Holmes, A.J.[93], Holt R.[1], Huang H.[94,95], Im K.[96,97], Ipser J.[98], Jack Jr, C.R.[99], Jackowski, A.P.[100,101], Jia T.[102,103,104], Johnson, K.A.[105,106,76,107], Jones, P.B.[5,108], Jones, D.T.[109,110], Kahn R.[111,112], Karlsson H.[113,114], Karlsson L.[113,114], Kawashima R.[115], Kelley, E.A.[116], Kern S.[117], Kim K.[118,119,120,121], Kitzbichler, M.G.[5], Kremen, W.S.[63], Lalonde F.[122], Landeau B.[34], Lee S.[123], Lerch J.[124,125,126], Lewis, J.D.[127], Li J.[128], Liao W.[128], Liston C.[129], Lombardo, M.V.[130,1], Lv J.[45,131], Lynch C.[59], Mallard, T.T.[132], Marcelis M.[133], Markello, R.D.[134], Mathias, S.R.[75], Mazoyer B.[44,135], McGuire P.[136], Meaney, M.J.[137], Mechelli A.[136], Medic N.[5], Misic B.[134], Morgan, S.E.[138,139,140], Mothersill D.[141,142,143], Nigg J.[144], Ong, M.Q.W.[145], Ortinau C.[146], Ossenkoppele R.[147,148], Ouyang M.[94], Palaniyappan L.[149], Paly L.[34], Pan, P.M.[150,151], Pantelis C.[152,153,154], Park, M.M.[155], Paus T.[156,157], Pausova Z.[46,158], Paz-Linares D.[15,159], Pichet Binette A.[160,161], Pierce K.[162], Qian X.[145], Qiu J.[163], Qiu A.[164], Raznahan A.[122], Rittman T.[165], Rodrigue A.[75], Rollins, C.K.[166,167], Romero-Garcia R.[5,168], Ronan L.[5], Rosenberg, M.D.[169], Rowitch, D.H.[170], Salum, G.A.[171,172], Satterthwaite, T.D.[7,3], Schaare, H.L.[173,174], Schachar, R.J.[46], Schultz, A.P.[105,175,76], Schumann G.[176,177,178], Schöll M.[179,180,181], Sharp D.[30], Shinohara, R.T.[33,182], Skoog I.[117], Smyser, C.D.[183], Sperling, R.A.[105,106,76], Stein, D.J.[184], Stolicyn A.[185], Suckling J.[5], Sullivan G.[26], Taki Y.[115], Thyreau B.[115], Toro R.[186,90], Traut N.[186,187], Tsvetanov, K.A.[165,188], Turk-Browne, N.B.[8,189], Tuulari, J.J.[190,191,192], Tzourio C.[193], Vachon-Presseau É.[194,195,196], Valdes-Sosa, M.J.[70], Valdes-Sosa, P.A.[197,198], Valk, S.L.[199], van Amelsvoort T.[200], Vandekar, S.N.[201,202], Vasung L.[203], Victoria, L.W.[12], Villeneuve S.[161,204,160], Villringer A.[23,205], Vértes, P.E.[5,206], Wagstyl K.[207], Wang, Y.S.[208,209,210,211], Warfield, S.K.[71], Warrier V.[1], Westman E.[212], Westwater, M.L.[5], Whalley, H.C.[185], Witte, A.V.[205,23,213], Yang N.[208,209,210,211], Yeo B.[214,215,216,217], Yun H.[218], Zalesky A.[219], Zar, H.J.[78,220], Zettergren A.[117], Zhou, J.H.[145,221,214], Ziauddeen H.[5,222,223], Zugman A.[224,225], Zuo, X.N.[208,209,210,211,226], 3R-BRAIN*, AIBL**, Alzheimer's Disease Neuroimaging Initiative***, Alzheimer's Disease Repository Without Borders Investigators****, UMN BCP*****, CALM Team******, Cam-CAN*******, CCNP********, COBRE*********, Developing Human Connectome Project, ENIGMA Developmental Brain Age working group**********, FinnBrain, Harvard Aging Brain Study***********, IMAGEN************, KNE96*************, The Mayo Clinic Study of Aging, NSPN**************, POND***************, The PREVENT-AD Research Group, VETSA, Bullmore, E.T.[5]†, Alexander-Bloch, A.F.[3,4]†

### contributed equally to the work

† jointly supervised the work

Corresponding authors:

Richard A.I. Bethlehem,

Jakob Seidlitz,

Common nomenclature 7

Supplementary Methods 8
1. Modelling lifespan trajectories of brain maturation 8
1.1 Model distributions 11
1.2 Convergence within GAMLSS 12
1.3 Fractional polynomial model set 13
1.4 Model simulations 14
1.5 Centile normalisation 16
1.6 Centile Mahalanobis distance 17
1.7 Longitudinal centiles 18
1.8 Out-of-sample estimation 20
2. Quality control 21
2.1 Euler index filtering 22
2.2 Expert visual quality control 24
2.3 Image quality and out-of-sample centile scoring 27
2.4 Euler index and neuroimaging phenotypes 27

Sensitivity Analyses 31
3. Model evaluation 31
3.1 Model diagnostics 31
3.2 Model sensitivity analyses 32
3.2.1 Leave-one-study-out 33
3.2.2 Bootstrap analysis 33
3.2.3 Parameter estimates 34
3.3 Study-specific curves 39
3.4 Brain weight, ultrasound, and head circumference validation 40
4. Out-of-sample centile scoring: bias, stability and reliability 41
4.1 Bias of out-of-sample centile scores: leave-one-study-out analyses for 100 studies 41
4.2 Stability of out-of-sample centile scoring: bootstrapped LOSO analyses for 100 studies 44
4.3 Test-retest reliability of out-of-sample centile scoring 46
4.4 Reliability of out-of-sample centile scoring across multiple versions of FreeSurfer 49
4.5 Effects of sample size on reliability of out-of-sample centile scores 50
5. Batch correction and site harmonisation 51
5.1 Modeling of between-site heterogeneity by GAMLSS: conceptual considerations in comparison to ComBAT batch-correction 51
5.2 Modeling of between-site heterogeneity by GAMLSS: empirical evaluation compared to ComBAT 53
6. Cohort effects 58

Supplementary Analyses 62
7. Extended global cortical phenotypes 62

7.1 Model optimisation 62
7.2 Normative trajectories of extended global MRI phenotypes 64
7.3 Quality control of extended global MRI phenotypes 69
7.4 Stability of out of-sample centile scoring for extended global phenotypes: LOSO analyses 70
8. Regional cortical volumetric trajectories and milestones 72
8.1 Charting development of regional volumes 73
8.2 Regional volumetric milestones 76
9. Developmental windows and milestones 78
9.1 Trajectories within developmental epochs 78
9.2 Grey-white matter differentiation 81
10. Clinical applications of centile scores 81
10.1 Case-control and between-disorder comparisons of centile scores on cerebrum tissue volumes and extended global MRI phenotypes 82
10.2 Multimodality of centile distributions in clinical disorders 87
10.3 Case-control differences on CMD 90
10.4 Summary centile comparison 91
11. Cross diagnostic analyses 93
11.1 Sliding window analyses of cross-disorder discriminability 93
11.2 Cross-diagnostic clustering 94
12. Associations of birth weight and gestational duration with centile scores on cerebrum tissue volumes 99
13. Twin-based heritability of centile scores 100
14. Longitudinal centiles 100
14.1 Longitudinal patterns in developmental epochs 100
14.2 Longitudinal variability across studies 102
14.3 Longitudinal variability in clinical samples 104
14.4 Longitudinal centile score changes and diagnostic progression 110
15. Interactions between cerebrum tissue volumes 111
16. Sex differences 113
Reference database details: demographics and processing pipelines 115
17. Demographics of reference database 115
18. Data processing 116
19. Primary dataset descriptions 117
3R-BRAIN - Brain Consortium for Reproducibility, Replicability and Reliability 117
ABCD - Adolescent Brain and Cognitive Development 117
ABIDE - Autism Brain Imaging Data Exchange 117
ABVIB - Aging Brain: Vasculature, Ischemia, and Behavior 117
ACE and IBIS 118
ADHD200 118
ADNI - Alzheimer's Disease Neuroimaging Initiative 118

AIBL - Australian Imaging. Biomarkers and. Lifestyle Flagship Study of Ageing 119
AOBA 119
AOMIC ID1000, PIOP1 & PIOP2 - Amsterdam Open MRI Collection 119
ARWIBO - Alzheimer's disease Repository Without Borders 120
ASRB - Australian Schizophrenia Research Bank 120
BCP - Baby Human Connectome Project 120
BGSP - Brain Genomics Superstruct Project 121
BHRCS - Brazilian High Risk Cohort Study for Mental Conditions 121
BioDep - Biomarkers of Depression 121
BSNIP - Bipolar & Schizophrenia Consortium for Parsing Intermediate Phenotypes 122
Calgary 122
CALM - Centre for Attention Learning and Memory 122
Cam-CAN - Cambridge Centre for Aging and Neuroscience 122
CAM-FT - Cambridge foetal testosterone 123
CCNP-devCCNP - Chinese Color Nest Project devCCNP 123
Cuban Human Brain Mapping Project (CHBMP) 123
CHILD - Cambridge Human Imaging and Longitudinal Development 123
COBRE - Center for Biomedical Research Excellence 124
CONTE 124
Cornell 125
CTAAC - Cape Town Adolescent Antiretroviral Cohort 125
cVEDA - Consortium on Vulnerability to Externalizing Disorders and Addictions 125
dHCP - Developing Human Connectome Project 125
DCHS - Drakenstein Child Health Study 126
DCHS Infants 126
DCHS Mothers 126
DLBS - Dallas Lifespan Brain Study 127
EDSD - European DTI Study on Dementia 127
EMBARC - Establishing Moderators and Biosignatures Of Antidepressant Response for Clinical Care 127
Female ASD 128
FinnBrain 128
Frankfurt 128
GOBS - Genetics of Brain Structure and Function study 128
GOSICH - Great Ormond Street Institute for Child Health 128
GUSTO - Growing Up in Singapore Towards Healthy Outcomes 129
HABS - Harvard Aging Brain Study 129
Harvard foetal 129
HBN - Healthy Brain Network 130
Human Connectome Project 130
Human Connectome Project Aging and Development 130

iADNI - Italian Alzheimer's Disease Neuroimaging Initiative 131
ICBM 131
IMAGEN 131
IMAP - Multi-modal Neuroimaging in Alzheimer's Disease 131
IXI 132
KNE96 - Korean normal elderly brain template study 132
LA5c - UCLA Consortium for Neuropsychiatric Phenomics LA5c Study 132
LATAM 132
LIFE - Leipzig Research Centre for Civilization Diseases Study 132
MCIC 133
MCSA (Mayo Clinic Olmsted Study of Aging) 133
MRi-Share 133
Narratives 134
NeuroScience and Psychiatry Network 134
NHGRI 134
NIH 134
NIHPD Infant and Adult 134
NKI - Nathan Kline Institute Rockland Sample 135
NTB_Yale 135
OASIS3 - Open Access Series of Imaging Studies 135
OHSU 136
OpenPain 137
Placebo 1 (PL1) 137
Subacute longitudinal study (SA1 & SA2): 138
Placebo predict Tetreault (PLOA) 138
Brain network change Mano (RS) 138
Accumbens Chronic Pain Signature (SAB) 138
Oslo 139
Oulu 139
Penn-CHOP Developmental Connectome (PCDC) 139
PING - Pediatric Imaging, Neurocognition, and Genetics 139
Pixar 140
PNC - Philadelphia Neuroimaging Cohort 140
POND - Province of Ontario Neurodevelopmental Disorders 140
PPMI - Parkinson's Progression Markers Initiative 140
PREVENT-AD - PRe-symptomatic EValuation of Experimental or Novel Treatments for AD 141
PSYSCAN Maastricht 141
PSYSCAN Dublin 141
RDB 142
SALD - Southwest University Adult Lifespan Dataset 142

SCZIowa 142
SLIM - Southwest University Longitudinal Imaging Multimodal Dataset 143
STRIvE - Stress in Eating 143
SYS Adults & Adolescents 143
TEBC - Theirworld Edinburgh Birth Cohort 144
TOPSY 144
UKB - UK Biobank 144
VETSA - Vietnam Era Twin Study of Aging 145
VITA - Vienna Transdanube Aging study 145

20. Replication/validation datasets 146
10k-in-a-day 146
Hsu et al. (Ultrasound) - estimated 146
Chang et al. (Ultrasound) - estimated 146
Roelfsema et al. (Ultrasound) - estimated 146
Brain Weight 147
International prenatal (HC) 147
WHO postnatal (HC) 147

21. A note on data sharing 148

22. Affiliations of authors 149

23. Acknowledgements 156

24. References 161

### Common nomenclature

AD - Alzheimer's disease
ADHD - Attention deficit hyperactivity disorder
ANX - Anxiety or phobic disorders
ASD - Autism spectrum disorder
BD - Bipolar disorder
BIC - Bayesian information criterion
CDF - Cumulative density function
CN - Control or cognitively normal
CSF - Ventricular cerebrospinal fluid (or ‘Ventricles’)
CMD - Centile Mahalanobis distance
CT - Cortical thickness
DK - Desikan-Killiany
DTOP - Detrended transformed Owen’s plot
DX - Diagnosis (or non-CN)
DZ - Dizygotic
EI - Euler index
Eq - Equation
FDR - False discovery rate
FTD - Frontotemporal dementia
GAM - Generalised additive model
GAMLSS - Generalised additive models for location scale and shape
GEEs - Generalised Estimating Equations
GG - Generalised Gamma
GMV - Total cortical grey matter volume
HC - Head circumference
ICC - Intraclass correlation coefficient
IQR - Interquartile range
LBD - Lewy body dementia
LOESS - Locally estimated scatterplot smoothing
LOSO - Leave-one-study-out
MCI - Mild cognitive impairment
MDD - Major depressive disorder
MRI - Magnetic resonance imaging
MZ - Monozygotic
OCD - Obsessive compulsive disorder
OoS - Out-of-sample
PD - Parkinson’s disease
QC - Quality control
RMR - Resting metabolic rate
TCV - Total cerebrum volume
sGMV - Total subcortical grey matter volume
SA - Surface area
SCZ - Schizophrenia
ST - Supplementary table
SI - Supplementary information
TCV - Total cerebrum volume
WHO - World Health Organisation
WMV - Total cortical white matter volume

### Supplementary Methods

#### 1. Modelling lifespan trajectories of brain maturation

Combining multiple cross-sectional studies covering a part of the lifespan to obtain a reference level lifespan curve, for a range of imaging phenotypes, requires a flexible modelling approach. In this section, we further describe the modelling issues and compare our chosen approach to potential alternative approaches. We also provide details about multiple sensitivity analyses using bootstrapping, study jackknifing and external validation. Furthermore, we outline several simulations that informed our choices of modelling in our final approach. We also note that figures are available in interactive format at www.brainchart.io.

The presence of study-specific effects requires the use of a random-effects modelling strategy, viewing each study as an example drawn from an infinite pool of potential studies rather than estimating a study fixed effect. Ignoring study-specific effects would fail to account for the substantial correlation between individuals within studies, resulting in substantial bias in any estimated parameters and covariances. A fixed-effect framework, modelling each study as an identifiable effect, would induce an ever-growing number of parameters within the model. It is more appropriate to view the set of observed studies as drawn from an infinite population of possible studies. A random-effects approach has the additional benefit of introducing smoothing, or equivalently shrinkage, across the multiple study estimates. There is a priori evidence of meaningful variation in higher moments across the lifespan—not just in the mean (first moment) but also in the variance (second moment)[1–5]. This would preclude, for example, mixed-effect models which only model the mean. Furthermore, there is no biological basis for the assumption that neuroimaging phenotypes follow a Gaussian distribution. This leads us to the set of 'generalised' extensions of common methods, for example generalised mixed-effects models.

The World Health Organization (WHO) recommended[6] framework to address these requirements within a growth chart context is the Generalised Additive Models for Location Scale and Shape (GAMLSS)[7–10] modelling approach. The flexible outcome distribution component allows us to consider non-Gaussian outcomes, for example the generalised gamma distribution[8]. Importantly the GAMLSS framework allows us to incorporate models for multiple moments of the outcome distribution (the Location [L], Scale [S] and Shape [S] within GAMLSS), for example modelling age-related changes in the mean, variance and skewness of imaging features. Depending on the outcome distribution, the moments may be defined in terms of the parameters of the distribution, e.g., the generalised gamma[11] is a three parameter distribution where the three moments are calculated in terms of the modelled parameters (rather than being directly modelled). See **SI1.1** for additional information about model distributions.

With respect to covariates (or independent variables, or predictors), the GAMLSS framework encompasses generalised additive models (GAMs), which allow a highly flexible format for the relationship between the covariates and the outcome (or dependent variable). This includes the ability to use flexible spline functions. However, the strong association between studies and period of the lifespan, specifically the coverage density and range of the lifespan by studies, would likely conflate with selecting knot locations for a spline model and may be unstable under the inclusion and exclusion of specific studies each covering specific ranges of the full lifespan. Despite having a substantial number of observations, allowing flexible splines within multi-parameter distributions with random-effects would induce too much flexibility. Instead, we consider a simpler additive construct, namely fractional polynomials[12]. Fractional polynomials can be viewed as a simpler form of spline modelling using a fixed set of polynomials. Following the standard approach for fractional polynomials, we consider a set of powers (the typical set of eight powers) across a number of orders of fractional polynomials. The GAMLSS framework includes a fractional polynomial function that automatically performs this model selection step within the fitting process, although in addition to this standard estimation we chose to evaluate model permutations of all possible combinations of the number of modelled polynomials in each term of the generalised gamma distribution (between 1–3 terms for each of the three parameters). In order to assess the uncertainty in our lifespan curves, we use a stratified bootstrapping procedure. It is important to stratify by study to ensure the bootstrap resampling maintains equivalent coverage of the lifespan. Further, we included a sex-effect as an a priori important feature and so also stratify on sex. See **SI1.3** for additional information about fractional polynomial model selection, and **SI1.4** for a discussion of the GAMLSS framework using simulated data.

Our modelling aim is to obtain a reference lifespan model for cognitively normal subjects (CN), onto which we can place individuals with various diseases and conditions. Furthermore, there could be variability between studies in the standards used for diagnosis of disorders and for ascertainment of healthy controls, and the non-CN individuals are spread across the studies. To that end, we require study-specific estimates from our model so that we can derive study-specific CN reference curves. This requirement excludes many non-parametric outcome distribution approaches and conditional inference approaches, for example Generalised Estimating Equations (GEEs)[13], since they explicitly avoid or side-step estimating the random-effects terms. Although many of these methods reduce bias under model mis-specification, we require the feature they integrate out. Similarly, this excludes approaches like the recent application of ComBAT[14] in neuroimaging from its origins in the genomics literature (see **SI5** for an in-depth comparison between the two approaches). The principal way in which we have accounted for inter-site variability issues is by using the GAMLSS modeling framework to correct for between-study differences in all studies, and to use only data from healthy controls to estimate the normative trajectories for all phenotypes. To demonstrate the robustness to study-specific variability of growth curves and the individual centiles derived from them, we conducted ‘leave-one-study-out’ (LOSO) analyses whereby the growth curves and centiles are repeatedly estimated after exclusion of each individual study (see **SI3.2** on model sensitivity analyses). These analyses confirm that trajectories from the total dataset are in general highly conserved after exclusion of each individual study, suggesting that study-specific differences do not materially influence model parameters.

Moving from Gaussian outcomes to general outcome distributions, with models for each parameter of the distribution, it becomes difficult to work in terms of the outcome measure directly. Rather, since we are working with probability distributions we consider the centile of the observation, the 50th centile being the median. This is in contrast to the commonly used approach of 'standardising' the predictions, for example with z-scores. This is important especially for non-symmetric or skewed distributions like the generalised gamma where the mean and median can differ, and 'z-scoring' (i.e., (x-μ)/σ)) is inappropriate or invalid.

We can obtain study-specific offsets and then derive reference-normalized outcomes for each individual's observation. Specifically, we can obtain a study-specific centile for the ith person as:

$$q_i = F'(y, x|\beta, z) \tag{1.1}$$

using the inverse cumulative density function, F', with the model fixed-effect parameters, $\beta$, and study random-effect, z, combined with the participant covariates, x, and outcome measure, y. The 'normalised' value, would be calculated using

$$y_{norm} = F(q_i|\beta). \tag{1.2}$$

These reference curves can be used to assess Dx individuals' centiles. If the non-CNs are comparable to the CNs then their centiles will span the zero to one range uniformly. Deviation from such a uniform distribution (within the study-specific centiles) indicates a deviation from the CN lifespan curve. See **SI1.4-1.5** for additional information about centile normalisation and centile deviation relative to reference curves.

Some of the included studies were longitudinal, including multiple follow-up scans for the same individuals. This enables us to investigate the validity of the cross-sectionally derived lifespan curves as longitudinal trajectories. For CNs, assuming a reasonable level of within-person variability, each individual's centile score should remain constant. This would be the equivalent of maintaining a specific centile while ageing on a growth chart. Significant changes in an individual's centile as they age, for example passing from 75th to 25th percentile in height or weight on the child growth chart is an indicator of a potential clinical issue. Our derived neuroimaging reference curves can be used in a similar fashion, using follow-up scans to assess whether an individual has crossed multiple deciles/quartiles/etc. of our lifespan curves. The utility of this approach for cases of extreme pathology may be limited, since they typically sit in an extreme part of the centile distribution (i.e., they are not uniformly distributed from zero to one). While their extreme place relative to the CN distribution is likely to be clinically meaningful, their amount of longitudinal variability might be compressed. To circumvent some degree of complexity we sought to quantify an individual's interquartile range of their position in the centile distribution as a proxy for longitudinal stability. See **SI1.7** for additional information about longitudinal centiles.

Once the GAMLSS model is fitted to a reference database we obtain a parameterisation of the fixed-effects of the curve, estimates of the random-effect covariance structure (the parameterisation of the population of potential studies), and random-effects terms for any study within the database. Using these estimates, specifically the fixed-effects and random-effect covariance structure, we can derive a random-effects term for a new study by using new data from that study and conditioning on the estimated parameters within the model. The stability of the newly estimated study random-effects will depend on the number of observations available. This can be partly quantified using the bootstrap parameter estimates, and we simulated several study sample sizes to determine an approximate minimum sample required for such out-of-sample (OoS) estimation. Crucially, the process described does not require refitting the model to the reference database, nor does it require revealing the original data from the reference database, making it both computationally effective as well as fitting within an open science framework. However, as more studies become available and sample sizes of new studies increase, we aim to periodically update the model fit to an updated/expanded reference database. See **SI1.8** for additional information about the application of models to OoS data.

Finally, it is worth noting that the strategic intent of this study (and some directly relevant prior work) was to quantify brain structural MRI phenotypes relative to age- and sex-specific norms, rather than to predict chronological or biological age of participants from their MRI data[15,16]. There is a large extant literature on attempts to predict “brain age” and compare brain age to the actual age of study participants[16–21]. In contrast we do not ask the question: what is a participant’s neurobiological age, or the difference between their neurobiological and chronological ages[22], given their brain morphology? Instead we ask: how (a)typical is a participant’s brain structure compared to their demographically matched peer group? More formally, we assess the vertical deviation of an individual scan from the normative trajectory of the corresponding phenotype in a reference population; whereas brain-age prediction attempts to quantify the horizontal deviation. Brain charting is more analogous than brain age prediction to the ways that traditional growth charts are used in pediatric practice for anthropometric variables. Additionally, normative growth curves allow us to benchmark even a single MRI phenotype – such as one of the global tissue volumes that are abundantly available across primary datasets – as opposed to brain age predictions that typically require a high-dimensional feature space comprising multiple MRI phenotypes[15,22]. In addition, several methodological critiques of brain age prediction are not relevant to the present approach[22–24]. Thus, we note that using GAMLSS to quantify centile dispersion of MRI phenotypes on age-normed and sex-stratified distributions shares conceptual goals with, but methodologically entirely distinct from, studies that seek to predict human age (or derive a ‘brain age gap’) from brain imaging data[25].

#### 1.1 Model distributions

The WHO provides several guidelines for choosing underlying model distributions for growth chart modelling[6,26] of anthropometrics such as head circumference, height and weight. For these phenotypes the guidelines suggest the Box-Cox t-distribution might be an appropriate starting place. Since we did not want to assume that the trajectories obtained from in vivo neuroimaging would automatically or by extension be best captured by the same underlying distributions, we decided to evaluate all possible distribution families. In that context it is worth noting that there is no specific statistical test to determine the best outcome distribution to use within the GAMLSS model, and the package provides a range of distribution families[8]. Standard practice is to fit multiple distributions, each consistent with the characteristics of the outcome, and to compare the distributions using an information criterion such as the Bayesian Information Criterion (BIC)[27]. We used this approach to select the optimal outcome distribution, keeping in mind the likely need for multi-parameter outcome estimates (so we only evaluated distribution families with 3 or more parameters for which GAMLSS fitting converged). **Fig. S1.1** shows the BIC for all evaluated models scaled to the lowest BIC for each phenotype. In all cases a generalised gamma distribution provided the best fit.

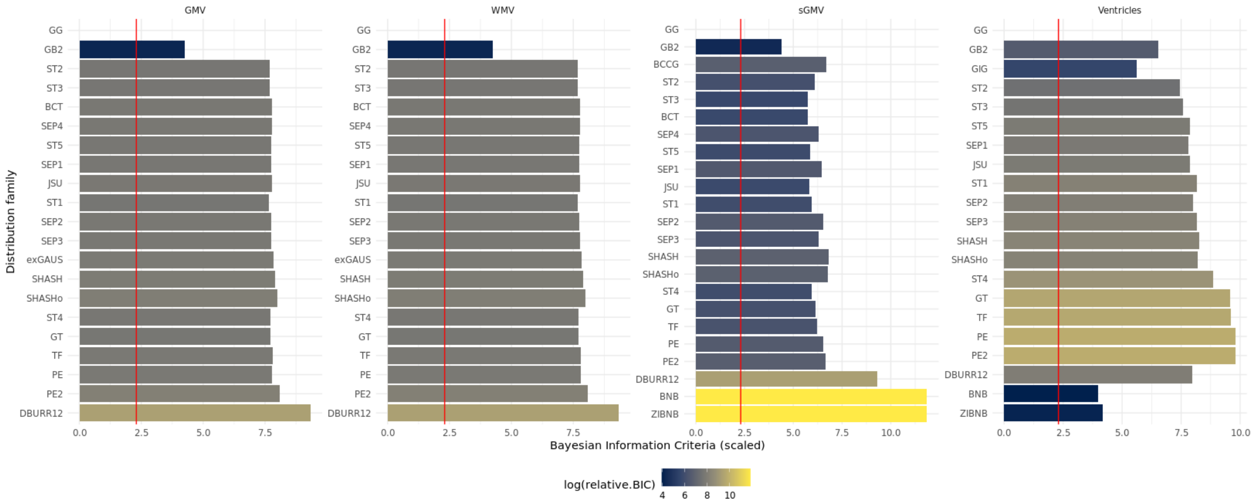


***Fig. S1.1. Relative Bayesian information criterion (BIC) for each family of distributions of cerebrum tissue volumes evaluated for GAMLSS modeling.*** *Log BIC scores are shown in terms of their difference from the lowest BIC score, corresponding to the best-fitting form of the outcome distribution. All BIC values were scaled to the lowest value for each cerebrum tissue volume. For all phenotypes, a generalised gamma (GG) distribution provided the best fit. Distribution family acronyms are adapted directly from the way they are listed within the GAMLSS package [8].*

#### 1.2 Convergence within GAMLSS

Model convergence within GAMLSS, like many iteratively fitting statistical models, is defined in terms of the estimated likelihood staying equivalent across several iterative steps, where equivalence is in terms of a defined convergence threshold. The threshold is with respect to changes in the (log-)likelihood between iterations (we use the default convergence criterion of 0.001)[7,8]. Instability, or non-convergence, is typically when the GAMLSS model cannot converge on a maximum likelihood estimate and jumps between multiple solutions, whose likelihood values differ by more than the threshold and hence the algorithm never converges.

If the model is over-parameterised there may be multiple solutions that fit the data, which will lead to non-convergence. Equivalently, within the bootstrapping procedure, it is possible for a bootstrap replication to become degenerate, meaning the resampled subset of data causes the model fitting to fail, e.g., the bootstrap replicate of a small study may, by chance, consist of copies of only one subject and have no variability with which to estimate the study random-effects. We employ a stratified bootstrap procedure to limit this issue (see **SI3.2.2 ”Bootstrap analysis”**); but given the sample size of some primary studies we experienced a small number (<1%) of model convergence failures across bootstrap replicates. A priori, we deemed the model unstable if more than 5% of bootstrap replicates failed to converge but this situation did not occur for any of the MRI phenotypes.

#### 1.3 Fractional polynomial model set

As noted above fractional polynomials can be viewed as a simpler form of spline modelling using a fixed set of polynomials (GAMLSS uses the standard set of polynomials powers: -2, -1, -0.5, 0, 0.5, 1, 2, 3, see Royston & Altman (1994) [12]). The GAMLSS framework includes a fractional polynomial function that automatically performs this model selection step within the fitting process. In addition to this standard estimation we chose to evaluate model permutations of all possible combinations of the number of modelled polynomials in each of the terms $(\mu, \sigma, \nu)$ of the generalised gamma distribution (between 1–3 terms for each of the three parameters). Across all four phenotypes this approach suggested 3rd order polynomial fits for $\mu$ and with the exception of subcortical grey matter volume (sGMV) also indicated 3rd order polynomial fits for $\sigma$ (**Fig. S1.3**). For white matter volume (WMV) and grey matter volume (GMV) this evaluation procedure suggested including a study random effect in both $\mu$ and $\sigma$, whereas for ventricular cerebrospinal fluid volume (CSF) and sGMV it indicated inclusion of a study random effect only for $\sigma$. Despite the size of the present dataset, we found that fractional polynomial modelling for $\nu$ resulted in model instability (e.g., the GAMLSS model selection did not converge on an optimal parameterisation). Polynomials for $\nu$ were therefore not evaluated as we did not have an a priori reason to assume age and random-effect dependent skewness.

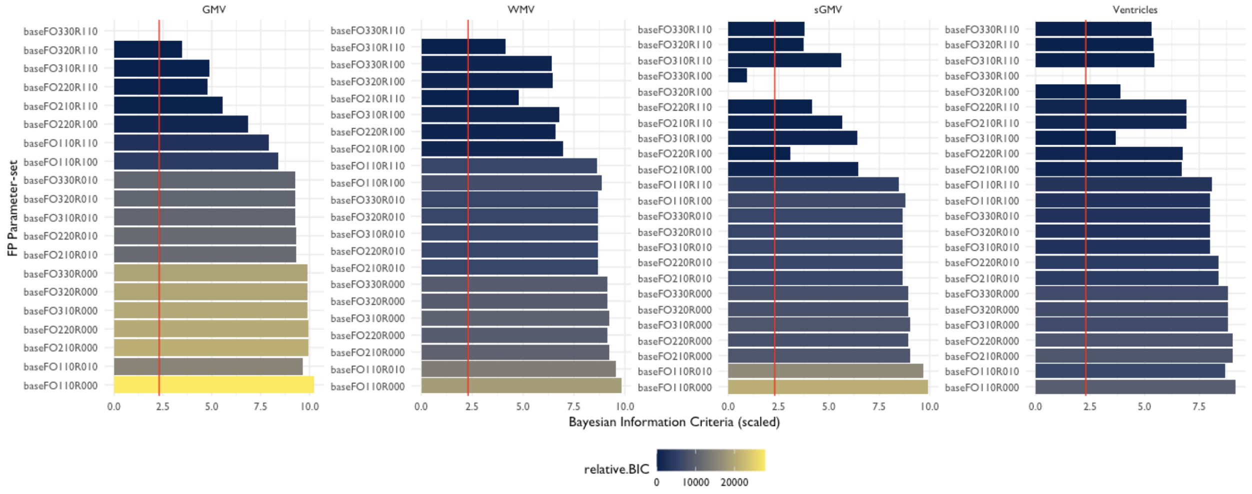


***Fig. S1.3. Optimization of GAMLSS model specification by analysis of the Bayesian information criterion (BIC) for multiple possible models on the generalised gamma distribution.*** *Here log BIC is plotted relative to the best-fitting model with lowest BIC for each combination of fractional polynomials and random effects for which the model converged. All BIC values were scaled to the lowest value for the set of models fitted to each cerebrum tissue volume (log-scored difference to the lowest scoring model). For all phenotypes, a model that included 3 polynomials for $\mu$ provided the best fit; and for all phenotypes other than sGMV the best fit also specified3 polynomials for $\sigma$. The various models fitted are summarised by y-axis labels denoting the base fractional polynomial configuration ("baseFO") that are structured as follows: baseFO[a][b][c]R[x][y][z], where a-c denote the number of fractional polynomials included in the age term on $\mu$, $\sigma$, and $\nu$ respectively, and x-z denote whether a study random effect was estimated for each of the model components (1 means a study random effect was included, 0 means no study random effect was included).*

#### 1.4 Model simulations

In order to motivate the specific use of GAMLSS for lifespan modelling as done here, we designed a simulation scenario that matches our use case for a single outcome or MRI phenotype. Specifically, we simulated data from twenty studies across the lifespan. We simulated data on both healthy controls (CN) and diagnosed cases (DX), some with longitudinal follow-up, as well as study-specific random-effects. We chose the generalised gamma for the true outcome distribution with age and sex fixed-effects, random-effects within the $\mu$-component, and constant $\sigma$- and $\nu$-components. The lifespan relationship was quadratic with age. Importantly, the simulated data also included a subject-level random-effect, which is fitted by the GAMLSS model. This allowed us to set the within- and between-subject covariance, which in turn allowed us to assess the utility of the longitudinal centiles (see **SI1.7 "Longitudinal centiles"**).

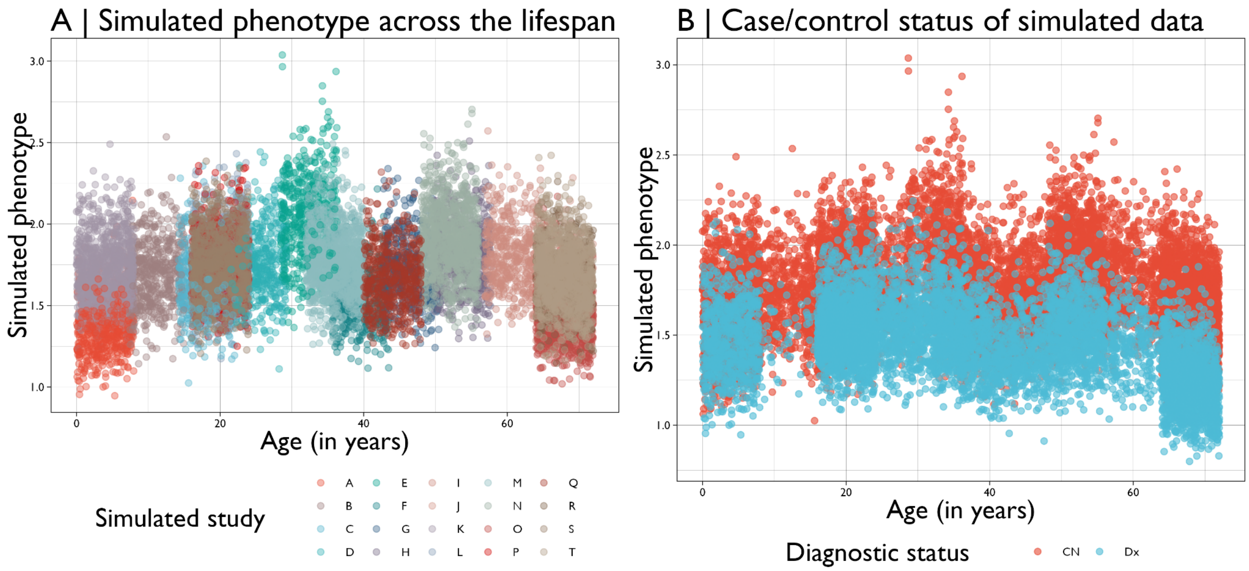


***Fig. S1.4.1. Simulated data for baseline observations.*** *A | Female and male healthy controls (CN) coloured according to 20 simulated studies, highlighting the coverage of the lifespan and the within- and between-study variability. These simulated observations (N=13,500) were used to estimate lifespan curves with GAMLSS in order to motivate the application to real data. B | Healthy controls (CN) and diagnosed (Dx) individuals from each study (black and red respectively) (n=20,250). This simulation posits a diverging lifespan trajectory for Dx individuals, such that at the start of the lifespan CN and Dx overlap but gradually separate, which is induced by using different true age-related quadratics. The specific functional form of the CN and Dx curves are* $((0.4 - x) * (0.5 - x) + 1.8)$ *and* $((0.35 - x) * (0.3 - x) + 1.55)$ *respectively. (values were scaled for computational stability and visualisation purposes).*

The inclusion of individual-level random-effects within the simulation is necessary to induce a dependence between longitudinal observations. While the analysis shown in **Fig. S1.4.1** only uses baseline observations, **Fig. S1.4.2** illustrates the longitudinal follow-up for a subset of individuals across five of the twenty simulated studies for CN and Dx individuals to assess the capacity to model longitudinal trajectories. The simulated dataset also mimics the real-world data with an uneven coverage of the lifespan, as shown in comparing **Fig. S1.4.3** to **Fig. 1A**.

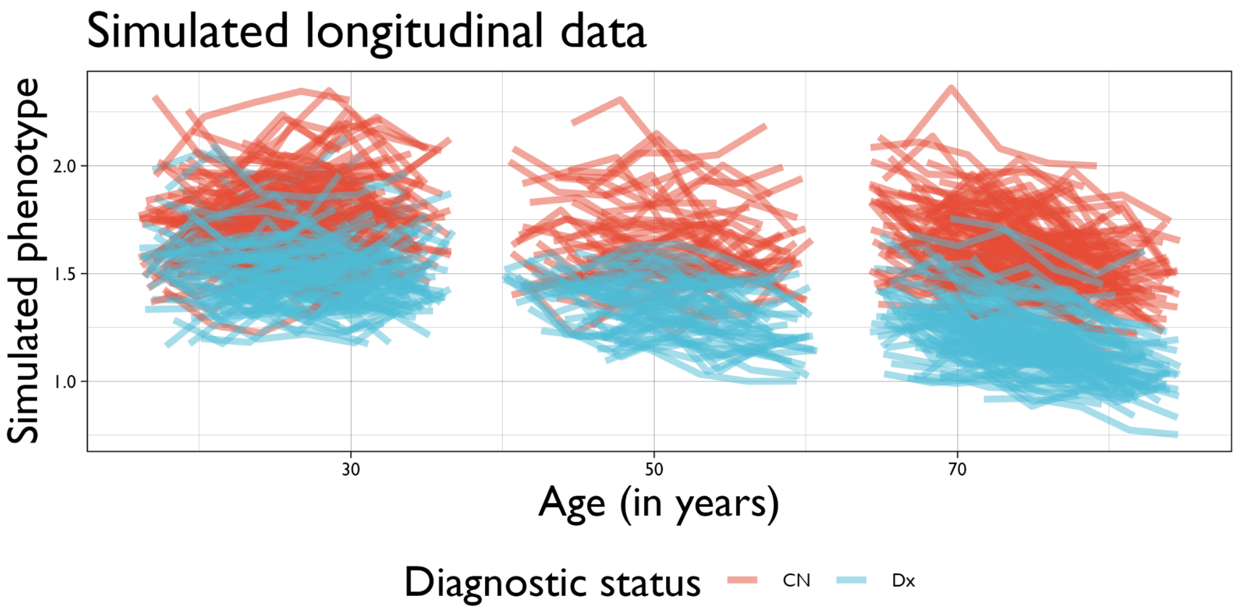


***Fig. S1.4.2. Simulated data for longitudinal observations.*** *An illustrative sample of individuals (250 from among 1,500 for clarity) with longitudinal follow-up within the simulated data coloured by CN (black) or cases (Dx: red). Within the simulation individuals have relatively stable longitudinal trajectories relative to the between person variation, implying longitudinal centiles will be relatively stable for both CN and Dx.*

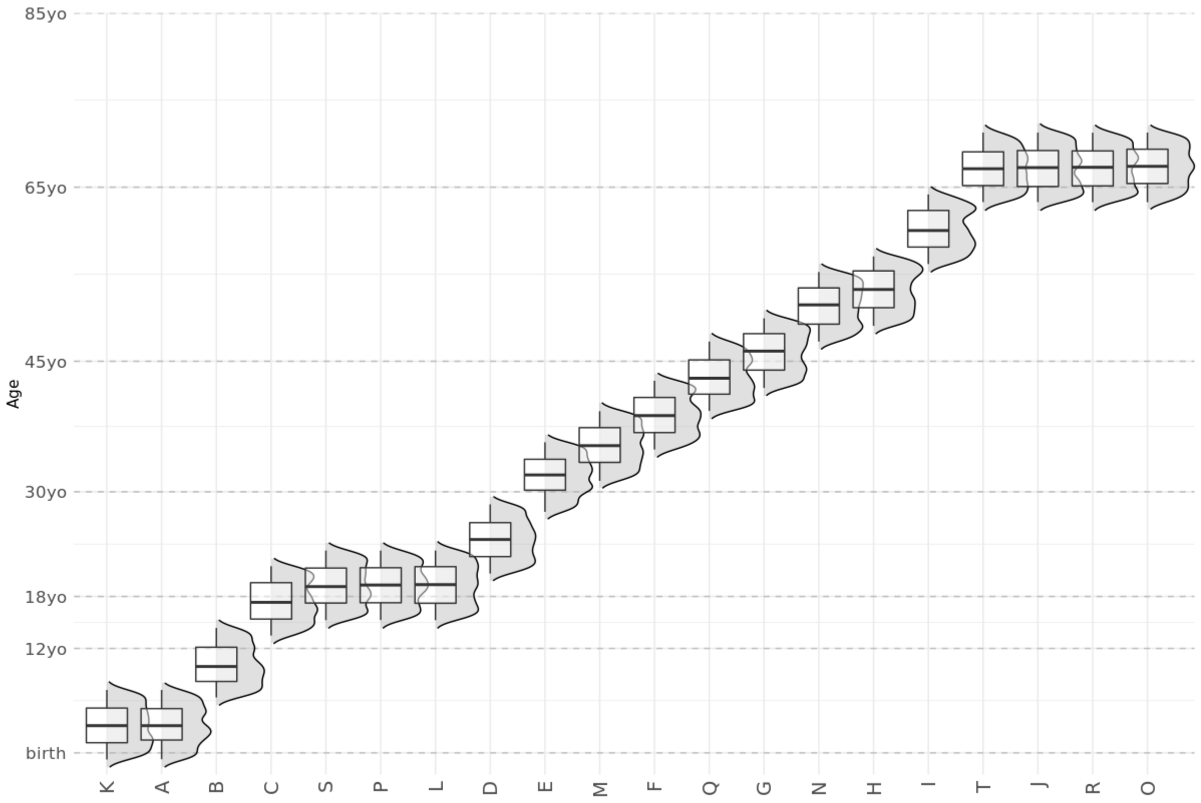


***Fig. S1.4.3. Box-violin plots show age distributions (log-scaled) of twenty simulated studies.*** *The design of the simulation mimics the structure of the observed datasets, with some periods of the lifespan*

*being represented by multiple studies, for example adolescence (studies C,S, P and L), while other periods have sparser coverage with fewer studies.*

#### 1.5 Centile normalisation

The GAMLSS framework allows us to fit complex outcome distributions by parameterising a distribution into multiple components. Each component is then modelled as a regression (with appropriate link functions to ensure valid parameters, for example exponential links for sigma to ensure non-negative sigma values if it corresponds to a variance). As described in the **Online Methods**, we may consider the outcome, Y, to follow a distribution, $F$, parameterised by up to four parameters, $(\mu, \sigma, \nu, \tau)$, with each component itself a regression on potential covariates. Importantly, the regression equations are not required to use the same set of covariates.

Model specification procedures indicated including a study random-effect. Specifically, within the component regressions there is a random-intercept per study. If there are multiple components with random-effects, they are uncorrelated between components. Hence, there will be random-effect estimates, z, drawn from a component specific random-effect distribution, Z (within GAMLSS random effects follow a normal distribution). Namely,

$$\begin{aligned} Y &\sim \mathcal{F}(\mu, \sigma, \nu, \tau) \\ g_\mu(\mu) &= X_\mu \beta_\mu + Z_\mu \gamma_\mu + \sum_i s_{\mu,i}(x_i) \\ g_\sigma(\sigma) &= X_\sigma \beta_\sigma + Z_\sigma \gamma_\sigma + \sum_i s_{\sigma,i}(x_i) \\ g_\nu(\nu) &= X_\nu \beta_\nu + Z_\nu \gamma_\nu + \sum_i s_{\nu,i}(x_i) \\ g_\tau(\tau) &= X_\tau \beta_\tau + Z_\tau \gamma_\tau + \sum_i s_{\tau,i}(x_i) \end{aligned} \tag{1.5.1}$$

where each component is linked to a linear equation through a link-function, $g.()$, and each component equation may include three terms: typical covariates and coefficients, $X.$and $\beta.$; random-effects, $\gamma.$ (which may include covariates, $Z.$); and non-parametric smoothing functions, $s_{\cdot,i}$ applied to the $i^{\text{th}}$ covariate (GAM aspect of GAMLSS). The nature of the outcome distribution, i.e., the generalised gamma distribution, determines the appropriate link-functions and which components are used.

The GAMLSS framework uses the likelihood of the observed data under the above model to estimate the coefficients of each component's regression equation. This formulation implies that we can obtain the centile of an observation using the cumulative density function (CDF). Let f be the probability density function (PDF) of the distribution $F$, $F$ be the CDF, and $F^{-1}$ be the inverse-CDF, such that

$$F(y) = \int_{-\infty}^{y} f(v)dv \text{ such that } 0 \leq F(y) \leq 1 \tag{1.5.2}$$
$$F^{-1}(q) = inf\ \{y: F(y) \geq q\}.$$

Within GAMLSS all outcome distributions are univariate, meaning the CDF and inverse-CDF are well defined, and we can obtain the centile, $q$(lying within the range zero to one), as

$$q = F(y|\mu(X,Z), \sigma(X,Z), \nu(X,Z), \tau(X,Z)). \tag{1.5.3}$$

Using the observed centile, which is dependent on the study random-effects, we can further obtain reference normalised values, $w$, by finding the matching centile in the reference distribution, that is without any random effects (i.e., $Z = 0$):

$$w = F^{-1}(q|\mu(X), \sigma(X), \nu(X), \tau(X)). \tag{1.5.4}$$

The reference normalised values are the matching centiles on the reference outcome distribution.

These normalised values, $w$, are on the same scale as the original values, $y$, having been corrected for the study-specific effects: namely, the $\mu$-component and $\sigma$-component study random-effects. However, these corrections are only appropriate for scoring scans that were included in the reference dataset, i.e., healthy controls, and normalised values are therefore not useful for scoring scans from cases of clinical disorder or for out-of-sample scoring of “new” scans. We have included a brief consideration of normalised values, $w$, for completeness and because they may be more interpretable than centile scores in some contexts, since they are scaled to the same units as the scored phenotypes. However, for most applications (including the case-control comparisons and out-of-sample analyses reported in this paper), we therefore strongly recommend the use of centiles.

#### 1.6 Centile Mahalanobis distance

To create an integrated measure of normative deviation across all centile scores we computed a Mahalanobis distance[28] in the 4-dimensional feature space relative to the normative mean across those phenotypes. This centile Mahalanobis distance (CMD), $D_M$, can be formalised as follows:

$$D_M(x) = \sqrt{(x-\mu)^T S^{-1} (x-\mu)} \tag{1.6.1}$$

where $x$ denotes the set of observations across multiple phenotypes, μ denotes the mean across those observations, and *S* denotes the covariance matrix across both. The squared Mahalanobis distance is also equivalent to the sum of squares of all non-zero standardised principal components scores (as illustrated in **Fig. 4B**). As such, CMD provides an indication of the distance of an individual from the centre of the normative multi-dimensional (multi-phenotype) space, taking into account the potential correlated structure of the dimensions (and thereby being arguably less sensitive to outliers along a single dimension than other possible distance metrics). The scale-invariant nature of CMD also makes it generalisable to centile scores on additional MRI phenotypes as they are included in the future.

#### 1.7 Longitudinal centiles

Using the CDF (as defined in **Eq1.5.3**), we can obtain the centile for any observation. Specifically, if an individual has multiple observations we can obtain the longitudinal centiles. It is important to stress that the fitted model only uses cross-sectional data and thus is not accounting for within-subject correlation. This means that the model is not optimized for individual longitudinal predictions. However, under our hypothesis that for nominal subjects within-subject covariance is dominated by between-subject covariance (in other words, there is more variation from person to person, but repeated measures within a person will remain fairly consistent), then an individual's centile should remain consistent over time. In effect, each individual will sit off the reference curve, but will track the curve over time. Whereas, for individuals that experience significant clinical changes, for example the onset of a disorder that affects neuroanatomy, will change their centiled position relative to the reference curve. This is the foundation of the use of growth curve charts in clinical practice, where we track an individual over time with two aspects of interest: 1) on initial assessment whether the individual is in an extreme (top 1%, 5%, etc.), and 2) given an initial assessment whether an individual 'jumps' deciles of the growth chart or remains consistent in their decile (which can also be formalised as thrive lines in the case of consistent longitudinal data [29]).

Hence, for individual j from study i, with observations at $t_1, t_2, \cdots, t_m$, we can obtain the centiles:

$$q_{ij1}, q_{ij2}, \cdots, q_{ijm}. \qquad (1.7.1)$$

For healthy controls these centiles will be an appropriate model. For other individuals (across the range of diseases and conditions in all our studies) these centiles may be biased if the cohort has a systematic deviation from the normative reference or follows a systematically different trajectory. Hence, we have two dimensions of interest for an individual's longitudinal centiles: 1) the distribution of centiles across classes of individuals, and 2) the stability of centiles within individuals within a class. For the first dimension, if diagnosed cases are all substantially lower on the outcome, then their normative control-referenced centiles will all be low. More generally, they will not be uniform across the range zero to one, as they are for the healthy controls comprising the reference dataset. For the second dimension, we may consider a summary of the within-subject variation.

Thus, comparing longitudinal centiles, with varying numbers of observations per individual, is approached via a univariate summary statistic. A univariate summary for variation across observations will assess the stability of the centiles within an individual. The summary must be valid for two or more observations, the minimal longitudinal follow-up period, and be comparable across individuals. The range, i.e., $max(q_{ij1}, q_{ij2}, \cdots, q_{ijm}) - min(q_{ij1}, q_{ij2}, \cdots, q_{ijm})$, would be well defined for two or more observations; however, the range is susceptible to outliers and statistically unstable under small samples. Instead, the interquartile range (IQR) acts as a robust equivalent of the range (in the same way that the trimmed mean is a robust version of the mean). However, unlike the trimmed mean which requires a “large enough” sample, the IQR is valid for small samples. Given the variable number of longitudinal data-points available for different participants, we chose to use a measure that was consistent for participants that only had 2 observations as well as for participants with more than 2 observations. Unfortunately, there is not a single definition of the IQR (there are 9 different definitions available within GNU R), and some versions are not defined for two observations. We estimated IQR as a continuous value by linear interpolation (within GNU R the default version of IQR, type 7), which is well defined for two (or more) observations.

Specifically, we summarise all individuals with longitudinal observations by,

$$IQR(\, q_{ij1}, q_{ij2}, \cdots, q_{ijm} \,). \qquad (1.7.2)$$

As a summary measure this gives us a single value per subject and incorporates both dimensions of interest. However, there are some substantial aspects that are obscured, for example, the temporal nature of the repeated measures. Note that,

$$IQR(\, q_{ij1}, q_{ij2}, q_{ij3} \,) \equiv IQR(\, q_{ij3}, q_{ij1}, q_{ij2} \,). \qquad (1.7.3)$$

That is, the IQR is indifferent to the temporal order of the centiles. This will obscure certain types of temporal divergence from the reference curve that are likely to be of interest in future work. It does, however, guard against individual outliers over multiple time-points. IQR assessed on the simulated data described above confirms this (**Fig. S1.7.1-1.7.2**).

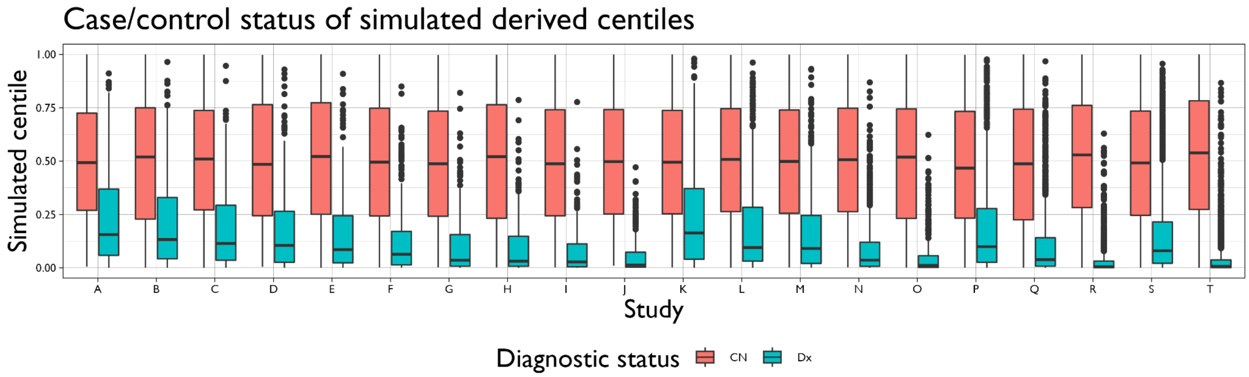


***Fig. S1.7.1. Comparing baseline centiles between healthy controls (CN) and diagnosed cases (Dx) in simulated data.*** *The CN and Dx simulations follow two distinct lifespan trajectories, both quadratic in shape and starting slightly offset in early life, both peaking in mid-life with growing divergence, and fully diverging in later life. The analysis of simulated data is formally equivalent to the analysis of observational data and the GAMLSS model is fitted to only simulated data of CN baseline scans. The figure shows the distribution of baseline centile scores across the twenty simulated studies (spanning different ranges of the lifespan, in four groups: A-J, K-O, P-R, S-T; see* ***Fig. S1.4.3****). We note that the Dx centiles are not uniformly*

*distributed between zero and one, but are skewed to the lower end of the distribution as expected from the simulation scenario: namely, that the Dx simulations are always below the fitted CN lifespan trajectory. Further, the skewness of the Dx centiles increases later in the lifespan (compare study J to study A). Conversely, the CN centile distributions are uniformly distributed from zero to one as expected.*

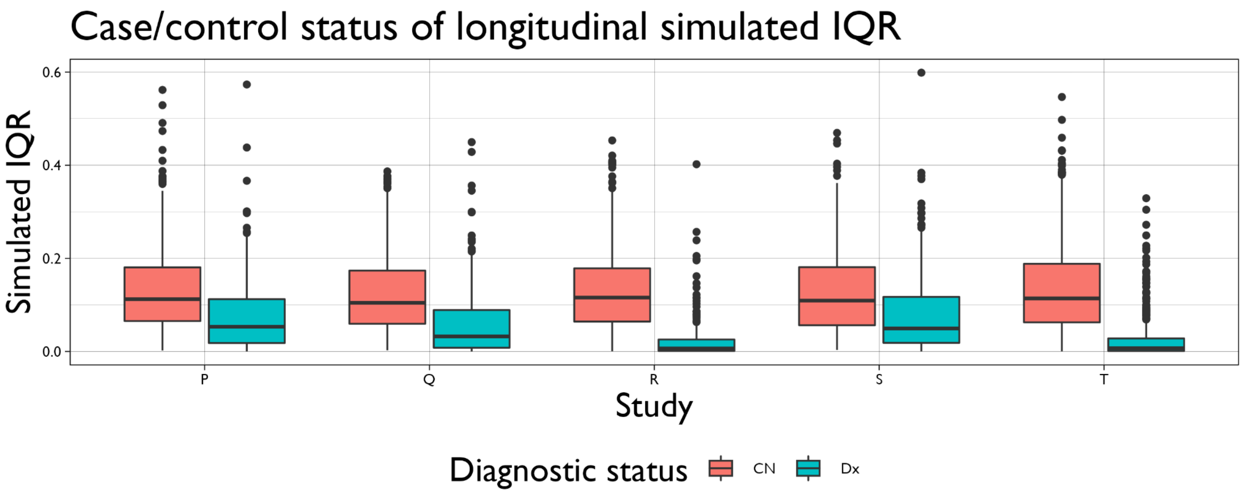


***Fig. S1.7.2. Comparison of interquartile range (IQR) of observed longitudinal centiles between healthy controls (CN) and cases (Dx) in simulated data.*** *Simulations P-T included longitudinal follow-up data for CN and Dx (non-CN) individuals. As described, the simulated analysis model (fit to the CN baseline observations) is used to derive centile scores for all observations, Dx and longitudinal. Taking the IQR as a summary statistic of within-subject variability of longitudinal centiles, the boxplots for CN simulations highlight the stability of longitudinal centiles over follow-up. For the Dx simulations, we see an echo of the effect from* ***Fig. S1.7.1****; importantly the collapse of IQR variability towards zero does not imply the Dx centiles are more stable per se but rather the Dx status might coincide with more limited variability by being confined to the tail end of the distribution. This plot confirms that cross-sectional brain charts can be used to benchmark longitudinal measurements.*

#### 1.8 Out-of-sample estimation

The GAMLSS framework uses iterative maximum likelihood to obtain estimates of the component regression coefficients (and any associated random-effects). The theoretical basis of a random-effect concerns the concept of an infinite population of possible random-effects, in our case all possible studies. To obtain observed centiles we require the study-specific random-effects, which are obtained during the model fit. For a new study, not included within the fitting process, we do not have an estimate of the random-effects and so cannot obtain observed centiles (or normalised outcome values).

However, estimates for novel study random-effects can be obtained conditional on the fitted fixed-effects. Specifically, for a novel study the only unknown (in terms of model estimators) are the random-effects. Using the likelihood as defined by the main model fit, we can obtain maximum likelihood estimates (MLEs) of the random-effects conditional on the fitted fixed-effects. To assess uncertainty within the novel random-effects, we utilise the bootstrap replications (each with their own fixed-effects estimates) to obtain uncertainty within the novel estimates.

It is worth noting that these novel random-effect estimates are conditional on the fitted model. This approach avoids having to refit the entire model. In fact these calculations are computationally trivial. However, this approach implicitly assumes that the novel data does not drastically diverge from the current model fit. In other words, the novel data does not substantially change our understanding of the reference lifespan curve, which would thereby invalidate the original model fit. Further, if we refit the model incorporating the novel data directly, we cannot expect the latter estimates to be identical since the novel data will now be affecting the entire fit, including the estimation of fixed-effects and the whole estimation of the random-effects.

Let $D = \{D_1, D_2, D_3, \ldots, D_k\}$ be the combined datasets used to estimate the model parameters, specifically the fixed-effects for each component of the GAMLSS model, $\beta = (\beta_\mu, \beta_\sigma, \beta_\tau, \beta_\nu)$, and the study-specific random-effects for each component, $\gamma = (\gamma_\mu, \gamma_\sigma, \gamma_\nu, \gamma_\tau)$, where each $\gamma$ contains a parameter for each dataset $D_i$ ,i.e., $\gamma_\mu = (\gamma_{\mu,1}, \gamma_{\mu,2}, \gamma_{\mu,3}, \ldots, \gamma_{\mu,k})$.

In symbolic terms, we may consider the set of fixed- and random-effects from our model to be obtained from fitting the GAMLSS model,

$$(\beta_D, \gamma_D) = GAMLSS(D) \qquad (1.8.1)$$

where $\beta_D$ and $\gamma_D$ are the maximum likelihood estimates of the fixed- and random-effects, respectively, from the GAMLSS model conditional on a given dataset, $D$. Note that the GAMLSS model includes specification of the functional form, namely the fractional polynomial specification; however, during OoS estimation the fractional polynomial specification of the GAMLSS model is fixed and hence has been omitted here for clarity.

For a "new" dataset, say $D_m$, we require inference on its study-specific random-effects parameters. However, we condition on the fixed-effects parameters from **Eq 1.8.1**, namely $\beta_D$. We can obtain these estimates from a conditional maximum likelihood estimator (MLE).

$$\gamma_{:,m} = (\gamma_{\mu,m}, \gamma_{\sigma,m}, \gamma_{\nu,m}, \gamma_{\tau,m}) = MLE\ (D_m \mid \beta_D). \qquad (1.8.2)$$

Combining the OoS estimate of study-specific random-effects with the fixed-effects, we can derive centile scores for the new study in the same way as centile scores are calculated for studies that were included in the reference dataset.

#### 2. Quality control

While developmental and ageing trajectories of cerebrum tissue volumes were expected to be relatively robust to data quality issues [30], controlling the quality of data is an important step in any neuroimaging analysis pipeline. We conducted several complementary analyses to evaluate the robustness of our procedures and results to variable image quality defined by the Euler Index (EI)[31] and other quality control (QC) metrics.

#### 2.1 Euler index filtering

First, we examined the effect of image quality on estimated brain phenotypes and GAMLSS model parameterisation using EI, an automated, quantitative measure of data quality in scans processed by FreeSurfer (~95% of the reference dataset)[31,32]. The EI metric we used was defined as the sum across hemispheres of the number of surface 'holes' or topological defects in the cortical surface reconstruction prior to a topological correction performed as part of the FreeSurfer pipeline (usually due to errors in white matter segmentation). Although cerebrum tissue volumes are expected to be less sensitive to cortical surface topology, compared to surface-based measures such as indices of cortical folding (see **SI18 "Data processing**"), EI has previously been used as a measure of the quality of "raw", unprocessed scans [31]. Thus, for the large majority of studies where EI was available (N=101,708 total scans on N=82,023 unique subjects), we assessed the impact on reference models of excluding high-magnitude EI scans. Given that no single EI threshold is expected to be generalizable across studies[31] (**Fig. S2.1.2)**, in this sensitivity analysis we excluded scans that had EI magnitude greater than 2 median absolute deviations from the primary study-specific median EI. This QC threshold, which is adaptive to the variable quality of scans between primary studies, excluded approximately 9-10% of scans from the original dataset. However, as can be seen in **Fig. S2.1.3,** the resulting model parameters were highly correlated with parameters estimated from the reference dataset without applying any EI-based QC threshold. The developmental trajectories estimated for all 4 cerebrum tissue volumes were highly correlated with their trajectories estimated on the basis of the full dataset (all $R^2 > 0.999$ for parametric [Pearson's] and non-parametric [Spearman's] correlations between EI-filtered versus EI-unfiltered median trajectories and lower (2.5%) and upper (97.5%) centiles). Identical parameterisation of fractional polynomials for each random effect was identified by the same model selection procedure was found in both EI-filtered and EI-unfiltered datasets. Importantly, EI-filtered and unfiltered datasets also showed a high degree of overlap in subsequently estimated model parameters (correlation of study-specific mean (mu) components > 0.99; correlation of study-specific variance (sigma) components > 0.93). Model specification thus appeared to be robust to the presence of the poorer quality data.

In addition, we examined the relationships between image quality measured by EI and individual centile scores of each brain phenotype. Both for the full dataset and the EI-filtered subset of higher quality scans, we found no significant associations between EI and individual centile scores (**Fig. S2.1.1**), nor did we find evidence for a non-linear relationship (quadratic, cubic, logarithmic) between EI and centiles**.**

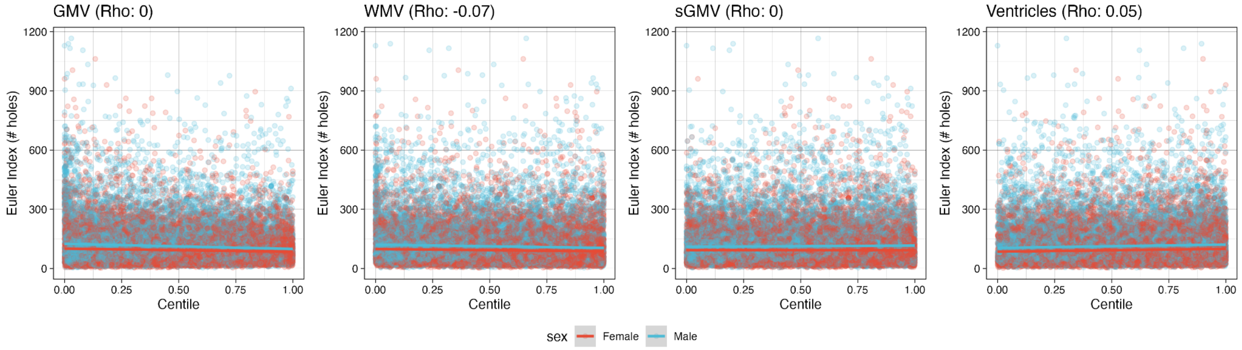

***Fig. S2.1.1 Associations between centile scores and MRI scan quality defined by EI***. *Panel depicts the relation between Euler indices (EI)* [31] *and centile scores for each of 4 cerebrum tissue volumes estimated by GAMLSS. The Spearman correlations between EI and centile scores were negligible (GMV, ρ<0.01; WMV, ρ=-0.07; sGMV, ρ<0.01; Ventricles, ρ=0.05). All linear mixed effect models examining non-linear (quadratic, cubic or logarithmic) relationships between EI and centile scores for each phenotype were P > 0.1.*

To assess whether there were any age-related differences that could influence model estimation, we evaluated the linear effect of age (in years) on EI in healthy controls in the reference dataset used to estimate normative lifespan trajectories. Using linear regression stratified by sex and accounting for study-specific random effects, we found no evidence for an age-related bias in image quality as assessed with EI ($t$ = -1.244, $P$ = 0.213). **Fig. S2.1.2** shows the median and standard deviation of age and EI and highlights the top 10 studies with the highest median EI.

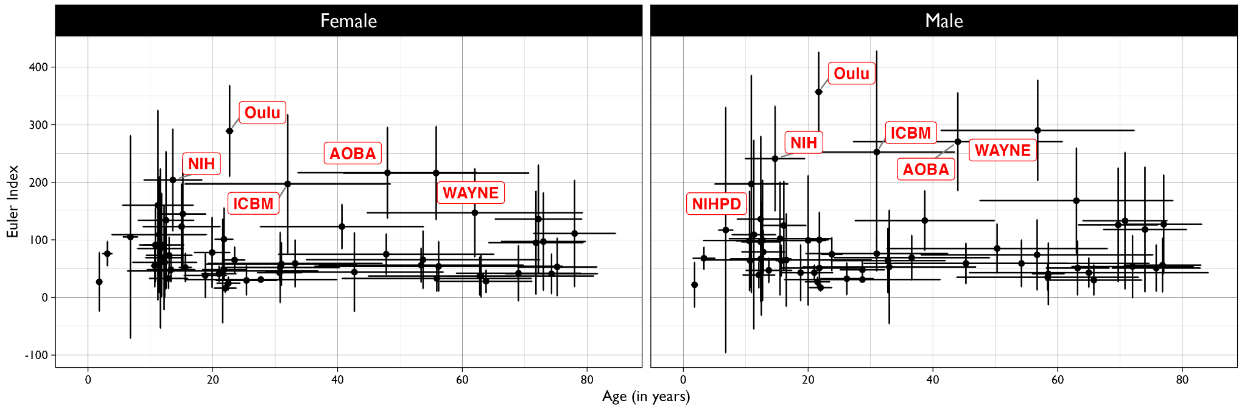


***Fig. S2.1.2 Age-related variation in image quality measured by the Euler index in female (left panel) and male (right panel) control subjects.*** *Median age (in years) and median EI are shown per study with cross-hairs indicating the standard deviations for age and EI per study. In red the top ten studies with the highest median EI are highlighted. There is no significant relationship between image quality and age at scanning.*

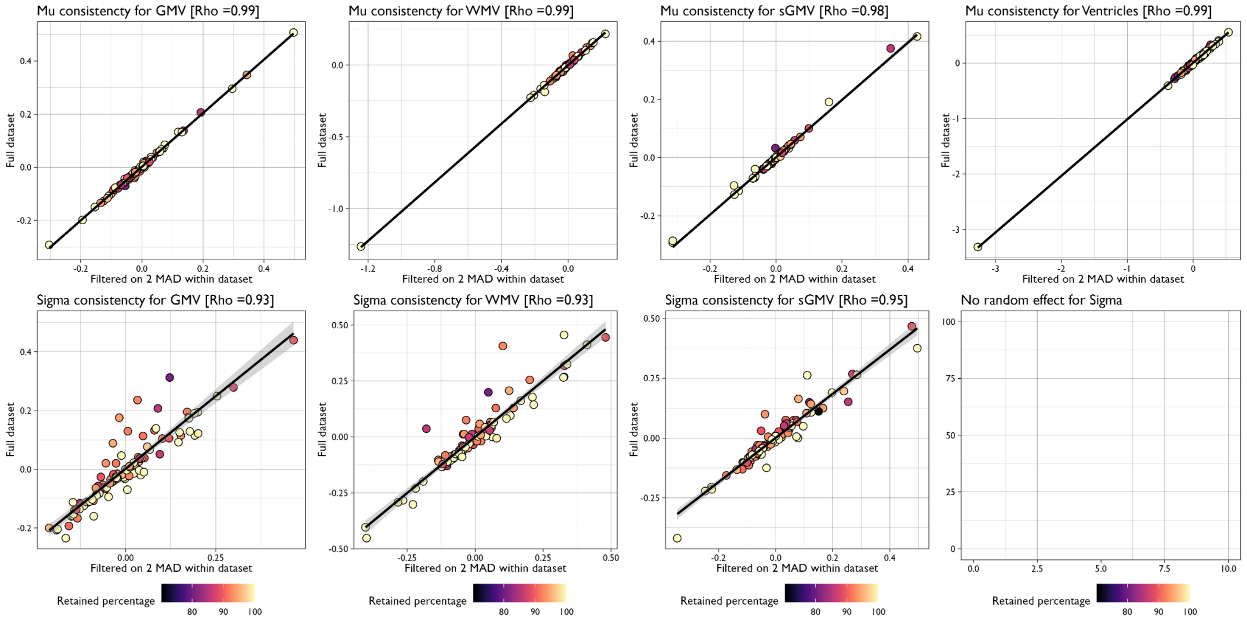


***Figure S2.1.3 Robustness of GAMLSS parameters to quality control by exclusion of scans with EI greater than twice the median absolute deviation (MAD) from the median EI in the corresponding primary study.*** *Scatterplots show the relationships between random effects (mu on the top row and sigma on the bottom row) estimated for each primary study without exclusion of poor quality scans (y-axis) and for each primary study after exclusion of scans with EI > 2 MAD, relative to the primary study's median EI. Colored points indicate the relative percentage of primary studies retained after filtering (darker means for subjects were removed) and Rho values in the titles indicate Spearman's correlations between parameters estimated from the unfiltered and EI-filtered datasets. As with the absolute QC threshold of EI < 217 (**SI 2.1**), the biggest discrepancy in study-specific random effects as a result of excluding poor quality scans was observed for the variance (Sigma) parameters, especially those estimated from the ICBM, HBN and EDSD datasets, which all included a relatively high proportion of excluded scans. We note that EI > 2 MAD filtering removed a lower proportion of data in primary studies where the distribution of EI was skewed towards higher quality/lower EI across the whole dataset (e.g., HCP, ABCD and UKB all have high data quality with low EI, and 2 MAD filtering in these studies only removed around 6-7% of data). In general, random effect parameter estimation was highly robust to adaptive EI thresholding for quality control.*

#### 2.2 Expert visual quality control

Recognising that EI is but one metric of image quality, and mainly based on the capacity of FreeSurfer to correctly process the scans, we also leveraged visually-rated image quality performed for a subset of 9,704 raw scans. These scans were provided by openly available datasets and are marked as having "Manual" quality control in the "Extracted QC" column in **ST1.1**. For each subject a slice stack of images was generated across the three axes, after bias field correction and intensity normalization, so that they were all easily comparable by visual inspection, and subsequently rated on motion corruption and other failure modes (artefacts, missing brain parts etc). Visual inspection then rated each image on the following questions: is the brain fully covered by the scan; is there visible noise (due to aliasing, motion etc.), blurriness, or ringing; is there acceptable tissue contrast and image orientation? Based on these criteria, each raw scan was expertly classified on a 6-point scale as perfect (1), very good (2), good (3), bad (4), very bad (5) or unacceptable (6). Only 3% of scans (N=374) were assigned to the two worst quality categories (5 and 6). Each image was rated by a single rater. We analysed centile scores for each of the 4 cerebrum tissue volumes in each of these 6 classes of visually curated image quality (**Fig. S2.2.1**). Centile scores for all 4 phenotypes were consistent across the top 4 classes of image quality but significantly variable for the minority of scans with very bad or unacceptable image quality. However, when we excluded these scans from re-analysis of this expertly QC'ed dataset, we found that the median trajectories and 95% confidence intervals for all 4 brain phenotypes were very highly correlated between the results of model fitting to all 9,704 scans and the results of fitting to the 9,380 scans assigned to the top 4 quality classes (all $R^2 > 0.999$ for both Pearson's and Spearman's correlations for all 4 phenotypes). Additionally excluding the 4th category for GMV did also not impact the stability of the resulting trajectories ($R^2 > 0.999$ for both Pearson's and Spearman's correlations).

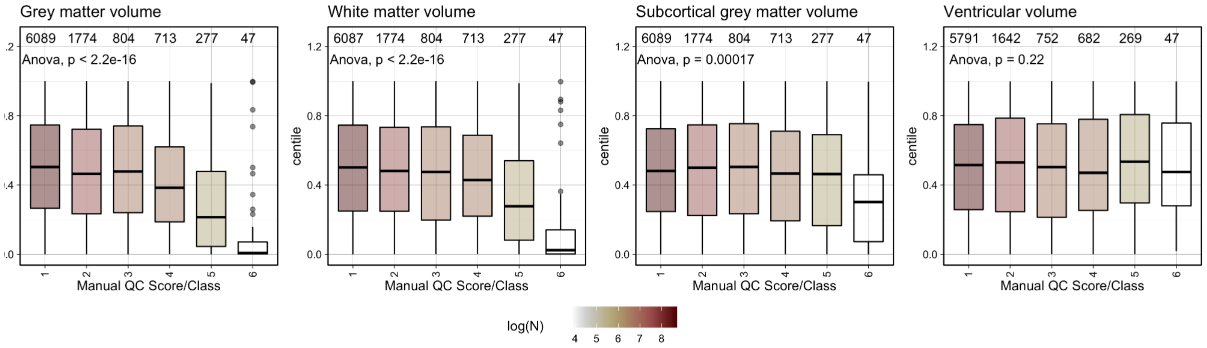


***Fig. S2.2.1. Centile scores for images categorized by expert visual quality assessment of 9,704 unprocessed scans.*** *A small subset (~3%) of the raw data were assigned to the two worst categories of data quality (QC class 5 or 6) and differed significantly from the other QC classes of data in terms of centile scores for cortical grey matter volume, white matter volume, and subcortical grey matter volume. Bars are coloured by log-scaled sample size.*

Using the subset of data with visually-rated image quality ratings, we also performed a limited validation of the use of EI as an automated metric of image quality. As shown in **Fig. S2.2.2**, EI magnitude was strongly associated with manual reviewer ratings, such that lower quality scans had a higher number of surface 'holes' or topological defects in the cortical surface reconstruction prior to a topological correction. Moreover, scans that had EI magnitude greater than 2 median absolute deviations above the primary study-specific median EI (the criterion applied in the sensitivity analysis described above) were significantly lower quality as determined by manual rating (**Fig. S2.2.2**).

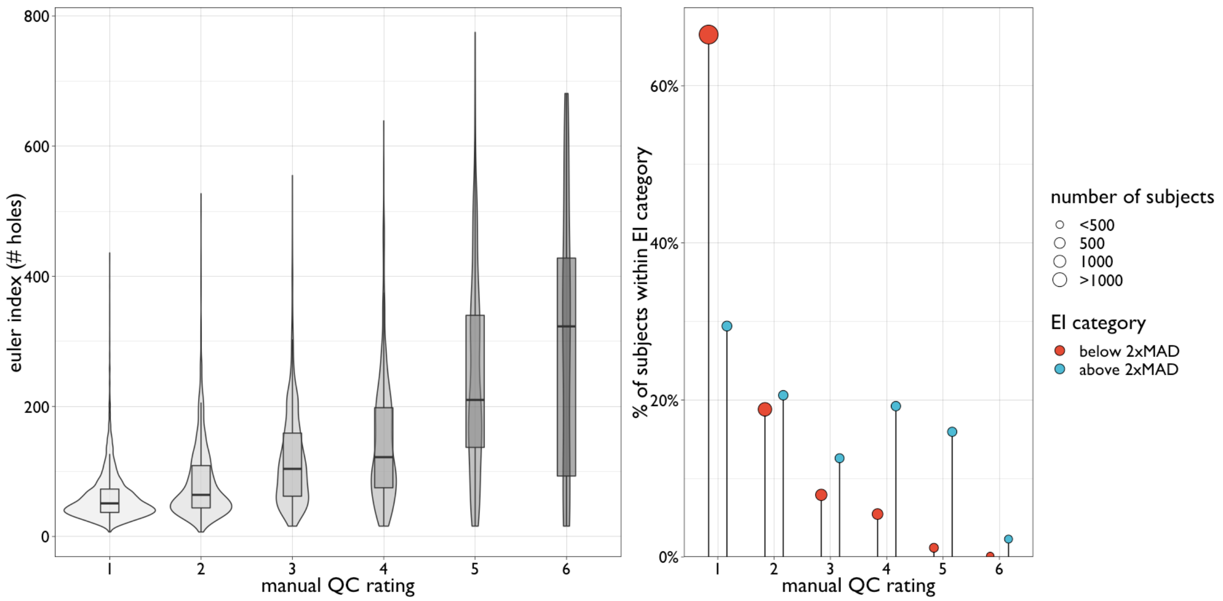


***Fig. S2.2.2. Comparison between manual quality control ratings and Euler index.*** *(Left) Box-violin plots showing the distribution of Euler index (EI; calculated as the number of holes or topological defects in cortical surface reconstruction) as a function of manual quality control (QC) ratings by a single rater (t=56.44, P<2e$^{-16}$). Increasing values for both variables are associated with worse image quality. (Right) Lollipop plot showing differences between EI category, binarised based on 2 median absolute deviations (MAD) above the primary study-specific median EI (t=36.36, P<2e$^{-16}$). Size of dots represent the absolute number of subjects per ordinal QC rating. Total number of subjects with both EI and manual QC ratings, N=9,704.*

For foetal and some other primary studies where MRI data were not reconstructed with FreeSurfer, and the EI was therefore not available, scan quality had previously been assessed by expert visual curation as part of primary study procedures (Table **ST1.1** lists the QC steps for each combination of dataset, sex, site and processing pipeline). We re-analysed data from these studies stratified by their prior QC ratings. For example, the Harvard foetal cohort conducted independent visual inspection of image reconstruction quality and classified each of the images as 'great', 'good' or 'bad'. Only the best two categories were included in analyses. We found no significant difference in centile scores for each of the 4 phenotypes between 'great' and 'good' images (GMV, *P*=0.58; WMV, *P*=0.34; sGMV, *P*=0.14; CSF was not available for these foetal scans).

Similarly, the ABCD study provided expert visual counts of artefacts identified by their inspection of FreeSurfer-processed data. For the ABCD data (N=9,056) included in our reference dataset, the majority of images had been rated as containing zero artefacts; a small subset (<0.5%) of scans had been rated as containing one or more artefacts. As shown in **Fig. S2.2.3**, there was some variability of centile scores in the small number of scans with high artefact scores, but there was no significant group level difference in centile scores for any of the four cerebrum tissue volumes between scans with zero artefacts and scans with one or more artefacts (ANOVA, *P*>0.05).

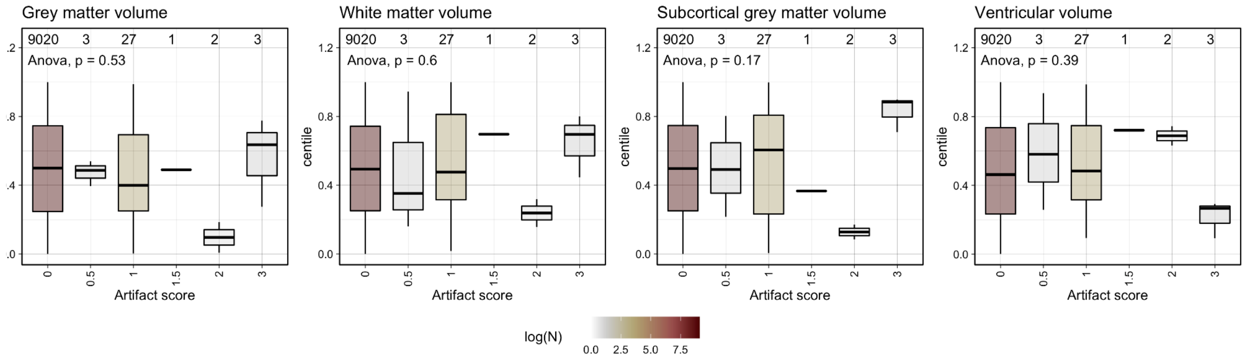


***Fig. S2.2.3. Centile scores for ABCD scans previously assigned artefact scores by expert visual QC.*** *The majority (>99%) of ABCD scans included in the aggregated dataset had zero artefacts; for scans with more than one artefact detected there was some variability in estimated centile scores. Bars are coloured by log-scaled sample size.*

#### 2.3 Image quality and out-of-sample centile scoring

Recognising that image quality would likely be most influential for out-of-sample centile scoring of scans that were not included in the reference dataset, we analysed N=72 scans from an open test-retest dataset [30] which had been quantitatively QC'd (by 5 independent raters using *Braindr* [33]) but had not previously been included in our analysis (https://anisha.pizza/braindr-results/#/). We found that *Braindr* QC scores were not substantially correlated with centile scores for each of the 4 cerebrum tissue volumes (Pearson's r; GMV=0.034, WMV=0.002, sGMV=0.007, Ventricles=0.004). In the same dataset, we did find that prospective motion correction [34] somewhat improved the intra-class correlations of GMV centile scores (which changed from r=0.91 for prospectively uncorrected data to r=0.98 for prospectively corrected data). We note that these beneficial effects of prospective motion correction on test-retest reliability of centile scores derived by OoS analysis using our model are consistent with comparable improvements in test-retest reliability of FreeSurfer-derived phenotypes, as previously reported[30].

#### 2.4 Euler index and neuroimaging phenotypes

To further examine the potential influence of quality control on the quantification of MRI phenotypes, we evaluated the relationship between Euler index (EI) and the four main global tissue volumes (GMV, WMV, sGMV, CSF) within each study with available EI data. We observed high variability in the range of EI within and between primary studies (**Fig. S2.4.1**). However, using linear models to assess the relationship between EI and non-centiled ("raw") tissue volumes for the healthy controls within each primary study (controlling for age and sex), we found that the relationship between EI and tissue volume was generally weak, with only a small subset of primary studies showing significant effects of image quality on MRI phenotypes ($P_{Bonferroni} < 0.05$, corrected for the number of studies of each phenotype). Critically, the sign of this relationship varied across studies and was zero-centred, with the significant effects observed in primary studies with greater sample size (linear mixed effects model with phenotype as a random effect, comparing sample size and -$\log_{10}$(*P*-values) for association with EI: $t = 8.77$, $P = 6e^{-16}$; **Fig. S2.4.2**). We stress that while global measures appear to be relatively robust to variation in image quality, finer-grained imaging phenotypes are likely to be more sensitive to variation in image quality, and the impact of QC must be assessed on a phenotype-specific basis in the future.

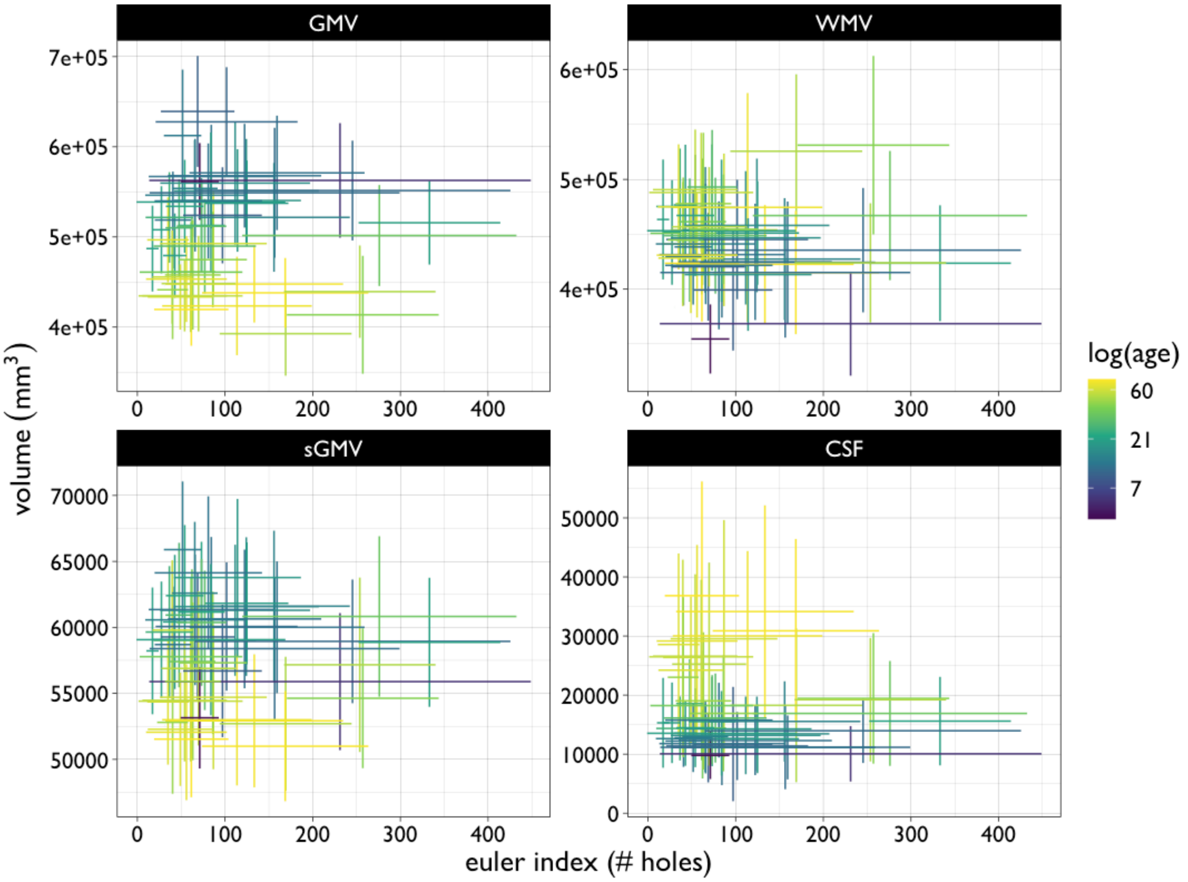


***Fig. S2.4.1. Relationships between the distributions of non-centiled ("raw") cerebrum tissue volumes and Euler index within each primary study.*** *Crosshair plots show the range of values (mean +/- 1 standard deviation) for the Euler Index (EI) and cerebrum tissue volumes for each primary study: clockwise from top left, grey matter volume (GMV), white matter volume (WMV), ventricular cerebrospinal fluid volume (CSF) and subcortical grey matter volume (sGMV). The colour scale represents median log age of participants in each primary study.*

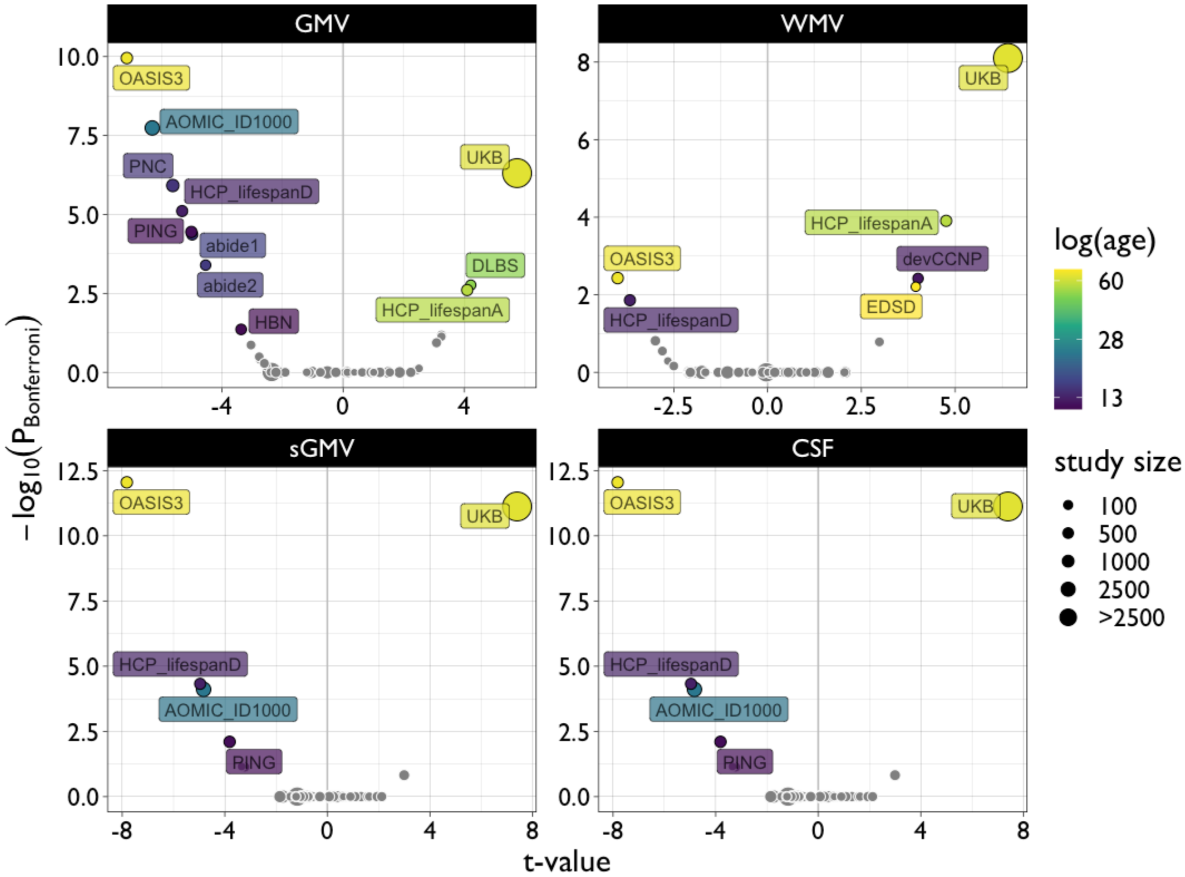


***Fig. S2.4.2. Model statistics examining the relationships between non-centiled ("raw") cerebrum tissue volumes and the Euler index within each primary study.*** *Volcano plots show the t-statistics (x-axis) versus negative log-scaled Bonferroni corrected P-values (y-axis) estimated from linear models of the relationship between Euler Index (EI) and cerebrum tissue volumes: clockwise from top left, grey matter volume (GMV), white matter volume (WMV), ventricular cerebrospinal fluid volume (CSF) and subcortical grey matter volume (sGMV). Each dot represents a single primary study and is coloured to represent the median log age of participants, and scaled to represent the sample size, in a study where there was a significant relationship between cerebrum tissue volume and EI ($P_{Bonferroni} < 0.05$). It is clear that the sign of association between EI and volumetrics was inconsistent between primary studies and the association tended to be significant for primary studies with larger sample sizes.*

In short, we have demonstrated by multiple complementary QC studies that our principal results, and additional out-of-sample results for new data not previously analysed, are remarkably robust to image quality across a range of assessments. We conclude that our results are not confounded by uncontrolled image quality issues; but proper QC procedures should, of course, be implemented on all scans before they are submitted for OoS centile scoring on the basis of our model and aggregated reference dataset. In the absence of a single gold standard for automated assessment of imaging data quality, we strongly recommend using a combination of approaches to determine inclusion/exclusion of MRI data for brain charting. In future, as these methods may be extended to more fine-grained structural MRI phenotypes that are likely to be more sensitive to variation in image quality, and/or to benchmark phenotypes measured in fMRI or more innovative modalities of MRI data more likely to be measured in small samples (N<100), we should be prepared for GAMLSS modelling to be significantly less robust to image quality in comparison to the case of global MRI phenotypes, like cerebrum tissue volumes. The importance of rigorous quality control therefore remains paramount.

### Sensitivity Analyses

#### 3. Model evaluation

##### 3.1 Model diagnostics

In addition to simulation, leave-one-study-out (LOSO) jacknife, bootstrap and validation analyses outlined below, we also utilised more traditional ways to assess model goodness of fit (e.g., inspecting the model residuals)[7,35]. Detrended transformed Owen's plots[36] of the ordered centile residuals clearly showed that the confidence intervals crossed the zero line, indicating normally distributed residuals (**Fig. S3.1.1**). Detrended transformed Owen's plots (DTOPs) are an alternative visual approach to assessing the adequacy of a fitted distribution, derived from a non-parametric approach to the data that uses the empirical samples to derive uncertainty intervals. DTOPs have the slight advantage over the traditional Q-Q (quantile-quantile) plots of being more flexible in relation to the form of the distribution and thus provide a way to compare goodness-of-fit across different distributions. Q-Q plots for GAMLSS fits are derived using transformations of the residuals, from the uniform 0–1 scale to the more familiar normal (Gaussian) distribution, hence they are based on a parametric approach. Neither approach alone is definitive for assessing GAMLSS fits, and Stasinopoulus[7] recommends a variety of approaches including both Q-Q plots and DTOPs.

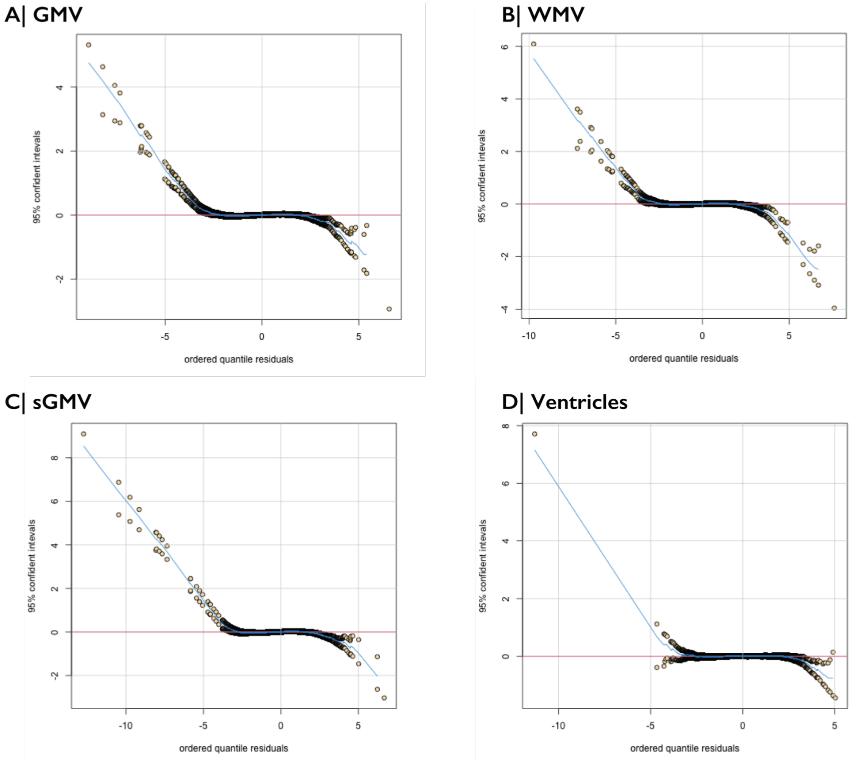


***Fig. S3.1.1. Detrended transformed Owen's plots of model residuals.*** *Visual inspection indicates that model residuals were normally distributed and supports the adequacy of the fitted generalised gamma distributions.*

Further evaluation of the model fits using more traditional QQ-plots (**Fig. S3.1.2**) and residual distribution measures, such as skewness, also showed that residuals were normally distributed and highly correlated[37] with model-predicted normality: GMV (skewness=-0.025, kurtosis=3.69, Filliben correlation coefficient=0.99), WMV (skewness=0.005, kurtosis=3.60, Filliben correlation coefficient=0.99), sGMV (skewness=-0.011, kurtosis=4.29, Filliben correlation coefficient=0.99) and Ventricular CSF (skewness=0.008, kurtosis=3.37, Filliben correlation coefficient=0.99).

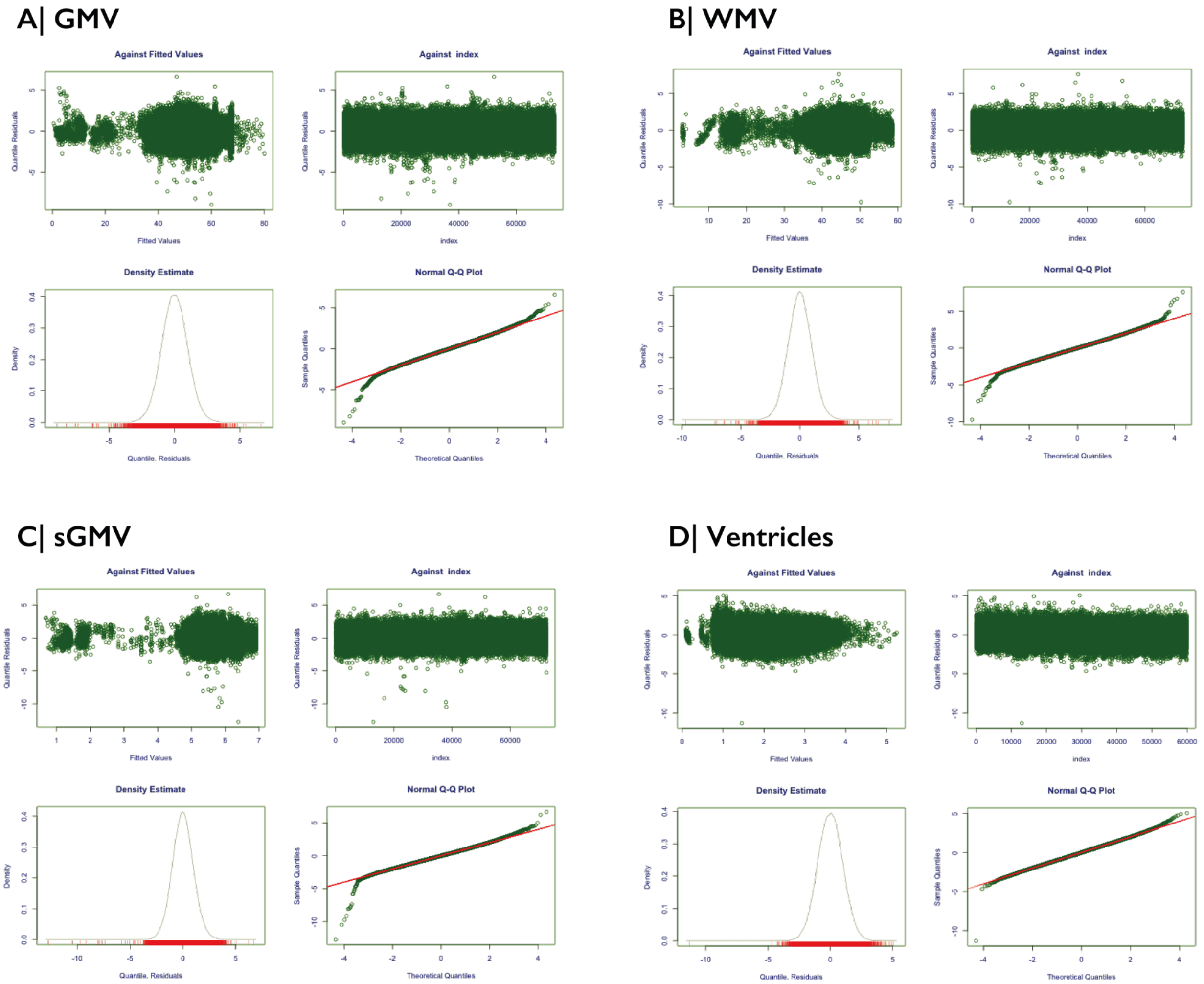


***Fig. S3.1.2. Model fit diagnostic Q-Q plots.*** *Visual inspection indicates that model residuals were normally distributed and supports the adequacy of the fitted distributions.*

#### 3.2 Model sensitivity analyses

With the GAMLSS implementation outlined above (**SI1**), we optimised the choice of model parameters including the outcome distribution and choice of fixed and random effects. In addition, GAMLSS provides automated parameter optimisation to obtain the best fit given the included data, covariates and random effects. Nonetheless we performed several sensitivity analyses to test the robustness and reliability of the optimised GAMLSS models.

##### 3.2.1 Leave-one-study-out

Although the current sample is the largest structural neuroimaging sample reported to date, a large proportion of this data is derived from two population-representative cohorts: the UK BioBank[38] and the ABCD study[39]. To test whether our model's reliability was skewed toward any particular study, we performed a leave-one-study-out (LOSO) analysis. Specifically, we iteratively subsetted our dataset, leaving out one study, re-estimated all model parameters and extracted the fitted trajectories. Given that these models were each derived from different datasets it was not possible to conduct a quantitative comparison of the models in terms of their Bayesian Information Criteria[27] as we did when evaluating the optimal underlying distribution (**SI 1.1 and 1.3**). Instead, we compared the resulting fits for consistency by computing a LOSO confidence interval based on the standard deviation across all LOSO iterations; see **Fig S3.2.1**.

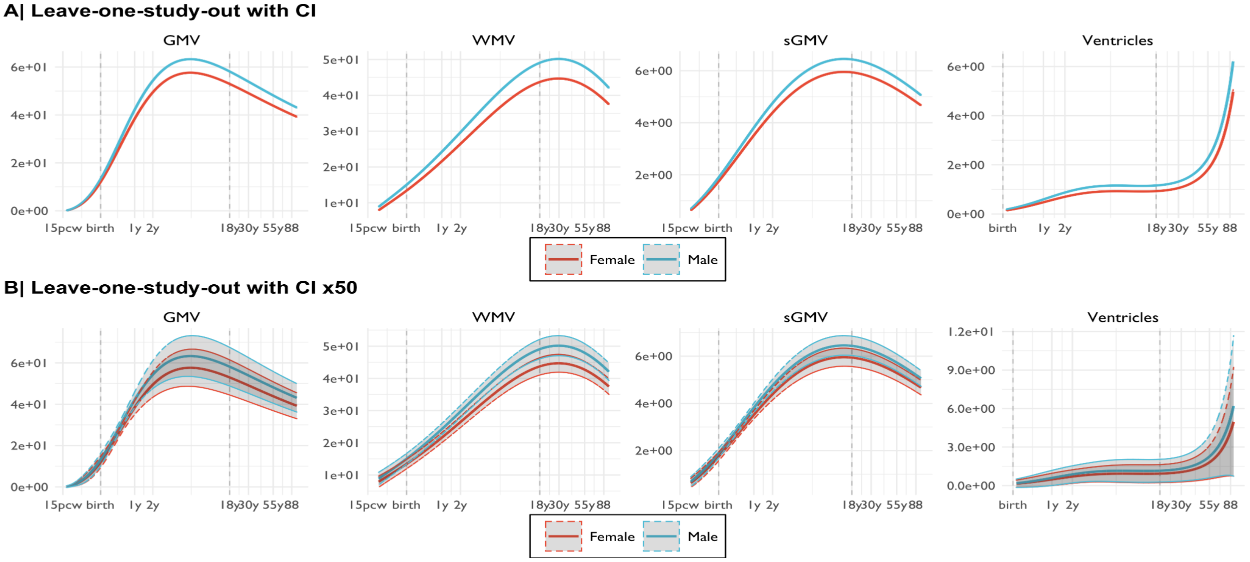


***Fig. S3.2.1. Leave-one-study-out (LOSO) analyses of normative trajectories for cerebrum tissue volumes.*** *A | Confidence intervals (representing the 95% confidence intervals) were computed from the mean and standard deviation of normative trajectories repeatedly estimated after leaving out each primary study in turn: from left to right, grey matter volume (GMV), white matter volume (WMV), subcortical grey matter volume (sGMV) and ventricular CSF volume (Ventricles). B | The same data are shown with the 95% confidence intervals magnified by a factor of 50 to enhance their visibility.*

##### 3.2.2 Bootstrap analysis

To determine reliability and stability of our GAMLSS fitted trajectories, and to obtain confidence intervals on all parameter estimates obtained from the GAMLSS fitting procedure as described above, we ran 1,000 bootstrap iterations with stratified sampling with replacement. The bootstrap replicates were stratified by study and sex, which maintains the relative proportions of the original datasets. Specifically, our process of random resampling of aggregated data was constrained by the relative size of each study compared to other primary studies, and by the sex ratio of each primary study, so that the bootstrap replicates conserved the same proportionality and sex balance as the observed primary studies. We considered it was important to ensure that the bootstrap resampling was representative of the relative sex proportions within studies because we have chosen to stratify normative trajectories by sex, including it as one of the fixed effects in the GAMLSS model. With regard to constraining bootstrap resampling by primary study, there are two inter-linked considerations: between-study differences in sample size and lifespan coverage. Failing to constrain resampling by study sample size could cause a study to be omitted entirely from a bootstrap replicate, or more typically to have a smaller or greater number of observations, meaning the bootstrap intervals would be incoherent for study-level inference. More importantly, the normative trajectories are derived from studies across the lifespan, but each study only partially covers the lifespan; hence failing to stratify by study age-range could alter the bootstrap distribution and lead to incoherent confidence intervals for the lifespan curves. The foetal and early postnatal periods  of the lifespan would be particularly vulnerable to this effect because relatively few primary studies have covered this age range. Our LOSO analysis showed that the lifespan curves were not in fact unduly affected by the removal of any single study (even large ones, for example ABCD and UK-Biobank); see **Fig. S3.2.2**.

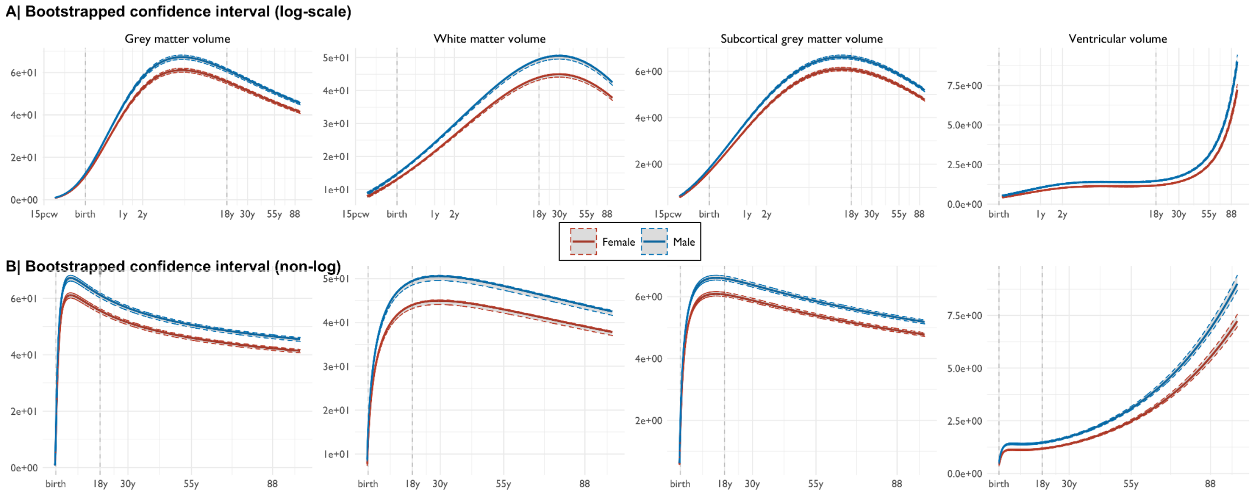


***Fig. S3.2.2. Bootstrap resampling of confidence intervals on normative trajectories for cerebrum tissue volumes.*** *A | 95% confidence intervals (estimated across random bootstrap iterations resampling with replacement) were computed from the mean and standard deviation of normative trajectories (with age on log scale, x-axis) after 1000 iterations of a bootstrapping procedure designed to conserve the relative proportion of primary studies, and the sex balance of each primary study, in each resampling with replacement from the representative dataset: from left to right, grey matter volume (GMV), white matter volume (WMV), subcortical grey matter volume (sGMV) and ventricular CSF volume (Ventricles). B | The same data are shown with age on a natural scale (x-axis).*

##### 3.2.3 Parameter estimates

From our bootstrapping approach, we can also derive confidence intervals for the models' parameter estimates (e.g., the μ and σ terms) for study-specific random effects. Qualitatively we observed very narrow confidence intervals on the estimated μ term, with some smaller sample foetal studies (e.g., CHILD and Harvard foetal cohorts) showing wider intervals, likely commensurate with the smaller sample size and general lack of reference data in that age range (**Fig. S3.2.3**). While there were generally wider confidence intervals on the σ term offsets, across studies all estimated random effect parameters were well contained within their bootstrapped confidence bounds.

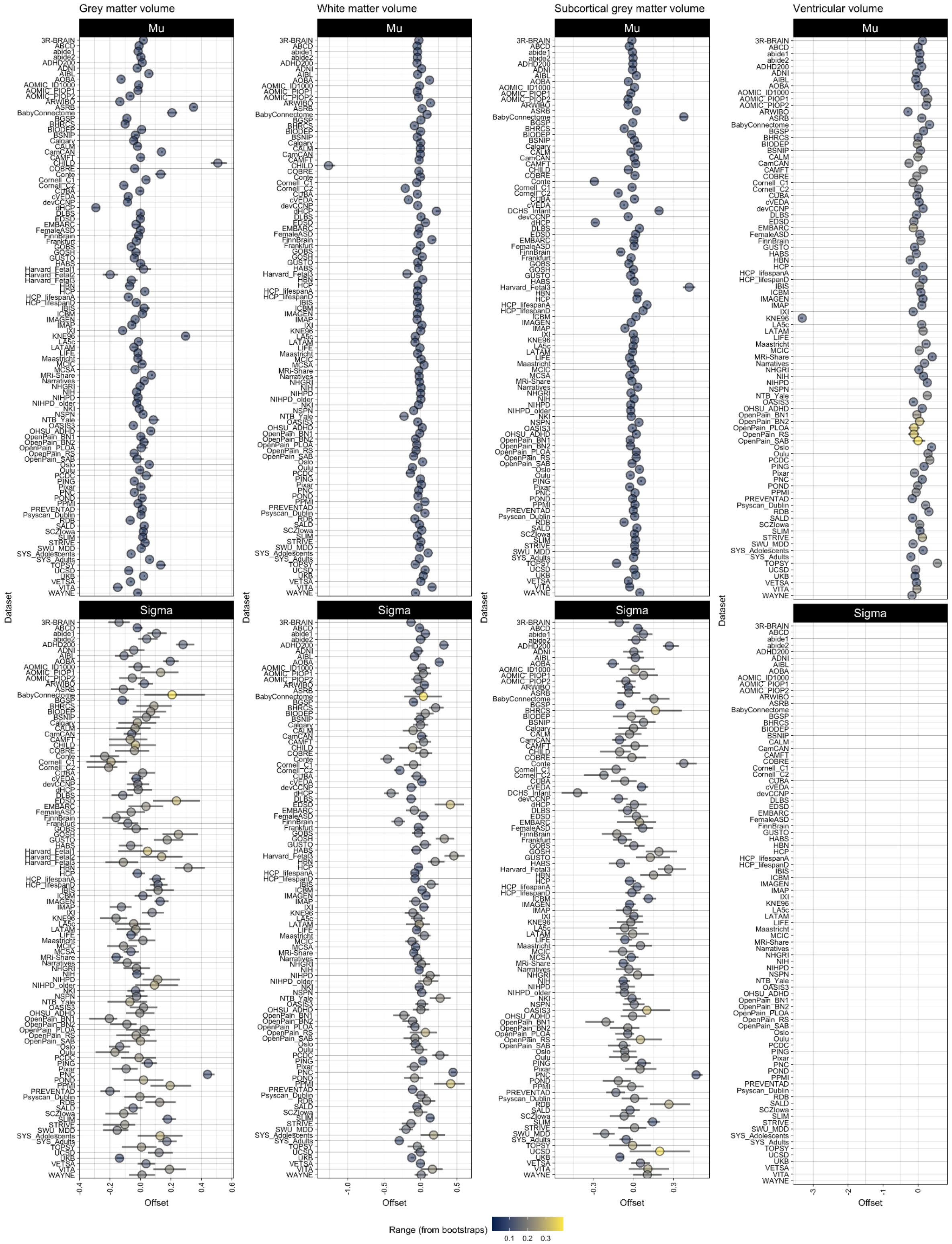


***Fig. S3.2.3.1 Point-range plots of study-specific random effects on the first (Mu) and second (Sigma) moments of the generalised gamma distribution for cerebrum tissue volumes and study-specific random effects on Mu only for ventricular CSF volume.*** *Bootstrapped 95% confidence intervals are shown and point estimates (dots) are coloured by the range of the confidence interval. Where not*

*observable, the confidence intervals are smaller than the size of the dots. There is no Sigma offset for the Ventricular volume as the data-driven process for GAMLSS model specification (**SI 1**) indicated that the best-fitting model did not include a study-specific random effect on the Sigma term.*

We further evaluated the potential impact of various technical and demographic covariates on the random effect parameters estimated by GAMLSS as a measure of each primary study's offset from the normative trajectories of each MRI phenotype. Specifically, we used linear models to estimate the strength of association between random effects (on Mu and Sigma) and median age, standard deviation of age, sample size, scanner manufacturer, and MRI field strength, for each cerebrum tissue volume; see **Figs S3.2.3.2 - S3.2.3.6**. We found only limited evidence for significant effects of any of these covariates on any of these random effect parameters. Other technical covariates, e.g., MRI sequence parameters, were too heterogeneous between primary studies to be assessed for impact on random effects in this way; but full technical specification of all primary studies is detailed in **ST 1.1.**

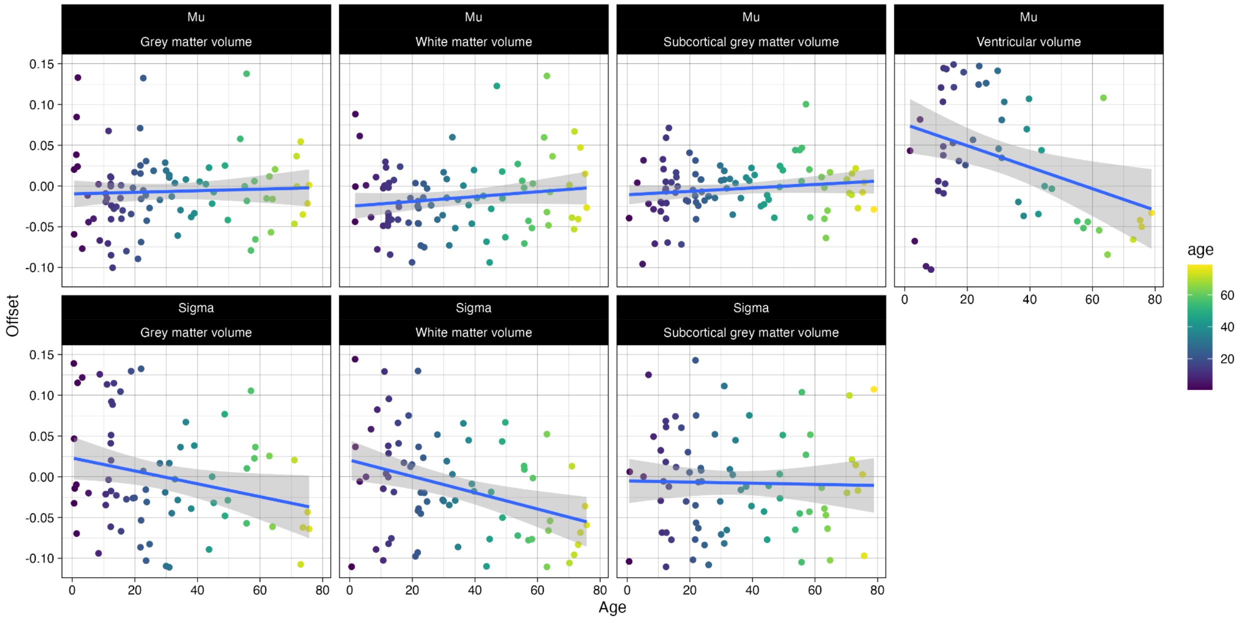


***Fig. S3.2.3.2. Association between median age of participants and random effect parameters estimated by GAMLSS modelling of cerebrum tissue volumes for each primary study.*** *Top row: random effects on Mu (y-axis) are plotted versus median age (x-axis) for each global MRI phenotype, left to right: grey matter volume (GMV), white matter volume (WMV), subcortical grey matter volume (sGMV) and ventricular CSF volume (Ventricles). Fitted lines and confidence intervals indicate the strength of association estimated by linear modeling. Bottom row: random effects on Sigma (y-axis) are plotted versus median age for the same set of global MRI phenotypes (except Ventricular volume for which Sigma was not estimated). There were larger random effects on Mu and Sigma in some of the primary studies of younger participants, as expected by the greater technical and biological variability of studies in childhood. The association between random effects and median age was only significant on the Mu parameter (after FDR correction for multiple comparisons) for ventricular CSF volume ($P_{fdr}$ = 0.007, $R^2$ = 0.12, $F_{(1,82)}$ = 12.9).*

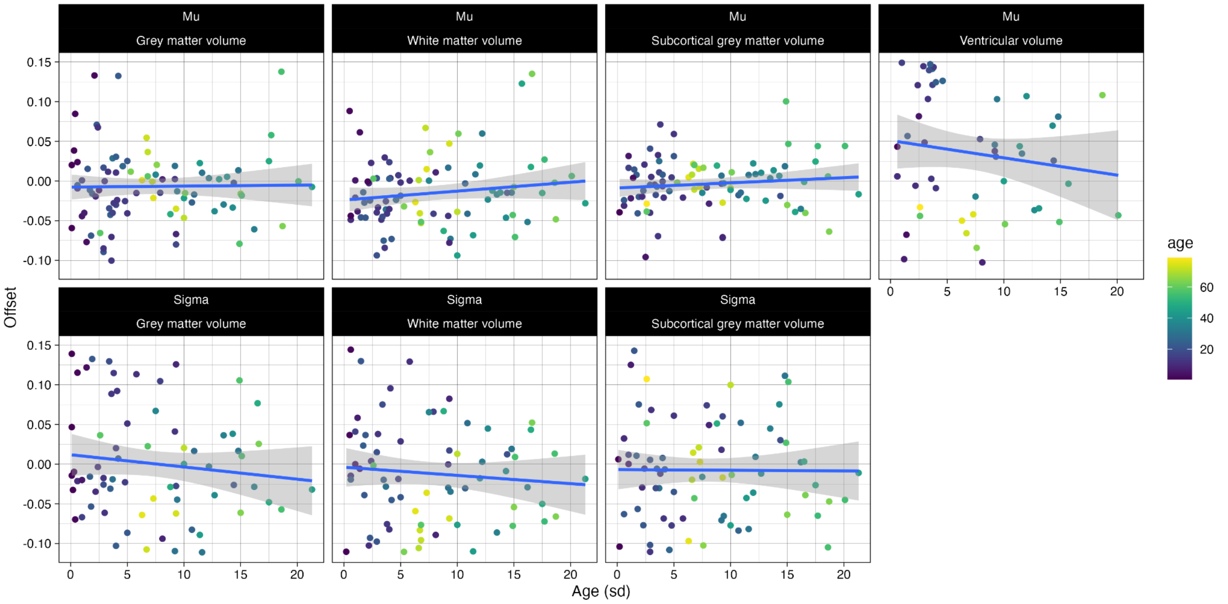


***Fig. S3.2.3.3. Association between the standard deviation of the age of participants and random effect parameters estimated by GAMLSS modeling of cerebrum tissue volumes for each primary study.*** *Top row: random effects on Mu (y-axis) are plotted versus standard deviation of age (x-axis) for each global MRI phenotype, left to right: grey matter volume (GMV), white matter volume (WMV), subcortical grey matter volume (sGMV) and ventricular CSF volume (Ventricles). Fitted lines and confidence intervals indicate the strength of association estimated by linear modeling. Bottom row: random effects on Sigma (y-axis) are plotted versus standard deviation of age for the same set of global MRI phenotypes (except Ventricles for which Sigma was not estimated). The association between random effects and standard deviation of age was not significant (after FDR correction for multiple comparisons) for any of these global MRI phenotypes.*

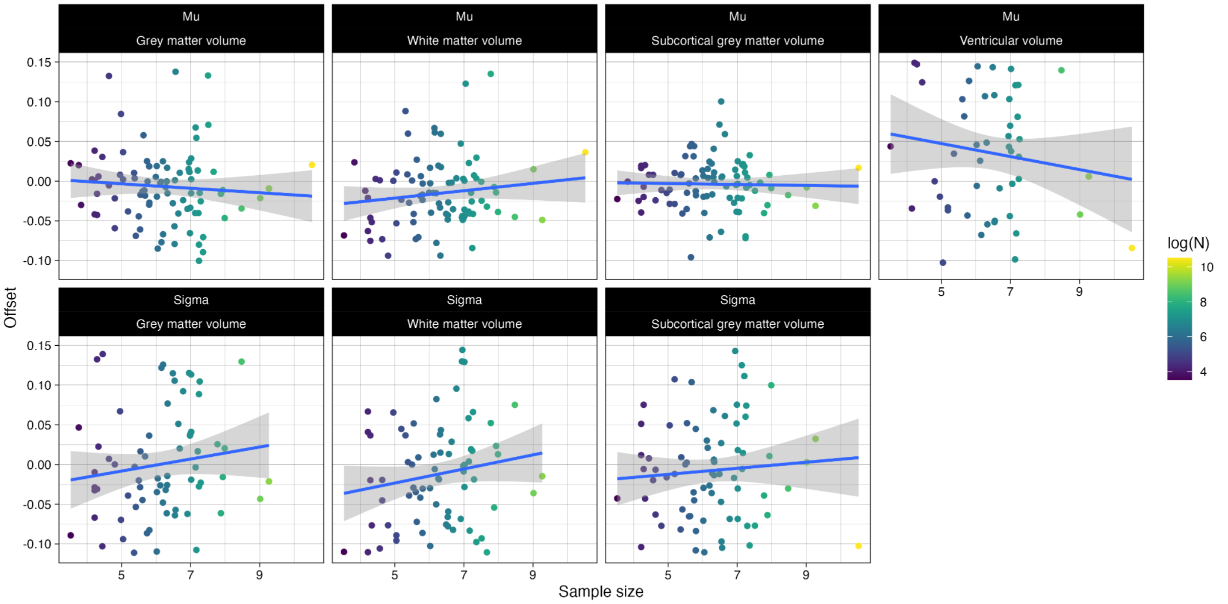


***Fig. S3.2.3.4. Association between sample size and random effect parameters estimated by GAMLSS modelling of cerebrum tissue volumes for each primary study.*** *Top row: random effects on Mu (y-axis) are plotted versus sample size (x-axis) for each global MRI phenotype, left to right: grey matter volume*

*(GMV), white matter volume (WMV), subcortical grey matter volume (sGMV) and ventricular CSF volume (Ventricles). Fitted lines and confidence intervals indicate the strength of association estimated by linear modeling. Bottom row: random effects on Sigma (y-axis) are plotted versus sample size for the same set of global MRI phenotypes (except Ventricles for which Sigma was not estimated). The association between random effects and sample size was not significant (after FDR correction for multiple comparisons) for any of these global MRI phenotypes.*

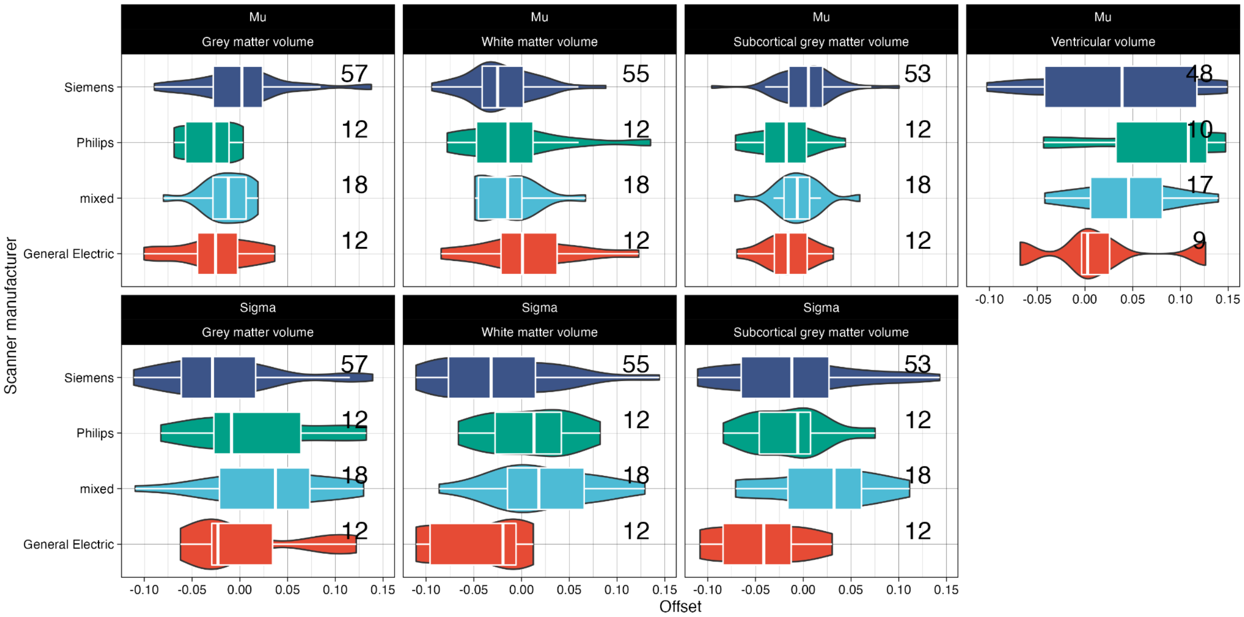


***Fig. S3.2.3.5. Association between the scanner manufacturer and random effect parameters estimated by GAMLSS modeling of cerebrum tissue volumes for each primary study.*** *Top row: boxplots of Mu (x-axis) are plotted for primary studies using scanners manufactured by General Electric (GE, red), Siemens (purple), Philips (green), or a mixture of different scanners (cyan), for each global MRI phenotype, left to right: grey matter volume (GMV), white matter volume (WMV), subcortical grey matter volume (sGMV) and ventricular CSF volume (Ventricles). Bottom row: boxplots of Sigma (x-axis) are plotted for primary studies stratified by scanner manufacturer (with the same colour coding) for the same set of global MRI phenotypes (except Ventricles for which Sigma was not estimated). There was no evidence for a significant difference in mean random effects of primary studies using different scanners (after FDR correction for multiple comparisons) for any of these global MRI phenotypes.*

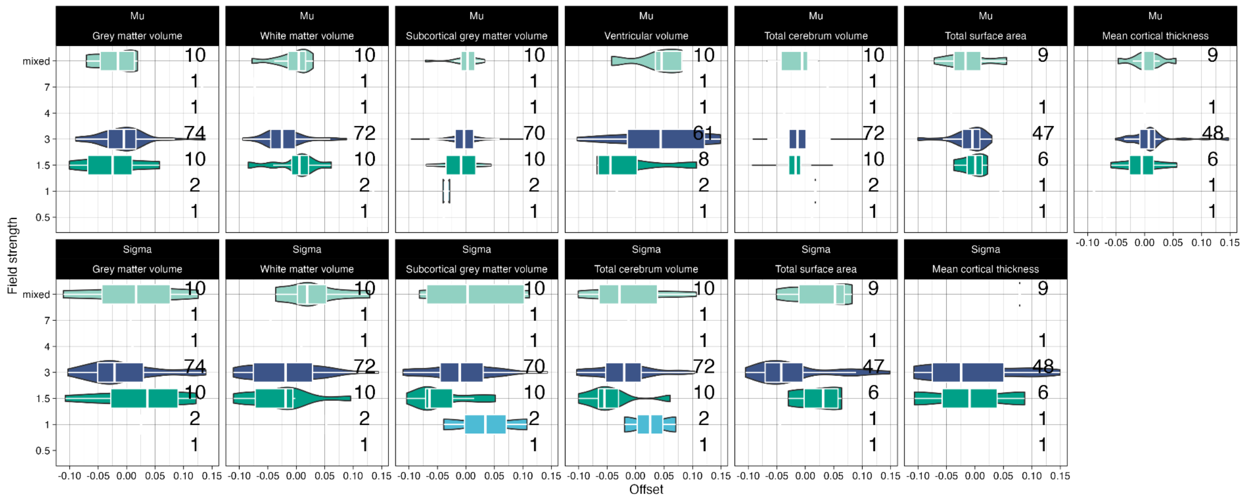


***Fig. S3.2.3.6. Association between the scanner field strength and random effect parameters estimated by GAMLSS modeling of cerebrum tissue volumes for each primary study.*** *Top row: boxplots of Mu (x-axis) are plotted for primary studies using scanners at different field strengths for each global MRI phenotype, left to right: grey matter volume (GMV), white matter volume (WMV), subcortical grey matter volume (sGMV) and ventricular CSF volume (Ventricles). Bottom row: boxplots of Sigma (x-axis) are plotted for primary studies stratified by scanner field strength (with the same colour coding) for the same set of global MRI phenotypes (except Ventricles for which Sigma was not estimated). There was no evidence for a significant difference in mean random effects of primary studies using scanners operating at different field strengths (after FDR correction for multiple comparisons) for any of these global MRI phenotypes. Numbers denote the number of studies included at this field strength.*

#### 3.3 Study-specific curves

Using the fitted model, specifically the fixed- and random-effects estimates, we can also derive study-specific prediction curves. On www.brainchart.io there are interactive plots of these studies' specific curves as the granularity of that figure is not suited for print format.

The study-specific prediction curves are obtained using the same method as the reference prediction curves described in **SI1.5**, using the mu-, sigma- and nu-component equations (**Eq1.5.1**) to calculate the predicted median (i.e., 50th percentile of the outcome distribution) across age and sex. However, there are two important differences. Firstly, we include a study-specific random-effect (where present) within the prediction calculations (i.e., random-effect terms within the component equations; **Eq1.1-1.2**), whereas in the reference prediction curves these are all set to zero (effectively not included). Secondly, the study-specific predictions are for the most common FreeSurfer version used within that study (if multiple FreeSurfer versions were used), whereas in the reference prediction curves the FreeSurfer contribution is equivalent to the grand-mean across all versions (across all studies), meaning the reference prediction curves correspond to a weighted average of FreeSurfer versions. All study- and individual-level analyses appropriately adjust for the specific version of FreeSurfer used. The study-specific prediction curves could be extrapolated across the entire lifespan, but we consider the age-censored versions to provide a more informative representation of study effects across the lifespan, which are normalized by the modelling approach. While most curves fall somewhere within the 95% centile boundaries of the population reference, clear deviations can be observed (see www.brainchart.io). Although each study-specific curve may be representative of that specific sample, this highlights that they may not always be appropriately extrapolated to available neuroimaging data at large.

##### 3.4 Brain weight, ultrasound, and head circumference validation

We performed a series of analyses to provide additional tests of functional interpretation of centile scores derived from the lifespan models, including a quantitative comparison to other data types traditionally used to measure brain size and growth (see dataset description below for further details on the data included in **Fig. S3.4**).

Historically, the international standards for growth modeling have been led by global initiatives and institutions, most notably the WHO[6]. These standards have been used and maintained for decades, but have been mainly developed for anthropometric measurements of height/length, weight, and head circumference. Until recently, ultrasound was the only non-invasive method for quantifying brain growth in utero, with head circumference and biparietal diameter being the measures of choice based on the ability to estimate in 2D. Brain size has been quantified via postmortem brain weight estimates for centuries, with conversions to mass being possible through scaling factors based on tissue density (1.03 kg/l). As such, we sought to provide quantitative relationships between model derived population reference values and each of the comparable estimates of brain size (head circumference, ultrasonic brain volume, and brain weight). We also compared the results of the lifespan model to an MRI dataset (10k-in-a-day) that could not be incorporated into the reference dataset because only age ranges were available due to privacy restrictions (not the precise age of individual participants).

To compare the GAMLSS trajectories to each of the traditional brain size features, we first determined the closest tissue types for each of the features: TCV (see supplementary figures below) for head circumference and brain weight, and GMV for the respective GMV estimates from one of the foetal ultrasound studies and the 10k-in-a-day MRI dataset. Next, due to the differences in age range and distribution of each of the replication studies, we extracted the median age and sex appropriate population reference points for the respective tissue classes from the GAMLSS model in each of the replication datasets. We then computed the correlation between the GAMLSS predicted and original sex-stratified mean values (**Fig. S3.4**), using log-scaled measures of brain weight and ultrasound-derived volumes (Pearson's *r*), and the naturally-scaled measure of head circumference (winsorized Pearson's *r*), due to the differences in units and methods of measurement [40].

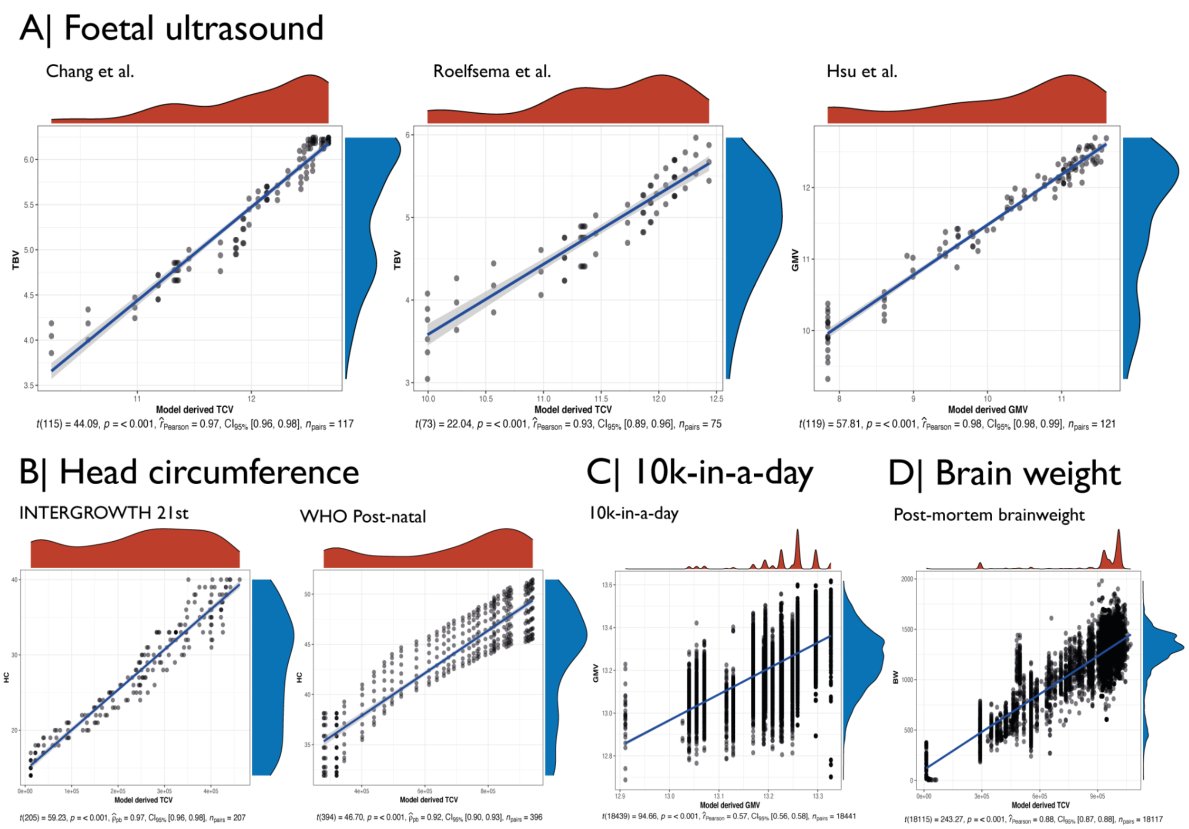


***Fig. S3.4. Validation of lifespan model-predicted values in independent datasets and modalities****. A | Three foetal ultrasound datasets, B | two head circumference reference norms (foetal=INTERGROWTH consortium, postnatal=WHO), C | a brain MRI dataset not included in the present models with only binned ages available, and D | four independent post-mortem brain weight datasets across the postnatal lifespan[41] (GTEx: https://gtexportal.org/home/, PsychENCODE: https://psychencode.synapse.org/). The neuroimaging models demonstrated high correlations (predicted vs. empirical values) across each of these modalities, thus showing the potential for inter-modal aggregation in future work.*

#### 4. Out-of-sample centile scoring: bias, stability and reliability

##### 4.1 Bias of out-of-sample centile scores: leave-one-study-out analyses for 100 studies

To further evaluate the robustness and consistency of centile scoring of OoS MRI data that were not included in the reference dataset used to estimate population trajectories, we performed a comprehensive series of leave-one-study-out (LOSO) analyses. For each one of the 100 studies in the reference dataset, we removed the study from the reference dataset, re-fitted the GAMLSS model to the remaining dataset of 99 studies, computed the OoS centile scores for the excluded study, and compared the OoS centile scores to the in-sample centile scores computed for the same study from the complete dataset including all 100 studies. Supplementary tables **ST7.1-7.4** list the correlations between OoS and in-sample centile scores for all 4 cerebrum tissue volumes in each of 100 primary studies. Overall, we found very high levels of correlation (Pearson's $r$ ~ 0.99) for almost all studies, indicating that centile scores can be estimated accurately for most studies even if they were not included in the reference dataset used to define population norms. Correlations between OoS and in-sample centile scores were lower than $r$ = 0.99 for only 3 out of

100 studies in the reference dataset: namely, the FinnBrain (*r* = 0.93), UCSD (*r* = 0.96) and NIHPD (*r* = 0.95) studies. These studies were characterised by relatively small sample size, foetal or early postnatal age range of participants, or idiosyncratic processing pipelines.

In addition to demonstrating high correlations between OoS and in-sample centile scores, we also evaluated their relative bias, defined as the difference between in-sample estimated centiles and OoS estimated centiles. The median bias in centile scores was generally low (GMV = -1.7e-06; WMV = 1.1e-04; sGMV = 3.8e-05; Ventricles= -7.3e-05, all with a standard deviation of ~0.01 centile). However, it is worth noting that the studies characterised by relatively small sample size, foetal or early postnatal age-range of participants, or idiosyncratic processing pipelines, appeared at the extreme ends of the distributions of the primary studies rank-ordered by the difference between in-sample and OoS centile scores (**Fig.S4.1.1**), indicating greater bias of OoS centile scoring, as expected, under these conditions.

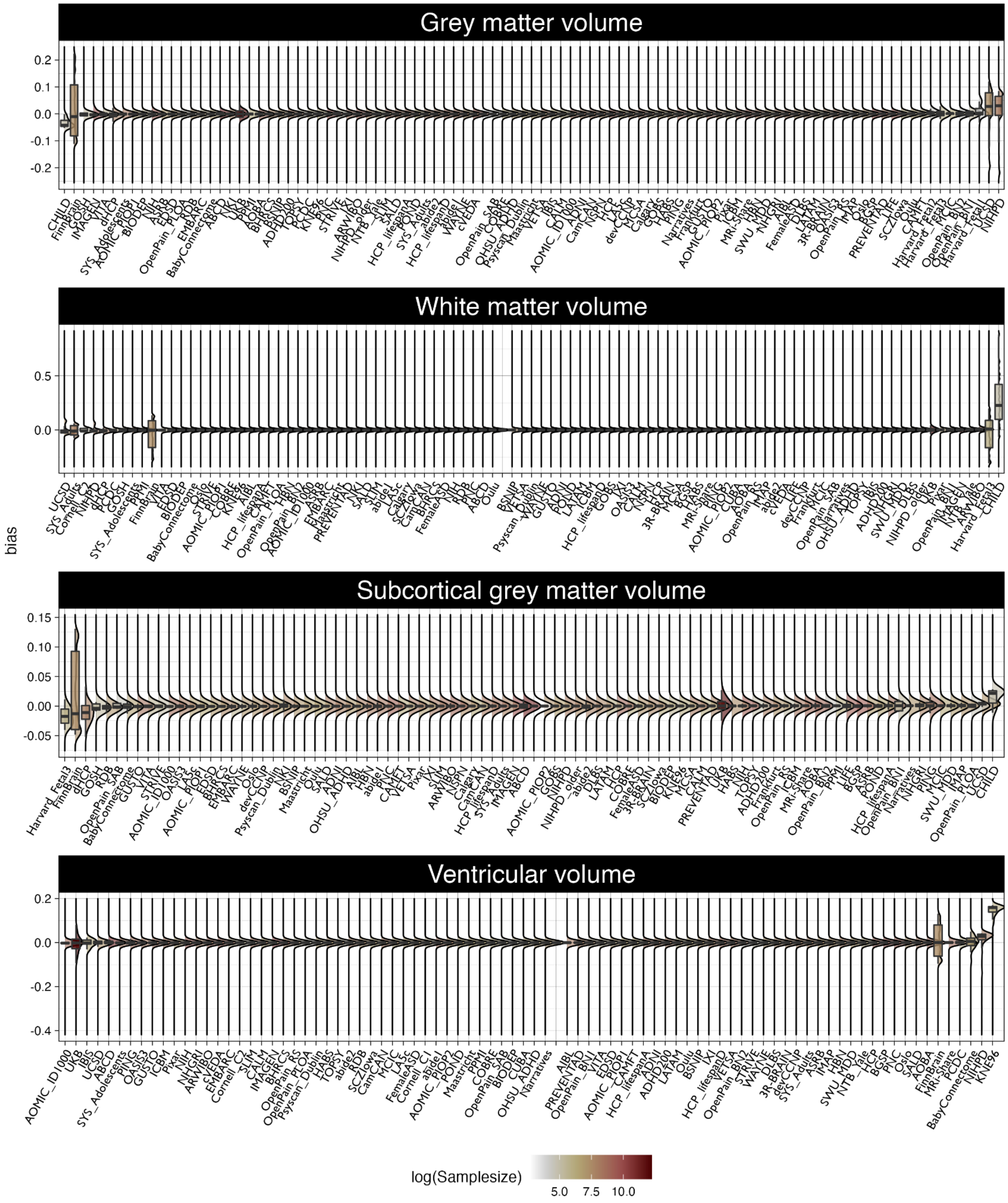


***Fig. S4.1.1. Bias of out-of-sample centile scores for four cerebrum tissue volumes.*** *Each panel shows boxplots of the bias in OoS centile scores (the signed difference between OoS and in-sample centile scores; y-axis) estimated for each primary study when it was excluded from the reference dataset. Studies are ordered on the x-axis from most negatively biased (left) to most positively biased (right) OoS centile scores. Boxplots are colour-coded according to log sample size, indicating that OoS centile scores tend to be most*

*biased for smaller primary studies. From top to bottom, panels represent the bias in OoS centile scores for grey matter volume, white matter volume, subcortical grey matter volume, and ventricular CSF volume.*

#### 4.2 Stability of out-of-sample centile scoring: bootstrapped LOSO analyses for 100 studies

In addition, we tested the reliability of OoS centile scores for each individual participant by bootstrapping. Specifically, for each LOSO sample, bootstrapped model parameters were generated (see **SI3.2.2 "Bootstrap analysis"**), resulting in 1,000 bootstrapped models with maximum likelihood estimated parameters for each bootstrap iteration of each left-out study. From this we obtained a bootstrapped distribution of out-of-sample centile scores for each individual subject in each individual iteration of left-out studies, thus providing a stability assessment in the form of the standard deviation of individual OoS centile scores across 1,000 bootstrap iterations. Across the datasets included in the model, we found that the average standard deviation of (bootstrapped) OoS centiles was 0.014, which is well below the level of within-subject longitudinal variation (see **Fig. S4.2.1** and **SI14 "Longitudinal centiles"**). Furthermore, we found increased standard deviation of OoS centile scores for datasets with comparatively small sample sizes (e.g., the OpenPain cohorts, Cambridge foetal Testosterone and CHILD studies; see **Fig. S4.2.2**). OoS centile scores were also more variable for datasets that had a more unique combination of age range, acquisition and processing pipelines (e.g., FinnBrain, IBIS and HBN; see **Fig. S4.2.2**). These observations reinforce the recommendation -- see main text, '**Out-of-sample centile scoring of "new" MRI data**' -- that OoS centile scoring is reliable for studies comprising N>100 scans. It was also notable that the reliability of OoS centile scores was weakly correlated with data quality as quantified by the Euler index (EI). So studies with higher number [31], indicating poorer image quality, tended to have higher variability of bootstrapped OoS centile scores (Pearson's *r* for all 4 cerebrum tissue volumes: GMV=0.05, WMV =0.11, sGMV =0.14, and Ventricular volume = 0.13). These results were not substantially different when the whole set of analyses was repeated with the dataset filtered on 2 median absolute deviations of the median EI. We conclude that OoS estimation of centile scores is generally reliable at the level of individual scans, and (as expected) reliability is greater for higher quality scans.

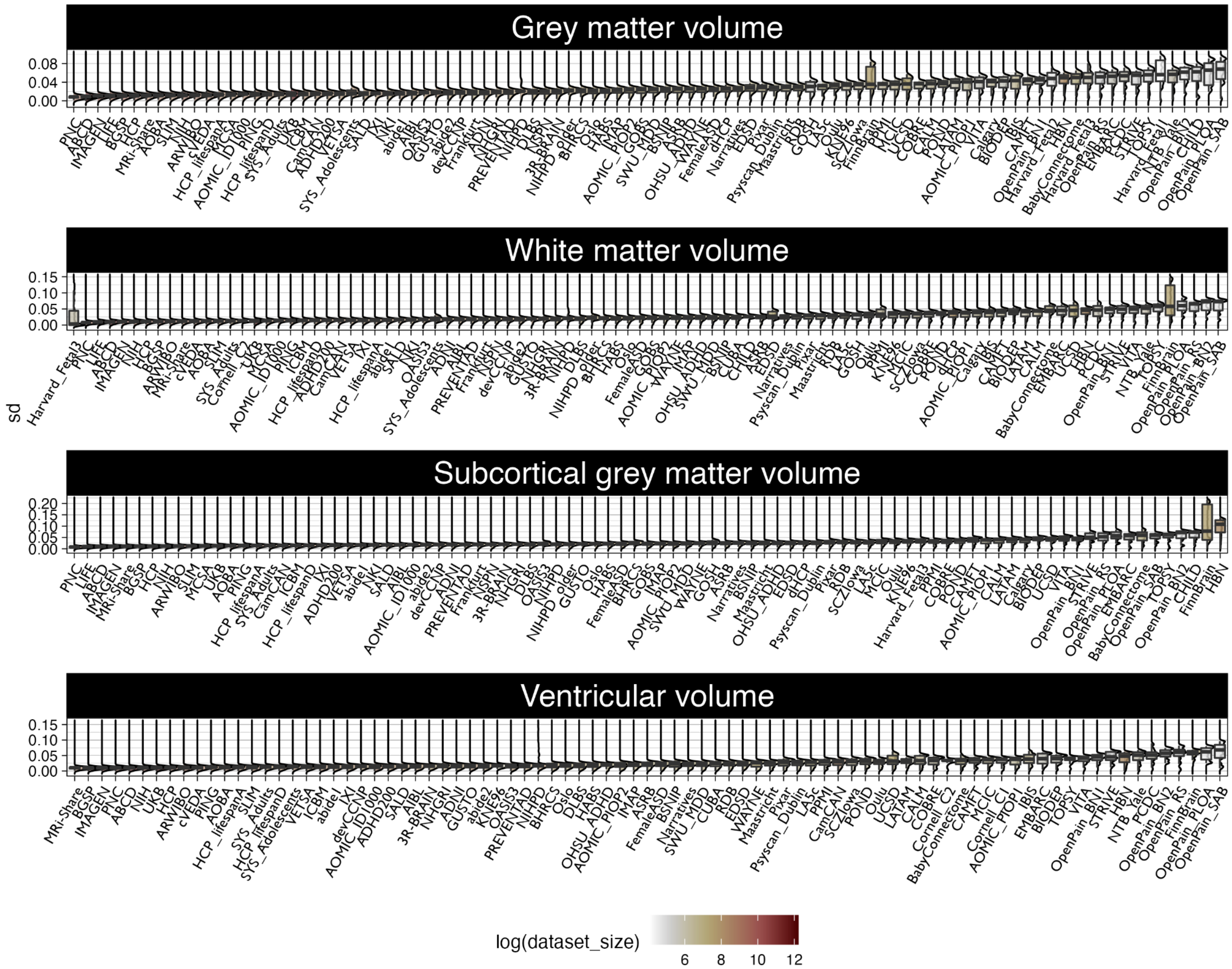


***Fig. S4.2.1. Stability of out-of-sample centile scores for four cerebrum tissue volumes when each of 100 studies was excluded from the reference dataset before bootstrapping.*** *The standard deviation of bootstrapped centile scores (y-axis) is plotted for each study (x-axis) for each phenotype, from top to bottom panels: total cortical grey matter volume, total cortical white matter volume, subcortical grey matter volume, and ventricular volume. Each study- and phenotype-specific boxplot is coloured according to log sample size. For each study, we estimated the normative model leaving that study out of the reference dataset and repeated this procedure after iteratively bootstrapping the reference dataset 1,000 times. This procedure allowed us to summarise the reliability of the out-of-sample estimates of centile scores in terms of the standard deviation of the 1,000 centile scores generated for each bootstrapped resampling of the reference dataset. Studies are ordered by median standard deviation of out-of-sample centile scores (small to large) indicating that scans are reliably assigned centile scores with the out-of-sample approach.*

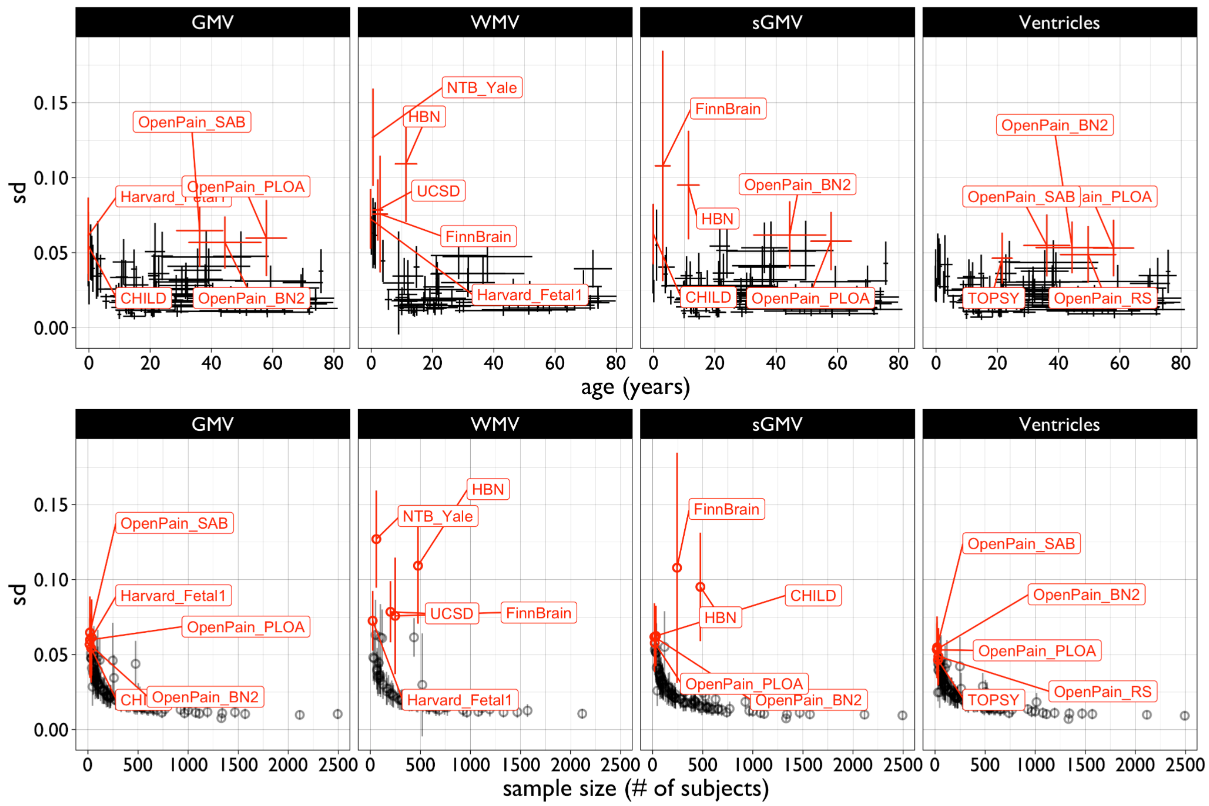


***Fig. S4.2.2. Stability of out-of-sample centile scores as a function of age and sample size.*** *The standard deviation (SD) of bootstrapped centile scores for four cerebrum tissue volumes (y-axis) is plotted against mean age of study participants (top row) or sample size (bottom row). Studies with the most unstable OoS centile scores (SD>0.05) are highlighted in red and labelled (see **ST1.1** for study details).*

#### 4.3 Test-retest reliability of out-of-sample centile scoring

We also assessed the reliability of OoS centile scoring in three independent datasets that acquired multiple MRI scans within a single session or two closely spaced sessions [30,42–44]. We analyzed each scan as a novel OoS dataset, then compared the consistency of centile scores across different scans of the same subject. We similarly compared the consistency of the uncentiled volumetric data and found that the out-of-sample centile scores were as consistent between scans in the same session as the “raw” volumetric data generated by FreeSurfer.

First, we analysed test-retest reliability using the multimodal MRI reproducibility resource[42], which provides two sessions of MRI data for multiple modalities. This dataset comprising 21 subjects was specifically designed for assessment of test-retest reliability as all subjects were scanned in two sessions separated by a one-hour break and the whole cohort was completed within a two week period. We analyzed each session of 21 scans as an independent OoS study (**Fig. 5**) and then estimated intra-class correlation coefficients (ICCs) to assess the between-session or test-retest reliability of individual centile scores for four cerebrum tissue volumes[45]. All ICCs were ~0.99 (**Fig. S4.3.1**).

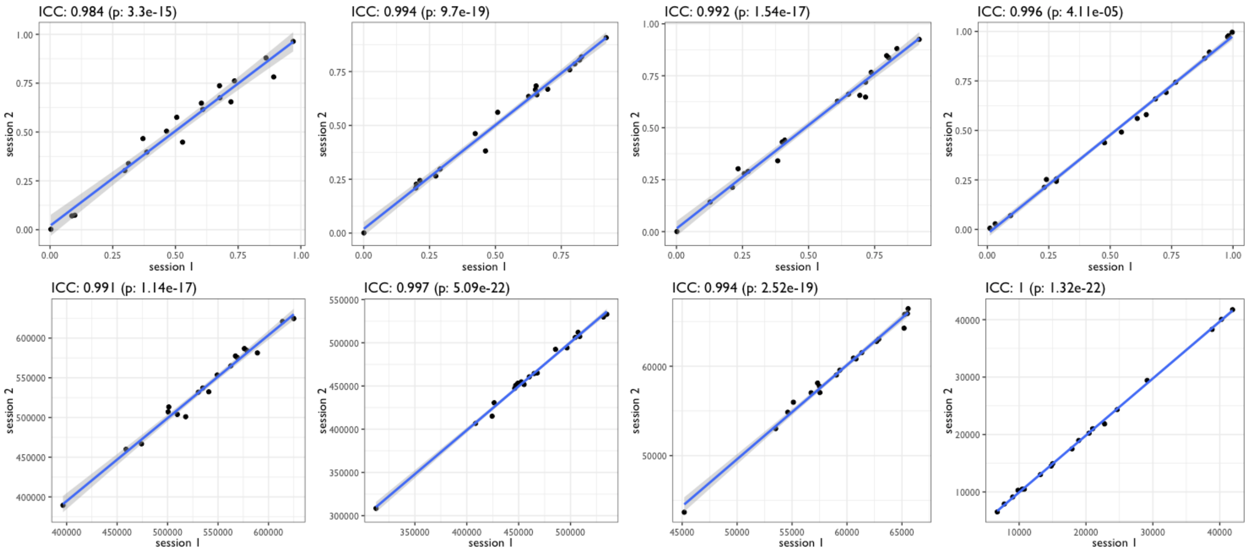


***Fig. S4.3.1. Test-retest reliability of out-of-sample centile scores for cerebrum tissue volumes.*** *MRI data were collected in two separate scanning sessions from N=21 participants and each session was analysed as an independent out-of-sample study using GAMLSS. Scatterplots represent OoS centile scores for session 1 (y-axis) versus OoS centile scores for session 2 (x-axis) for each brain tissue volume, from left to right: GMV, WMV, sGMV, Ventricular CSF. Data points represent individual subject centile scores. Test-retest reliability was consistently very high (all ICCs > 0.99) for all cerebrum tissue volumes. The bottom panel shows the identical analysis for non-centiled, "raw" volumetric data. P-values represent the significance of the intraclass correlation coefficient between two sessions.*

Second, we analysed the test-retest reliability of OoS centile scoring using MRI data on N=72 participants in the Healthy Brain Network (HBN) cohort[30], which was not originally included in the reference dataset. The HBN cohort was designed to assess the influence of an alternate MRI data acquisition protocol, which included prospective motion correction[34] to improve quality and reliability of MRI. The study protocol included 2 sessions of scanning using a conventional MPRAGE sequence for T1-weighted data acquisition and another 2 sessions of scanning using an innovative, prospectively motion-corrected sequence, VNaV, for T1-weighted imaging[34]. For all 72 individuals each session of each sequence was analysed as an OoS study (**Fig. 5; SI1.8 "Out-of-sample estimation"**) and then we estimated ICCs as a measure of the test-retest reliability of individual centile scores for each brain tissue volume derived from each sequence (MPRAGE or VNaV). Test-retest reliability was uniformly high (ICCs > 0.95) for all OoS centile scores on all cerebrum tissue volumes estimated from both MPRAGE and VNaV sequences (**Fig. S4.3.2**). Reliability was incrementally higher for OoS centile scores derived from the VNaV sequence, under-scoring the importance of high-quality data especially for OoS analysis of datasets with N<100. However, we note that this increased reliability of centile scoring was most likely driven by a comparably increased consistency of the raw volumes estimated by FreeSurfer (as also noted in the original paper describing the impact of prospective motion correction[30]).

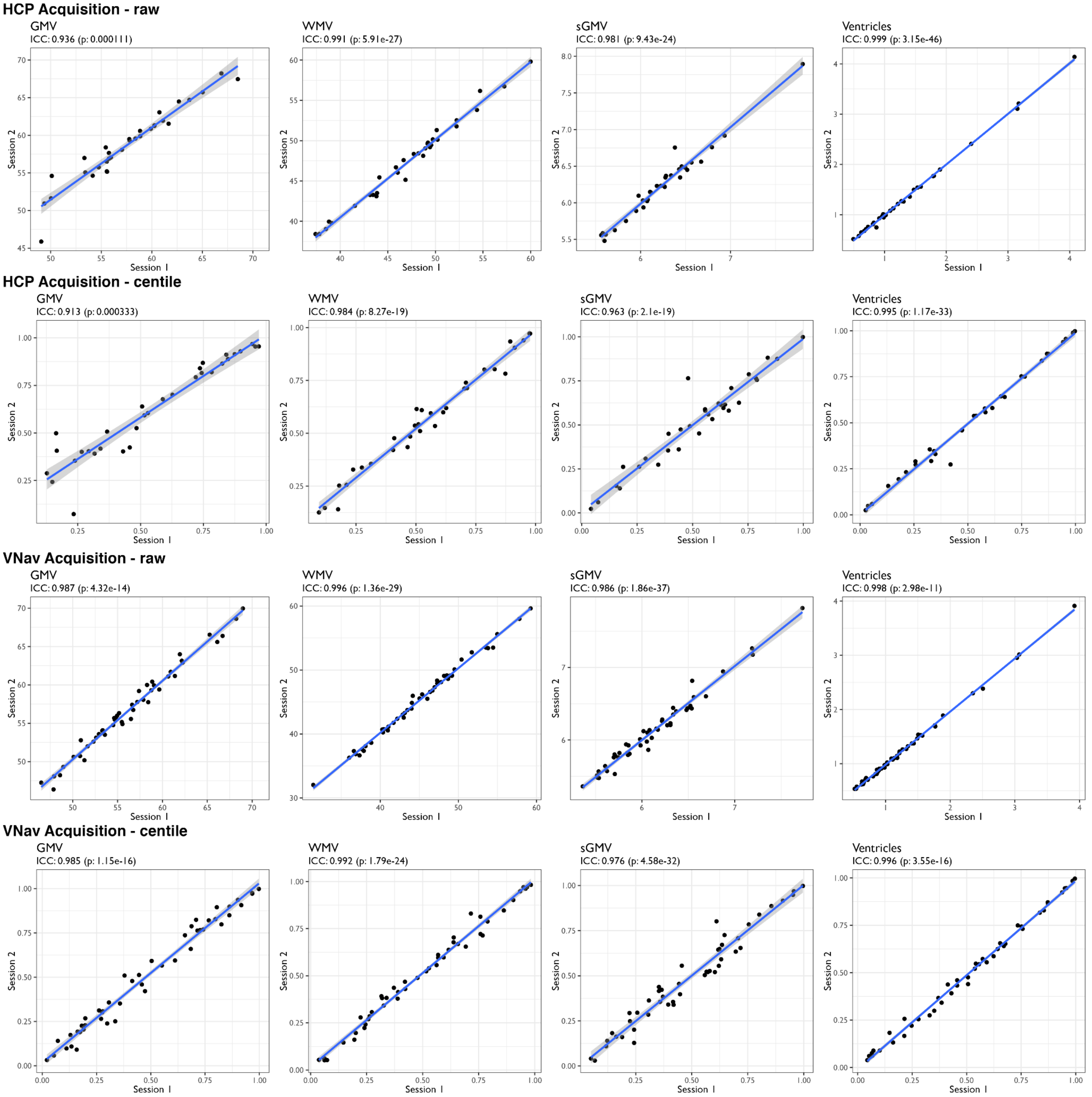


***Fig. S4.3.2. Test-retest reliability of out-of-sample centile scores for cerebrum tissue volumes measured twice in the same N=72 participants using two T1-weighted sequences, MPRAGE and VNaV.*** *For each type of acquisition, the top row shows out-of-sample centile scores for session 1 (y-axis) versus out-of-sample centile scores for session 2 (x-axis) for cerebrum tissue volumes. For each type of acquisition, the bottom row shows the unprocessed ("raw") scores for session 1 (y-axis) versus session 2 (x-axis) for cerebrum tissue volumes estimated from VNaV data, from left to right: GMV, WMV, sGMV, Ventricles. In all plots, data points represent individual subject scores. Test-retest reliability was uniformly high (all ICCs > 0.95) and generally somewhat higher for volumetrics derived from prospectively motion-corrected data (VNaV). P-values represent the significance of the intraclass correlation coefficient between two sessions. HCP session refers to the Human Connectome Project MPRAGE acquisition used.*

Third, we assessed the test-retest reliability of OoS centile scoring using the Vietnam Era Twin Study of Ageing (VETSA) study cohort[43]. VETSA is a longitudinal study following 1,200 twins from the Vietnam Era Twin Registry, which includes two technically identical MPRAGE acquisitions within the first (baseline) scanning session. Both these scans were processed with FreeSurfer 6.0.1 for all participants, then the two sets of scans were each analysed as an independent OoS study, and ICCs were estimated to assess the test-retest reliability of individual centile scores on all four cerebrum tissue volumes. Test-retest reliability of OoS centile scores was uniformly very high (all ICCs > 0.98) across all phenotypes, comparable to the high reliability of the uncentiled volumetric data generated by FreeSurfer 6.0.1 (all ICCs > 0.95), and in line with the constraints on reliability expected from technical sources of noise[46] (**Fig. S4.3.3**).

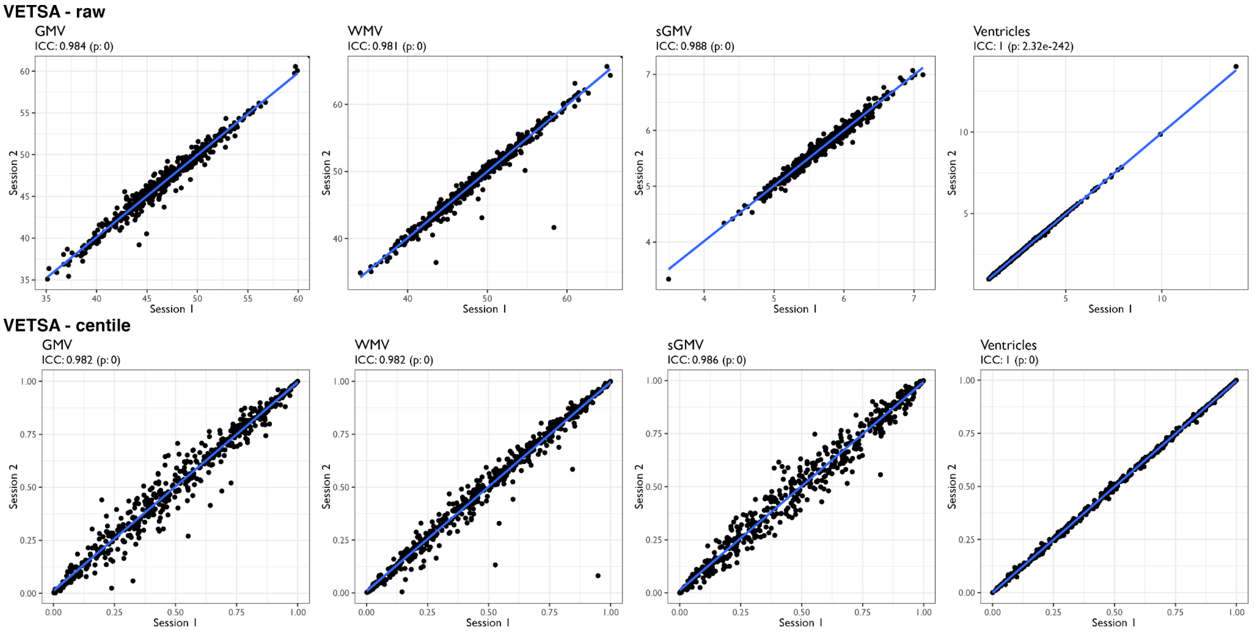


***Fig. S4.3.3. Test-retest reliability of out-of-sample centile scores for cerebrum tissue volumes measured twice in the same 1,200 participants (600 twin pairs).*** *The top row shows scatterplots of unprocessed ("raw") volumes for scan 1 (y-axis) versus scan 2 (x-axis) for cerebrum tissue volumes estimated from MPRAGE data from the same subject, from left to right: GMV, WMV, sGMV, Ventricles. Data points represent individual subject centile scores. The bottom row shows the consistency of centile scores for the same subjects and same phenotypes. Reliability was uniformly high across all phenotypes (ICCs > 0.95) and comparable to reliability of uncentiled volumetric measurements from the same set of scans. P-values represent the significance of the intraclass correlation coefficient between two sessions.*

#### 4.4 Reliability of out-of-sample centile scoring across multiple versions of FreeSurfer

Knowing that a large majority (~95%) of primary studies in the reference dataset used one of a series of versions of FreeSurfer for image analysis, we also evaluated the impact of these incrementally different image analysis pipelines on reliability of OoS centile scores. To do this we repeatedly re-analysed a single NIH dataset[44] (see **SI 19** "NIH" for a fuller description) using 4 different versions of FreeSurfer (5.1, 5.3, 6.01, and 7.1). Each version of the processed dataset was treated as an independent OoS study for GAMLSS modeling and then we estimated ICCs between individual centile scores for each possible pair of FreeSurfer pipelines and for each of four cerebrum tissue volumes. This analysis demonstrated generally high within-subject reliability of OoS centiles across all four pipelines: ICCs for GMV=0.978, WMV=0.972, sGMV=0.816 and

Ventricles=0.982 (**Fig. S4.4**). We noted that there was somewhat reduced reliability of subcortical grey matter volume in both raw and centiled data from FreeSurfer version 5.1 in comparison to later FreeSurfer versions. While the reasons for this are unclear, none of the studies included in the principal dataset were processed with FreeSurfer 5.1, or any version of FreeSurfer older than 5.3. Furthermore, we found the highest between-pipeline reliability for both raw volumetric data and centile scores derived from the two most recent versions of FreeSurfer, 6.0.1 and 7.1, suggesting that minor inconsistencies due to FreeSurfer pre-processing are becoming less problematic as this widely used software package incrementally evolves.

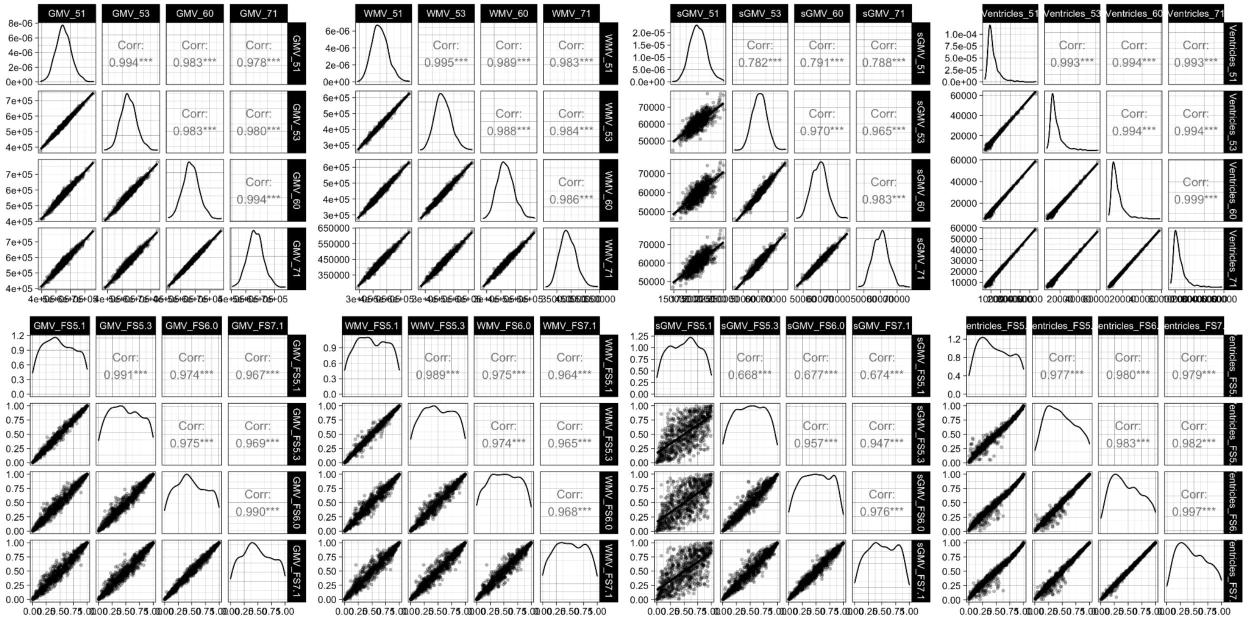


***Fig. S4.4. Between-pipeline reliability of volumetric data and out-of-sample centile scores for four cerebrum tissue volumes measured in the same set of N=1,468 scans re-analysed using 4 different versions of FreeSurfer (5.1, 5.3, 6.01, and 7.1).*** *Top row shows scatterplot matrices representing the correlations between raw volumetric data derived from each possible pair of FreeSurfer pipelines, from left to right: GMV, WMV, sGMV, Ventricles. Bottom row shows scatterplot matrices representing the correlations between out-of-sample centile scores derived from each possible pair of FreeSurfer pipelines, from left to right: GMV, WMV, sGMV, Ventricles. Intra-class correlations of out-of-sample centile scores and uncentiled volumetric data, on average over all pairs of four pipelines, were generally high (GMV=0.978, WMV=0.972, sGMV=0.816 and Ventricles=0.982). Although the reliability of sGMV volumetrics and centile scores was somewhat lower due to discrepant measurements by the oldest version of FreeSurfer, v5.1, this version of FreeSurfer was not used to analyse any of the scans included in the reference dataset.*

#### 4.5 Effects of sample size on reliability of out-of-sample centile scores

To further assess the validity of the OoS estimates, we generated “clones” of existing datasets. Clones were resampled subsets (without replacement, no duplicate subjects per clone) of studies included in the reference dataset used to estimate the study specific GAMLSS parameters. Each clone was then treated as if it was a “new” study using the methods for out-of-sample centile scoring. This allows us to compare the OoS estimates to a relative truth, i.e., from the original, non-cloned version of the study included in the reference dataset, we know what the GAMLSS parameters ‘truly’ are, and we have an estimation of their ‘true’ uncertainty from the bootstrap resampling distributions. Thus for a given study dataset, $D_m$, we generate a cloned copy $D_1$, and if our approach is unbiased we expect the out-of-sample parameter estimates for $D_1$ to be equal to the in-sample parameters estimated for $D_m$, i.e., $\gamma_{\cdot,m}$ (representing the set of random effects estimated by in-sample analysis of the original study treated as part of the reference dataset) should approximate $\gamma_{\cdot,1}$ (representing the set of random effects estimated by OoS analysis of the cloned study treated as a new dataset): see **SI1.8 "Out-of-sample estimation"** and **Fig. S4.5**.

In other words, we validated the OoS estimation by simulating a "new" study with the same underlying distribution as one of the studies included in the reference dataset. Hence, we expect the OoS random-effect estimates for this 'clone' to agree with the in-sample random-effect estimates. More formally, we are comparing $\gamma = MLE_{\beta,\gamma}(D)$ and $\gamma^{Clone} = MLE_{\gamma}(D_{Clone}|\beta(D))$, where the clone is contained within the data, i.e., $D \cap D_{Clone} = D_{Clone}$; see **SI1.8 "Out-of-sample estimation"** for further details on OoS MLE estimation. We used the Neuroscience in Psychiatry Network dataset (NSPN) to evaluate at what relative sample size the OoS estimation approaches the true parameter offset. As illustrated in **Fig. S4.5**, these simulations indicated good performance for the OoS approach for "new" study sizes greater than N=100 scans. In addition to the internally collected NSPN dataset, we also confirmed this optimal sample size in the ADNI cohort.

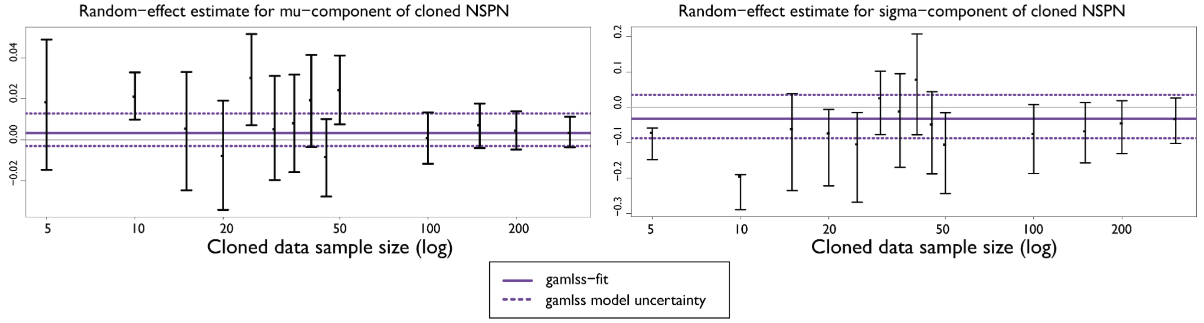


***Fig. S4.5. Out-of-sample estimates of cloned study random-effect parameters compared to in-sample estimates of random-effect parameters in the original or non-cloned study.*** *The plot shows random-effects estimated using the out-of-sample approach across a range of possible sample sizes for a "new" study, generated by taking subsets of the same cloned study with uncertainty intervals derived from the bootstrap replicates. The purple horizontal lines are the equivalent in-sample estimates of the random-effects parameters. We see that the out of sample estimates are somewhat unreliable below N=100 subjects, but with larger samples the out-of-sample estimates from the cloned data converge with the in-sample estimates from the original data for both $\mu$-component and $\sigma$-component random effects. NSPN refers to the Neuroscience in Psychiatry Network study included in the reference dataset.*

#### 5. Batch correction and site harmonisation

##### 5.1 Modeling of between-site heterogeneity by GAMLSS: conceptual considerations in comparison to ComBAT batch-correction

Batch effects, or heterogeneities between sites or primary studies, are a challenging issue for estimating generalisable results from multi-site or multi-study neuroimaging data. In recent years, methods such as ComBAT[14,47] have been translated from their primary application for whole genome transcription (microarray) analysis to achieve harmonisation of MRI data acquired across multiple sites. For our principal analysis, however, we preferred to use GAMLSS, a conceptually similar mathematical framework, to account for between-site or between-study heterogeneity. We made this choice *a priori* for several reasons. Firstly, GAMLSS explicitly includes the possibility of accounting for non-linear age effects (including age-related changes to higher order moments such as variance) during the harmonisation process. Adaptations of traditional ComBAT harmonisation have recently been developed that also allow the inclusion of non-linear age-trends as well as longitudinal, within-subject effects[48,49]; but these refinements of ComBAT remain somewhat restricted to batch correction of the mean and are not trivial to extend to batch correction of higher order moments, such as the variation across sites. Secondly, we chose to use GAMLSS because it is flexible with regards to the underlying distribution of the data that is to be harmonised; thirdly, because GAMLSS is the WHO-recommended statistical framework for growth chart modelling[26]; and finally because GAMLSS allows a flexible modelling capacity that would facilitate scaling of this framework to growth charting of additional MRI phenotypes in the future.

Conceptually, normalised centiles derived from the GAMLSS model (see **SI1.5**) are analogous to normalised scores derived from ComBAT. Specifically, multiple groups of observations have an induced co-dependence, arising in the context of our analysis from common study-specific factors, which leads to a common measurement bias. The aim of both ComBAT and GAMLSS is to correct that common measurement bias. However, whereas ComBAT is derived from a conjugate Bayesian approach and hence restricted to a Gaussian distribution of phenotypes, GAMLSS uses a frequentist, iterative maximum likelihood approach that allows a range of distributions including those with non-zero third and fourth statistical moments (the Gaussian distribution by definition has third and fourth moments equal to zero). Flexibility in the distribution is important, especially for potentially highly skewed measures (with non-zero third moments), and to allow distributions that conform with the distributions of the measurements. ComBAT assumes that these distributions are naturally Gaussian or can be rendered approximately Gaussian by a simple (e.g., log) transformation. However, even if working with Gaussian measurements, the mean and variance may require non-constant terms to account for heteroskedasticity, and the resulting models are dependent on non-intuitive transformations for Gaussianisation.

In the context of the present study, we used the Bayesian information criterion (BIC) to assess the goodness-of-fit of GAMLSS models making different assumptions about the form of the phenotypic distributions. We found that not only was the Gaussian a suboptimal distribution, but that the optimal choice was the generalised gamma distribution, which includes a third order moment. Although we found no evidence of an age-related change in the third order moment, it was different from unity and hence there was evidence of skewness (otherwise we could reduce it to the gamma distribution, which is the simplified form of the generalised gamma). The (generalised) gamma distribution is also defined only on the positive real line, negating the need to perform any transformations (apart from multiplicative scaling for computational stability), meaning the fitted model coefficients are on the same scale as the original phenotype.

The GAMLSS and ComBAT approaches to batch correction differ substantially in a few other ways. Whereas GAMLSS directly uses centiles and medians of the phenotypic distribution, ComBAT uses the mean and variance. Hence, when comparing these methods, we cannot expect exactly the same results, even if we enforce a Gaussian outcome distribution within GAMLSS. Another substantial difference between the GAMLSS and ComBAT approaches is that GAMLSS requires a substantial amount of data. Even with the number of observations available for our analysis, it has been necessary to use restricted forms, i.e., fractional polynomials, for the normative lifespan trajectories rather than more flexible forms, e.g, splines. Furthermore, ComBAT is defined on a multivariate (Gaussian) phenotype distribution, whereas we used GAMLSS to model multiple univariate phenotypes. (GAMLSS does have some capability to model multivariate distributions, but this area is currently under-developed.) Therefore, ComBAT is able to adjust for batch effects with fewer observations on the assumption that the batch effect is shared across multiple phenotypes. Running ComBAT in a univariate mode would be most directly equivalent to the GAMLSS approach but this is not how it is used in the wider literature. This implies that multivariate normalisation by ComBAT is to some extent dependent upon the set of phenotypes included; if a new phenotype is included the ComBAT correction for batch effects would need to be re-run.

#### 5.2 Modeling of between-site heterogeneity by GAMLSS: empirical evaluation compared to ComBAT

To empirically evaluate the capacity of GAMLSS to account for batch effects or between-site variation, we analysed the well-known multi-site ABCD study[50] and compared the results of between-site harmonisation by GAMLSS to the results of a standard ComBAT harmonisation pipeline. We specifically chose the ABCD study to test the capacity of GAMLSS and ComBAT to remove between-site noise because it is a demographically harmonised multi-site cohort. In addition the large sample size of healthy controls per site makes ABCD highly suitable for GAMLSS harmonisation of between-site differences (see also **SI 4.5**). This means that in the context of ABCD any residual significant differences between sites are less likely to be due to true site variation or recruitment differences and more likely to be due to noise (technical or otherwise), though even in this study recruitment bias can not be fully eliminated. In addition, the ABCD dataset also provided a wide range of non-MRI phenotypes to test any downstream impact of batch-effect correction approaches on analyses of association between MRI centile normalised scores and non-MRI phenotypes. Despite being a technically harmonised cohort, and despite using acquisition protocols that included prospective motion correction, the uncorrected ABCD imaging data still show clear and significant differences between sites across all MRI phenotypes. Both ComBAT and GAMLSS efficiently removed these batch effects in the normalised (site-corrected) data, but both harmonisation pipelines retained a high degree of variation at the level of individual scans (**Fig. S5.2.1-5.2.2**).

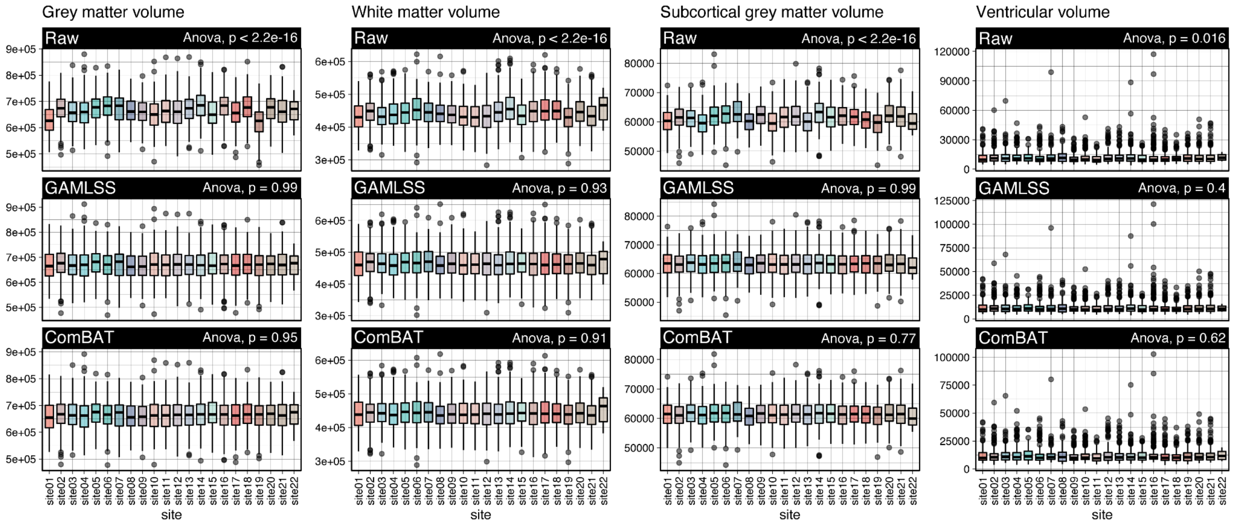


***Fig. S5.2.1. Raw volumetric data and centile scores for male subjects from the ABCD cohort.*** *The top row shows raw volumetric data across the 22 sites included in ABCD, the middle row shows centile normalised data by GAMLSS and the bottom row shows data normalised using ComBAT. ANOVA P-values refer to one-way analyses of variance across sites for each individual phenotype. Bars are coloured by site. ComBAT and GAMLSS are both able to substantially mitigate batch effects in multi-site MRI data.*

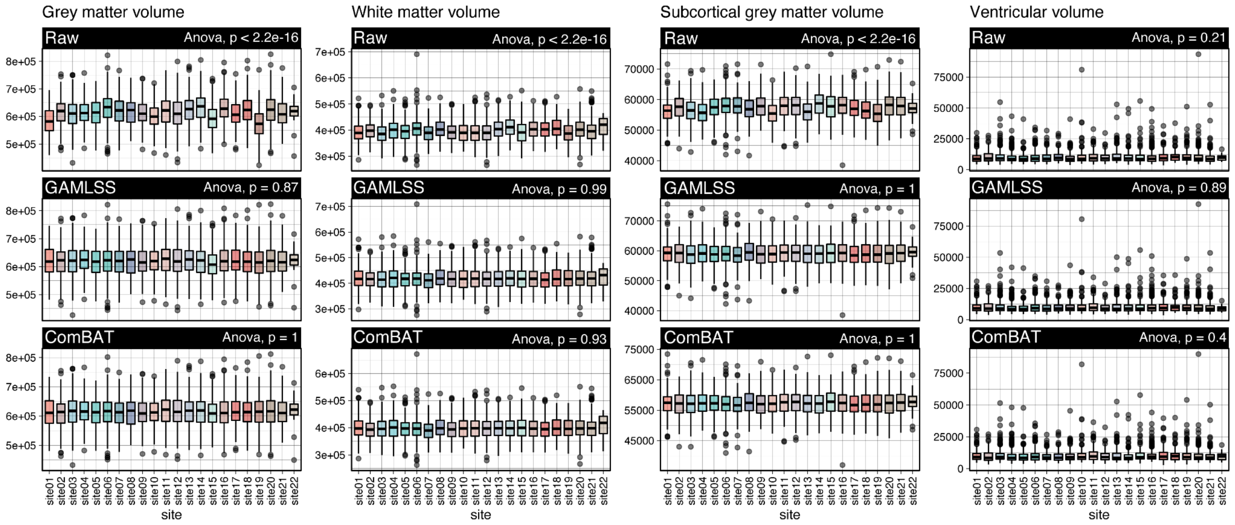


***Fig. S5.2.2. Raw volumetric data and centile scores for female subjects from the ABCD cohort.*** *The top row shows raw volumetric data across the 22 sites included in ABCD, the middle row shows centile normalised data by GAMLSS and the bottom row shows data normalised using ComBAT. ANOVA P-values refer to one-way analyses of variance across sites for each individual phenotype. ComBAT and GAMLSS are both able to substantially mitigate batch effects in multi-site MRI data.*

To further assess whether batch-corrected MRI data derived from both ComBAT and GAMLSS pipelines would generate convergent results in subsequent analyses, we estimated the correlations between total cerebrum volume (TCV) and fluid intelligence or birth weight, after TCV had been batch-corrected by either GAMLSS or ComBAT. Both these psychological and biological factors have previously been shown to be correlated with similar brain volumetrics[51–53]. We were able to replicate these significant associations with uncorrected TCV, as well as after both GAMLSS and ComBAT batch correction, all largely showing consistent effects across sites (**Fig. S5.4-5.5**).

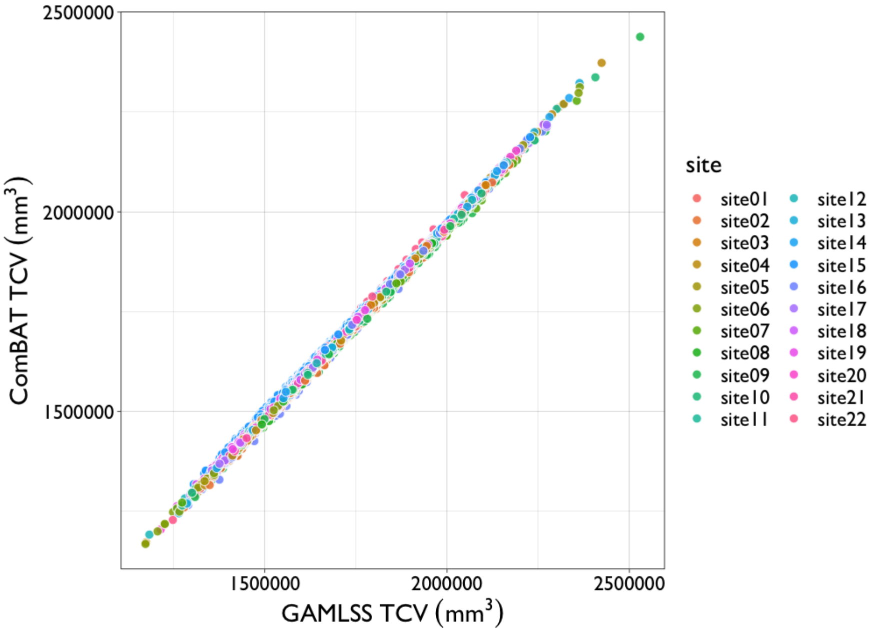


***Fig. S5.2.3. Comparing effects of GAMLSS versus ComBAT batch correction on estimation of total cerebrum volume.*** *TCV was estimated for N=10,583 participants in the ABCD multi-site study after MRI data had been batch-corrected for between-site differences by ComBAT (y-axis) or GAMLSS (x-axis). Estimated TCV was highly correlated (r > 0.99) downstream of these two batch correction procedures. Scans are point-coloured according to site.*

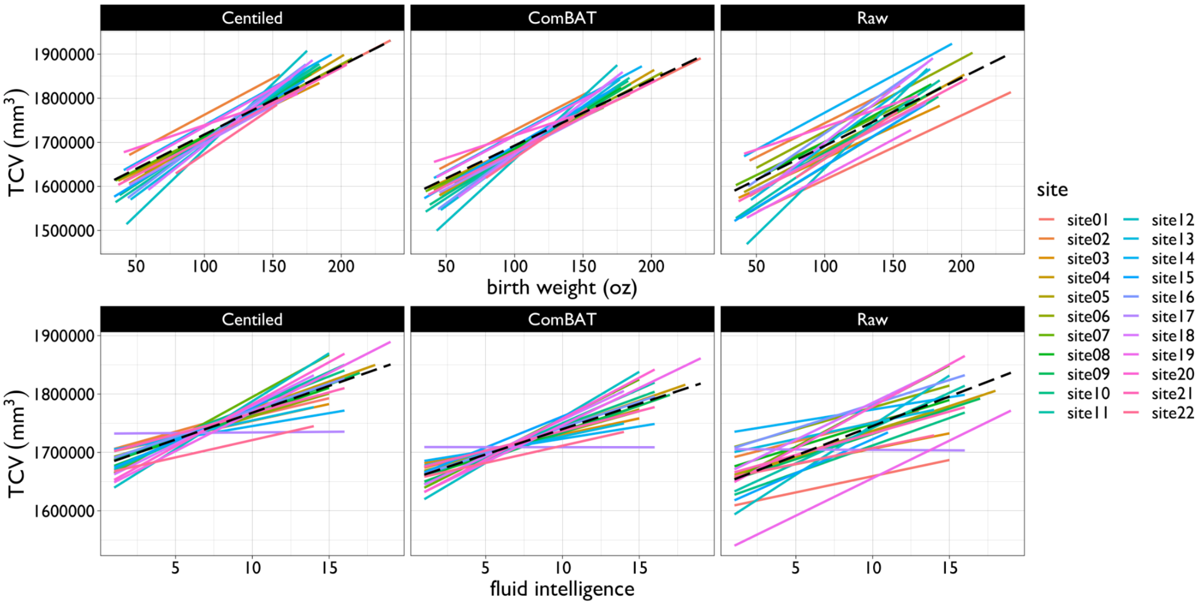


***Fig. S5.2.4. Associations between total cerebrum volume (TCV) and birth weight (top) or fluid intelligence (bottom) after batch correction by GAMLSS (left), by ComBAT (middle), or without batch correction (raw, right).*** *Linear relationships for each of the 22 sites in the ABCD study are in coloured solid*

*lines; dashed lines signify overall model fit across sites; fluid intelligence was assessed using the NIH Toolbox[54]. These results show that predicted relationships between TCV and both birth weight and fluid intelligence are more convincingly replicated in these N=10,583 scans from the ABCD multi-site study when the MRI data have been batch-corrected by either GAMLSS or ComBAT compared to when the MRI data have been analysed without correction of between-site differences.*

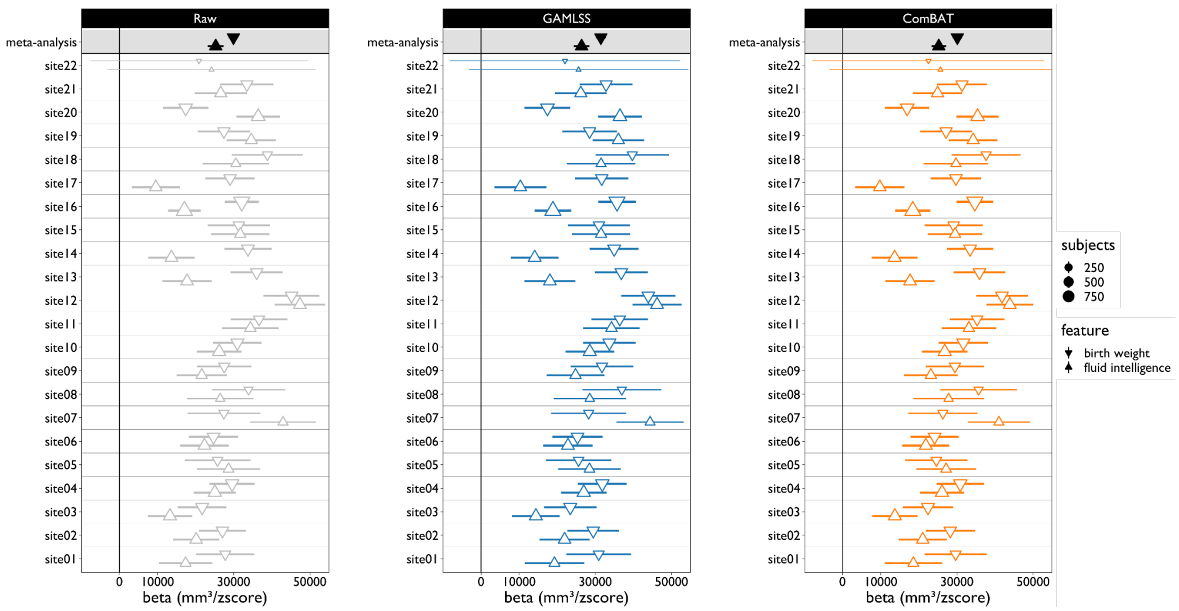


***Fig. S5.2.5. Consistency of behavioural (fluid intelligence) and biological (birth weight) associations with total cerebrum volume (TCV) estimated at 22 MRI acquisition sites in the ABCD cohort, either without batch correction (left) or after batch correction for site effects by GAMLSS (middle) or ComBAT (right).*** *Regression coefficients and standard errors from linear regression models of TCV on birth weight or fluid intelligence are plotted using point-ranges for each site. Meta-analytic coefficients and errors, combining all primary study effects, are shown in black at the top of each column. Coefficients (triangles) are scaled based on sample size at each site within the ABCD study.*

Few other datasets fit the selection criteria used for the specific comparison between GAMLSS and ComBAT approaches to normalisation (i.e., N >100 healthy control participants per site, aligned recruitment criteria, and broadly aligned MRI data acquisition protocols). The only other multi-site datasets fitting these criteria in our aggregated dataset were the IMAGEN and UK BioBank cohorts. To explore whether the harmonisation approach worked well in a cohort other than ABCD we chose the IMAGEN cohort as UK BioBank implements an extremely well-harmonised acquisition and recruitment strategy across its 3 sites. While we did not have access to the same extensive set of non-neuroimaging based phenotypes in the IMAGEN dataset as we had for the ABCD dataset, we observed that both GAMLSS and ComBAT were highly effective at removing large site-related variation from raw neuroimaging phenotypes (**Fig. S5.2.6-5.2.7**).

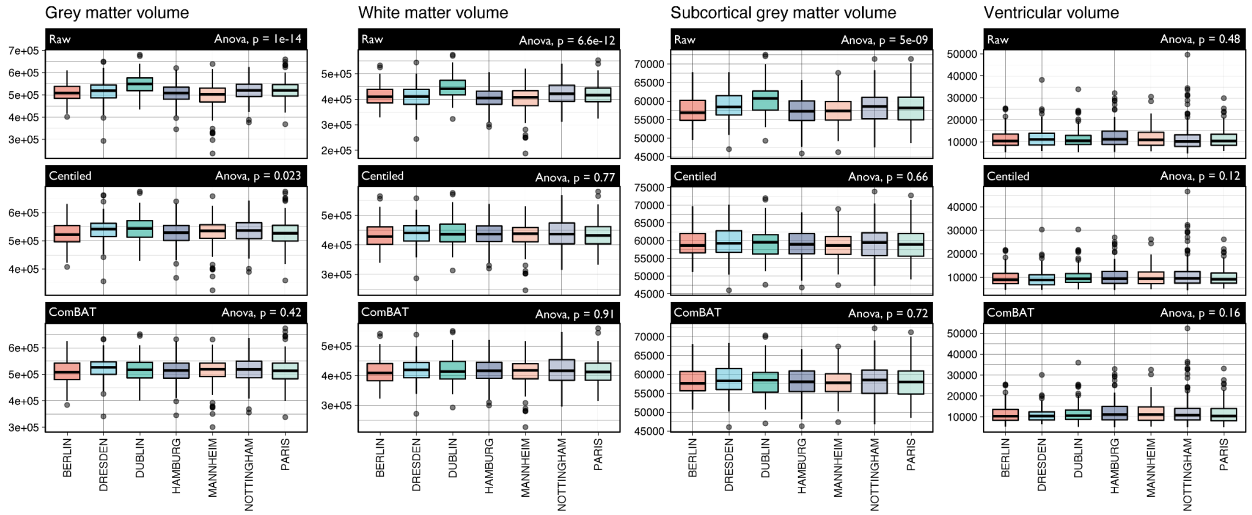


***Fig. S5.2.6. Raw volumetric data and centile scores for female participants from the IMAGEN cohort.*** *Top row shows raw volumetric data across the different sites included in IMAGEN, the middle row shows centile normalised data by GAMLSS, and the bottom row shows data normalised using ComBAT. ANOVA uncorrected P-values refer to one-way analyses of variance across sites for each individual phenotype. ComBAT and GAMLSS are both able to substantially mitigate batch effects in multi-site MRI data from the IMAGEN study (as well as the ABCD study).*

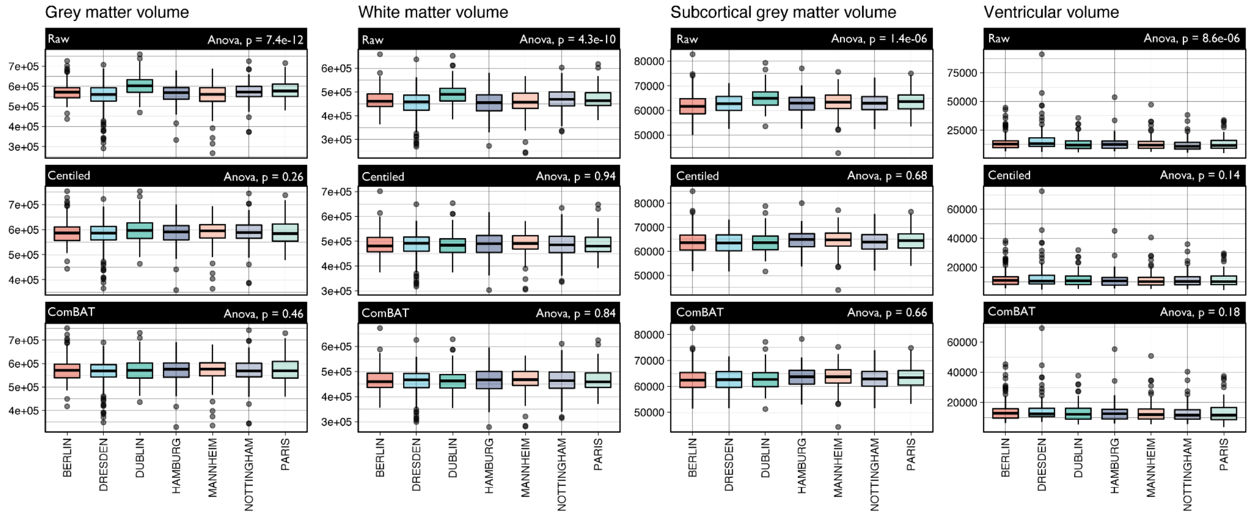


***Fig. S5.2.7. Raw volumetric data and centile scores for male participants from the IMAGEN cohort.*** *The top row shows raw volumetric data across the different sites included in IMAGEN, the middle row shows centile normalised data by GAMLSS and the bottom row shows data normalised using ComBAT. ANOVA uncorrected P-values refer to one-way analyses of variance across sites for each individual phenotype. Bars are coloured by site. ComBAT and GAMLSS are both able to substantially mitigate batch effects in multi-site MRI data from the IMAGEN study (as well as the ABCD study).*

In short, there are pros and cons to both harmonisation strategies: ComBAT is better suited for smaller datasets, normalised distributions and multivariate phenotypes; whereas GAMLSS is well suited for large datasets, non-Gaussian distributions and univariate phenotypes. We preferred GAMLSS on the grounds of its greater scalability and flexibility to match the distributional properties of the reference data and the objectives of this project. It is beyond the scope of the present work to provide an exhaustive review on batch correction methods or to evaluate the performance of GAMLSS (or ComBAT) for correction of batch effects under all possibly relevant experimental conditions. We emphasize that our use of GAMLSS for between-site or between-study harmonisation may not be optimal for studies with small (N<100) numbers of healthy control participants per site (**SI 4.5**). In addition, GAMLSS will not mitigate study- or site-specific effects driven by ascertainment bias or variability in diagnostic criteria between sites. Adaptations of ComBAT have been proposed for batch effect correction of multi-site data where such factors are likely to be problematic[55]. However, these approaches may not be suitable for harmonisation of datasets with partially or totally non-overlapping age-ranges, as required for integration of primary studies to estimate brain charts over the entire lifespan.

Finally, while we principally modeled lifespan brain trajectories with primary study (not scanning site) as “the batch” to be corrected by GAMLSS or ComBAT, we also modelled trajectories treating both study and site as batch effects. The results were nearly identical for study-batch corrected or study-and-site batch corrected trajectories (all $r^2 > 0.99$ for both parametric [Pearson’s] and non-parametric [Spearman’s] correlations). This near-perfect agreement is likely due in part to the partitioning of variation. The study and study-site random-effects covariance structures are both dominated by the sigma-component, i.e., phenotype variance. Essentially once we increase the resolution of batch effects to study-and-site specific random-effects, we have reduced the sample size to estimate each random-effect and hence this uncertainty is unable to compete with the raw observation noise (captured by the sigma-component). In an ideal scenario one would use a site within study nested random-effects structure. However, the co-dependence of variation in processing pipelines, MRI acquisition parameters, lifespan coverage, and small site-specific sample sizes, combined with the inherent observation noise, means such a covariance specification is unlikely to be viable with the currently available data (also, GAMLSS does not currently support nested covariance structures).

#### 6. Cohort effects

As is the case for traditional growth charts, reference norms for brain charts may change over time, underscoring the need for “front work” on constructing normative reference models that are adaptive to future trends. Our choice of GAMLSS as the preferred modeling framework was in part motivated by its ability to provide a flexible and scalable basis that could support ongoing updates to the reference data. Likewise, our effort to share these models on an interactive web-platform ([www.brainchart.io](http://www.brainchart.io) & [https://github.com/ucam-department-of-psychiatry/Lifespan](https://github.com/ucam-department-of-psychiatry/Lifespan)) was also motivated by the likely need for continuous updates to the reference dataset as and when more MRI data become available.

To assess the potential risk of cohort effects, or population norms shifting over historical time and biasing estimation of centile scores in future, we used a single (NIH) study already included in our aggregated dataset, which collected data from 1991 onwards in a constrained age range (5–25 years; N=1,468 scans). While MRI is a comparatively novel methodology (~30 years), it is possible that there may be systematic cohort effects within studies that have sampled individuals over prolonged periods of time[56], or between measurements aggregated in different age bins at different times. To quantitatively assess this possibility and the robustness of our procedures and results against such cohort effects, we analysed this NIH study containing longitudinal scans collected over two decades, from 1991 to 2011. During this time there were multiple upgrades to the hardware and software, but the core system remained a 1.5T GE Signa platform throughout:

| Label | Scanner ID | Description | Date of upgrade |
|---|---|---|---|
| 1 | S1-1 | GE Signa 1-1 (Hardware 1) | 6/9/90 |
| 2 | S1-2a | GE Signa 1-2a (Hardware 2--Software a) | 3/19/02 |
| 3 | C1-1 | CRADA magnet upgrade (Hardware 1) | 12/16/03 |
| 4 | C1-1b | CRADA magnet (Hardware 1--Software b) | 5/15/07 |

We found no evidence for significant variation of centile scores on any of the 4 cerebrum tissue volumes as a function of year-of-scanning or between these four major eras defined by the upgrade history of the NIH study scanning platform (**Fig. S6.1-2**). Thus, there was no clear evidence of cohort effects in one of the few large studies to have sustained scanning over a long period of time, and there was no evidence of measurement biases related to technical development of image analysis software that potentially could contribute to cohort effects in large, aggregated MRI datasets. However, the ongoing technical development of MRI scanners and image analysis software, as well as the possibility of more general secular trends in brain growth over time, mean that the risk of cohort effects should nonetheless be iteratively re-evaluated as the currently available reference dataset continues to be updated in the future.

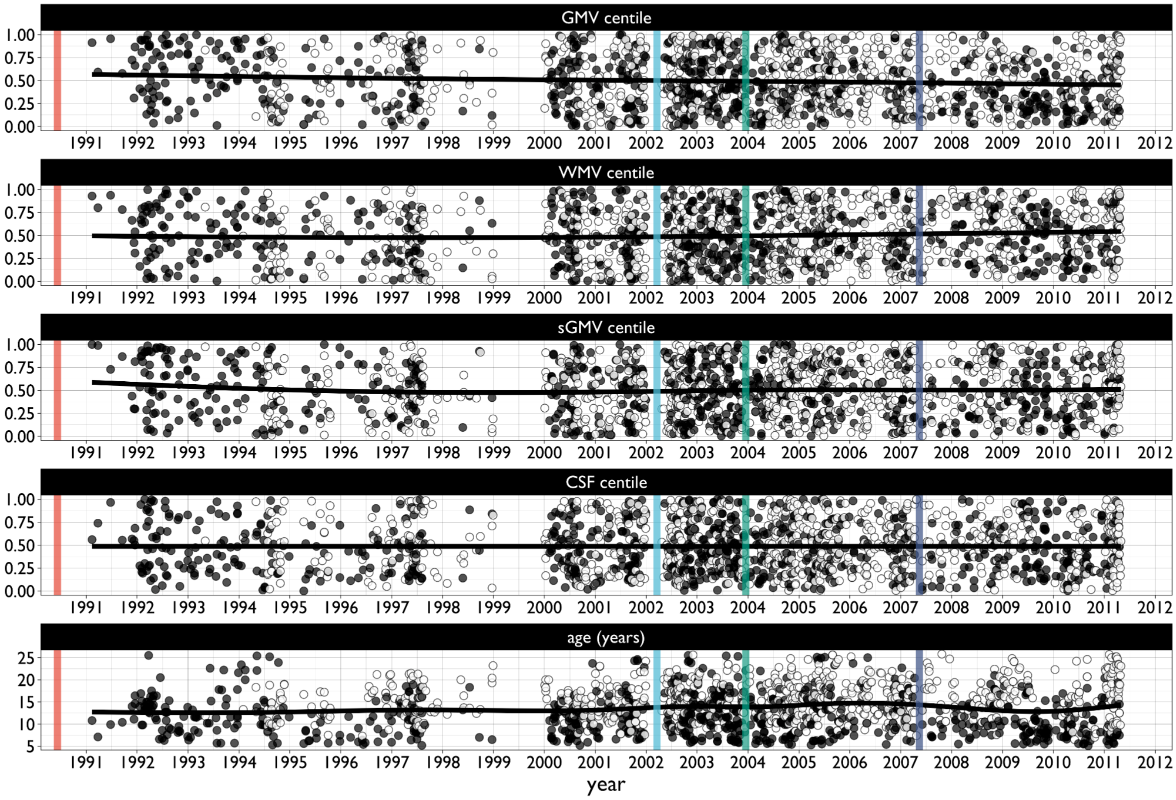


***Fig. S6.1. Assessment of potential cohort effects based on date of scanning over two decades.*** *The longitudinal study at the National Institutes of Health (NIH) contains N=1,468 longitudinal scans (N=788 subjects) collected across the age range 5–25 years and over the historical period 1991–2011. Scatterplots represent individual centile scores (y-axis), ordered by date of scanning (x-axis), for each of the four cerebrum tissue volumes (top four rows); and age at scan (y-axis) versus date of scanning (x-axis) (bottom row). Lines represent locally-weighted regression lines (LOESS regression) for qualitative analysis of possibly non-linear cohort effects on brain phenotypes or age at scanning. Filled circles denote baseline scans, empty circles denote follow-up scans in this longitudinal dataset; vertical lines indicate the timing of scanner upgrades over the course of the study (see also* ***Fig. S6.2****).*

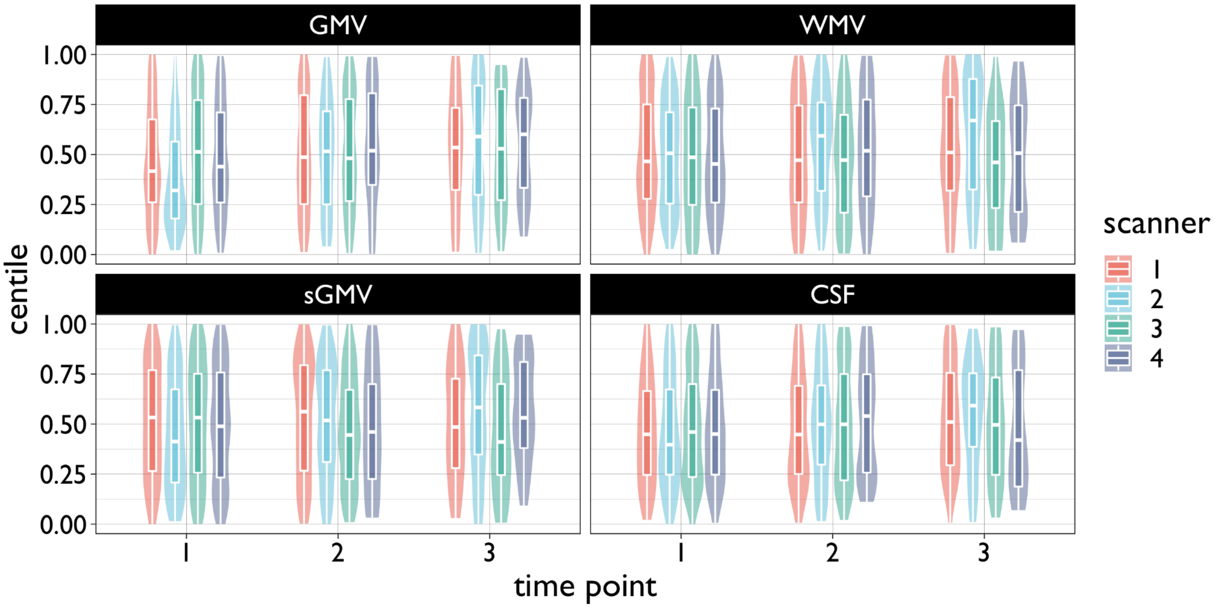


***Fig. S6.2. Assessment of potential cohort effects related to scanner upgrades in the NIH longitudinal study.*** *Centile scores for all four cerebrum tissue volumes estimated at baseline (time point 1) or two follow-up assessments (time points 2 and 3) were assigned to one of four epochs partitioned by the timing of upgrades to the 1.5T MRI scanner used for data collection. Box-violin plots show the distribution of centiles, and the range (whiskers) and 25th, 50th, and 75th percentiles of the centile distributions (boxes). Linear mixed effect modeling demonstrated no evidence of a significant effect (t=-1.577, P=0.115). This analysis was restricted to time points with N > 100 subjects.*

### Supplementary Analyses

In addition to sensitivity analyses aimed at validation of the modelling framework, primary output and out-of-sample stability and reliability, we conducted several analyses that extend the work presented in the main manuscript.

#### 7. Extended global cortical phenotypes

In addition to the principal results based on cerebrum tissue volumes, we also developed brain charts, based on the same GAMLSS modeling strategy, for other global phenotypes. This set of extended phenotypes including mean cortical thickness (CT) and surface area (SA) which we refer to as 'cortical geometric phenotypes'. Geometric cortical phenotypes are likely to be useful in addition to, and complementary to, cerebrum tissue volumes that can theoretically be derived from MRI data without cortical surface reconstruction and are expected to be more robust to estimation in MRI data of marginal image quality. CT and SA were estimated from a subset of the representative dataset for which we had access to quality-controlled, surface-reconstructed MRI data suitable for cortical geometry ($N_{total}$=97,980, $N_{unique\ subjects}$= 75,889 and $N_{unique\ CN\ subjects}$= 59,643 for SA and $N_{total}$=97,933, $N_{unique\ subjects}$= 75,847 and $N_{unique\ CN\ subjects}$= 59,599 for CT; see **ST1.6-1.9** for demographic and other details for each study included; see also **SI19 "Primary dataset descriptions"**). Another extended phenotype was total cerebrum volume (TCV)—a composite metric defined as the aggregate volume of GMV and WMV (measurable in $N_{total}$=121,650 and $N_{unique\ subjects}$=98,724). TCV estimated by combining all 4 cerebrum tissue volumes, i.e., inclusive of sGMV and CSF as well as GMV and WMV, was highly similar to TCV = GMV+WMV (*r*=0.99); but a smaller subset of the reference cohort had analysable data for all 4 tissue classes.

##### 7.1 Model optimisation

CT, SA and TCV were all analysed using the same GAMLSS modeling strategy (see **SI1-6**) as we originally used for growth charts of cerebrum tissue volumes. For 2 extended phenotypes (TCV and SA), optimal GAMLSS model specification converged on 3rd order polynomial fits for $\mu$ and $\sigma$ and a 2$^{nd}$ order polynomial fit for mean thickness on the $\mu$ and $\sigma$terms (**Fig. S7.1.1**). We found that fractional polynomial modelling for $\nu$ resulted in model instability, i.e., the GAMLSS model specification process did not converge on an optimal parameterisation, and these terms were therefore not included as fixed-effects of time in the GAMLSS model. On the other hand, model specification endorsed the inclusion of study-specific random effects on both mean and variance ($\mu$ and $\sigma$ terms) of all extended phenotypes. We note the discontinuity in the raw, non-centiled CT data that is most likely driven by a combination of brain tissue-related changes in the MRI signal due to the ending of the phase of GMV:WMV differentiation (and peak velocity in WMV) and technical differences in pre-processing of images around this age. Two-three years is often used as the cutoff to determine application of different pre-processing pipelines, e.g., infant FreeSurfer versus adult FreeSurfer. However, multiple studies and associated pre-processing pipelines were extended across this age-range in our aggregated dataset (and we also observed this discontinuity within studies that were processed homogeneously across the 2-3 year span), hence we consider it is unlikely that the transition between pre-processing pipelines can entirely explain the discontinuity in CT. We also emphasise that while there is a relative paucity of data in the <2yr age-range compared to other epochs, and for extended global MRI phenotypes (including CT) compared to cerebrum tissue volumes, the current trajectories are based on the largest early developmental dataset reported to date. Future work will hopefully be able to more precisely disambiguate to what extent this discontinuity is driven by MRI signal changes representative of maturation of brain tissue composition and to what extent the discontinuity represents technical factors such as adoption of infancy-specific pre-processing pipelines.

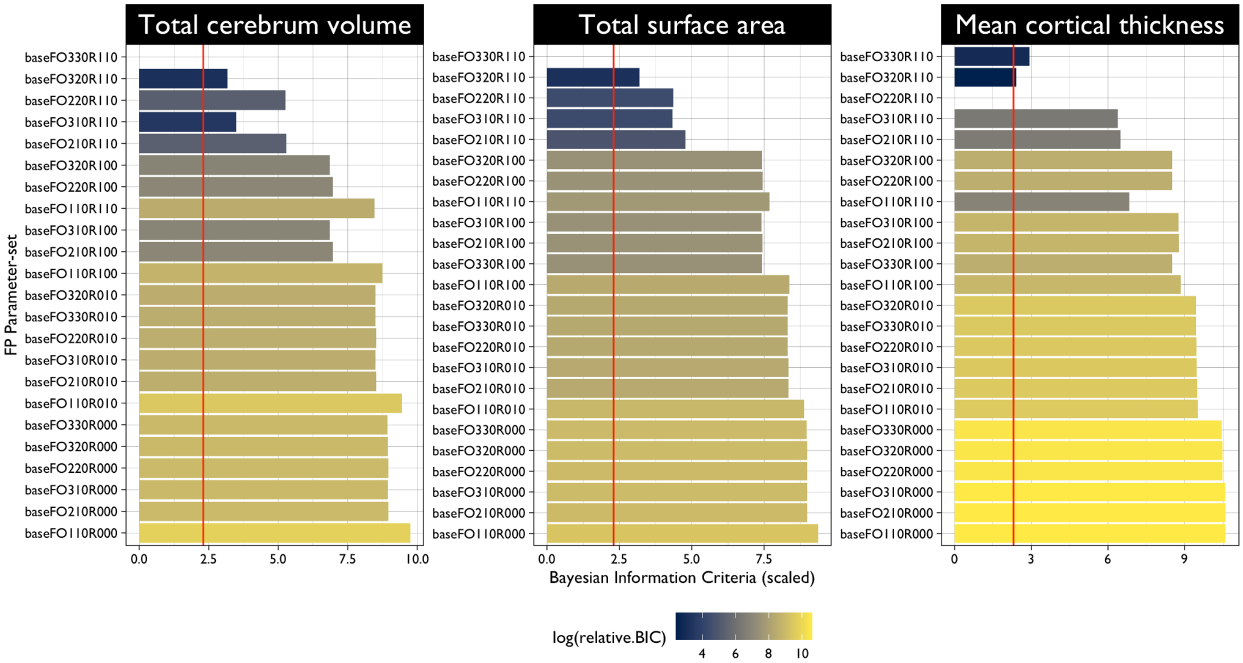


***Fig. S7.1.1 Optimization of GAMLSS model specification by analysis of the Bayesian information criterion (BIC) for multiple possible models on the generalised gamma distribution.*** *For each of three global metrics – TCV, total SA and mean CT – we compared model fit across multiple possible models combining fractional polynomial fixed effects of time and study-specific random effects on statistical moments of MRI phenotypes. Model goodness was quantified by the Bayesian information criterion (BIC) with greater log BIC indicating better-fitting models. Here log BIC is plotted relative to the best-fitting model with lowest BIC for each combination of fractional polynomials and random effects for which the model converged. All BIC values were scaled to the lowest value for the set of models fitted to each cerebrum tissue volume (log-scored difference to the lowest scoring model). For all phenotypes the best-fitting model included 3 fractional polynomials for* $\mu$*; and for all but CT the ordering also suggested 3 polynomials for* $\sigma$. *The various models fitted are summarised by y-axis labels denoting the base fractional polynomial configuration (“baseFO”) that are structured as follows: baseFO[a][b][c]R[x][y][z], where a-c denote the number of fractional polynomials included in the age term on* $\mu$, $\sigma$, *and* $\boldsymbol{\nu}$ *respectively, and x-z denote whether a study random effect was estimated for each of the model components (1 means a study random effect was included, 0 means no study random effect was included).*

#### 7.2 Normative trajectories of extended global MRI phenotypes

Following the data-driven determination of the optimal GAMLSS specification of the number of random-effect fractional polynomials on each of the distribution parameters, normative trajectories were generated using the same framework as outlined in **SI1-6** including the same bootstrapping procedure. Briefly, we generated 1,000 bootstrap iterations with stratified (by study and sex) sampling with replacement. The figure below (**Fig. S7.2.1**) shows the mean trajectory across bootstraps with a shaded region indicating the 95% confidence intervals (across the bootstrap replicates). In addition, and analogous to our primary phenotypes, we evaluated the stability of all GAMLSS derived study specific parameters (**Fig. S7.2.2**). Again, we find that smaller studies in specific age ranges tend to have somewhat wider confidence intervals on both mean and variance parameters.

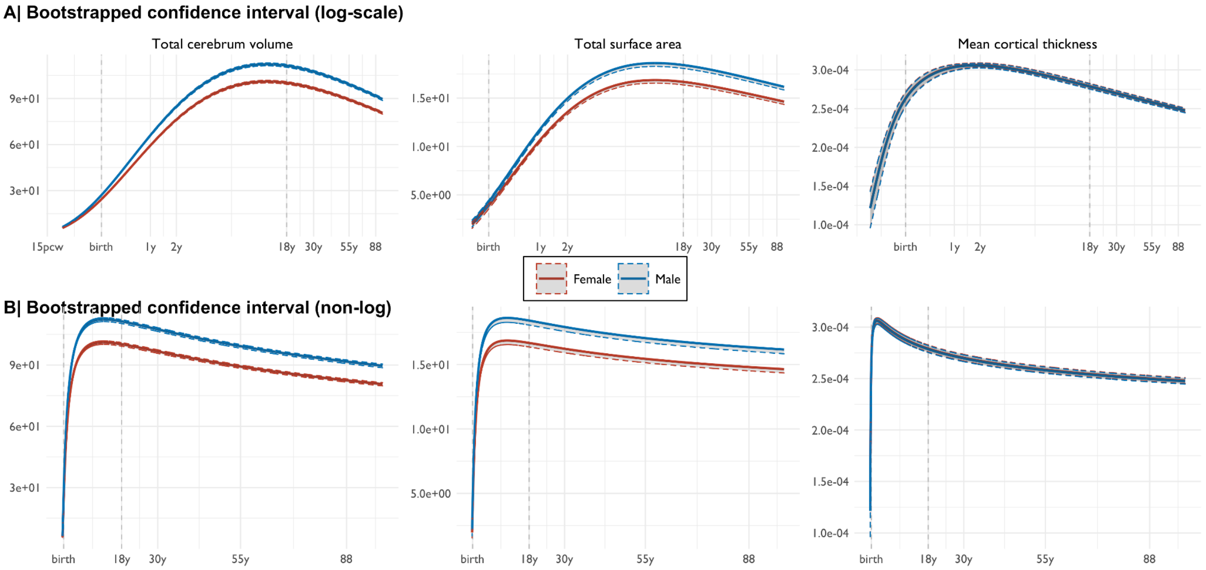


***Fig. S7.2.1. Normative trajectories of median and bootstrapped confidence intervals for three extended global MRI phenotypes, from left to right: TCV, SA and CT.*** *A | Sex-stratified curves plotted on a log scale. B | Sex-stratified curves plotted on natural scale.*

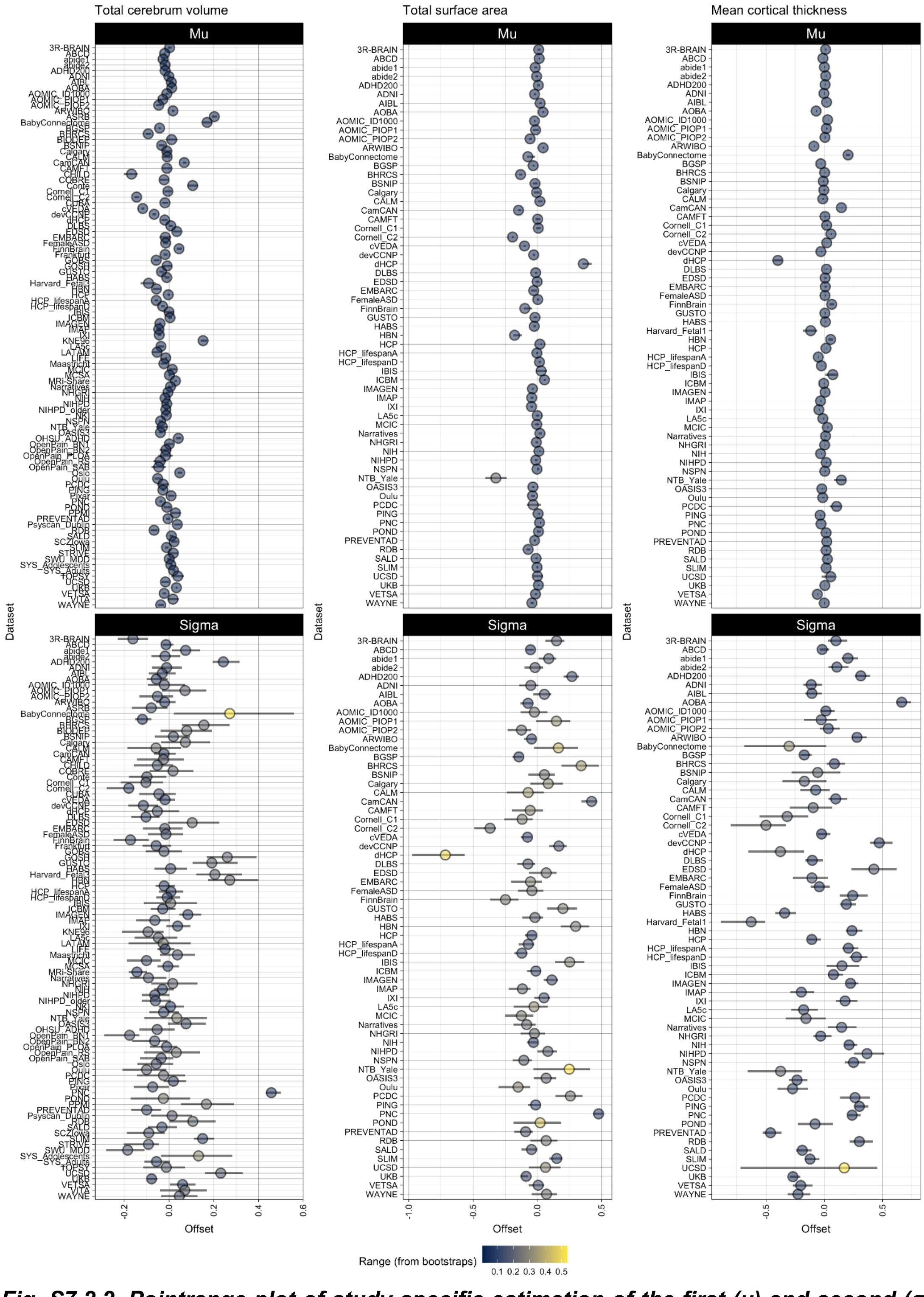


***Fig. S7.2.2. Pointrange plot of study-specific estimation of the first (μ) and second (σ) parameters of the generalised gamma fitting (where present in the selected model).*** *Confidence intervals across bootstraps (see above) are shown and dots are coloured by the range of the confidence interval. Where not observable, the confidence intervals are smaller than the size of the dots.*

Analogous to the analyses reported in SI 3.2.3 for cerebrum tissue volumes, we also examined the linear relationships between study-specific random parameters estimated in the analysis of other global MRI phenotypes and 5 demographic or technical covariates: median age, standard deviation of age, sample size, scanner manufacturer, and scanner field strength; see **Figs. S7.2.3-7.2.7.** We found only limited evidence for significant effects of any of these covariates on any of these random effect parameters.

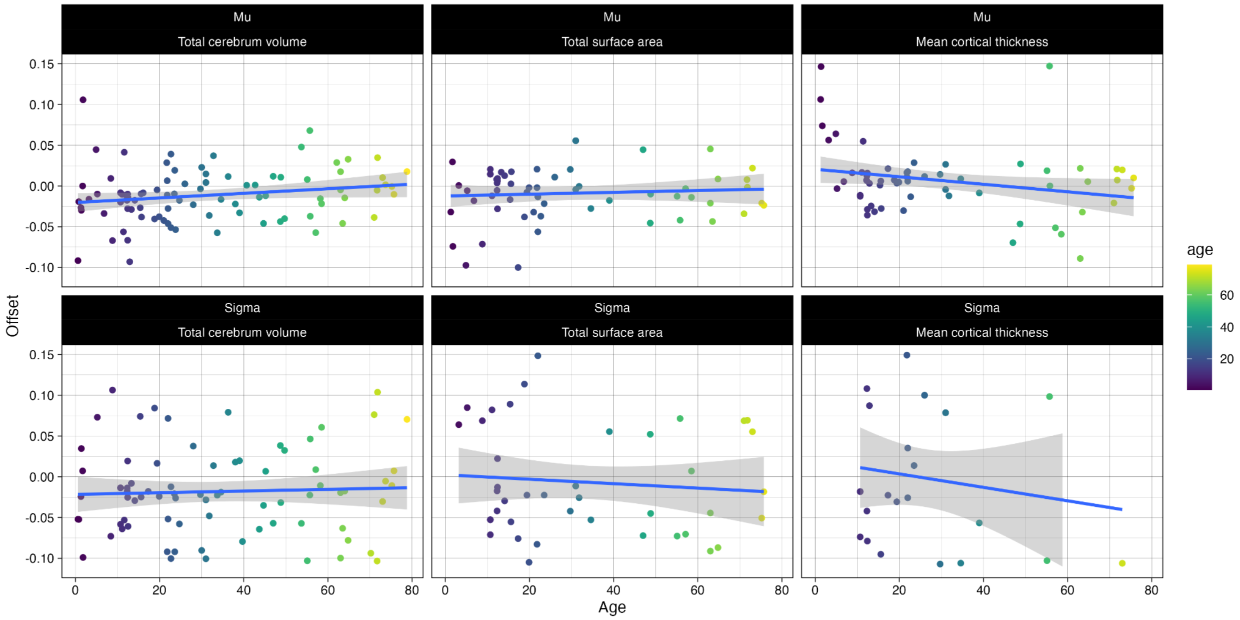


***Fig. S7.2.3. Association between median age of participants and random effect parameters estimated by GAMLSS modeling of extended global MRI phenotypes for each primary study.*** *Top row: random effects on Mu (y-axis) are plotted versus median age (x-axis) for each global MRI phenotype, left to right: total cerebrum volume, total surface area, mean cortical thickness. Fitted lines and confidence intervals indicate the strength of association estimated by linear modeling. Bottom row: random effects on Sigma (y-axis) are plotted versus median age for the same set of global MRI phenotypes. The associations between random effects and median age were not significant for any of these global phenotypes.*

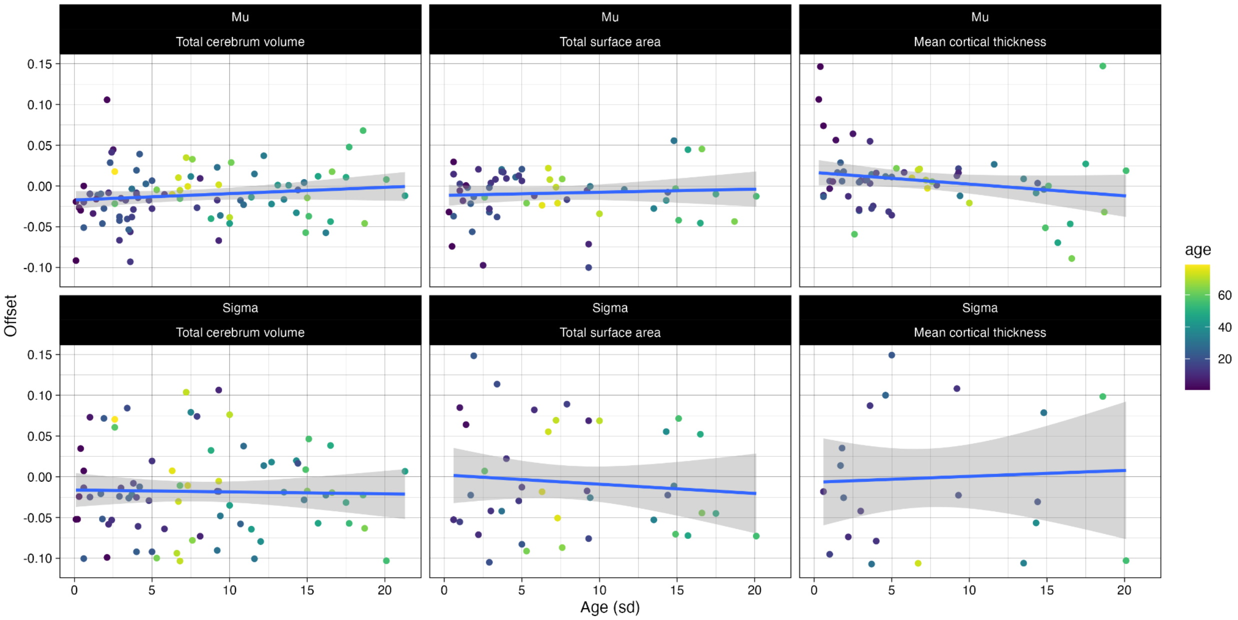


***Fig. S7.2.4. Association between the standard deviation of the age of participants and random effect parameters estimated by GAMLSS modeling of extended global MRI phenotypes for each primary study.*** *Top row: random effects on Mu (y-axis) are plotted versus standard deviation of age (x-axis) for each global MRI phenotype, left to right: total cerebrum volume, total surface area, mean cortical thickness. Fitted lines and confidence intervals indicate the strength of association estimated by linear modeling. Bottom row: random effects on Sigma (y-axis) are plotted versus standard deviation of age for the same set of global MRI phenotypes. The associations between random effects and standard deviation of age were not significant (after FDR correction for multiple comparisons) for any of these global MRI phenotypes.*

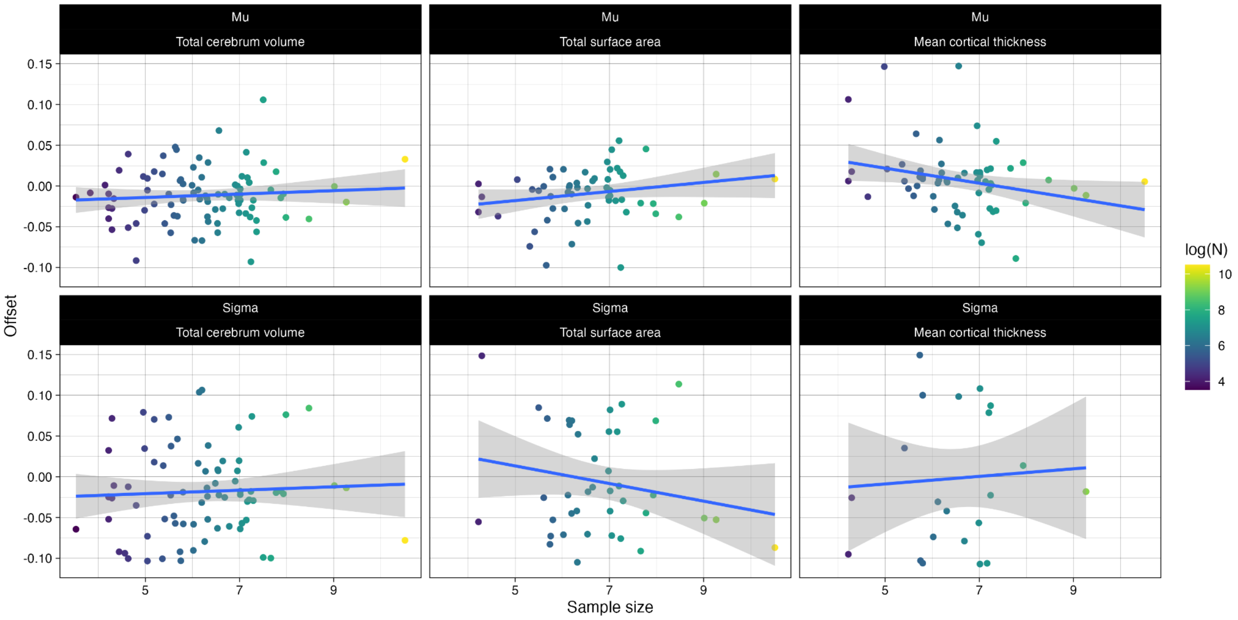


***Fig. S7.2.5. Association between sample size and random effect parameters estimated by GAMLSS modeling of extended global MRI phenotypes for each primary study.*** *Top row: random effects on Mu (y-axis) are plotted versus sample size (x-axis) for each global MRI phenotype, left to right: left to right: total cerebrum volume, total surface area, mean cortical thickness. Fitted lines and confidence intervals indicate*

*the strength of association estimated by linear modeling. Bottom row: random effects on Sigma (y-axis) are plotted versus sample size for the same set of global MRI phenotypes. The associations between random effects and sample size were not significant (after FDR correction for multiple comparisons) for any of these global MRI phenotypes.*

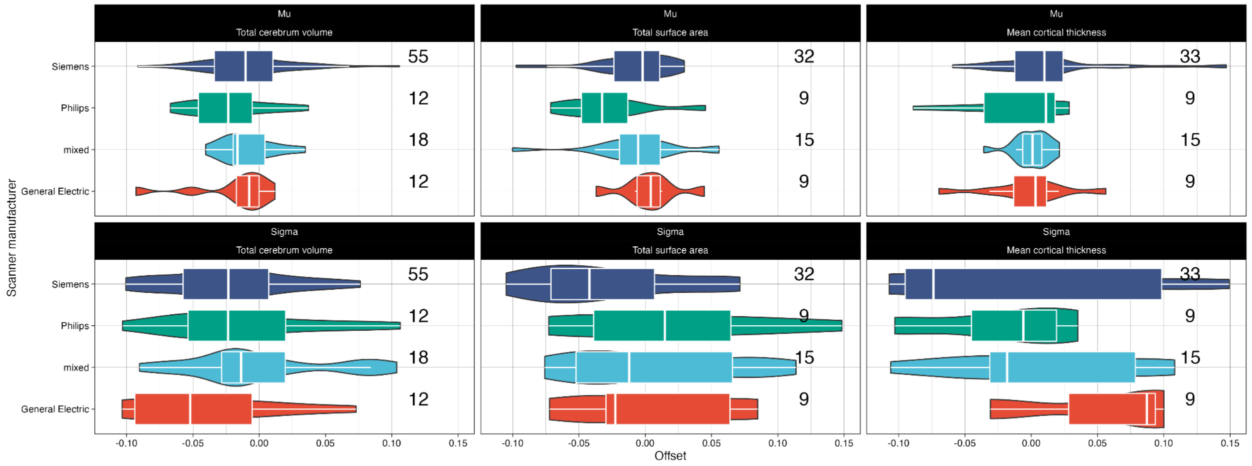


***Fig. S7.2.6. Association between the scanner manufacturer and random effect parameters estimated by GAMLSS modeling of extended global MRI phenotypes for each primary study.*** *Top row: boxplots of Mu (x-axis) are plotted for primary studies using scanners manufactured by General Electric (GE, red), Siemens (purple), Philips (green), or a mixture of different scanners (cyan), for each global MRI phenotype, left to right: total cerebrum volume, total surface area, mean cortical thickness. Bottom row: boxplots of Sigma (x-axis) are plotted for primary studies stratified by scanner manufacturer (with the same colour coding) for the same set of global MRI phenotypes. There was no evidence for a significant difference in mean random effects of primary studies using different scanners (after FDR correction for multiple comparisons) for any of these global MRI phenotypes.*

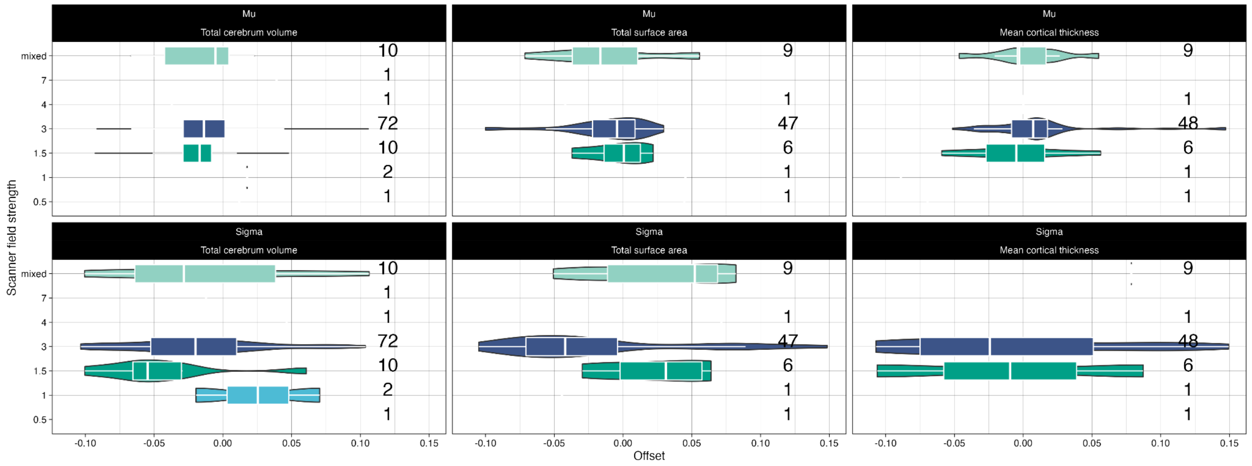


***Fig. S7.2.7. Association between the scanner field strength and random effect parameters estimated by GAMLSS modeling of extended global MRI phenotypes for each primary study.*** *Top row: boxplots of Mu (x-axis) are plotted for primary studies using scanners at different field strengths (1T, red; 1.5T, purple; 3T, green; or 7T, cyan) for each global MRI phenotype, left to right: total cerebrum volume, total surface area, mean cortical thickness. Bottom row: boxplots of Sigma (x-axis) are plotted for primary studies stratified by scanner field strength (with the same colour coding) for the same set of global MRI phenotypes. There was no evidence for a significant difference in mean random effects of primary studies using scanners*

*operating at different field strengths (after FDR correction for multiple comparisons) for any of these global MRI phenotypes.*

#### 7.3 Quality control of extended global MRI phenotypes

We applied similar quality control procedures for extended global MRI phenotypes as for cerebrum tissue volumes (**SI2**), but excluded individuals with below 2 median absolute deviation (~5%). No large effect of EI on centiles was found, nor did visual classification of a subset of raw images reveal centile differences across included QC classes—apart from in the 2 worst rated classes of images that constituted less than 5% of the data, exclusion of which did not affect models. We note, however, that especially for phenotypes extracted from the reconstructed surfaces, averaging (as in the case of mean thickness) and summing (as in the case of total surface area) likely mitigated the impact of regional reconstruction inaccuracies driven by bad data quality (see also **SI8** on regional variability).

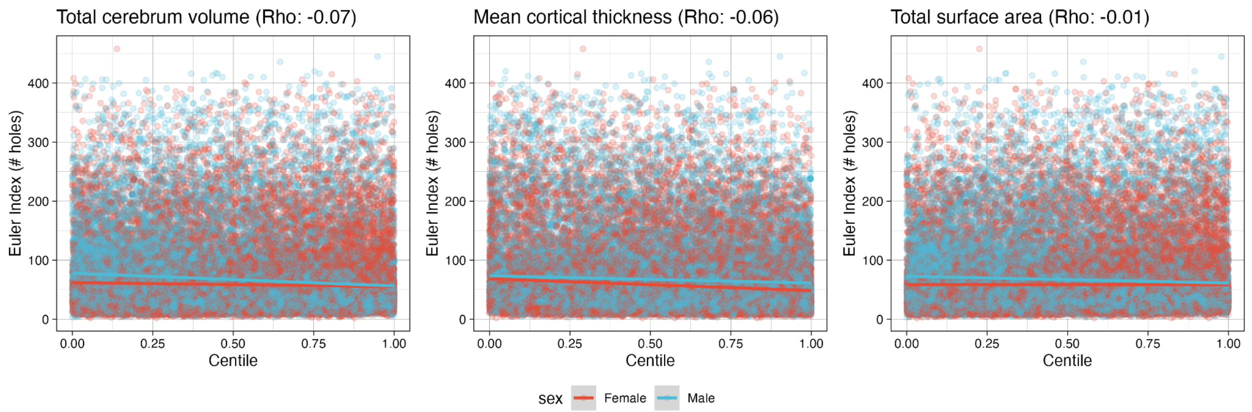


***Fig. S7.3.1. Association between EI and estimated centiles.*** *Spearman correlations between Euler Index (EI) and centiles for extended phenotypes revealed a negligible association between EI estimated image quality and derived centiles.*

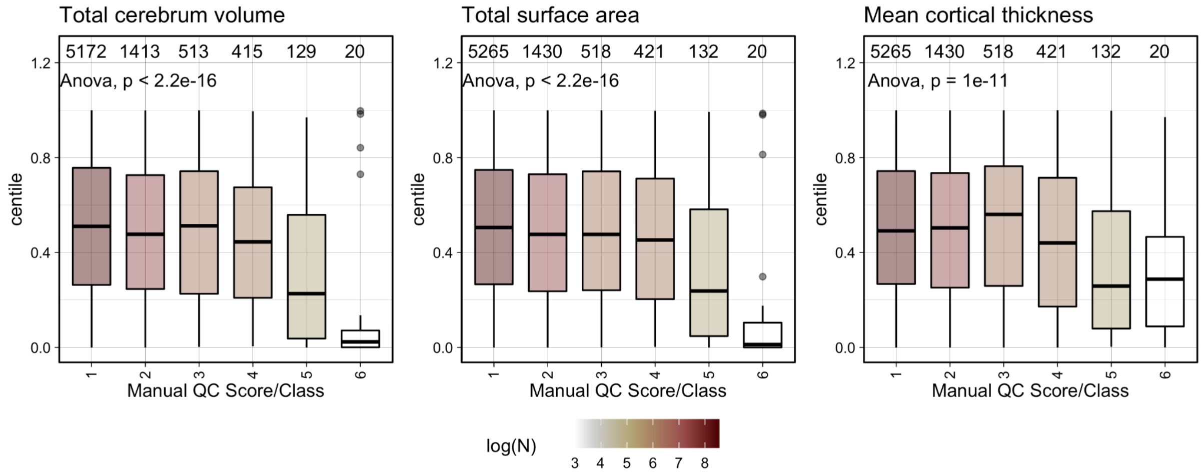


***Fig. S7.3.2. Manual quality control rating from visual inspection of raw data.*** *A small subset (~5%) of the two worst categories of raw data showed significant deviations in their estimated centiles. Excluding this subset from model estimation did not impact the model. Bars are coloured by log-scaled sample size.*

#### 7.4 Stability of out of-sample centile scoring for extended global phenotypes: LOSO analyses

Analogous to the primary four phenotypes (**SI4**) we conducted a LOSO analysis of all studies that included the extended phenotypes. While the overall variability, i.e., standard deviation across bootstrap iterations, across studies was low (<0.05 centiles), a similar pattern of increased variability of OoS estimation emerged whereby smaller studies or those in a narrow age range in a period of rapid change were slightly more variable (**Fig. S7.4.1-2**).

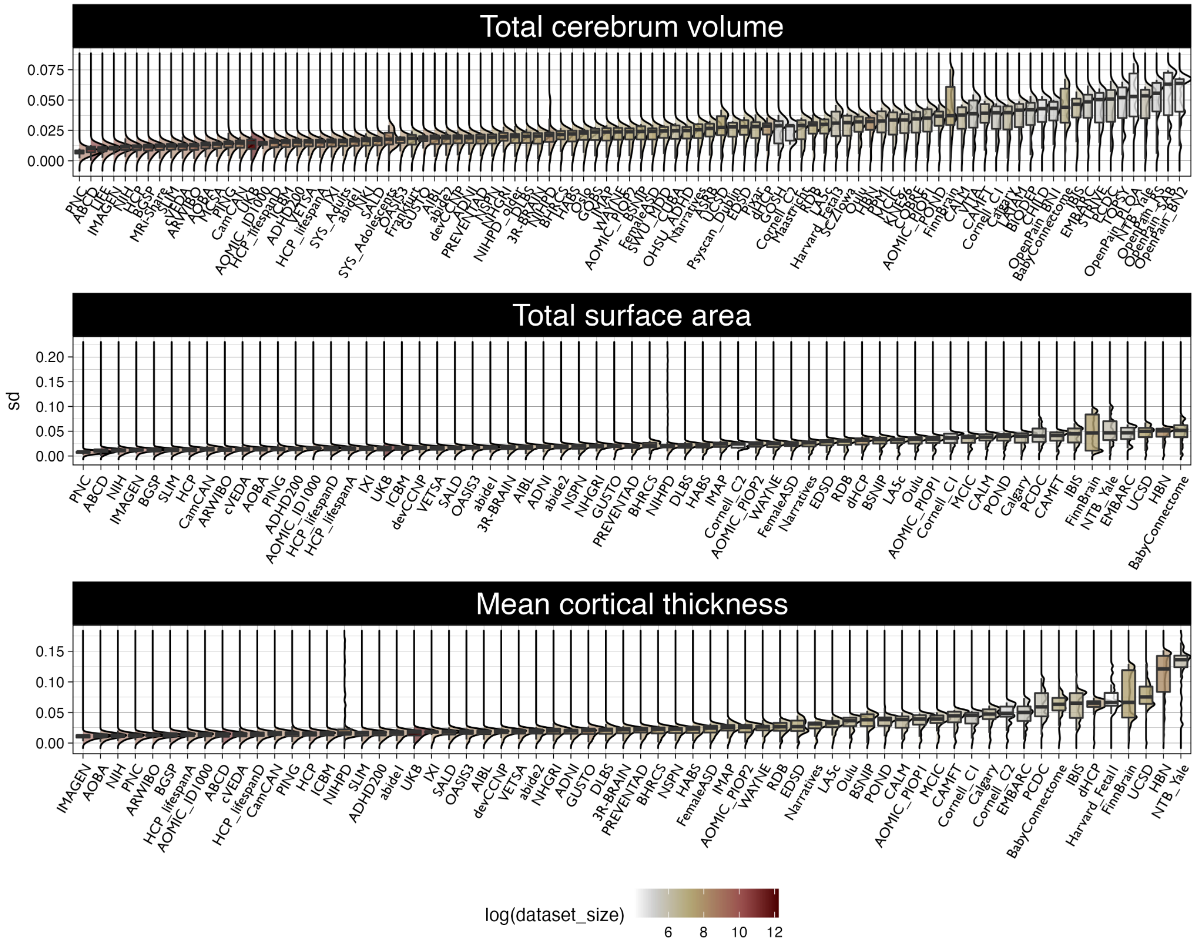


***Fig. S7.4.1. Stability of OoS estimates of centile scores on three extended global MRI phenotypes when each study was excluded from the reference dataset before bootstrapping.*** *The standard deviation of bootstrapped centile scores (y-axis) is plotted for each study (x-axis) for each phenotype, from top to bottom panels: total cerebrum volume, mean cortical thickness and total surface area. Each study- and phenotype-specific boxplot is coloured according to log sample size. For each study, we estimated the normative model leaving that study out of the reference dataset and repeated this procedure after iteratively bootstrapping the reference dataset 1,000 times. We estimated the OoS centile scores for each individual in the left-out study, normalised by each of the bootstrapped normative trajectories. This procedure allowed us to summarise the reliability of the OoS estimates of centile scores in terms of the standard deviation of the 1,000 centile scores generated for each bootstrapped resampling of the reference dataset. Studies are*

*ordered by median standard deviation of out-of-sample centile scores (small to large) indicating that scans are reliably assigned centile scores with the out-of-sample approach.*

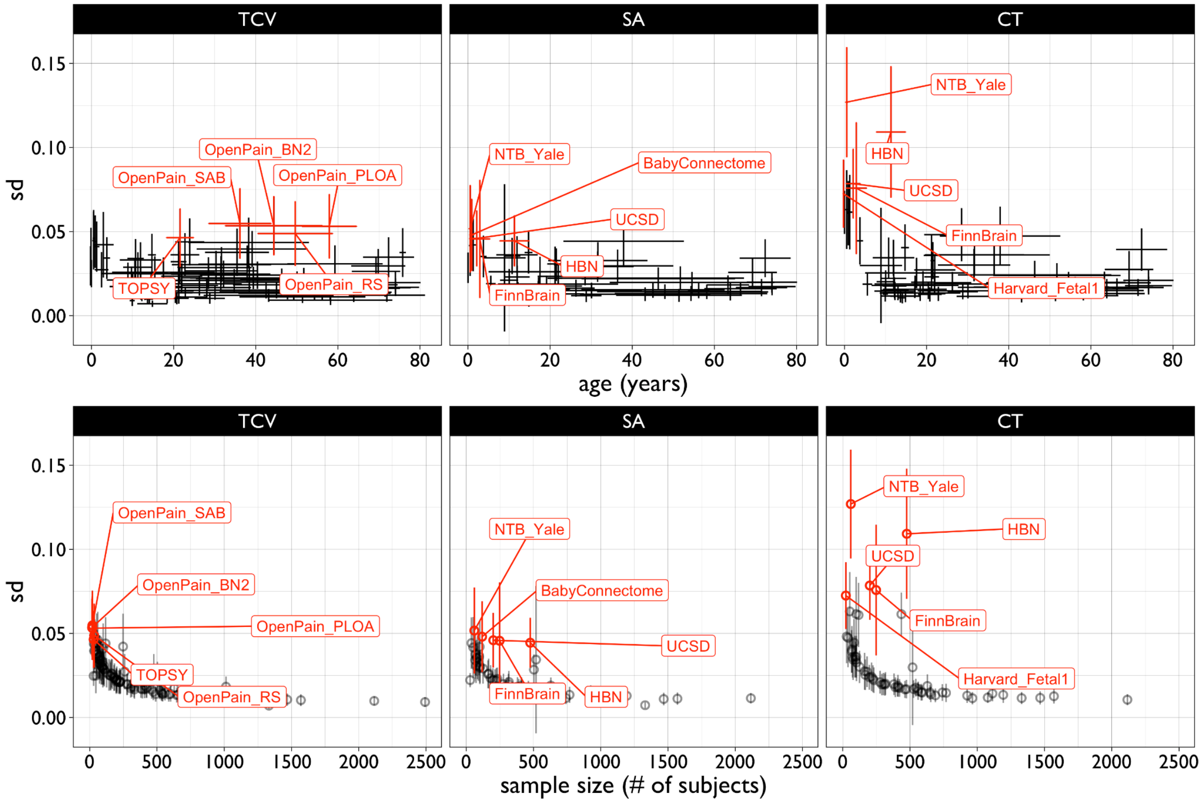


***Fig. S7.4.2. Stability of out-of-sample estimates of centile scores on extended global MRI phenotypes across age and sample size.*** *Standard deviation (sd) of individual centile scores for the extended neuroimaging phenotypes were computed across leave-one-study-out lifespan models, and plotted as a function of age (top) and sample size (bottom) for each study.*

Analogous to our assessment of bias in centile scores of cerebrum tissue volumes in **SI 4.1,** we also assessed bias of centile scores of extended global MRI phenotypes, i.e., the difference between OoS-estimated and in-sample estimated centiles. Bias was generally very low except for a few studies (i.e., CHILD, NIHPD, FinnBrain) with smaller sample size or younger participants (**Fig. S7.4.3**).

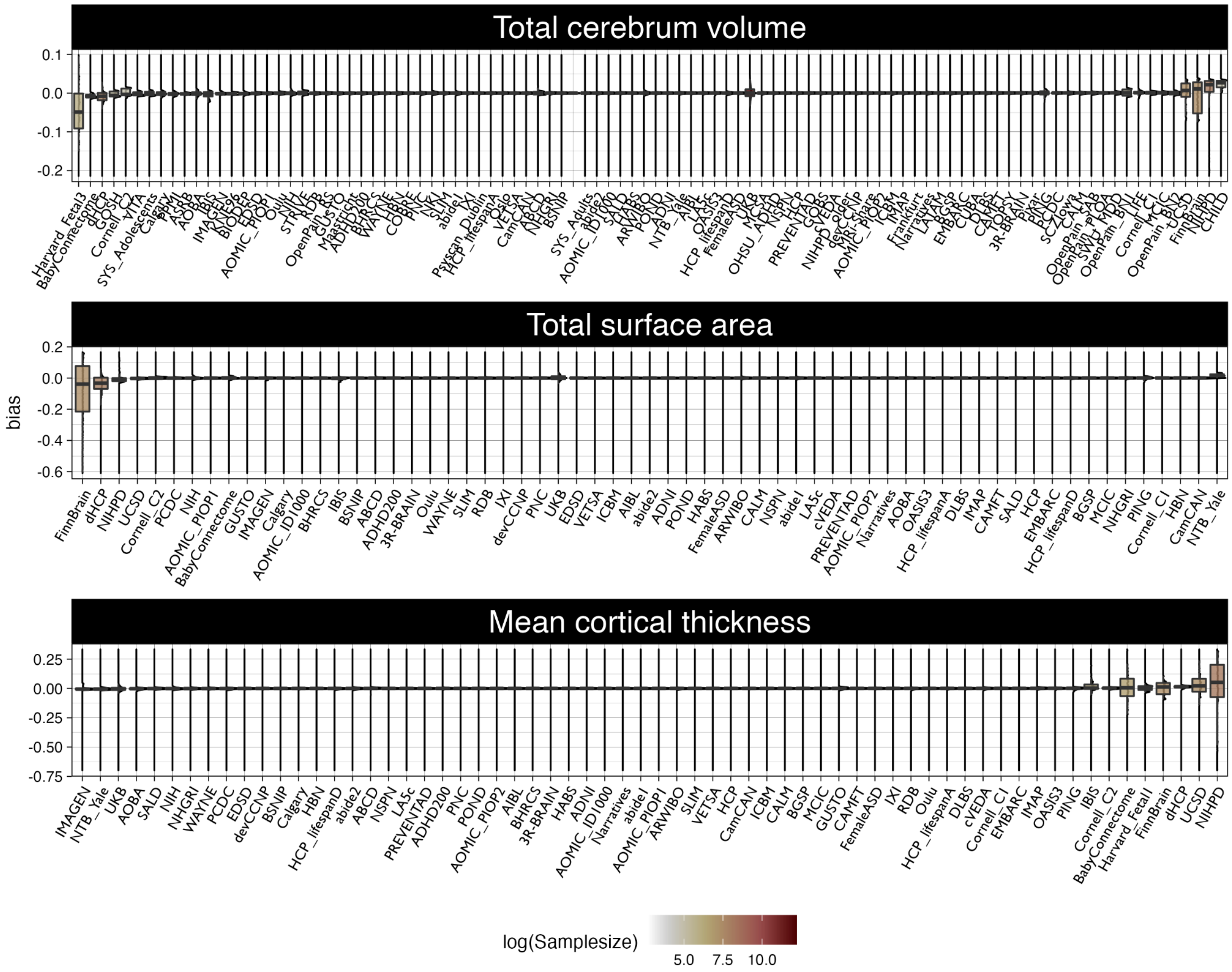


***Fig. S7.4.3. Bias of out-of-sample centile scores for extended global MRI phenotypes.*** *Each panel shows boxplots of the bias in OoS centile scores (the signed difference between OoS and in-sample centile scores; y-axis) estimated for each primary study when it was excluded from the reference dataset. Studies are ordered on the x-axis from most negatively biased (left) to most positively biased (right) OoS centile scores. Boxplots are colour-coded according to log sample size, indicating that OoS centile scores tend to be most biased for smaller primary studies. From top to bottom, panels represent the bias in OoS centile scores for total cerebrum volume, total surface area, and mean cortical thickness.*

#### 8. Regional cortical volumetric trajectories and milestones

To analyse trajectories and milestones of brain development with finer-grained anatomical resolution, we extracted volumetric information from 34 bilateral regions in the Desikan Killiany parcellation[57] for a subset of ~65,000 unique individuals (depending on the region) from birth until 100 years (**ST1.9-1.42**). Since we expected data quality to have a greater impact on the accuracy of regional volumetrics, compared to the minor impact of data quality demonstrated for cerebrum tissue volumes (see **SI2 “Quality control”**), we only included quality controlled scans with (EI < 2 median absolute deviations within each study) in these analyses, or scans that had undergone prior visual inspection. We applied exactly the same modelling pipeline to these regional volumetric phenotypes as previously applied to cerebrum tissue volumes. Briefly, we first specified the optimal combination of fractional polynomials in each term of the model using BIC, then fitted the optimal model to the sex-stratified data and to 1,000 bootstrapped resamples of the original data, and finally plotted the trajectories for the median and between-subject variability (with confidence intervals) of each regional volume. This work extends previous work on developmental trajectories of brain regional volumes in several important ways. Most prominently, for the first time, these trajectories encompass the full age-range of the lifespan, including the earliest period of development before postnatal year 2. There is evidently considerable variation between cortical regions in their developmental trajectories, but all regions show peak volume, and peak rate-of-growth of volume, in the first decade, which is compatible with our results for global cortical volume estimated in a larger and more inclusive sample.

#### 8.1 Charting development of regional volumes

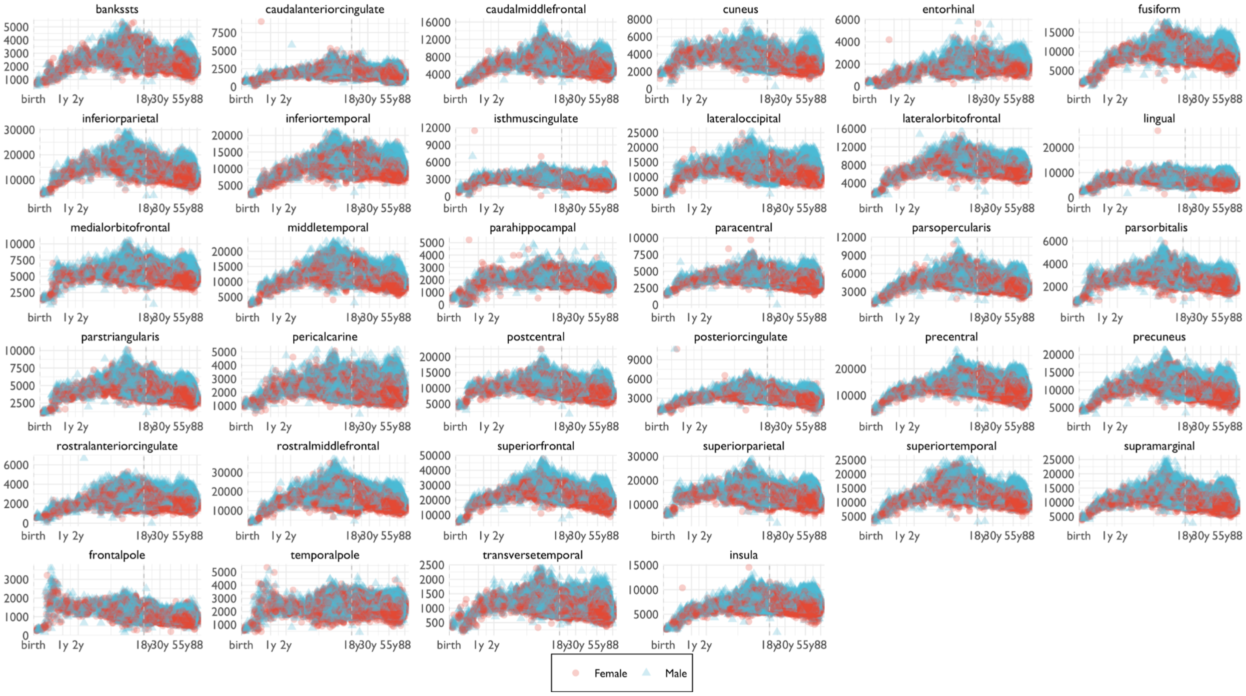


***Fig. S8.1.1. Raw regional volumetric data across the lifespan for 34 bilateral brain regions as defined in the Desikan-Killiany parcellation[57] (mm$^3$).*** *These data are analogous to the raw data depicted in* ***Figs. 1*** *and* ***2*** *for cerebrum tissue volumes and other global cortical metrics (SA and CT), respectively. Demographics for the QC'd sample available for estimation of each regional volume are provided in* ***ST1.9-1.42.***

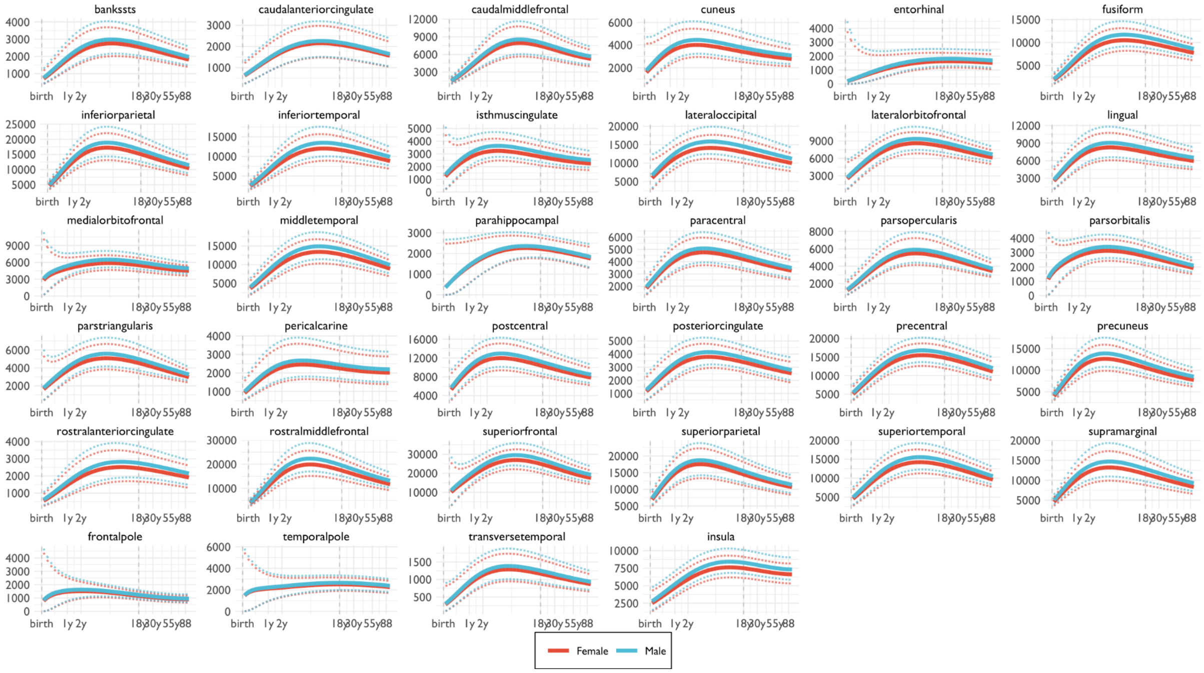


***Fig. S8.1.2. Normative trajectories of median regional volumes (and confidence intervals) across the lifespan for 34 bilateral brain regions as defined in the Desikan-Killiany parcellation[57] (mm$^3$).*** *Dotted lines indicate the 97.5% and 2.5% centile lines. These trajectories were fitted to the raw data in* ***Fig. S8.1.1*** *using the same GAMLSS model used for estimation of tissue volume trajectories, as shown in* ***Fig. 1*** *and* ***Fig. 2*** *of the main text Further details on milestones (age at peak volume and age at maximum rate-of-growth of volume) are provided for each region in* ***ST2.2*** *and* ***SI8.2*** ***"*Regional volumetric milestones"**.

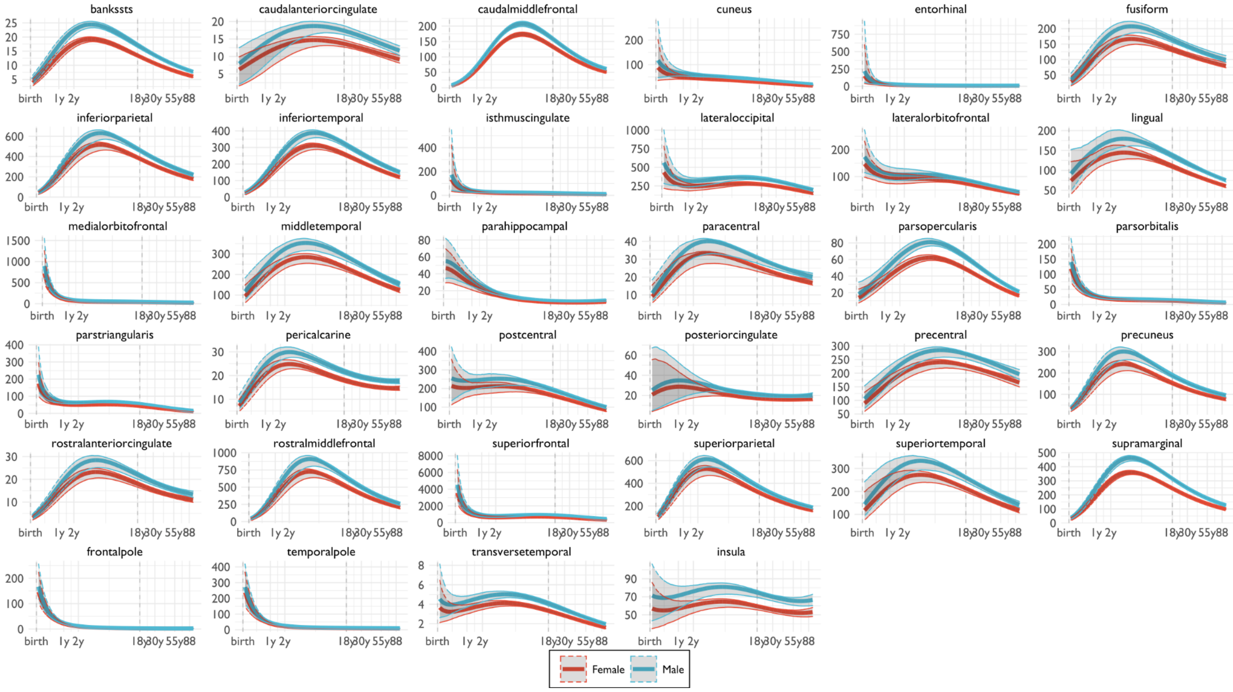

***Fig. S8.1.3. Normative trajectories of between-subject variation of regional volumes (and confidence intervals) across the lifespan for 34 bilateral cortical** regions as defined in the Desikan-Killiany parcellation[57](mm$^3$). Shaded areas represent the 95% confidence interval defined by 1,000 bootstrapped resamples of the original data, as identically done for estimation of between-subject variation in global brain phenotypes (**Figs. 1** and **2** in the main text). Further details on milestones (age at peak variation and age at maximum rate-of-growth of variation) are provided for each region in **ST2.2**.*

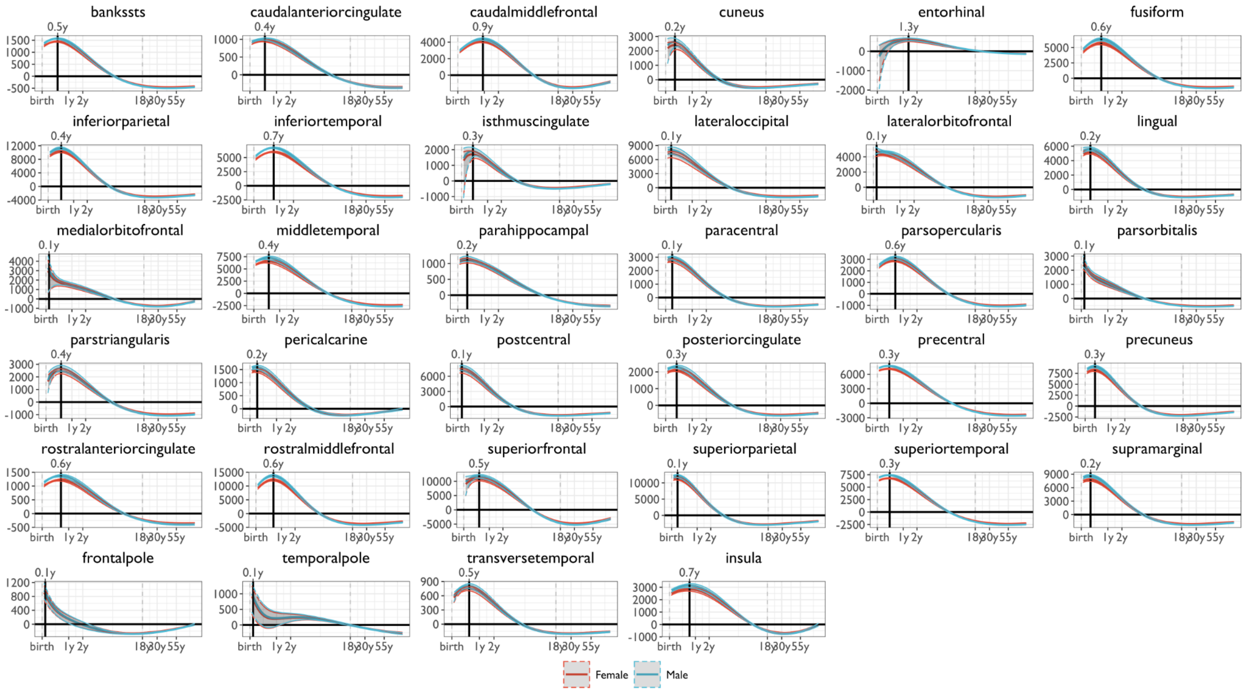


***Fig. S8.1.4. Estimated rates of change in regional volumes across the lifespan (first derivatives of the median trajectories) for 34 bilateral brain regions as defined in the Desikan-Killiany parcellation[57].** Shaded areas represent the 95% confidence interval defined by 1,000 bootstrapped resamples of the original data, as identically done for estimation of rate-of-growth curves for global brain phenotypes (**Figs. 1** and **2 in the main text)**. The numbers displayed at the top of each chart denote age at peak rate-of-growth for each regional volume and the solid horizontal line at y=0 indicates the age at which regional volumes stop growing and start to shrink. Further details on milestones (age at peak volume and age at maximum rate-of-growth of volume) are provided for each region in **ST2.2**.*

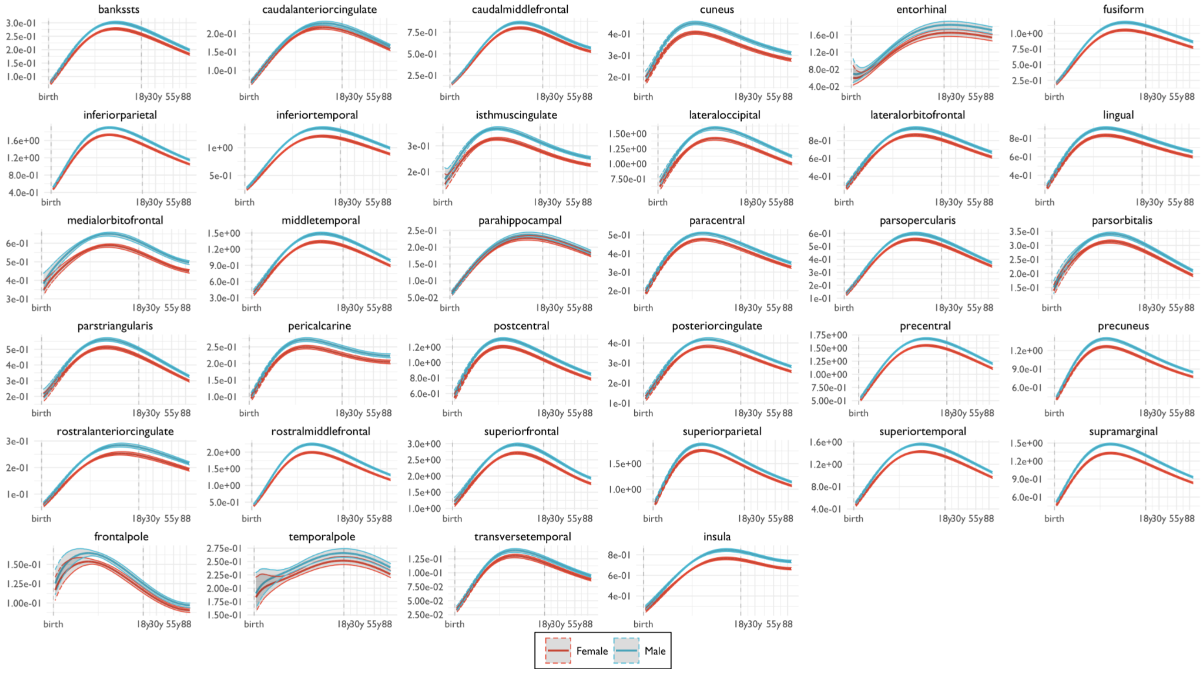


***Fig. S8.1.5. GAMLSS estimated confidence interval for model fits to regional volumes across the lifespan for 34 bilateral brain regions as defined in the Desikan-Killiany parcellation[57].*** *Shaded areas represent the 95% confidence interval estimated by 1,000 bootstraps. These results are analogous to the sensitivity analysis depicted in* ***SI3.2.2*** *and show that for most regions the confidence intervals are extremely narrow, i.e., it barely extends beyond the thickness of the lines. However, in entorhinal cortex, frontal pole and temporal pole the bootstrapped variability is considerably greater in early development, possibly indicating marginal quality of data or cortical surface reconstruction for these regions in this age range.*

#### 8.2 Regional volumetric milestones

We also estimated the developmental milestones of each region in terms of age at peak volume or peak between-subject variation, and age at peak rate-of-growth in volume or between-subject variation. **Fig. S8.2.1** shows the regions ordered by age at peak median volume alongside the bootstrapped confidence intervals of those milestones. The shaded grey bar shows the age at peak total cortical grey matter volume, with the width of the bar indicating the 95% confidence interval for that milestone. In the corresponding figure of the main text (**Fig. 2**), we excluded outlying data points, defined as age at peak volumes more than 2 median absolute deviations away from the median of the regional distribution of age at peak volume. This removed the 3 regions with the highest between-subject variability, especially in early development (entorhinal cortex, temporal and frontal poles). Perhaps unsurprisingly, both the temporal and frontal poles are regions with notoriously questionable signal quality[58]. The entorhinal cortex is the smallest cortical region defined by the Desikan-Killiany atlas and is often missing in parcellated foetal and neonatal MRI data for that reason. These results further underscore the need for conducting quality control on scanning data prior to estimation of brain charts at regional resolution. Further details on milestones (age at peak volume and age at maximum rate-of-growth of volume) are provided for each region in **ST2.2**.

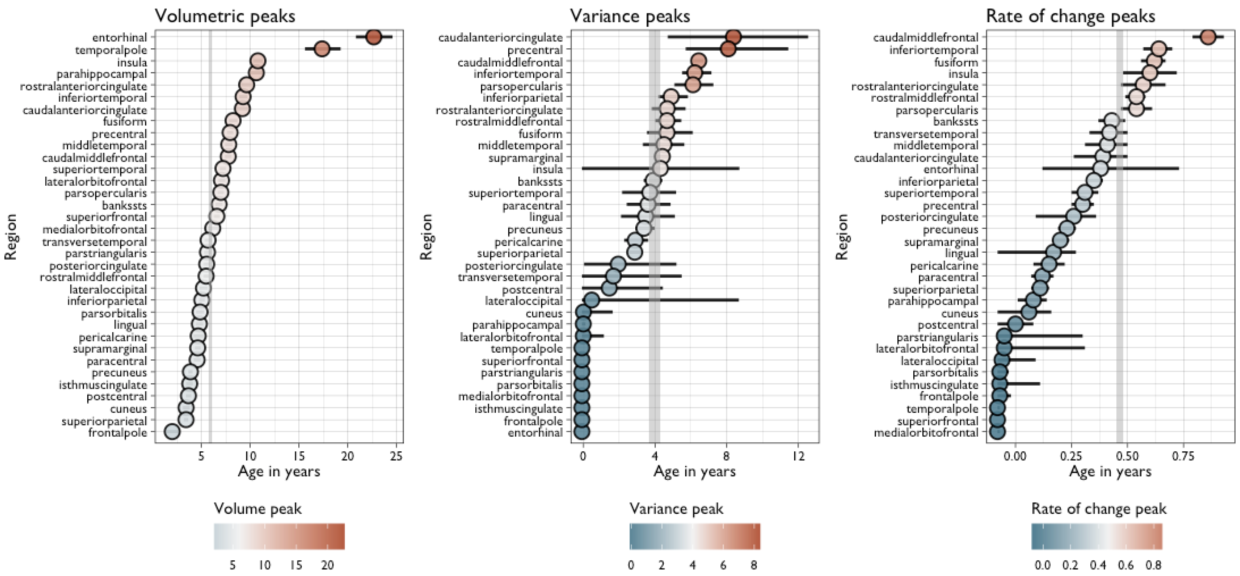


***Fig. S8.2.1. Milestones for development of regional volumes estimated from the first derivatives of the trajectories of median volume and between-subject variation for each of 34 cortical regions defined by the Desikan-Killiany parcellation.*** *Each point-range plot shows, from left to right, the age at peak volume, the age at peak between-subject variation, and the age at maximum rate-of-growth in volume. In each case, median milestones are shown in the context of their 95% confidence intervals, which are not always visible for narrow intervals. The shaded grey area in each panel shows the median and 95% confidence interval for the corresponding milestone for total cortical grey matter volume.*

To contextualise the spatial distribution of the regional volume peaks, we compared the age at peak volume to the x-, y-, and z-coordinates of the centroids of each region-of-interest in the Desikan-Killiany cortical parcellation. We observed a relatively wide distribution of age at peak regional volume, centred around the age of peak total cortical grey matter volume (grey dashed line in **Fig. S8.2.2**). Moreover, there was a clear trend for rostral and dorsal regions to have later peak volumes compared to caudal and ventral regions (**Fig. S8.2.2**). Regions in the cingulate and frontal cortices, which span greater distances (especially in rostral-caudal and dorsal-ventral dimensions), had a wider range of age peaks.

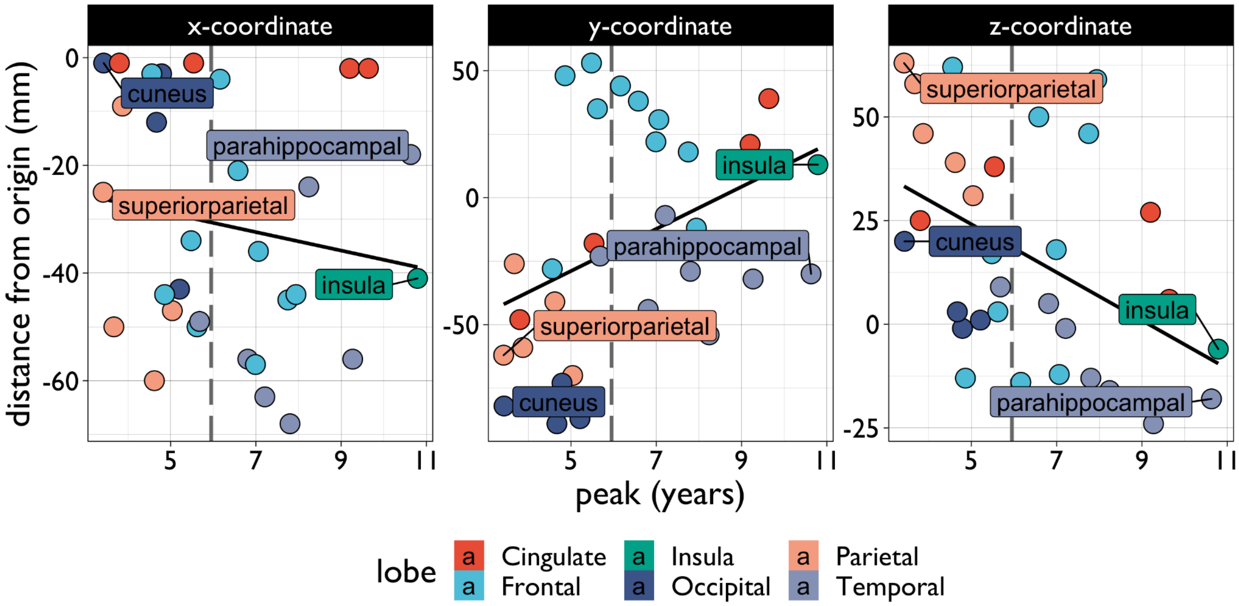


***Fig. S8.2.2. Relative timing of regional volume peak milestones, highlighting spatial gradients in timing of peak volumes.*** *Scatterplots show the relationship between age of peak volume for each region of the Desikan-Killiany parcellation (x-axes) versus x (left), y (middle), or z (right) coordinates in MNI space (y-axes). Coordinates are based on the left hemisphere, thus the interpretation (from negative to positive) is: x=lateral-to-medial, y=caudal-to-rostral, z=ventral-to-dorsal. Spearman's r was computed for each scatterplot, represented by black lines: x-coordinates were not significantly correlated with age at peak volume, r=-0.21, P=0.26; y-coordinates were positively correlated with age at peak volume, r=0.42, P=0.02; and z-coordinates were negatively correlated with age at peak volume, r=-0.50, P=0.004). Labels represent the most extreme (top two and bottom two) region peaks relative to peak total cortical grey matter volume. Grey dashed lines represent the age atpeak total cortical grey matter volume.*

### 9. Developmental windows and milestones

#### 9.1 Trajectories within developmental epochs

To clarify the developmental trajectories at different stages across the lifespan below we provide the fitted trajectories on a non-log scale for each of the lifespan windows defined by Kang et al.[59]

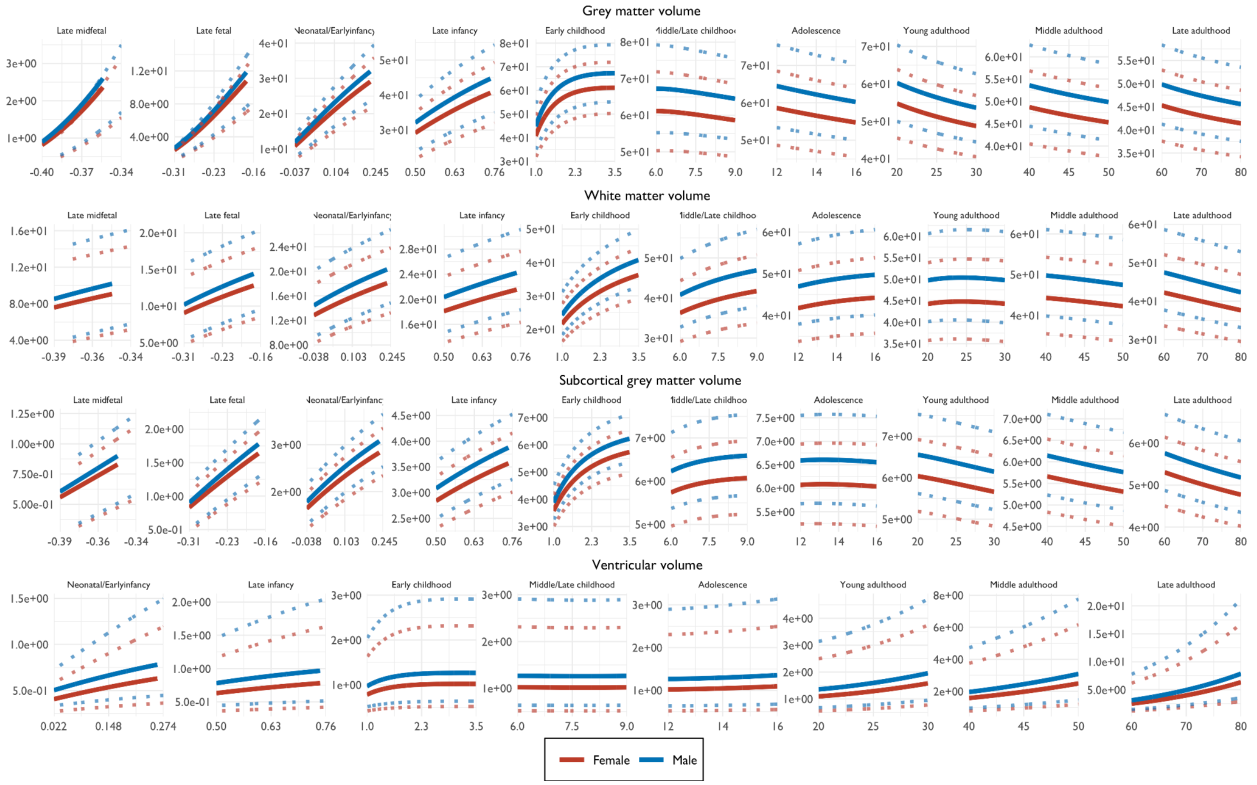


***Fig. S9.1.1. Normative trajectories of median (and 2.5-97.5% centile boundaries) of cerebrum tissue volumes.*** *As shown in main* ***Fig.*** *1, but stratified by age-defined developmental windows – from late midfoetal to late adulthood – and plotted on natural scale of age in years (x-axis) to allow further examination of the trajectory shapes over time.*

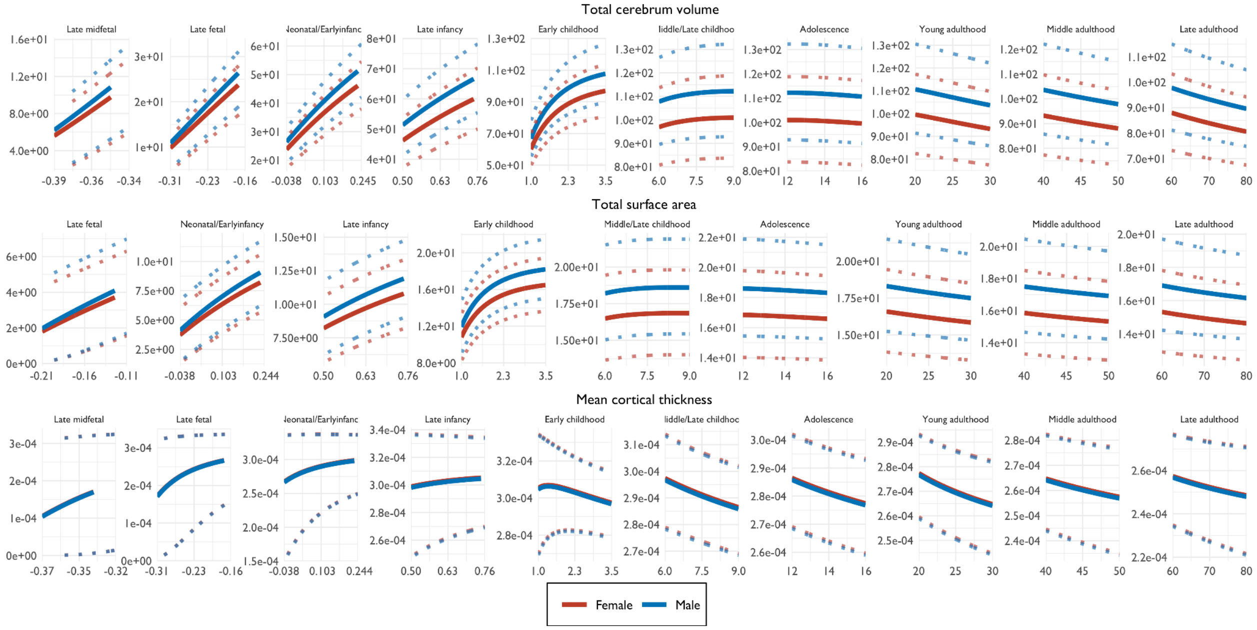


***Fig. S9.1.2. Normative trajectories of median (and 2.5-97.5% centile boundaries) of extended global MRI phenotypes stratified by age-defined developmental windows – from late midfoetal to late***

***adulthood – and plotted on natural scale of age in years (x-axis) to allow further examination of the trajectory shapes over time.***

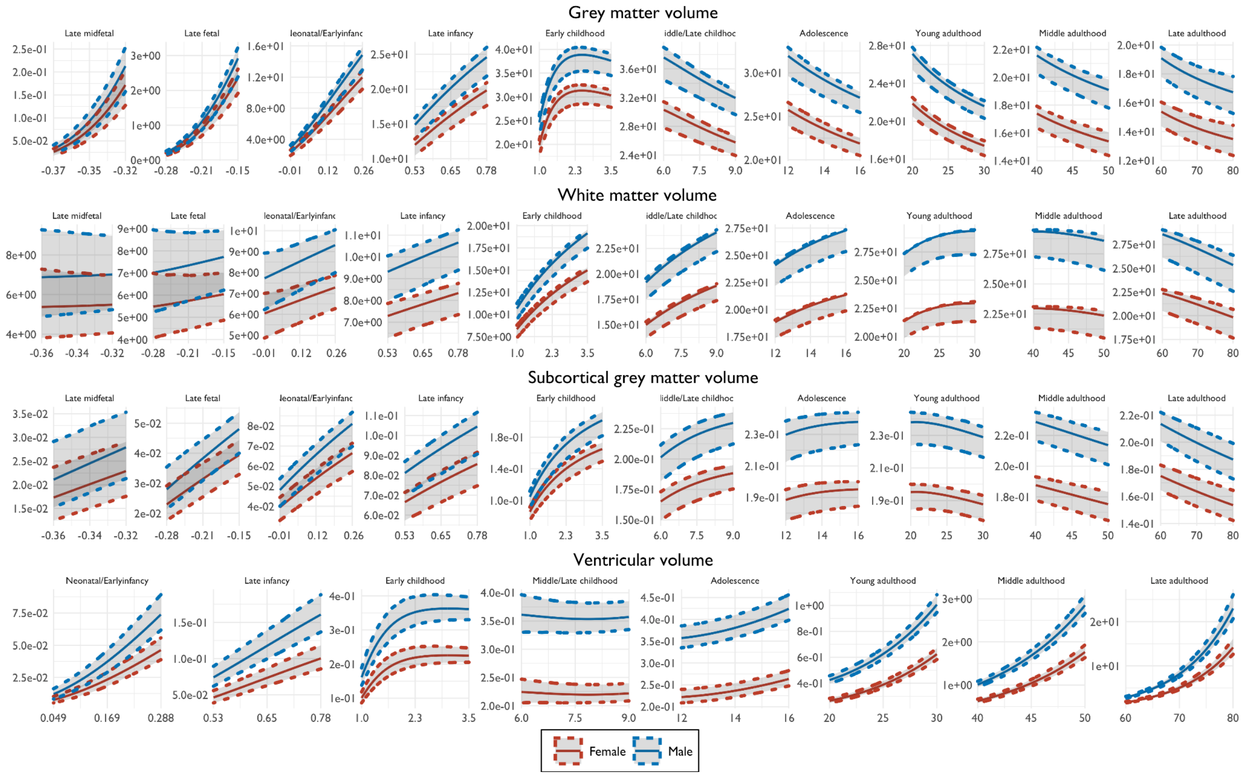


***Fig. S9.1.3. Normative trajectories of between-subject variability (and bootstrapped confidence interval) of cerebrum tissue volumes.*** *As shown in main* ***Fig.1****, but stratified by age-defined developmental windows – from late midfoetal to late adulthood – and plotted on natural scale of age in years (x-axis) to allow further examination of the trajectory shapes over time.*

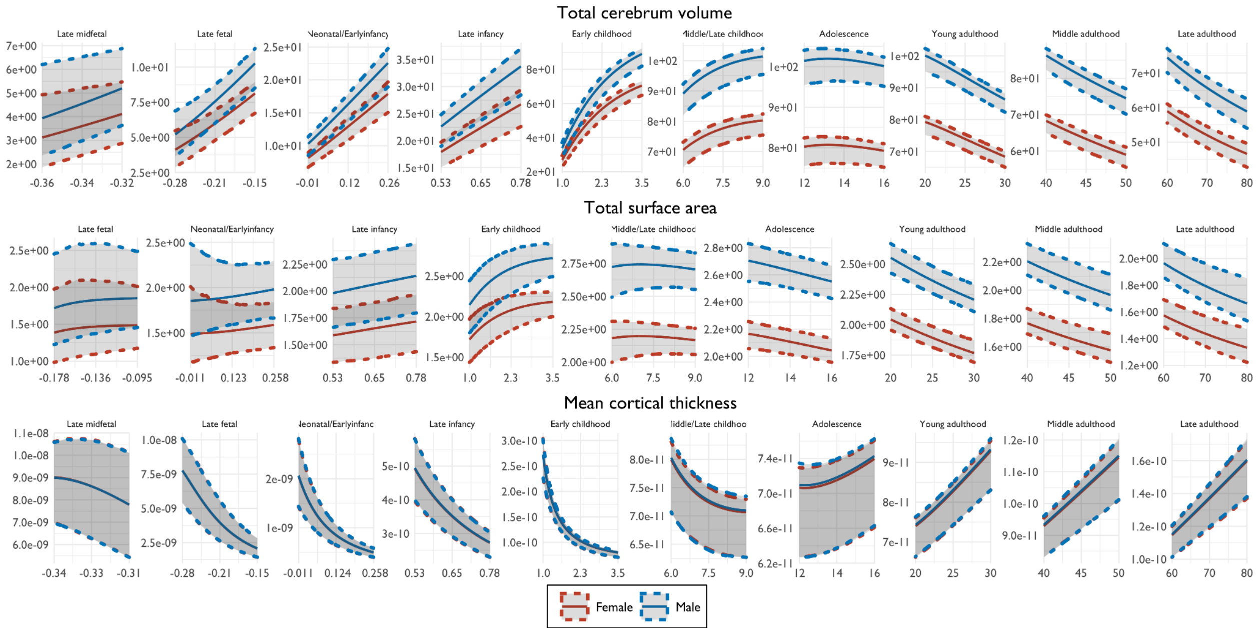

***Fig. S9.1.4. Normative trajectories of between-subject variability (and bootstrapped confidence interval) of extended global MRI phenotypes stratified by age-defined developmental windows – from late midfoetal to late adulthood – and plotted on natural scale scale of age in years (x-axis) to allow further examination of the trajectory shapes over time.***

#### 9.2 Grey-white matter differentiation

One of the emergent milestones that was delineated from the normative trajectories was the early period of grey-white matter differentiation. This was derived from the observation that these two major tissue classes show differential velocities in the early stages of development – GMV increases nonlinearly perinatally and WMV increases linearly through childhood and early adolescence. Thus, the intersection point of these two trajectories (around birth) marks the critical transition whereby GMV becomes the majority tissue compartment. Furthermore, this difference increases until ~3 years, when GMV velocity slows before peaking ~6 years. As such, the periods before and after this so-called differentiation can be characterised as grey-white consolidation and dedifferentiation, respectively.

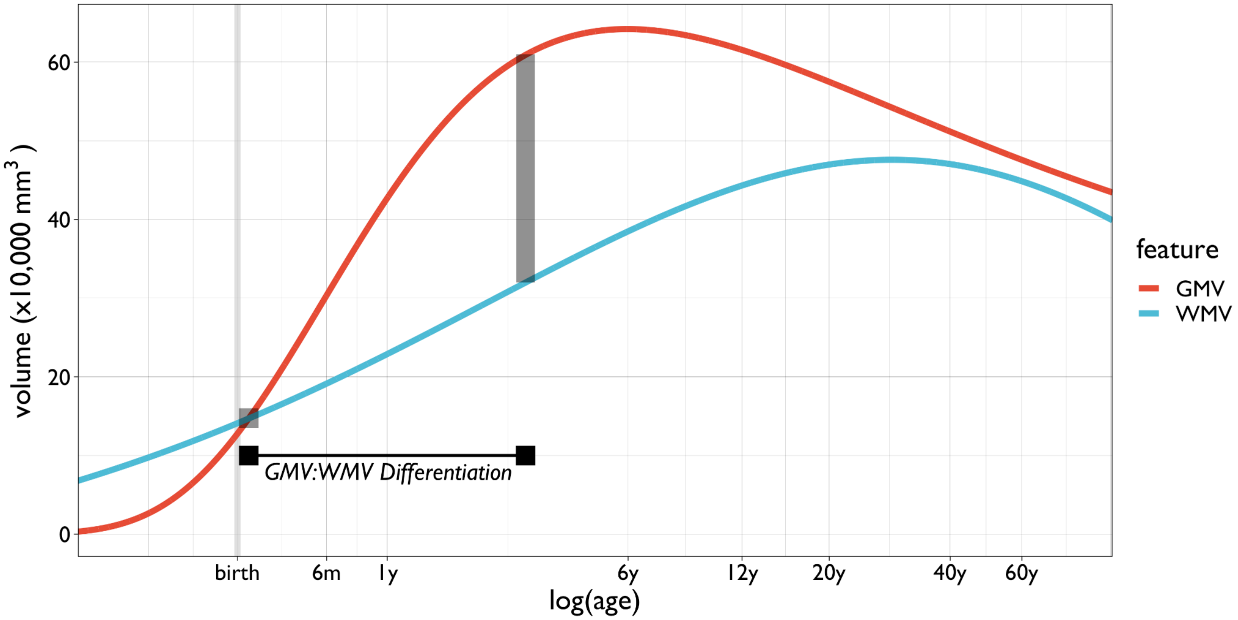


***Fig. S9.2. Definition of grey matter volume (GMV) and white matter volume (WMV) differentiation.*** *This early developmental period (filled black segment) demarcates the point of intersection between the trajectories of GMV and WMV (shaded square; 298 post-conception days) until the point of maximum absolute difference between GMV and WMV (shaded rectangle; 1395 post-conception days). X-axis denotes age, calculated as log-scaled post-conception days.*

### 10. Clinical applications of centile scores

In addition to the relevance of healthy brain growth to the many facets of behaviour and cognition, reference models of typical development are important for understanding disorders that may result from or be characterised by atypical maturational trajectories[60]. For example, neuropsychiatric conditions generally have behavioural and cognitive antecedents in earlier developmental periods prior to when they are diagnosed[61]. Thus, the ability to chart human brain morphology from the earliest developmental stages through to old age would be a major advance towards the identification of imaging markers across a multitude of neurological and psychiatric diseases [62,63]. Centiles provide a standardised measure that allows such clinical comparisons across the lifespan and across disorders.

#### 10.1 Case-control and between-disorder comparisons of centile scores on cerebrum tissue volumes and extended global MRI phenotypes

For brevity our main paper only shows pairwise comparisons of centile differences for the healthy control group compared to groups of at least N=500 diagnosed casest (**Fig. 4A**). Tables **ST3.1-3.7** list the full results for each pairwise comparison conducted with the same Monte Carlo permutation tests (10,000 permutations). The tables also include Cohen's *d* effect size estimates (including Hedges confidence intervals for these estimates[64]). Below we show all significant pairwise comparisons surviving FDR correction (corrected *P*<0.001). Clinical groups were aggregated by categorical diagnosis, per each individual study design, or for endorsement of symptomatology as in the case of population cohorts (e.g., ABCD and UKB).

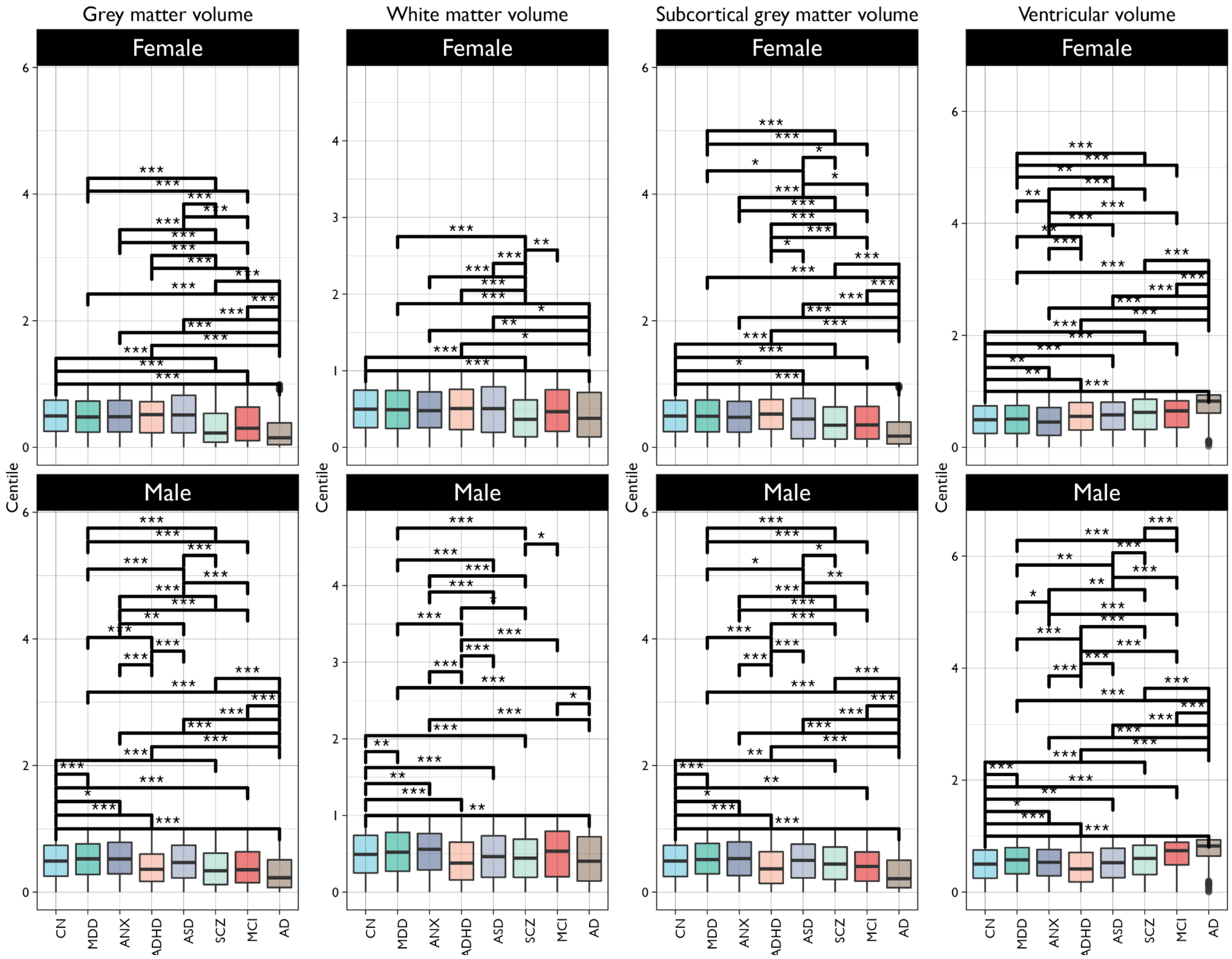


***Fig. S10.1.1. Case-control and between-disorder comparisons of centile scores on cerebrum volumes.*** *The same as shown in main* ***Fig. 4A*** *but not limited to comparison with the CN group only. Asterisks indicate significance after FDR correction (q<0.001) as computed using Monte Carlo permutation*

*tests and the Benjamini-Hochberg[65] procedure to correct for multiple comparisons entailed by all possible pairwise tests. Abbreviations; Control (CN), Alzheimer's Disease (AD), Attention Deficit Hyperactivity Disorder (ADHD), Anxiety or Phobia (ANX), Autism Spectrum Disorder (ASD), Anxiety/Phobia (ANX), Mild Cognitive Impairment (MCI), Major Depressive Disorder (MDD), Schizophrenia (SCZ); Grey Matter Volume (GMV), Subcortical Grey Matter Volume (sGMV), White Matter Volume (WMV), Ventricular Cerebrospinal Fluid (CSF).*

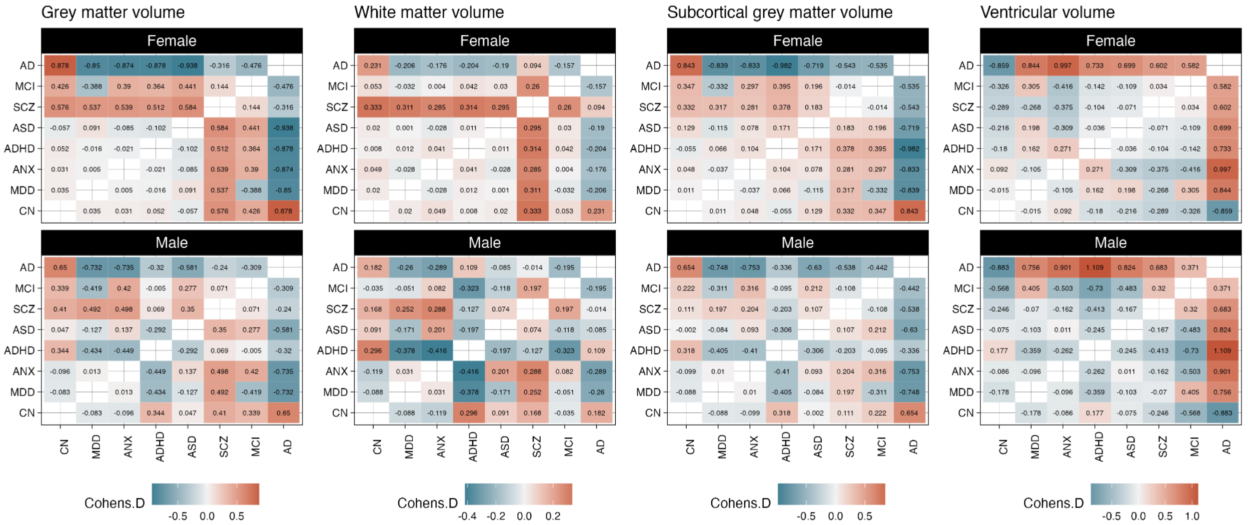


***Fig. S10.1.2. Case-control and between-disorder comparisons of centile scores on cerebrum volumes.*** *Matrix plots show the pairwise Cohen's d values for every combination. More positive d indicates that the centile score on the x-axis was higher relative to the corresponding label on the y-axis, more negative d indicates the opposite effects, i.e., CN > AD in both males and females. Abbreviations: control, CN; Alzheimer's disease, AD; attention deficit hyperactivity disorder, ADHD; autism spectrum disorder, ASD; anxiety/phobia (ANX), mild cognitive impairment, MCI; major depressive disorder, MDD; schizophrenia, SCZ; grey matter volume, GMV; subcortical grey matter volume, sGMV; white matter volume, WMV; ventricular cerebrospinal fluid volume, CSF.*

For example, we observed that men with a diagnosis of Alzheimer's Disease (AD) and Schizophrenia (SCZ) showed a median centile difference significantly below the CN group for GMV *(CN-AD=0.263, $P_{FDR}$=0.0003, Cohen's d=0.65; CN-SCZ=0.153, $P_{FDR}$=0.0003, Cohen's d=0.41)*, WMV *(CN-AD=0.089, $P_{FDR}$=0.0016, Cohen's d=0.18; CN-SCZ=0.049, $P_{FDR}$=0.0005, Cohen's d=0.17)* and sGMV *(CN-AD=0.279, $P_{FDR}$=0.0003, Cohen's d=0.65; CN-SCZ=0.048, $P_{FDR}$=0.0052, Cohen's d=0.11)*, while Ventricular CSF volume in AD was significantly larger (*CN-AD=-0.322, $P_{FDR}$=0.0003, Cohen's d=-0.88*). Interestingly, men with mild cognitive impairment (MCI), while overall showing a pattern similar to the AD group, did not differ significantly on WMV. For women this pattern was even stronger: GMV *(CN-AD=0.345, $P_{FDR}$=0.0003, Cohen's d=0.88; CN-SCZ=0.269, $P_{FDR}$=0.0003, Cohen's d=0.58)*, WMV *(CN-AD=0.12, $P_{FDR}$=0.001, Cohen's d=0.23; CN-SCZ=0.133, $P_{FDR}$=0.001, Cohen's d=0.33)*, sGMV *(CN-AD=0.317, $P_{FDR}$=0.0004, Cohen's d=0.84; CN-SCZ=0.145, $P_{FDR}$=0.0004, Cohen's d=0.33)* and Ventricular CSF volume (*CN-AD=-0.332, $P_{FDR}$=0.0004, Cohen's d=-0.86, CN-SCZ=-0.132, $P_{FDR}$=0.0004, Cohen's d=-0.29*). Although different mechanisms underlie the neuroanatomical abnormalities observed in AD and schizophrenia[66], and in the case of schizophrenia the cellular basis remains to be fully elucidated, cortical grey matter loss has been associated with cognitive impairment and psychiatric symptomatology in both disorders[67]. It would be premature to speculate on the clinical interpretation of these consistencies between effect direction and size in SCZ and AD, and it should be noted in particular that the AD cohort includes some younger individuals where clinical status was confirmed with post-mortem pathology.

Males with Attention Deficit Hyperactivity Disorder (ADHD) also showed evidence of overall lower GMV and WMV centiles (though note the discussion on centile distributions in **S10.3**): GMV *(CN-ADHD=0.131,* $P_{FDR}$*=0.0003, Cohen's d=0.34)*; WMV *(CN-ADHD=0.114,* $P_{FDR}$*=0.0005, Cohen's d=0.3)*. Ventricular volumes were slightly decreased in ADHD (*CN-ADHD= 0.087,* $P_{FDR}$*=0.0003, Cohen's d=0.18*). In addition, males with ADHD showed a reduced sGMV pattern (*CN-ADHD= 0.124,* $P_{FDR}$*=0.0003, Cohen's d=0.32*). These significant differences were less apparent in females and even showed a globally increased ventricular volume (*CN-ADHD= -0.06,* $P_{FDR}$*=0.0026, Cohen's d=-0.18*). Finally, it is interesting to note that the lower tissue volumes in ADHD were not accompanied by a commensurate increase in ventricular CSF as was observed in for example AD and SCZ.

Males with Autism Spectrum Disorder (ASD) also showed marginal evidence of overall lower WMV centiles (though again note the discussion on centile distributions in **S10.3**): WMV *(CN-ASD=0.03,* $P_{FDR}$*=0.0008, Cohen's d=0.1)*. This was not the case for females with ASD. Ventricular volumes were slightly decreased in females with ASD (*CN-ASD= 0.025,* $P_{FDR}$*=0.0056, Cohen's d=-0.08*). In addition, females with ASD showed a reduced sGMV pattern (*CN-ASD= 0.048,* $P_{FDR}$*=0.0246, Cohen's d =0.13*). These significant differences in sGMV were not observed in males.

The same case-control analysis was performed for the three extended global MRI phenotypes. **Fig. S10.1.3** shows significant differences relative to the CN group in a similar presentation as represented in **Fig. 4A**. All significant pairwise combinations are visualised in **Fig. S10.1.4** and all statistical pairwise effect-sizes and *P*-values are provided in **ST3.5-3.7**.

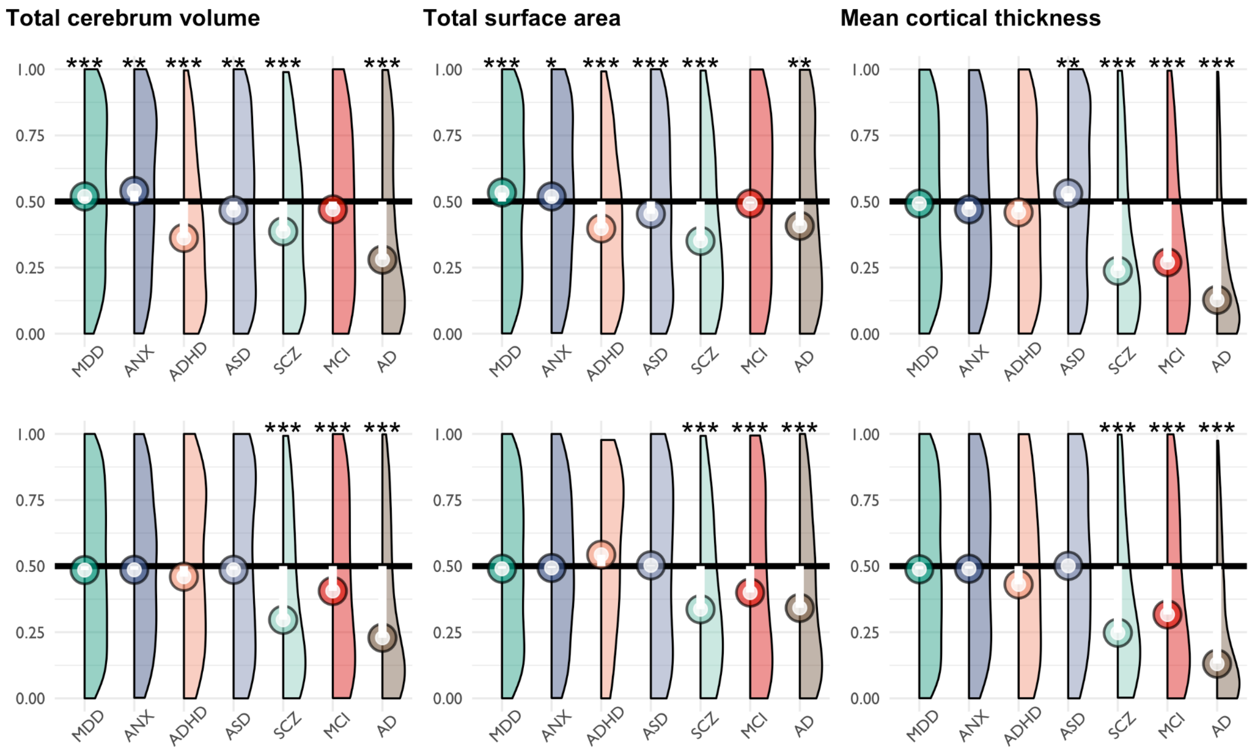


***Fig. S10.1.3. Case-control differences of centile scores on extended global MRI phenotypes.*** *Centile distributions for each of the clinical disorders with N>500 cases relative to the CN group median (depicted as a horizontal black line). The top row depicts the male only subset, the bottom the female only subset. The deviation in each clinical group is overlaid as a lollipop plot (white line with circle corresponding to the clinical group median). Pairwise tests for significance were done using Monte Carlo permutation (10,000 permutations) and P-values adjusted using the Benjamini-Hochberg FDR procedure for the multiple comparisons entailed by testing all possible pairs. Only significant differences to CN (corrected P<0.001) are depicted here and highlighted with an asterisk. For a complete overview of all pairwise comparisons, see supplementary tables* ***ST3.5-3.7****. Abbreviations; control, CN; Alzheimer's disease, AD; attention deficit hyperactivity disorder, ADHD; autism spectrum disorder, ASD; anxiety/phobia (ANX), mild cognitive impairment, MCI; major depressive disorder, MDD; schizophrenia, SCZ; grey matter volume, GMV; subcortical grey matter volume, sGMV; white matter volume, WMV; ventricular cerebrospinal fluid volume, CSF.*

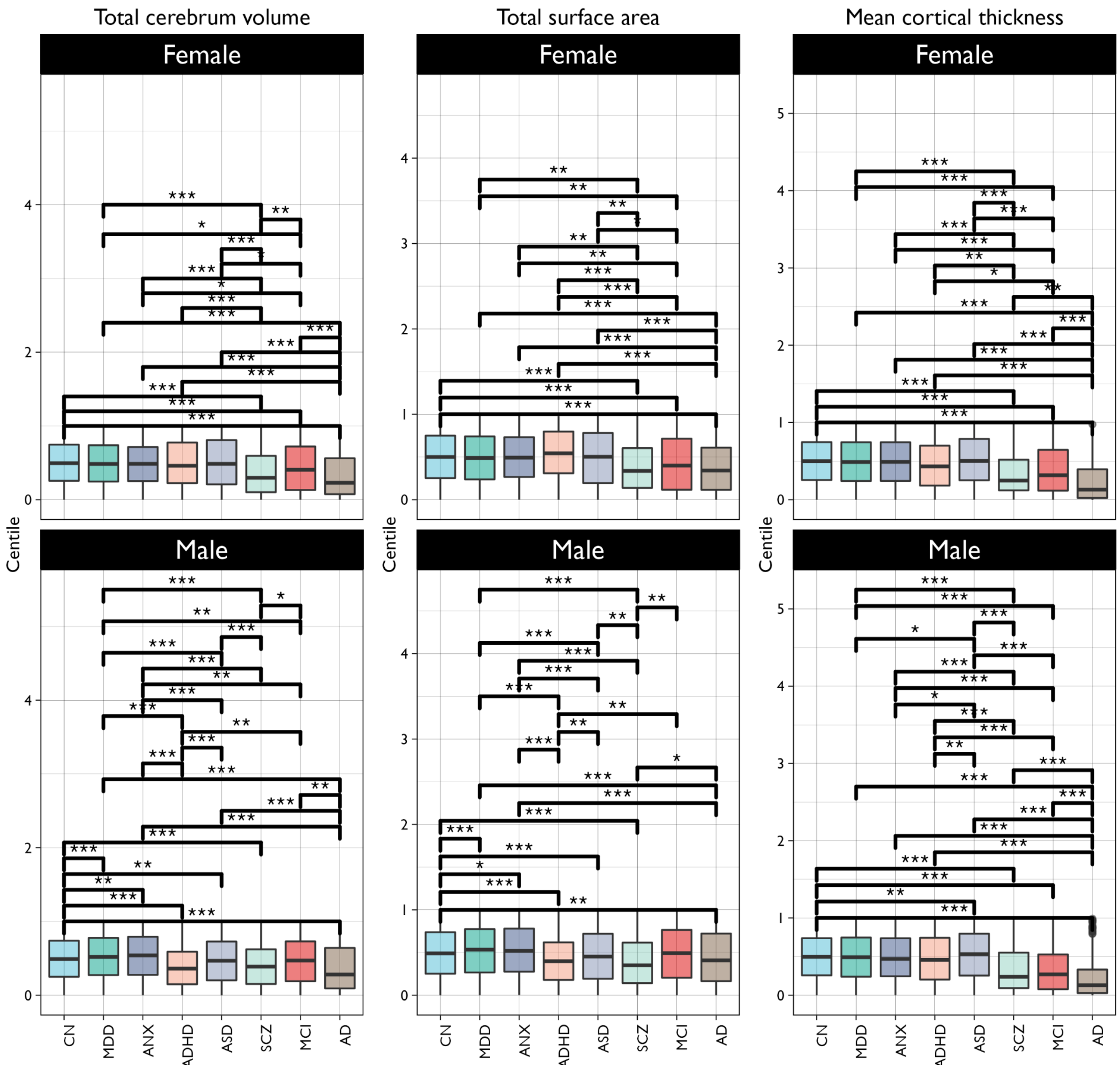


***Fig. S10.1.4. Case-control and between-disorder comparisons of centile scores on extended global MRI phenotypes.*** *The same as shown in main* ***Fig. 4A*** *and S10.1.1 but not limited to comparison with the CN group only. Asterisks indicate significance after FDR correction (q<0.001) as computed using Monte Carlo permutation tests and using a Benjamini-Hochberg[65] correction to correct for multiple comparisons accounting for all possible pairwise combinations. Abbreviations; control, CN; Alzheimer's disease, AD; attention deficit hyperactivity disorder, ADHD; autism spectrum disorder, ASD; anxiety/phobia (ANX), mild cognitive impairment, MCI; major depressive disorder, MDD; schizophrenia, SCZ.*

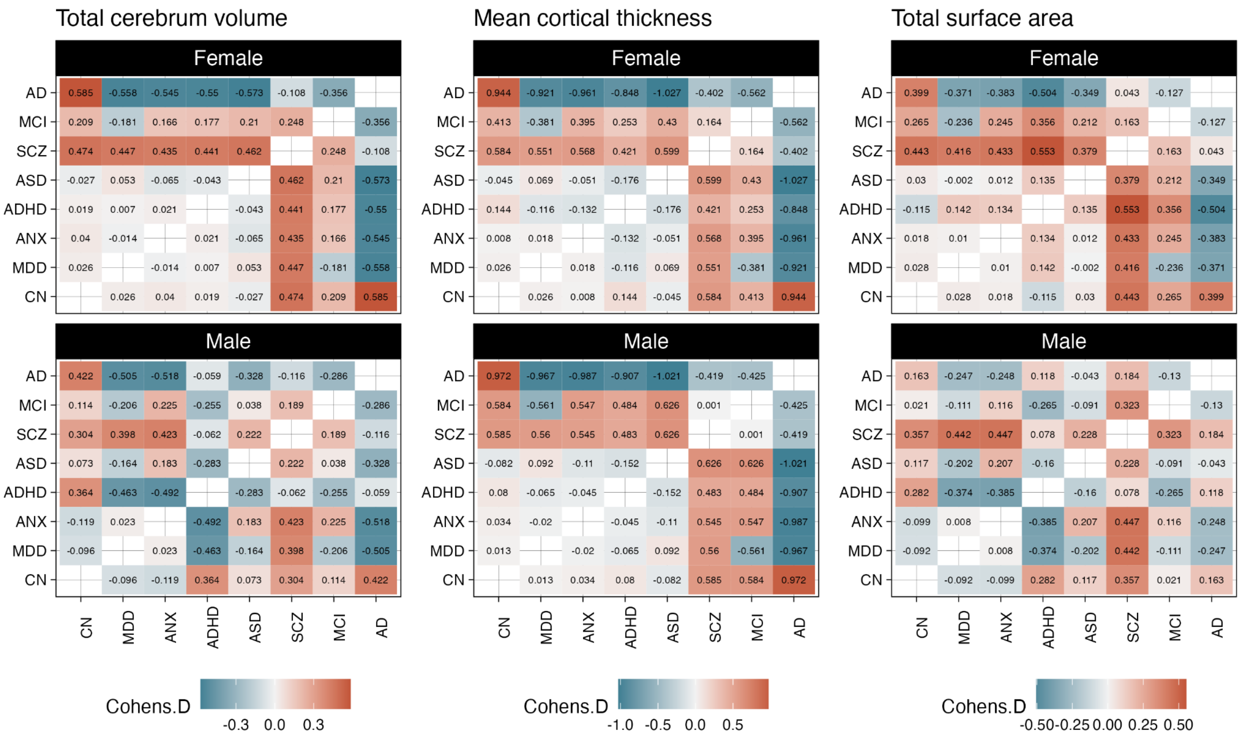


***Fig. S10.1.5. Case-control and between-disorder comparisons of centile scores on extended global MRI phenotypes.*** *Matrix plots show the pairwise Cohen's d values for every combination. More positive d indicates that the centile score on the x-axis was higher relative to the corresponding label on the y-axis, more negative d indicates the opposite effects, i.e., CN > AD in both males and females. Abbreviations; control, CN; Alzheimer's disease, AD; attention deficit hyperactivity disorder, ADHD; autism spectrum disorder, ASD; anxiety/phobia (ANX), mild cognitive impairment, MCI; major depressive disorder, MDD; schizophrenia, SCZ.*

The threshold of a minimal sample size of N=500 restricted our analyses to the reported 7 clinical conditions. To provide a comprehensive assessment across multiple minimum sample size thresholds for clinical groups, **ST3.8-3.28** lists the statistical values for these comparisons at thresholds of N=250, 100 and 50 (using non-parametric Monte Carlo permutation tests).

#### 10.2 Multimodality of centile distributions in clinical disorders

It should be noted that while centile scores provide an age-normalised and sex specific assessment, these broad case-control comparisons do not explicitly account for the possibility of developmentally specific subgroups and or trajectories, especially in cross-sectional data. For example, it is possible the early developmental differences in cerebrum tissue volumes normalise or become less apparent later in life. Indeed recent studies would suggest such age-related patterns[68,69]. To explore the possible or even likely existence of subgroups within the space of centile scores, we assessed the number of peaks in the probability density function. Density plots were generated with the 'geom_flat_violin' option from the Raincloud package[70]. Estimation of densities and the resulting number of peaks were done using the default settings of the '*density()*' function in the R stats package[71] using a Gaussian smoothing kernel[72,73] which defaults to 0.9 times the minimum of the standard deviation and the interquartile range divided by 1.34 times the sample size to the negative one-fifth power (Silverman's 'rule of thumb'[74]); unless the quartiles coincide, when a positive result will be guaranteed. The number of peaks was defined as the inflection point on these Gaussian smoothed density curves. Unimodality of smoothed density curves was tested using Hartigan's dip-test[75] which indicated that none of the distributions were perfectly unimodal (see **ST4.1-4.7**).

Most distributions revealed at least 2 peaks and varying levels of skewness in the direction of global case control differences highlighted in **Fig. 4** and below. For example, the AD cohort shows a clear peak and strong skewness toward the lower centile range for GMV and sGMV (and to a lesser extent for WMV), combined with a skew to the higher centile range for ventricular volume commensurate with the direction of main effects. The multiple peaks of these distributions are particularly salient in the context of ASD where there have been differing accounts of both micro- and macrocephaly[76]. Head circumference-based findings of macrocephaly in ASD were first reported in 2001 by Courchesne and colleagues[77] and subsequently confirmed in a meta-analysis[78]. This early overgrowth has also been thought to be accompanied by a later arrest in growth[69,79]. While our reference-based approach to derive centiles is age-agnostic to the extent that static time-points are normalised for age, the existence of peaks at both extreme ends and in the middle of the GMV and sGMV centile distribution would suggest that there may be both micro- and macro-cephalic subgroups of ASD. How these are clustered or aligned across the lifespan would be an interesting topic for follow-up research. One point worth highlighting is that the distributions in ASD do not show the level of extreme skewness observed in AD and so it is possible that canonical or mean-based case-control analyses may erroneously indicate little difference with typical development by the mere fact that the extremes balance each other out when calculating a mean difference. Furthermore, by aggregating subgroups all together, it is possible that case-control comparisons could show biased effects either driven by the predominance of a specific subgroup[76], the effect size of a specific subgroup[80] or the relative representation of specific age-ranges for specific clinical cohorts in the present dataset. Leveraging centiles to parse such within-disorder heterogeneity is an important area for follow-up research.

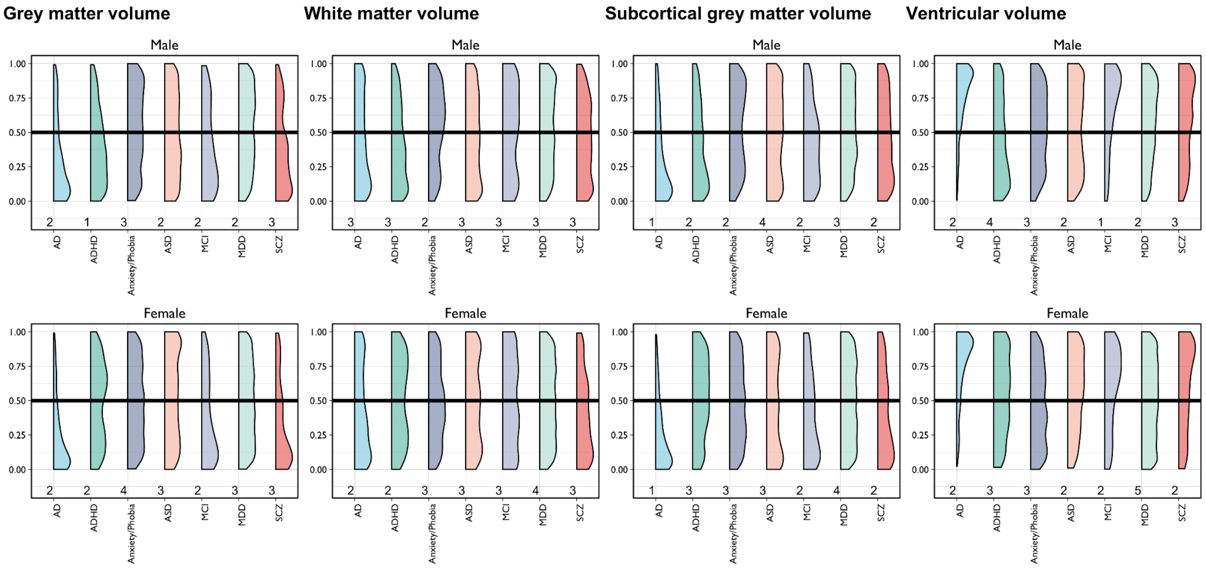


***Fig. S10.2.1. Probability density plots of centile scores on cerebrum tissue volumes for clinical cohorts with at least N=500 diagnosed cases****. Labels underneath each density plot show the estimated number of peaks or modes in the smoothed distribution. Abbreviations; control, CN; Alzheimer's disease, AD; attention deficit hyperactivity disorder, ADHD; autism spectrum disorder, ASD; anxiety/phobia (ANX), mild cognitive impairment, MCI; major depressive disorder, MDD; schizophrenia, SCZ; grey matter volume, GMV; subcortical grey matter volume, sGMV; white matter volume, WMV; ventricular cerebrospinal fluid volume, CSF.*

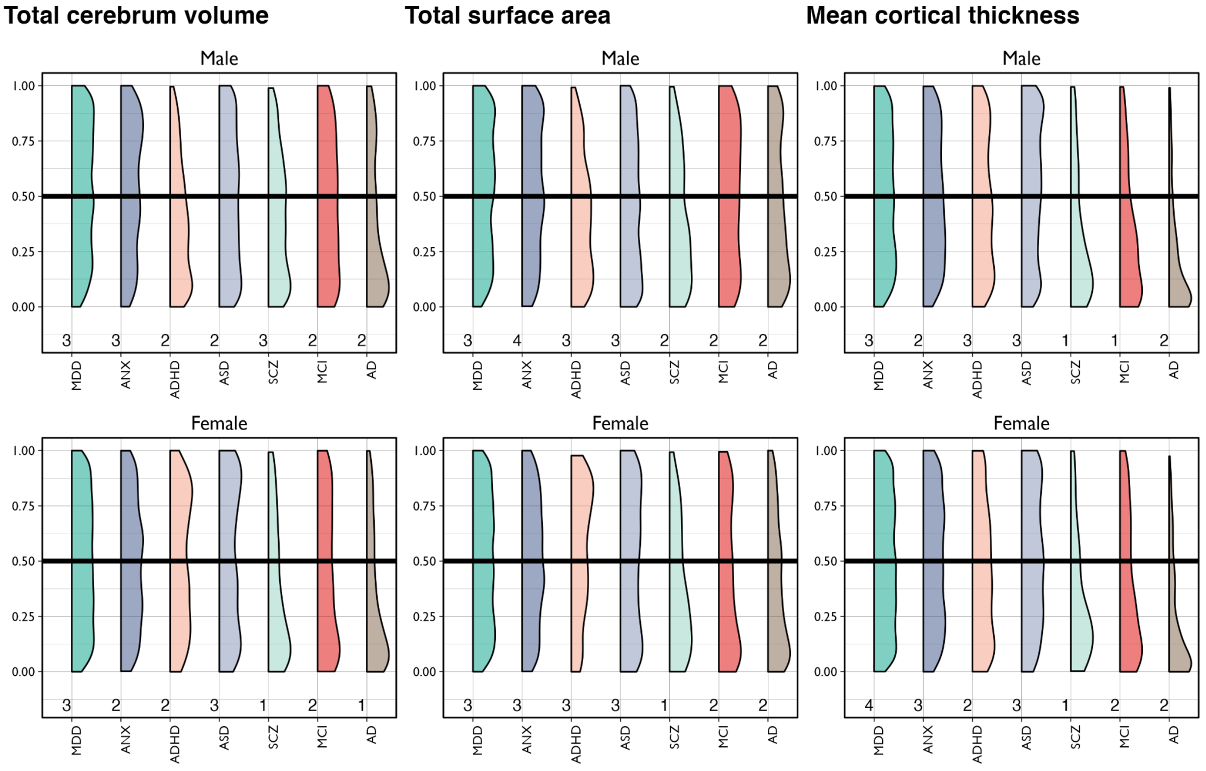


***Fig. S10.2.2. Probability density plots of centile scores on extended global MRI phenotypes for clinical cohorts with at least N=500 diagnosed cases****. Labels underneath each density plot show the*

*estimated number of peaks in the smoothed distribution. Abbreviations; control, CN; Alzheimer's disease, AD; attention deficit hyperactivity disorder, ADHD; autism spectrum disorder, ASD; anxiety/phobia (ANX), mild cognitive impairment, MCI; major depressive disorder, MDD; schizophrenia, SCZ; grey matter volume, GMV; subcortical grey matter volume, sGMV; white matter volume, WMV; ventricular cerebrospinal fluid volume, CSF.*

#### 10.3 Case-control differences on CMD

In order to determine whether centiles provided sensitivity to detect case-control differences over all clinical groups at specific developmental epochs, we conducted an exploratory analysis using developmental windows as defined by Kang et al.[59] Specifically, we re-coded all diagnostic labels to either healthy controls (CN) or diagnosed cases of any disorder (DX), then estimated the centile Mahalanobis distance (CMD; analogous to **Fig. 4**) across the four cerebrum tissue volumes relative to the CN group mean (0.5). Then we ran two-sided Monte Carlo permutation tests (10,000 permutations) on CMD within each developmental window. We found overall case-control differences in CMD across the lifespan (**Fig. S10.3**), indicating that relatively increased CMD - a multivariate marker of atypicality - was associated with DX status. These differences were most strongly pronounced in late adulthood (mean difference, 0.655, $P<0.001$; Cohen's $d=0.25$), middle/late childhood (mean difference=0.493, $P<0.001$; Cohen's $d=0.24$), adolescence (mean difference=0.512, $P<0.001$; Cohen's $d=0.24$), young adulthood (mean difference, 0.363, $P<0.001$; Cohen's $d=0.17$) and middle adulthood (mean difference, 0.133, $P<0.001$; Cohen's $d=0.06$). In foetal, neonatal, and very early childhood, the current dataset was insufficiently powered to determine gross differences on disease status (**Fig. S10.3**, panel B label provides the number of individuals with any kind of diagnostic label).

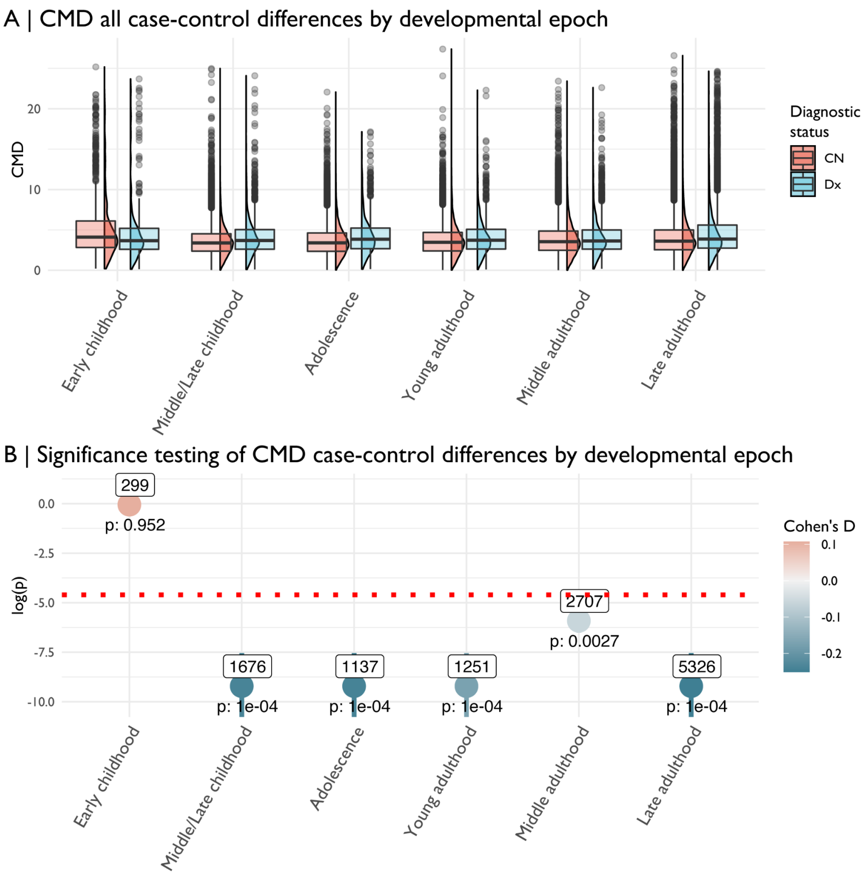


***Fig. S10.3. Case-control differences, between healthy controls (CN) and all diagnosed cases (DX), for centile Mahalanobis distance (CMD) over all four cerebrum tissue volumes at each developmental window over the lifespan.*** *A | The relative distributions of CMD for CN and DX groups in each developmental window. B | The point-range plot of the P-values and their confidence intervals as computed using a Monte Carlo permutation test (10,000 permutations). Labels above each point indicate the number of individuals in the DX group in each developmental window. The red-dotted line shows P=0.01*

#### 10.4 Summary centile comparison

Here we highlight the difference in two summary centile metrics that could be used to characterise (a)typicality across all neuroimaging phenotypes: the mean centile and the centile Mahalanobis distance. The mean centile is simply the average of the centile scores for all 4 cerebrum tissue volumes for a given subject. The centile Mahalanobis distance (CMD) is a summary dispersion metric, which is statistically distinct from the mean (see **Fig. S10.3, Fig. S10.4.1** and **SI1.6**). Whereas the mean centile score is normally distributed across subjects, CMD is skewed—biased towards lower estimates. Thus, while the mean centile can obscure correlated changes in phenotypes—such as increased CSF with decreased GMV in AD patients—CMD can directly capture this covariation. Overall, both metrics showed relatively similar distributions in diagnosed cases and healthy controls (**Fig. S10.4.1**), with highly varying estimates across diagnostic groups (medians for each category plotted in **Fig. S10.4.2** and **Fig. S10.4.3**).

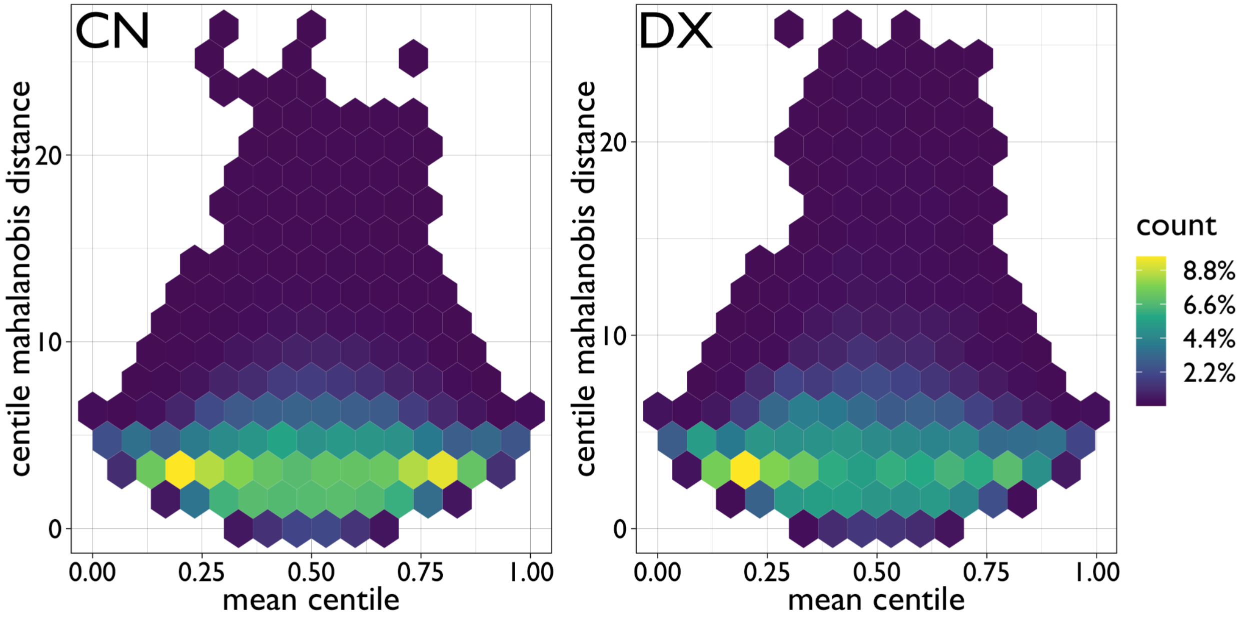


***Fig. S10.4.1. Hex plot showing all case-control differences (between healthy controls (CN) and all cases regardless of diagnostic category (DX) of centile scores averaged across phenotypes (mean centile) versus the preferred multivariate metric of centile dispersion (CMD: centile Mahalanobis distance).*** *Count refers to the hex-binning percentage of the total dataset within the CN and DX groups. Thus, as each coloured hexagon represents multiple data points (subjects), it is clear that both groups show a skewed distribution for CMD despite a relatively normal distribution for the mean centile (with DX having a preponderance of subjects with low mean centiles, see **Fig. S11.1**).*

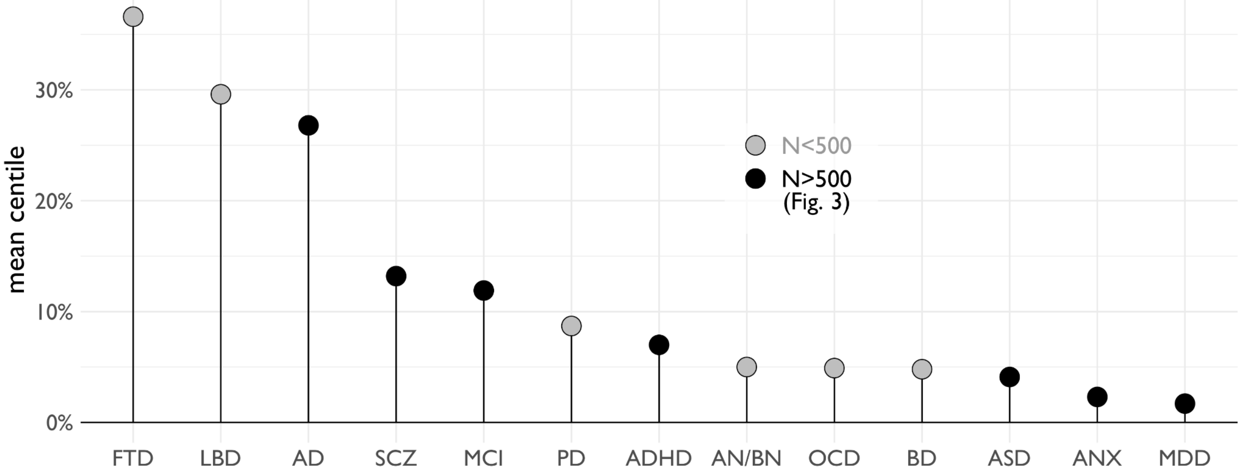


***Fig. S10.4.2. Case-control differences in mean centile scores averaged across all 4 cerebrum tissue volumes (GMV, WMV, sGMV, CSF).*** *Black circles represent clinical conditions included in the main analyses in **Fig. 4A** (N>500 per condition); grey circles represent clinical conditions represented in multiple datasets with a total N<500 subjects. Abbreviations: Abbreviations: Alzheimer's disease, AD; attention deficit hyperactivity disorder, ADHD; anorexia nervosa or bulimia nervosa, AN/BN; anxiety or phobia, ANX; autism spectrum disorder, ASD; bipolar disorder, BD; fronto-temporal dementia, FTD; Lewy body dementia,*

*LBD; mild cognitive impairment, MCI; major depressive disorder, MDD; obsessive-compulsive disorder, OCD; Parkinson's disease, PD; schizophrenia, SCZ.*

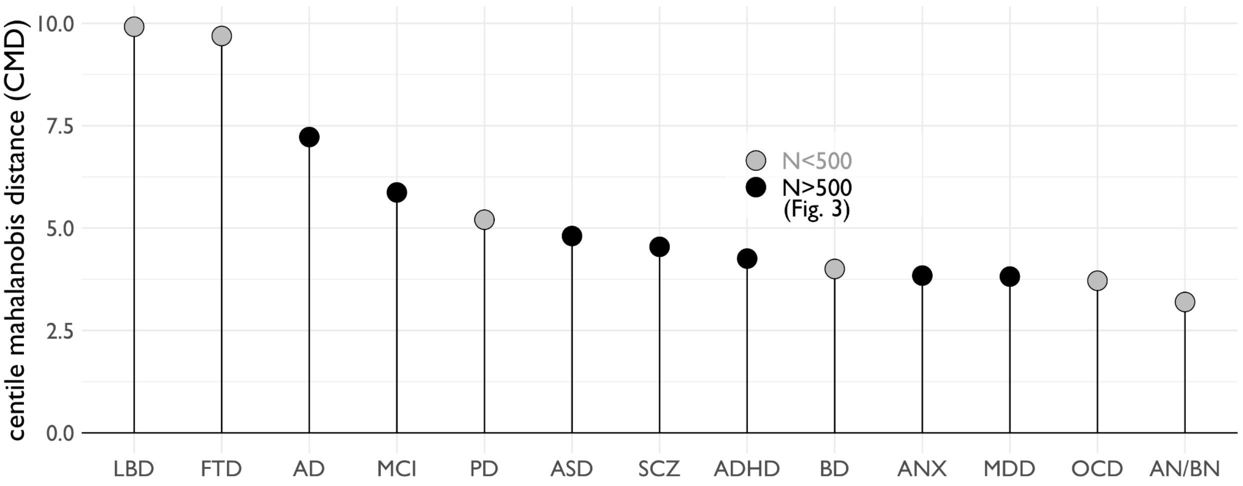


***Fig. S10.4.3. Case-control differences of median centile Mahalanobis distance (CMD) across tissue types (GMV, WMV, sGMV, CSF) for all clinical disorders.*** *Black circles represent clinical conditions included in the main analyses in* ***Fig. 4A*** *(N>500 per condition); grey circles represent clinical conditions represented in multiple datasets with a total N<500 subjects. Abbreviations: Alzheimer's disease, AD; attention deficit hyperactivity disorder, ADHD; anorexia nervosa or bulimia nervosa, AN/BN; anxiety or phobia, ANX; autism spectrum disorder, ASD; bipolar disorder, BD; fronto-temporal dementia, FTD; Lewy body dementia, LBD; mild cognitive impairment, MCI; major depressive disorder, MDD; obsessive-compulsive disorder, OCD; Parkinson's disease, PD; schizophrenia, SCZ.*

### 11. Cross diagnostic analyses

The functional interpretations of centile scores of brain volume will ultimately be dependent on, and informed by, the particular context in which brain charts are implemented. But the brain lifespan models we have presented here provide a standardised reference point, normalised for age, that could be relevant to many investigations of brain-behaviour relationships. Further work will be needed to fully explore the applicability of GAMLSS centile scoring for individual assessments in clinical contexts. Like traditional growth charts, an individual's centile score may not be clinically decisive on its own in many cases, i.e., someone could have a bigger or smaller brain, or be short or tall, without diagnostic implications. To explore the ability of GAMLSS derived centiles scores for case-control discriminability we conducted two cross-disorder analyses.

#### 11.1 Sliding window analyses of cross-disorder discriminability

We computed odds ratios for clinical disorders using a sliding window across the full range of centile scores for cerebrum tissue volumes (window size=0.1, increment size=0.05). Major diagnostic categories (as in **Fig. 4)** were combined to form one group of all diagnosed cases (DX or non-CN) and compared to healthy controls (CN) to estimate the odds ratio of being diagnosed with any clinical disorder. These analyses indicated that lower centile scores, especially <5%, on cerebrum tissue volumes, cortical surface area and cortical thickness were all significantly over-represented in individuals with neuropsychiatric disorders (**Fig. S11.1**). This means that a lower centile score on any or all of these brain MRI metrics was associated with a higher probability of any clinical disorder. It will be important to discover if low centile scores on brain MRI metrics are predictive of later clinical outcomes, meaning that brain charts could be used in future as paediatric growth charts are used now, to raise levels of clinical concern proportionately, rather than to make a specific diagnosis.

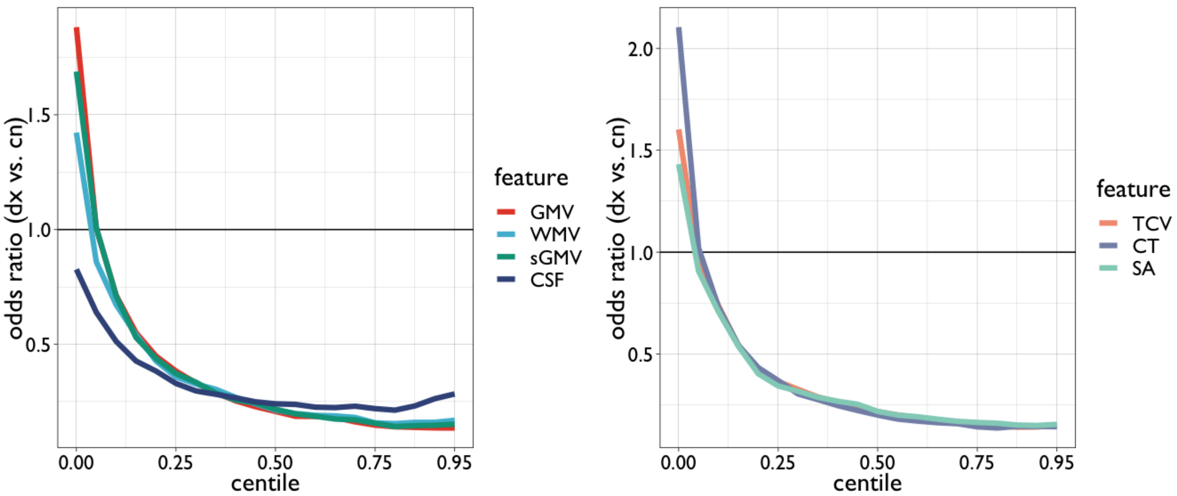


***Fig. S11.1. Brain MRI centile scores are related to the probability of any clinical disorder.*** *The odds ratio for clinical disorder (versus healthy control) is plotted on the y-axis of both panels; positive OR indicates greater risk of disorder. Centile scores by GAMLSS modeling are plotted (on the x-axis) for global brain MRI phenotypes: left panel, 4 cerebrum tissue volumes; right panel, total cerebrum volume, cortical surface area, and mean cortical thickness. Odds ratios were computed using a sliding window across centiles (window size=0.1, increment size=0.05). Diagnostic categories in* ***Fig. 4*** *were combined (i.e., binarised to make any diagnosis, or 'dx' vs. 'cn') to estimate the odds ratio of being in any clinical cohort. Scans with lower centile scores on all phenotypes, especially centiles <5%, have increased odds ratio for all clinical disorders.*

#### 11.2 Cross-diagnostic clustering

Multivariate approaches can also be used to delineate neuroanatomical signatures across metrics and to compare neuroanatomical signatures across clinical conditions, while maintaining diagnostic labels. As such, we performed data-driven hierarchical clustering based on the group average and standard deviation of centiles, combined with the overall deviation metric CMD. Centile scores were derived for each subject across the four primary lifespan phenotypes (GMV, sGMV, WMV, Ventricular CSF). We calculated the median and the variance (as standard deviation) across patients within a cohort. An overall deviation score was also calculated for each subject as CMD across all four features. These nine scores were z-scored within clinical groups, and hierarchical (k-means) clustering was used to determine patterns of these centile profiles across conditions. The silhouette coefficient was computed (across a range of k clusters between 1 and 12) to determine the optimal number of clusters for the extended clinical condition analysis (peak value, N=3).

These analyses partially capture an age-stratification and sexual dimorphism of diagnostic conditions, despite the fact that the centiles themselves were normalized for age and sex. The across-condition clustering of neuroanatomical signatures broadly recovers plausible nosological groupings (**Fig. S11.2.1-11.2.4**). Interestingly, one exception was the high similarity observed between neurodevelopmental disorders and MCI, and the fact that Parkinson's disease diverged from other neurodegenerative disorders. While these broad distributions of gross neuroanatomical centile scores may obscure specificity and interindividual variation, they showcase the potential for future investigations of disorder-specific neuroanatomical signatures defined relative to lifespan standards.

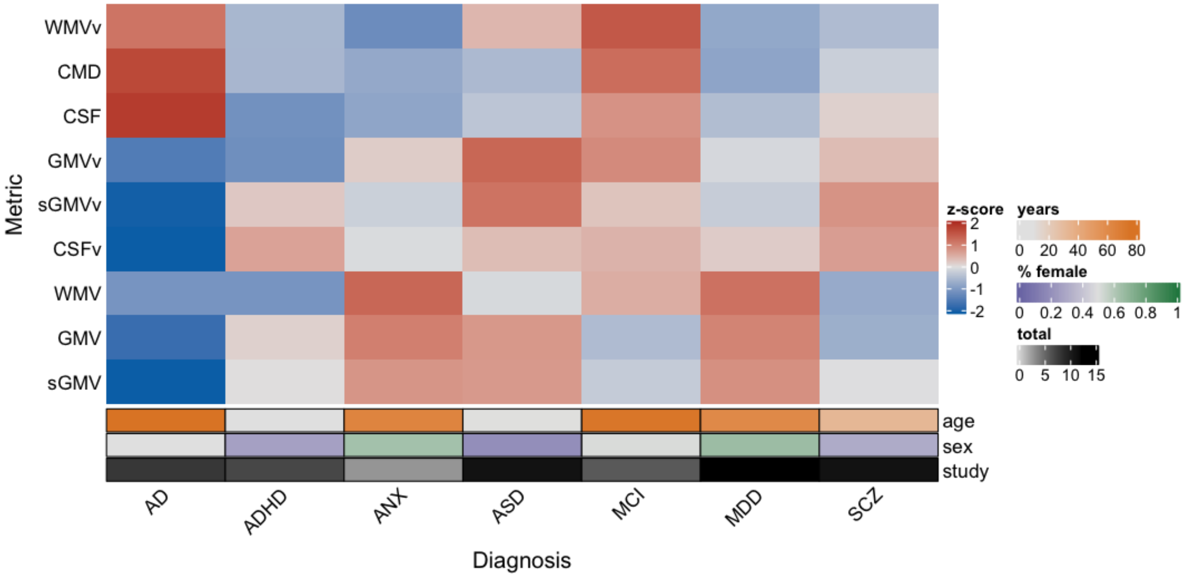


***Fig. S11.2.1. Profiles of centile scores for median cerebrum tissue volumes (GMV, WMV, sGMV, CSF), centile scores for between-subject standard deviation of cerebrum tissue volumes (GMVv, WMVs, sGMVv, CSFv), mean age, percentage female, and number of primary studies for 7 clinical disorders with N > 500 cases****, as per* ***Fig. 4****. Legend (x-axis and right annotation): 'years' corresponds to 'age' and represents median age of the diagnostic groups, '% female' corresponds to 'sex' and represents the percentage of female patients in each diagnostic group, 'total' corresponds to 'study' and represents the number of studies containing patients in the respective diagnostic group. Values of each cell represent z-scores of median centiles (row-wise across diagnostic groups) for visualisation. Clustering was determined using the gap statistic (k=1). Lowercase 'v' stands for the standard deviation and was Z-scored as per the median centiles for each phenotype across diagnostic groups. Abbreviations: Alzheimer's disease, AD; attention deficit hyperactivity disorder, ADHD; autism spectrum disorder, ASD; mild cognitive impairment, MCI; major depressive disorder, MDD; schizophrenia, SCZ; grey matter volume, GMV; subcortical grey matter volume, sGMV; white matter volume, WMV; ventricular cerebrospinal fluid volume, CSF.*

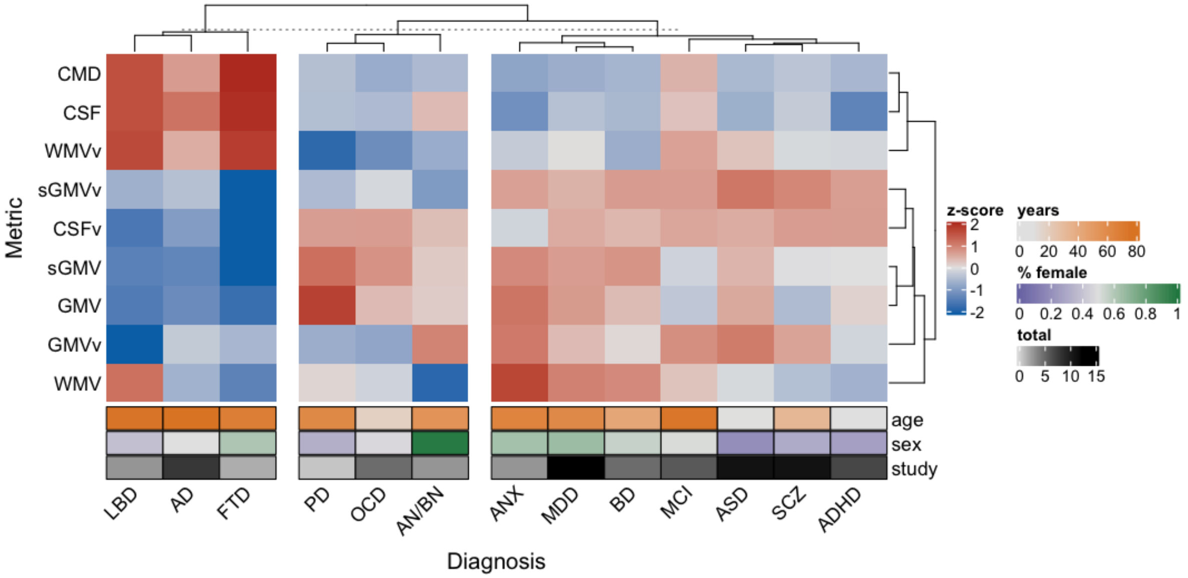


***Fig. S11.2.2. Hierarchical clustering of clinical disorder profiles of median and standard deviation of centile scores on cerebrum tissue volumes.*** *Centile profiles using a less-stringent N<500 cutoff for the number of patients with similar diagnoses, as per* ***Fig. S10.4.2*** *above. Legend (x-axis and right annotation): 'years' corresponds to 'age' and represents median age of the diagnostic groups, '% female' corresponds to 'sex' and represents the percentage of female patients in each diagnostic group, 'total' corresponds to 'study' and represents the number of studies containing patients in the respective diagnostic group. Values of each cell represent z-scores of median centiles (row-wise across diagnostic groups) for visualisation. Clustering was determined using the gap statistic (k=3). Lowercase 'v' stands for the standard deviation and was z-scored as per the median centiles for each feature across diagnostic groups. Abbreviations: Alzheimer's disease, AD; attention deficit hyperactivity disorder, ADHD; anorexia nervosa or bulimia nervosa, AN/BN; anxiety or phobia, ANX; autism spectrum disorder, ASD; bipolar disorder, BD; fronto-temporal dementia, FTD; Lewy body dementia, LBD; mild cognitive impairment, MCI; major depressive disorder, MDD; obsessive-compulsive disorder, OCD; Parkinson's disease, PD; schizophrenia, SCZ; grey matter volume, GMV; subcortical grey matter volume, sGMV; white matter volume, WMV; ventricular cerebrospinal fluid volume, CSF.*

The same analyses were repeated after the inclusion of the 3 extended brain phenotypes and broadly similar clustering patterns were preserved (**Fig. S11.2.3-11.2.4**).

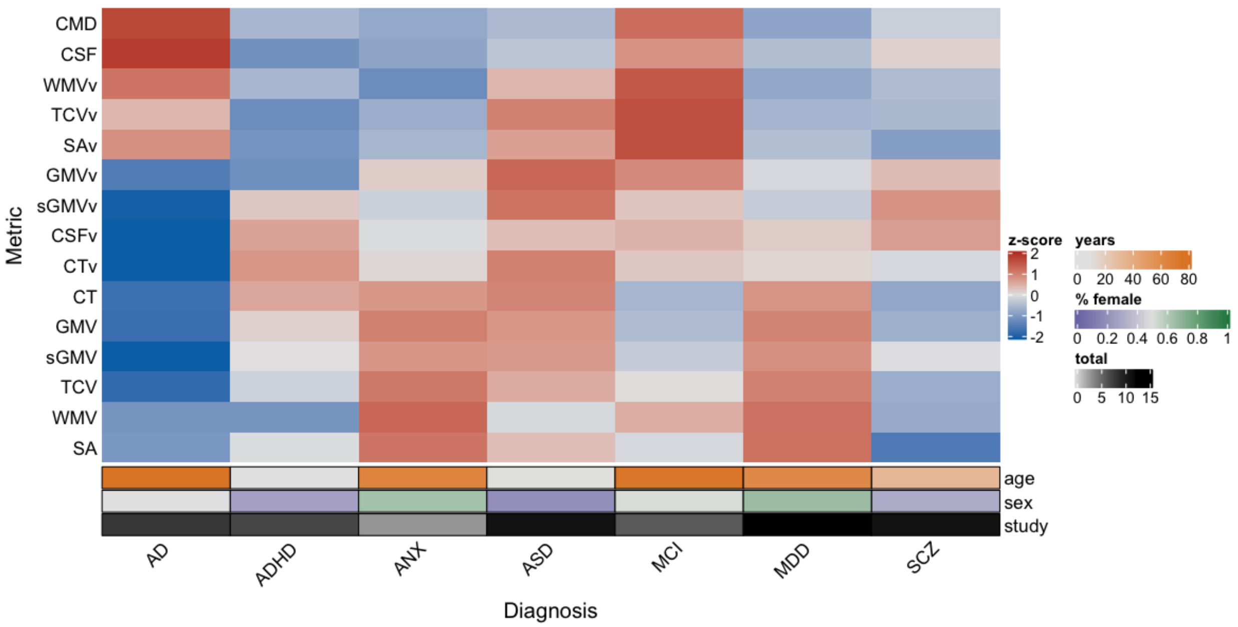


***Fig. S11.2.3. Profiles of centile scores for median cerebrum tissue volumes and extended global MRI phenotypes (GMV, WMV, sGMV, CSF, TCV, SA, CT), centile scores for between-subject standard deviation of cerebrum tissue volumes (GMVv, WMVs, sGMVv, CSFv, TCVv, SAv, CTv), age, percentage female, and number of primary studies, for 7 clinical disorders with N > 500 cases.*** *Centile profiles using a stringent N > 500 cutoff for the number of patients with similar diagnoses, as per* ***Fig. 4****. Legend (x-axis and right annotation): 'years' corresponds to 'age' and represents median age of the diagnostic groups, '% female' corresponds to 'sex' and represents the percentage of female patients in each diagnostic group, 'total' corresponds to 'study' and represents the number of studies containing patients in the respective diagnostic group. Values of each cell represent Z-scores of median centiles (row-wise across diagnostic groups) for visualisation. Clustering was determined using the gap statistic (k=1). Lowercase 'v' stands for 'variance' and was calculated as the standard deviation (rather than median), and was Z-scored as per the median centiles for each phenotype across diagnostic groups. Abbreviations: Alzheimer's disease, AD; attention deficit hyperactivity disorder, ADHD; anxiety or phobia, ANX; autism spectrum disorder, ASD; mild cognitive impairment, MCI; major depressive disorder, MDD; schizophrenia, SCZ.*

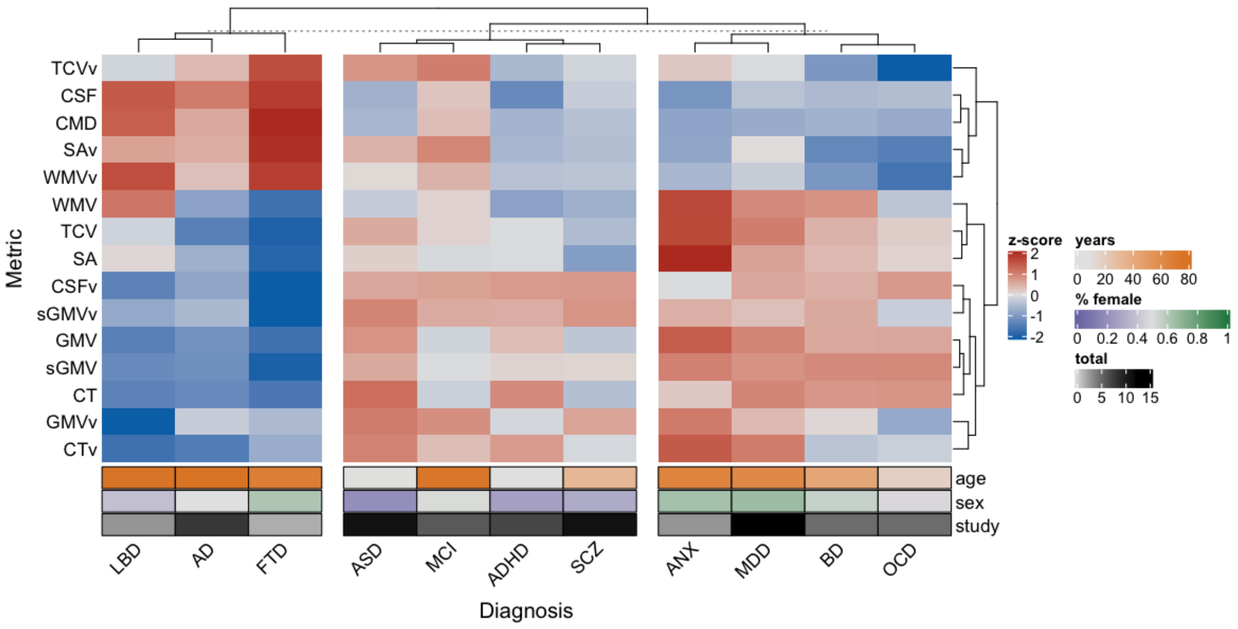


***Fig. S11.2.4. Hierarchical clustering of clinical disorder profiles of median and standard deviation of centile scores on cerebrum tissue volumes and extended global MRI phenotypes.*** *Centile profiles using a less-stringent N<500 cutoff for the number of patients with similar diagnoses, as per* ***Fig. S11.2.2*** *above. The Parkinson's and anorexia/bulimia clinical groups were removed from this analysis due to the lack of available surface-based MRI phenotypes (cortical thickness and surface area). Legend (x-axis and right annotation): 'years' corresponds to 'age' and represents median age of the diagnostic groups, '% female' corresponds to 'sex' and represents the percentage of female patients in each diagnostic group, 'total' corresponds to 'study' and represents the number of studies containing patients in the respective diagnostic group. Values of each cell represent Z-scores of median centiles (row-wise across diagnostic groups) for visualisation. Clustering was determined using the gap statistic (k=3). Lowercase 'v' stands for 'variance' and was calculated as the standard deviation (rather than median), and was Z-scored as per the median centile for each phenotype across diagnostic groups. Abbreviations: Alzheimer's disease, AD; attention deficit hyperactivity disorder, ADHD; anxiety or phobia, ANX; autism spectrum disorder, ASD; bipolar disorder, BD; fronto-temporal dementia, FTD; Lewy body dementia, LBD; mild cognitive impairment, MCI; major depressive disorder, MDD; obsessive-compulsive disorder, OCD; schizophrenia, SCZ.*

The clustering results remained relatively consistent with the inclusion of the extended global MRI phenotypes, which is likely due to the intrinsic covariance or correlation between the phenotypes, i.e., TCV is a sum of tissue class volumes, and SA and CT are geometrically related to GMV. However, despite this general consistency, some phenotypes (e.g., WMV) showed relative variability across diagnostic categories within each cluster (**Figs. S11.2.2** and **S11.2.4**) – speculatively due to the differential impact on brain tissue compartments based on variance in developmental trajectories. It is to be expected that hierarchical clustering results will evolve as more metrics, MRI modalities and diagnostic groups are included in the reference dataset in future.

#### 12. Associations of birth weight and gestational duration with centile scores on cerebrum tissue volumes

To examine the effects of early life stress on centile scores, we examined 5 independent samples across the lifespan with self-reported gestational age at birth and/or birth weight (dHCP, neonatal; UNC, neonatal and early infancy/childhood; ABCD, late childhood; NIH, childhood/adolescence/young adulthood; UKB, mid-late adulthood). Average centile scores on all four cerebrum tissue volumes were significantly related to multiple metrics of premature birth across datasets (gestational age at birth, $t$ = 13.164, $P$ < 2e-16; birth weight, $t$ = 36.395, $P$ < 2e-16). This corroborates previous work indicating the ability to capture relationships between early life factors such as birth weight and brain volumetrics measured several decades later[81].

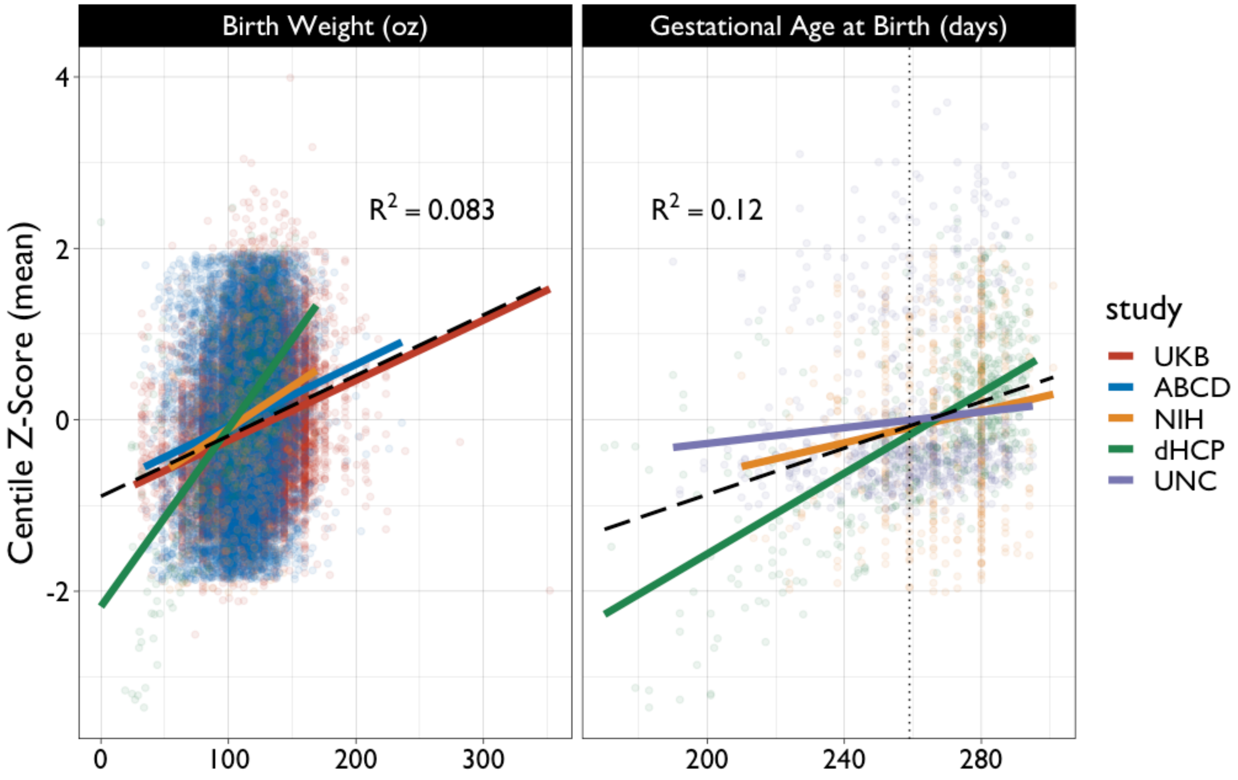


***Fig. S12. Relationships between centile scores on cerebrum tissue volumes and birth weight (left panel) and gestational age at birth (right panel) for each of 5 primary studies with relevant data available**. Centile-normalised Z-scores were computed for each global phenotype in each individual study and then averaged across phenotypes to compute a mean centile Z-score for each subject. The black dashed lines represent the relationships between mean centile scores and birth weight or gestational age at birth estimated by a linear mixed-effects model: for gestational age at birth, t = 12.624, P < 2e-16; for birth weight, t = 34.945, P < 2e-16. The black dotted line in the right panel denotes the commonly-used threshold for defining premature birth at 37 weeks post-conception. Conditional R-squared in each panel represents the variance explained by the entire model (black dashed lines).*

#### 13. Twin-based heritability of centile scores

We examined the heritability of centile scores on cerebrum tissue volumes, leveraging available data of monozygotic (MZ) and dizygotic (DZ) twins in the ABCD cohort of adolescents ( N=297 MZ, N=400 DZ pairs), and in the HCP cohort of adults (; N=138 MZ, N=78 DZ pairs). For both cohorts, zygosity was previously determined based on parental and/or self endorsement, and genetic kinship[82–84]. Heritability was estimated using Cholesky decomposition, allowing 'ACE' partitioning of the phenotypic variance into additive genetic (A), common environmental (C), and unique environmental (E) components, as implemented in the *umx* R package[85]. As shown in **Fig. 4**, we found greater heritability of centile normalised scores compared to the respective raw, non-centiled volumetric phenotypes.

#### 14. Longitudinal centiles

Due to the relative scarcity of longitudinal neuroimaging datasets (<10% of the total dataset), normative lifespan trajectories were generated with cross-sectional data. However, with the available longitudinal samples, we tested the generalisability of the models to capture within-subject variation of centile scores over time (quantified as the IQR). As described in the main text, IQR varied by only ~4% on average (see **ST5.1-5.4**) indicating highly stable centile scoring across multiple repeated scans.

##### 14.1 Longitudinal patterns in developmental epochs

Consistent with the idea of emerging variability in periods of highly dynamic change, we found that IQR was moderately related to the age distribution of longitudinal sampling (e.g., higher variability in younger samples; **Figs. S14.1** and **SI14.2**). This is in line with the level of variation expected from other anthropometrics derived from developmental longitudinal data[86] (e.g., ~30% variation on average with height during childhood). While longitudinal centile scores were generally stable, we found a number of small but significant differences in centile stability across clinical cohorts (See **Fig. S14.1** and **ST5.1-5.4**).

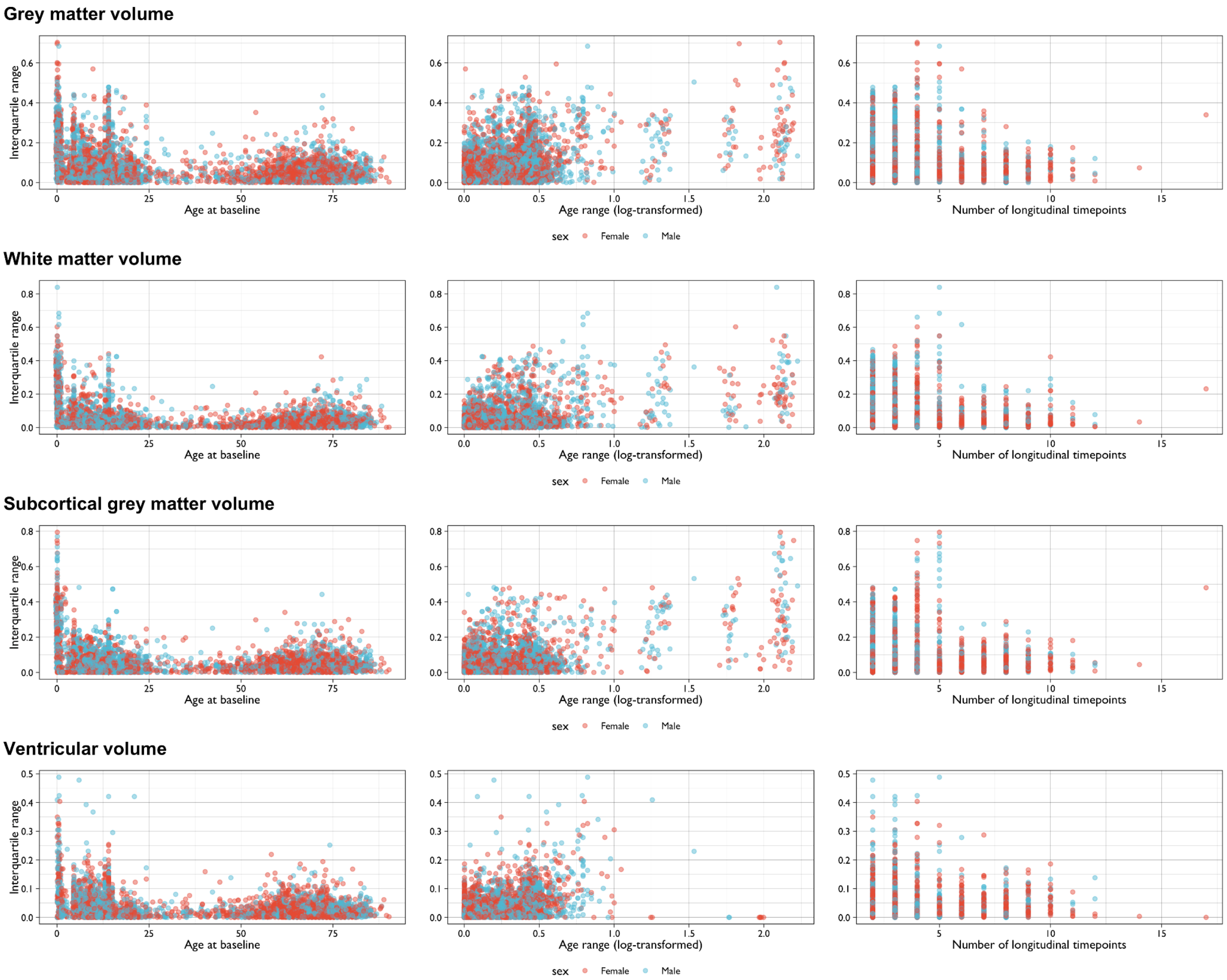


***Fig. S14.1.1. Overview of possible associations between within-subject variation (interquartile range, IQR) of longitudinal centile scores on cerebrum tissue volumes and factors that could influence longitudinal stability of centile scores.*** *First column shows the IQR in relation to the individual's age (in years) at the time of their baseline scan. Second column shows the length of follow-up (in years, log-transformed) between the baseline scan and the final follow-up scan. Third column shows the IQR in relation to the number of longitudinally repeated scans available per participant.*

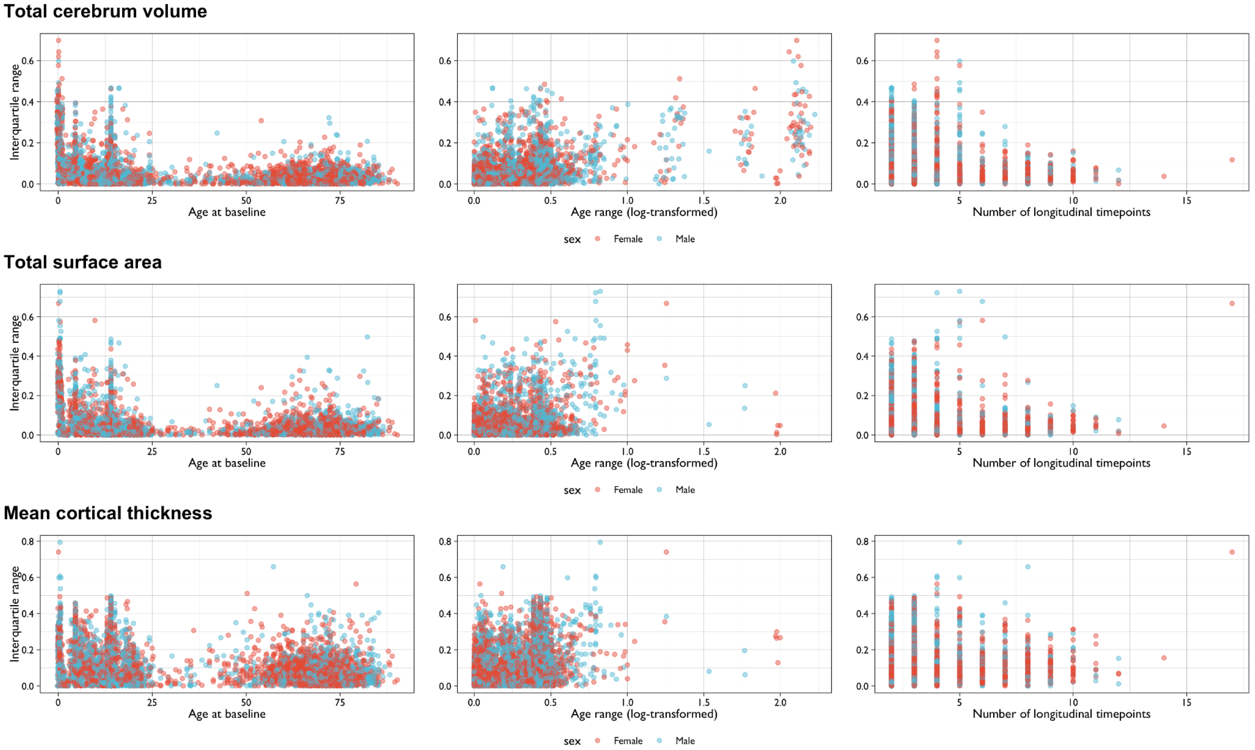


***Fig. S14.1.2. Overview of possible associations between within-subject variation (interquartile range, IQR) of the longitudinal centile scores on extended global MRI phenotypes and likely factors that could influence longitudinal stability of centile scores.*** *First column shows the IQR in relation to the individual's age (in years) at the time of their baseline scan. Second column shows the length of follow-up (in years, log-transformed) between the baseline scan and the final follow-up scan. Third column shows the IQR in relation to the number of longitudinally repeated scans available per participant.*

#### 14.2 Longitudinal variability across studies

In some studies, we observed that within-subject variability was greater compared to other studies over the course of longitudinal scanning. Without robust longitudinal reference data, it is not possible to disentangle whether this is meaningful biological variation or due to non-biological (e.g., technical) confounds. Increased longitudinal variability appeared more prevalent in younger cohorts, further emphasizing the need for robust and consistent high quality imaging data and unified processing protocols[87,88] in that age range. It is important to acknowledge that in contrast to other scientific scenarios, a fully longitudinal neuroimaging study across the human lifespan is practically impossible. However, the current results emphasize the need for longitudinal initiatives that consistently measure the same phenotypes at different intervals across the lifespan.

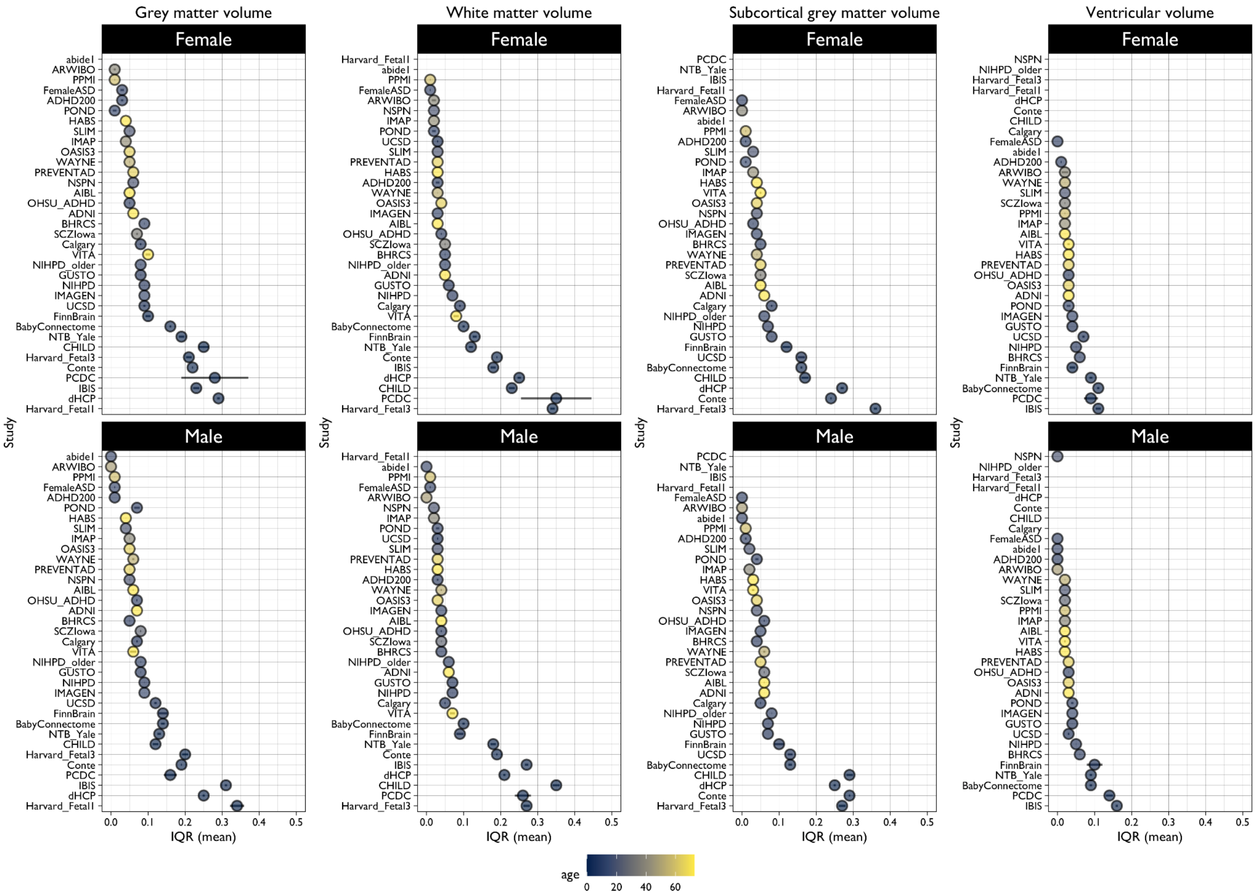


***Fig. S14.2.1. Overview of within-subject variability of longitudinal centile scores (IQR) on cerebrum tissue volumes for each primary study with repeated scans available for participants.*** *Colour scales indicate the mean age for a given study and studies are ordered by mean IQR, from the lowest within-subject variability at the top to the highest within-subject variability at the bottom.*

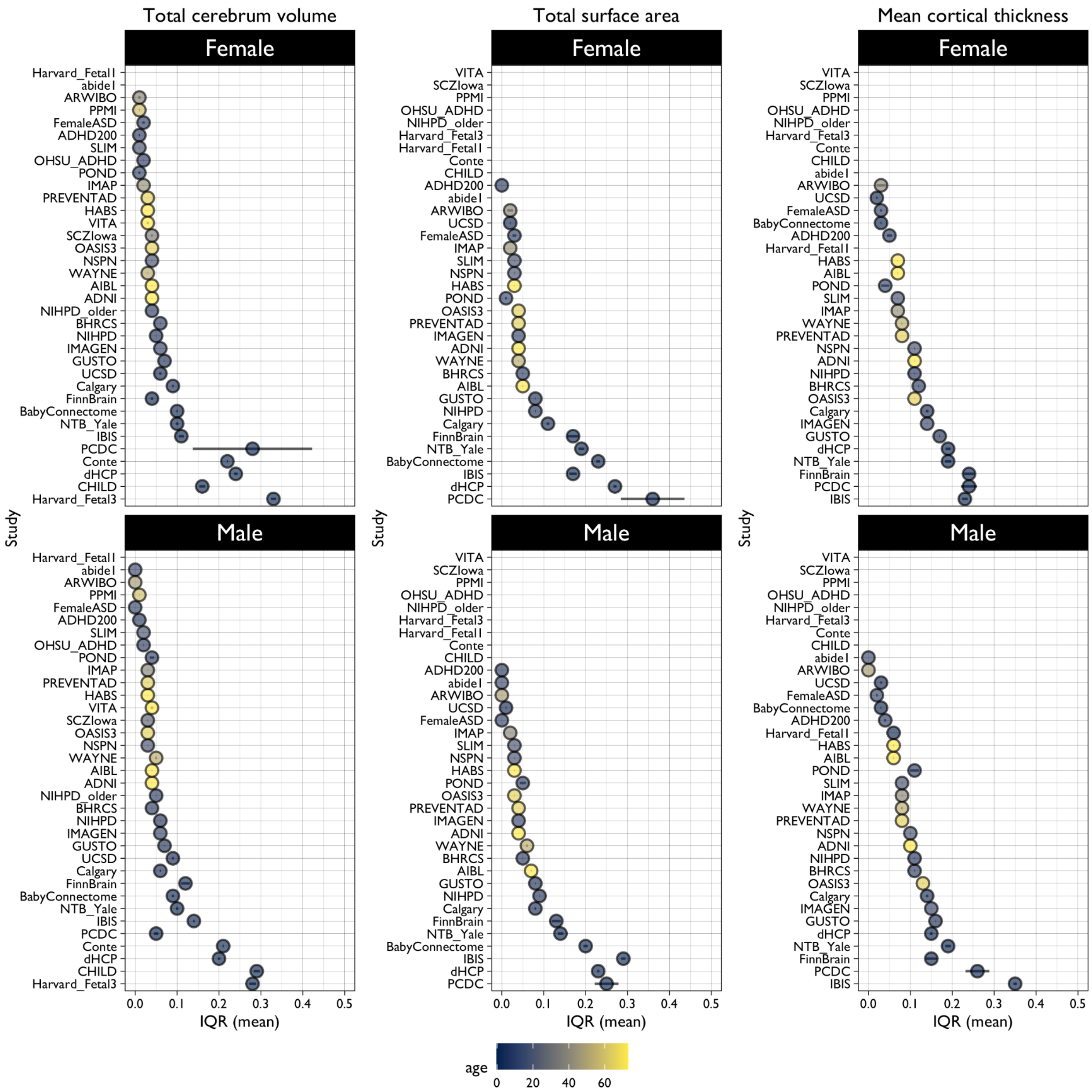


***Fig. S14.2.2. Overview of within-subject variability of longitudinal centile scores (IQR) on extended global MRI phenotypes for each primary study with repeated scans available for participants.*** *Colour scales indicate the mean age for a given study and studies are ordered by mean IQR, from the lowest within-subject variability at the top to the highest within-subject variability at the bottom.*

#### 14.3 Longitudinal variability in clinical samples

Most clinical groups showed significantly lower variability of centile scores across time compared to CN (**Fig. S14.3.1A**). This would suggest that differences in cross-disorder centiles largely persist over time. Indeed, centiles showed lower IQR within clinical groups than within the CN group. However, as described in **SI1.7**, because centiles were derived relative to a CN reference, the IQR within clinical groups may be expected to be compressed compared to the IQR within the CN group.

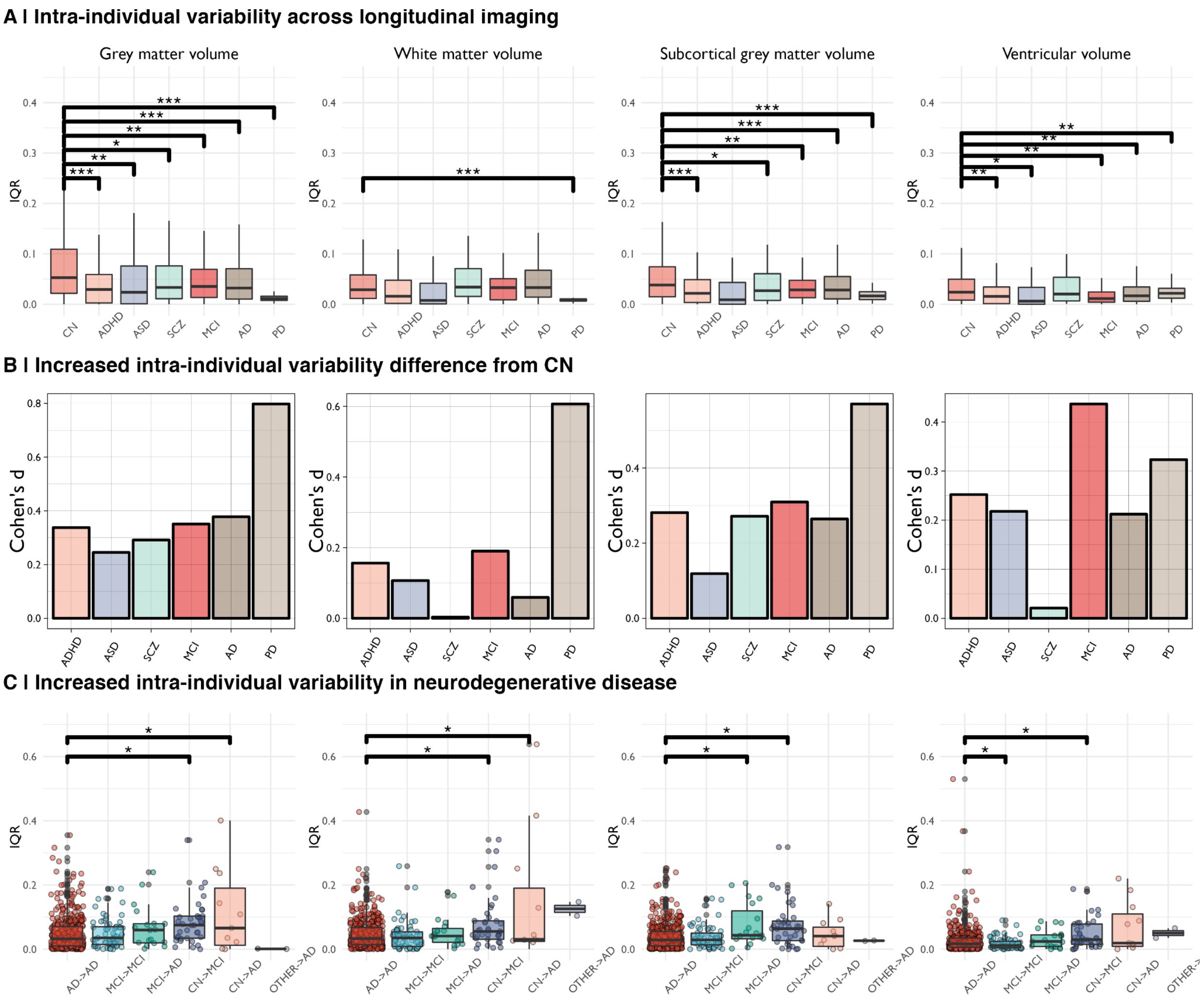


***Fig. S14.3.1. Within-subject variation (IQR) in longitudinal centile scores on cerebrum tissue volumes for control subjects (CN) and 6 clinical disorders.*** *A | All disorders compared pair-wise to the CN group. Asterisks indicate P-values of <0.001 (***), <0.01 (**) and <0.05 (*) as computed using Monte Carlo permutation tests and using the Benjamini-Hochberg[65] procedure to correct for multiple comparisons accounting for all possible pairwise combinations. B | The median difference of within-subject variability between each clinical disorder and the control groups. C | Comparison within-subject variability of longitudinal centile scores for cases of mild cognitive impairment (MCI) or Alzheimer's disease (AD) who maintained the same diagnostic status over the course of repeated scanning (MCI+MCI, AD+AD) and participants who changed diagnostic status (CN→AD, MCI→AD, etc) over the course of longitudinal assessments.*

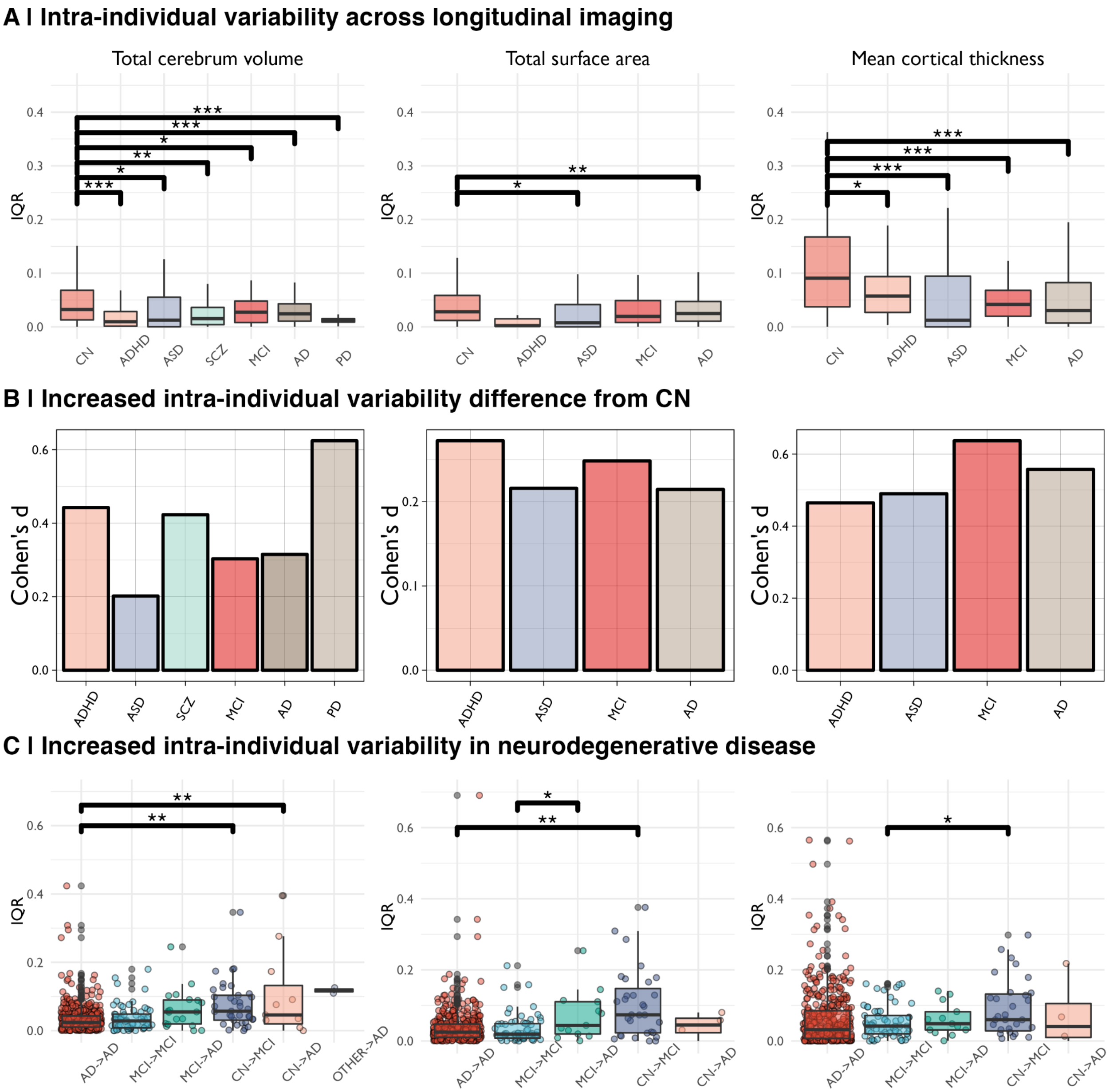


***Fig. S14.3.2. Within-subject variation (IQR) in longitudinal centile scores on extended global MRI phenotypes for control subjects (CN) and 6 clinical disorders.*** *A | All disorders compared pair-wise to the CN group. Asterisks indicate P-values of <0.001 (***), <0.01 (**) and <0.05 (*) as computed using Monte Carlo permutation tests and using the Benjamini-Hochberg[65] procedure to correct for multiple comparisons accounting for all possible pairwise combinations. B | The median difference of within-subject variability between each clinical disorder and the control group. C | Within-subject variability of longitudinal centile scores for cases of mild cognitive impairment (MCI) or Alzheimer's disease (AD) who maintained the same diagnostic status over the course of repeated scanning (MCI+MCI, AD+AD) and participants who changed diagnostic status (CN→AD, MCI→AD, etc) over the course of longitudinal assessments.*

Longitudinal centiles, **Fig. S14.3.1** shows only significant case-control differences in the IQR, although significance was corrected for multiple comparisons using FDR (q < 0.001) across all possible pairs. Statistics for all comparisons are provided in **ST5.1-5.7**, and **Figs. S14.3.3-14.3.4** provide visualisations of all significant pairs.

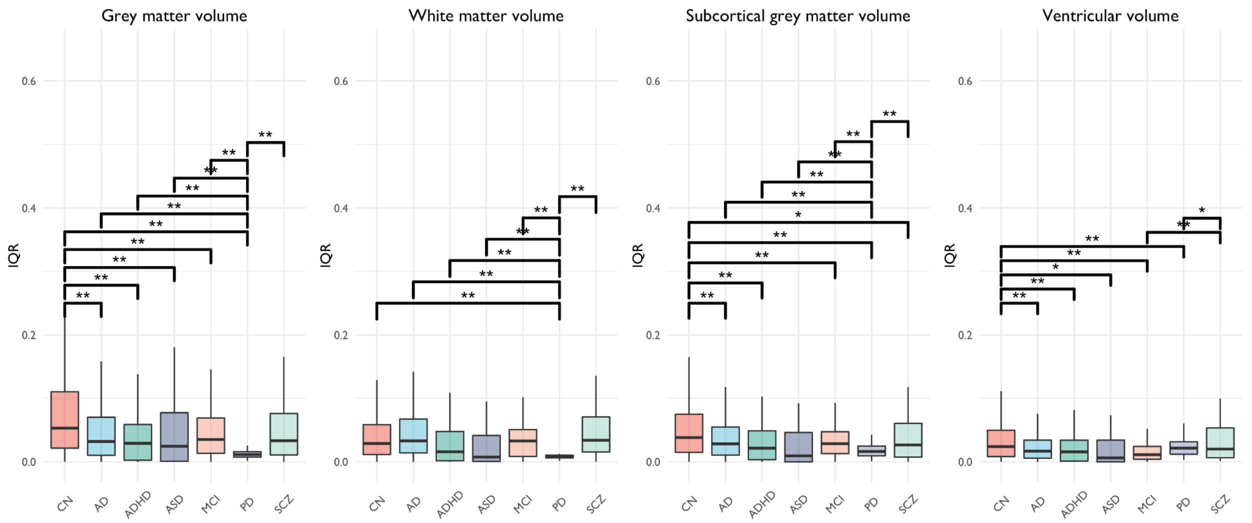


***Fig. S14.3.3. Case-control and between-disorder differences in within-subject variability of centile scores on cerebrum tissue volumes.*** *This is the same figure as depicted in* ***Fig. S14.3.1A****, but including all pairwise comparisons, i.e., not just comparisons relative to the CN group. Asterisks indicate P-values of <0.001 (***), <0.01 (**) and <0.05 (*) as computed using Monte Carlo permutation tests and using the Benjamini-Hochberg[65] procedure to correct for multiple comparisons accounting for all possible pairwise combinations.*

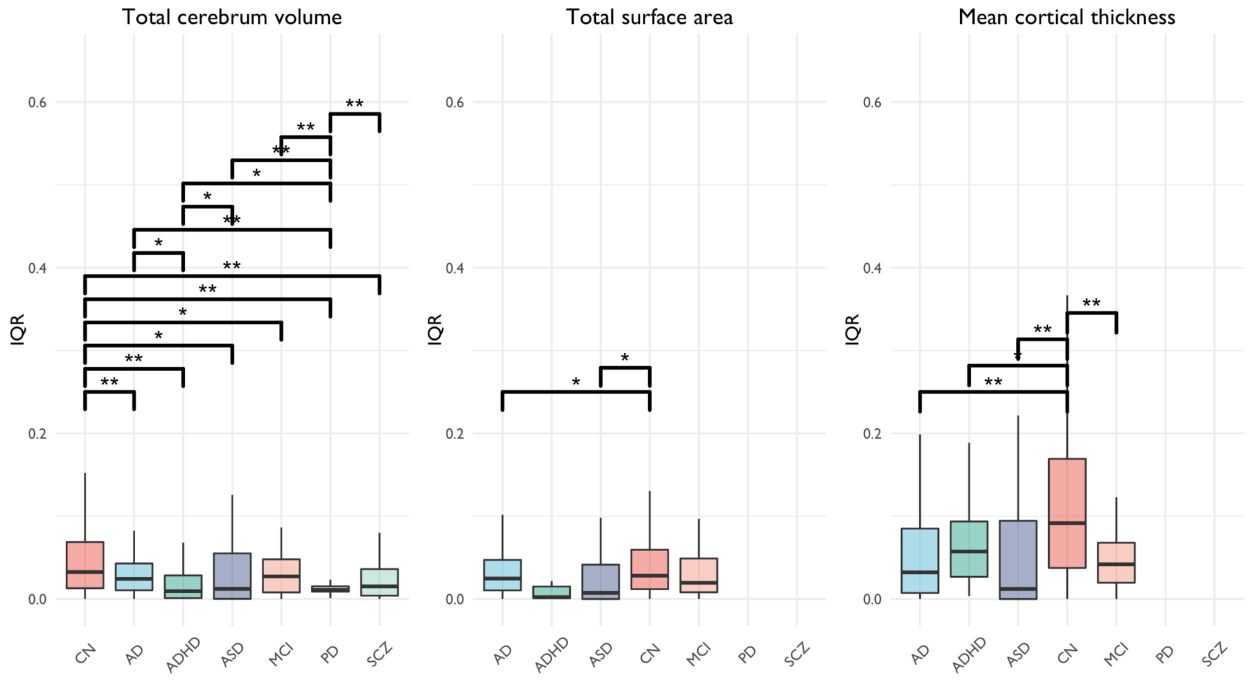


***Fig. S14.3.4. Case-control and between-disorder differences in within-subject variability of centile scores on extended global MRI phenotypes.*** *This is the same figure as depicted in* ***Fig. S14.3.1A****, but including all pairwise comparisons, i.e., not just comparisons relative to the CN group. Asterisks indicate P-*

*values of <0.001 (***), <0.01 (**) and <0.05 (*) as computed using Monte Carlo permutation tests and using the Benjamini-Hochberg[65] procedure to correct for multiple comparisons accounting for all possible pairwise combinations.*

To further visualise the stability of centiles across the lifespan in clinical cohorts, we filtered the longitudinal clinical data using a threshold of 3x the median absolute deviation within the cohort[89]. This filtering was mainly done to clarify the visualisation as the alternative is visually biased toward the few individuals that show an extreme change (which may be related to measurement error). We observed that between time points, individuals generally either stay within ~10% of their first centile measurement, or they progress towards the extreme end of the distribution (**Fig. S14.3.5**). For example, males with ASD tend to drop in their GMV centile scores in early development, consistent with the cross-sectional analysis showing that at the group level males with an ASD have lower GMV centile scores. A similar picture occurs for AD and MCI. For ADHD the changes appear more mixed. Note that in the CHILD study, individuals that were considered high risk for ASD are included here as ASD cases (hence the impression of individuals diagnosed with ASD in utero).

**Grey matter volume**

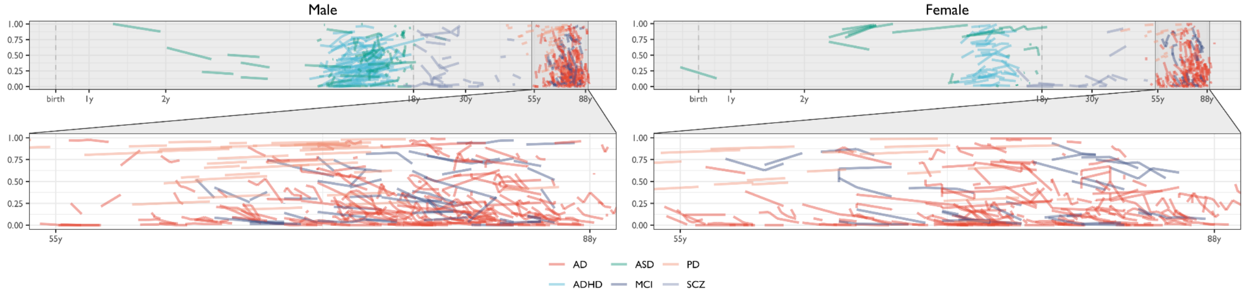


**White matter volume**

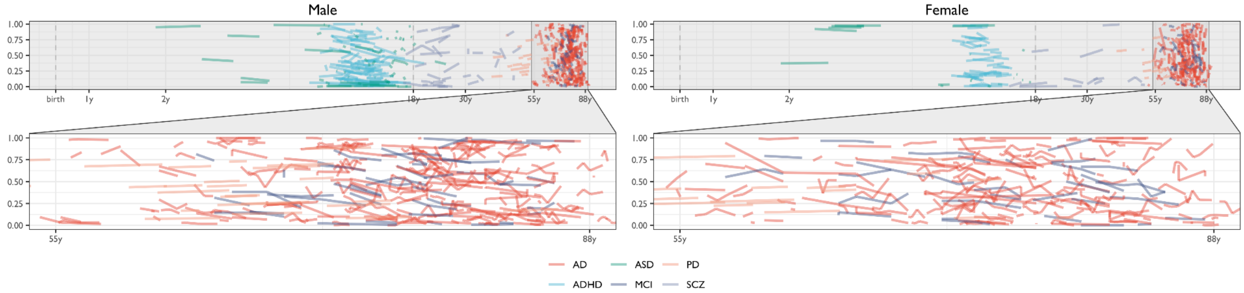


**Subcortical grey matter volume**

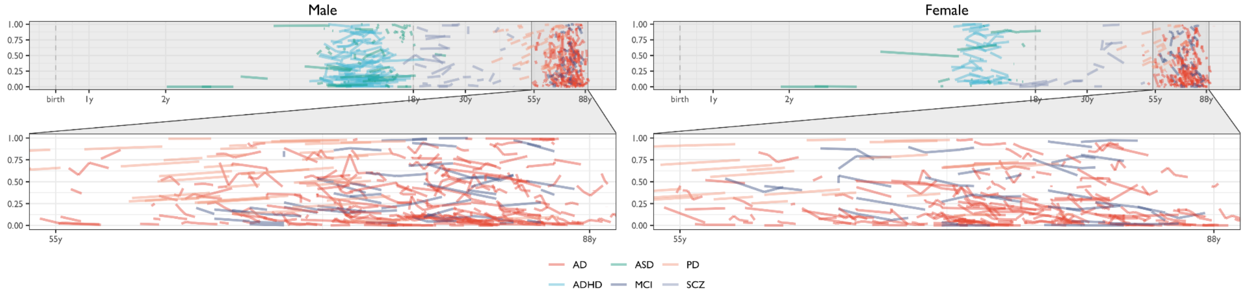


**Ventricular volume**

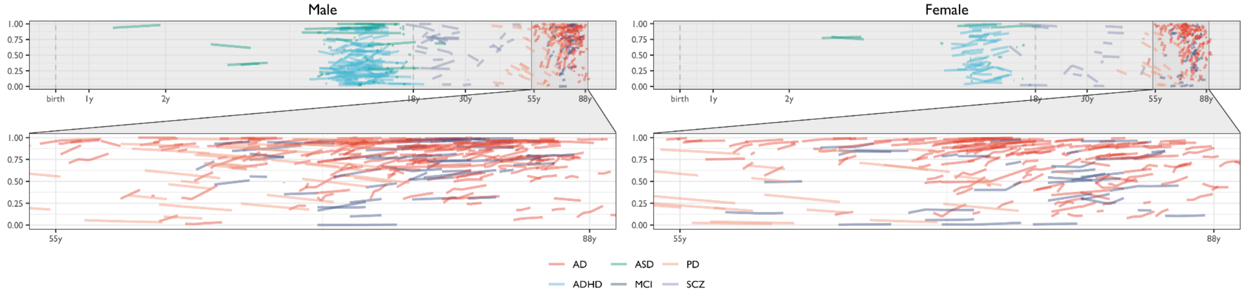


***Fig. S14.3.5. Within-subject changes in longitudinal centile scores on cerebrum tissue volumes.*** *Each line represents a single individual and their respective centile changes across the lifespan. Zoomed-in panels emphasise the age-span beyond 55-years as a large proportion of longitudinal data included individuals with Alzheimer's disease and were repeatedly scanned in this phase of the lifespan.*

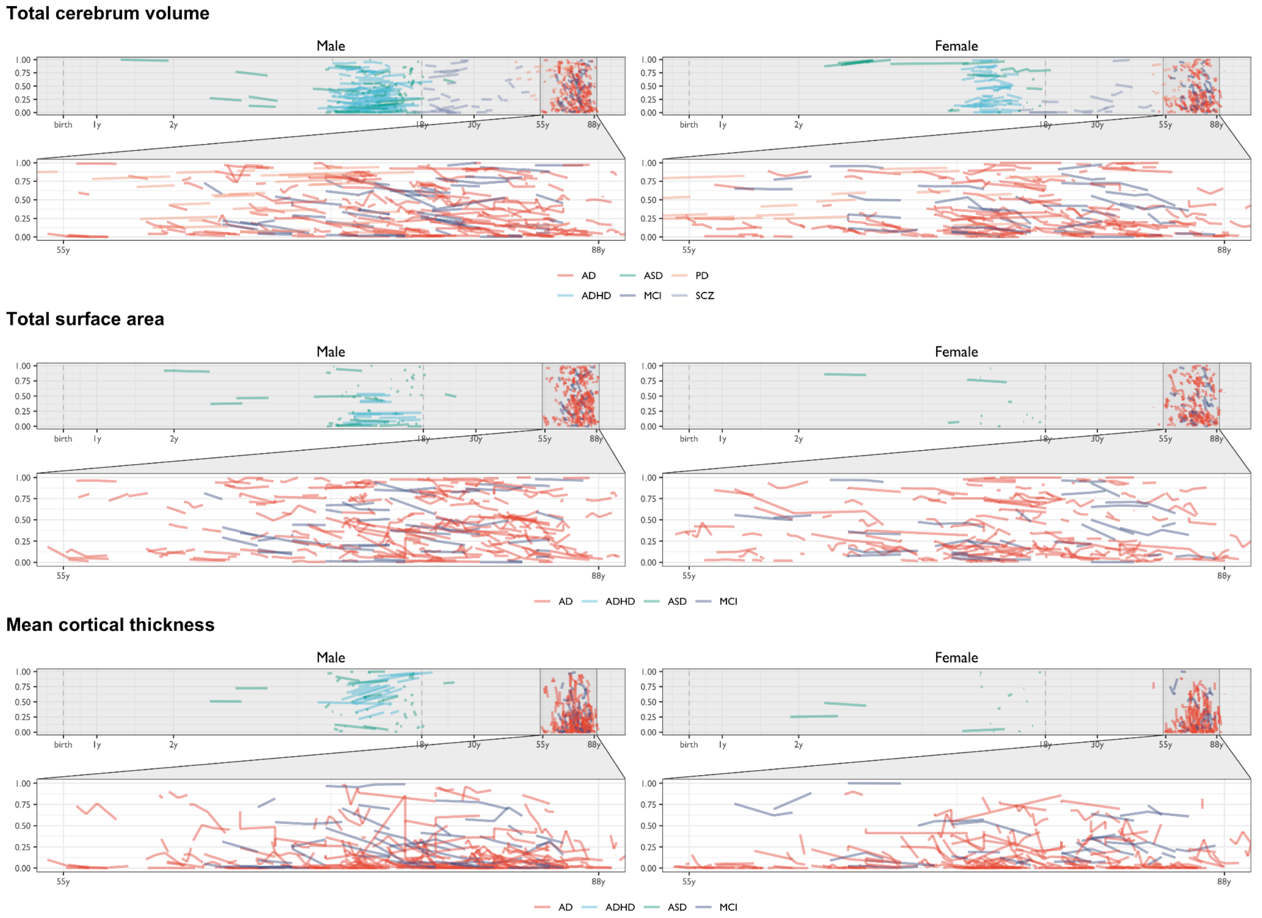


***Fig. S14.3.6. Within-subject changes in longitudinal centile scores on extended global MRI phenotypes.*** *Each line represents a single individual and their respective centile changes across the lifespan. Zoomed-in panels emphasise the age-span beyond 55-years as a large proportion of longitudinal data included individuals with Alzheimer's disease and were repeatedly scanned in this phase of the lifespan.*

#### 14.4 Longitudinal centile score changes and diagnostic progression

Similar to paediatric growth charts, further value from having age-appropriate standardised reference curves will likely come from the ability to more reliably detect atypical longitudinal changes in brain changes within individuals. As an example of this approach, we have tracked centile scores in longitudinal (repeated) cerebrum tissue volumes for a large cohort of older individuals, some of whom transitioned between diagnostic categories during the period of longitudinal follow-up from CN to MCI (CN → MCI), from CN to AD (CN → AD), or from MCI to AD (MCI → AD). Interestingly, in contrast to the lower within-subject variability (IQR) of cases compared to healthy controls in general, there was a reverse trend of increased within-subject variation in cerebrum tissue volumes (especially GMV and Ventricles) in the subset of cases that changed diagnostic status. Specifically, there was faster than normal decrease of grey matter volume, and faster than normal increase of ventricular CSF volume, among participants who transitioned from CN or MCI to AD over the course of repeated scanning (**Fig. S14.4.1** and **ST6.1-6.7**).

Analysis of within-subject changes in centile scores focused on individuals with the most frequently observed diagnostic transitions, all in the direction of greater severity or disability: from CN to MCI, from CN to AD and from MCI to AD (**ST6**). The longitudinal change in centile scores occurred in the same direction as predicted by the cross-sectional case-control differences (compare **Fig. 4A** and **SI10 "Clinical applications of centile scores"**). We rescaled the longitudinal data to generate a group-level trajectory for each transition (CNI → AD, CN → MCI, and MCI → AD) using linear mixed effects models. As shown in **Fig. S14.4.1,** all clinical transitions were associated with significantly increased rates of cortical and subcortical grey matter loss, and ventricular CSF volume expansion — both reflected by decreases in centile scores. Because the significant age-related changes expected in healthy older individuals are incorporated into the reference norms, centile scores provide a clear indication of a change in trajectory for individuals with neurodegenerative disease.

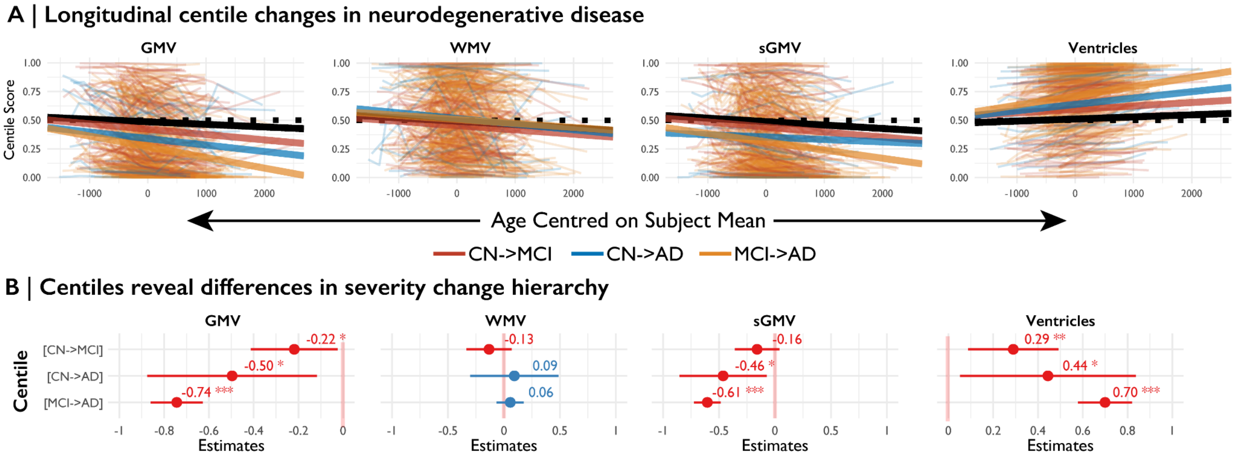


***Fig. S14.4.1. Longitudinal changes in centile scores are associated with diagnostic transitions between the groups of healthy controls (CN), mild cognitive impairment (MCI), and Alzheimer's disease (AD).*** *A | Shows the within-subject changes in centile scores for CN→MCI, CN→AD, and MCI→AD, with the dotted black lines showing the median slope for all controls that had longitudinal measurements and the solid black lines showing the median slope for all controls from the datasets that contributed to the diagnostic change group. B | Shows the model fixed effects standardised coefficients (e.g., model fixed effects divided by two standard deviations, to denote the slope differences in longitudinal changes in centile scores between the groups. Asterisks indicate the level of uncorrected significance (* is P<0.05, ** is P<0.01, *** is P<0.001) as tested with a linear mixed model restricted maximum likelihood (REML) fit that included a subject-level random effect, but not a random intercept as some individuals only had 2 observations and including random slopes would cause convergence issues. The error bars in panel B depict the confidence intervals around the beta coefficients. These results show that for both GMV and Ventricular CSF the rate-of-change in centile scores is significantly greater in individuals undergoing a clinically documented transition (from less to more severe diagnostic categories)*

#### 15. Interactions between cerebrum tissue volumes

It has been hypothesized that age-varying cellular processes could be captured by neuroimaging milestones, in terms of the growth trajectories of relative volumetric measurements[90]. In line with these expectations, we found an initial postnatal increase in GMV relative to WMV, likely due to increased complexity of neuropil including synaptic proliferation[91,92]. Subsequently, GMV declined relative to WMV (**SI9.2 "Grey-white matter differentiation"**), likely due to both continued myelination and synaptic pruning[93]. To further explore the patterning of tissue interactions, we performed supplementary analyses to empirically assess the correlations between global tissue classes. **Fig. S15.1** presents these inter-relationships as Pearson's correlation coefficients between each pair of global brain MRI phenotypes across participants within each study. These results highlight the variability of these relationships across studies (which themselves vary in terms of technical and biological variables – see **Fig. 1A, ST1.1**). However, it is also clear that there are generally high correlations between grey and white matter volumes and surface area (SA). Comparatively, GMV and WMV are less strongly correlated with CT and CSF. Additionally, we substantiated the prior consensus in the literature concerning the orthogonality of CT and SA by finding that these two global metrics were not correlated with each other (**Fig. S15.1**).

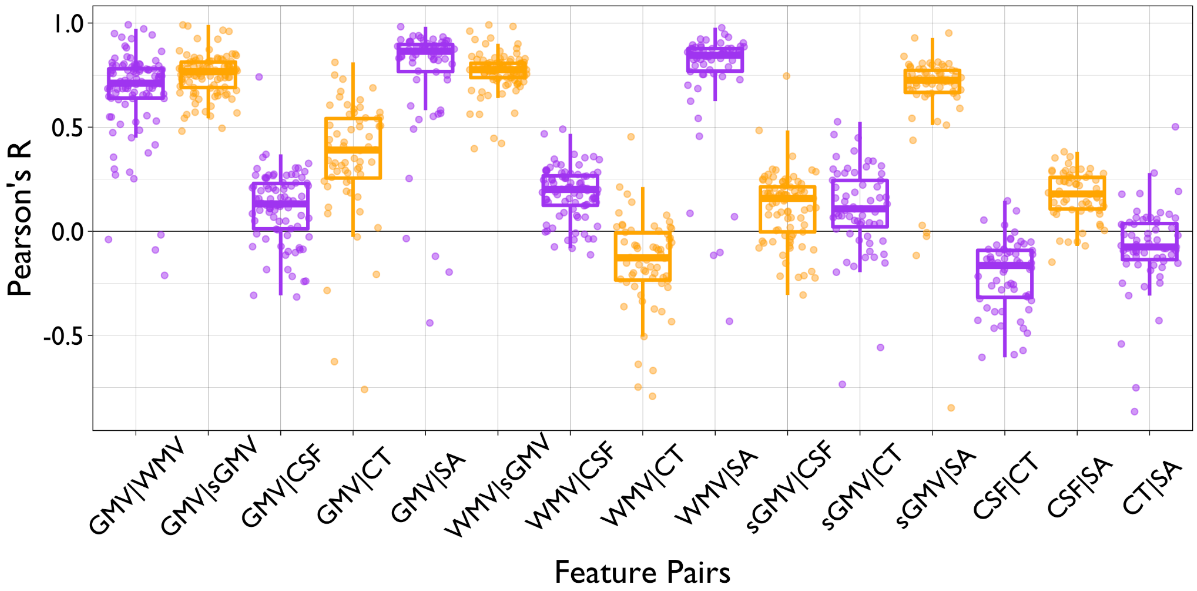


***Fig. S15.1. Box-plots of Pearson correlations between each pair of global neuroimaging metrics in each of the primary studies in the reference dataset.*** *Each datapoint represents a single primary study; boxes highlight the median and interquartile range of between-study variation in correlations of "raw", non-centiled volumetrics for all possible pairs of global MRI phenotypes. Alternating colours are for visualisation purposes only.*

Given these findings in the context of each study in our aggregated dataset, we examined the same inter-relationships between phenotypes across age, in line with previous work examining regional correlations of diffusion-weighted imaging phenotypes across age[94]. We used a sliding window approach to apply this framework to global MRI phenotypes, binning segments of the lifespan based on age (each window = 300 days, sliding by 50 days). Pearson's correlation between phenotypes was then calculated within each bin, and locally-weighted (LOESS) regression was used to fit a nonlinear curve to the age-related changes in each pair-wise phenotypic correlation (**Fig. S15.2**). These results recapitulate some of the findings of the correlational analyses within each primary study, e.g., the GMV/WMV correlation is consistently more strongly positive than the CT/SA correlation. However, there are also some age-related shifts in the strength and/or sign of these phenotypic correlations, especially in late gestation and early postnatal life, that will be interesting to investigate in more detail as additional early-life MRI data become available in future.

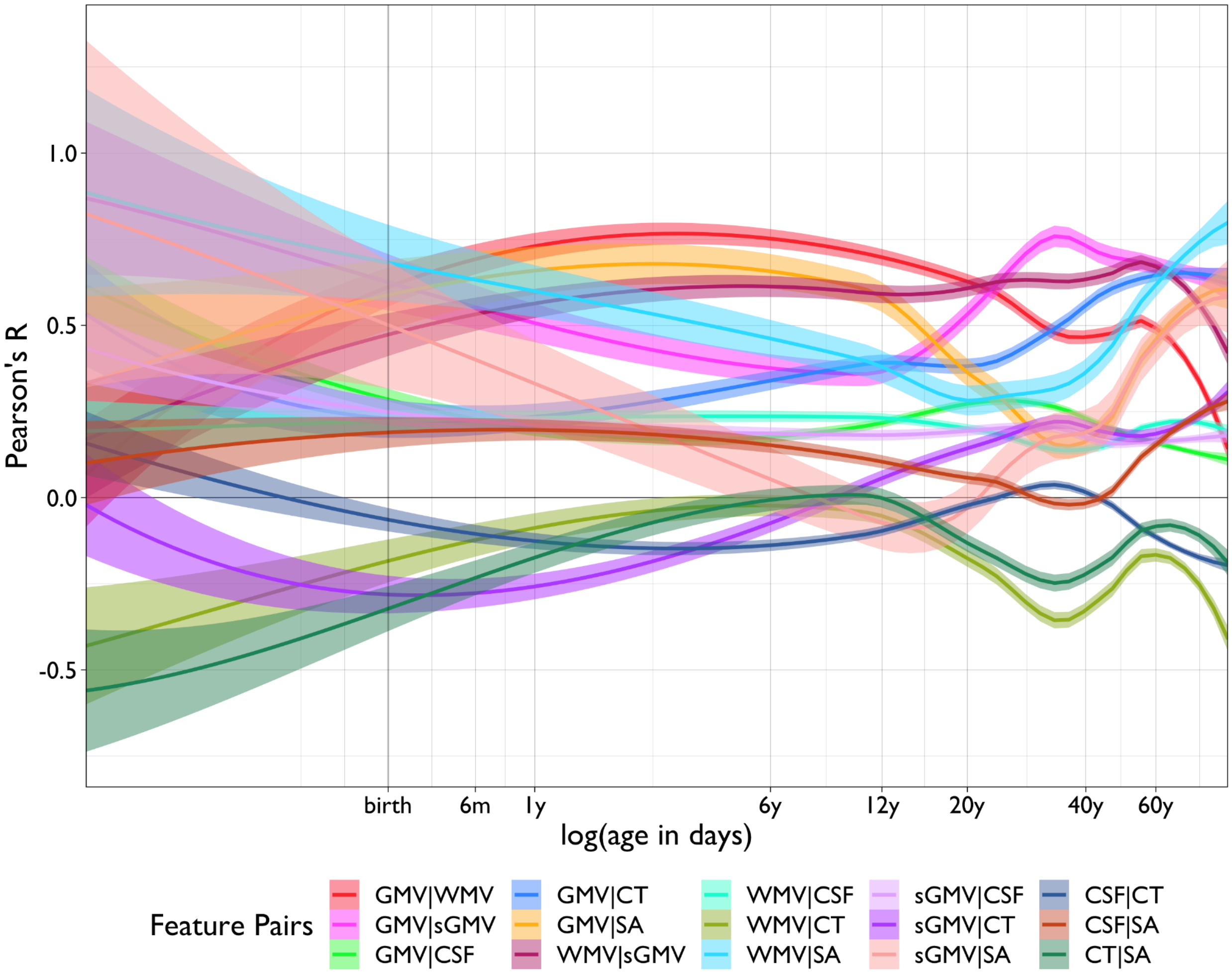


***Fig. S15.2. Sliding-window analysis of age-related changes in pairwise correlations for all possible pairs of 7 global MRI phenotypes (4 cerebrum tissue volumes and 3 extended global MRI phenotypes) over the course of the lifespan.*** *We used a window size of 300 days, sliding by 50 days. Plotted lines are colour-coded by pairwise correlation and represent the fitted lines and 95% confidence intervals from locally-weighted (LOESS) regression for each correlated pair of phenotypes.*

#### 16. Sex differences

In line with prior literature[95] we find that variance in males is higher than in females across imaging phenotypes. These variance differences across development demonstrate the importance of modelling age and sex-related differences in variability in addition to absolute size. Sex differences in brain development in relation to developmental psychopathology are an active field of research. Particularly in the case of ASD, ADHD, MDD, and SCZ, sex differences have been proposed to be of major importance for aetiology[96,97] and a key driver of the clinically observed sex bias in diagnoses. While we did not explicitly model sex differences beyond their main additive effects, the present results indicate that this could be an important area for future research, especially given differences in finer regionally-specific profiling of brain morphology[98]. For example, in light of the mixed literature on sex differences in SCZ including neuroanatomy[99,100], a post-hoc analysis showed a significant effect of sex in SCZ centiles—with biologically female SCZ patients showing significantly greater deviations in GMV, WMV, and sGMV (12.3%, 8.2%, 9.2%, respectively; all $P<0.01$ after permutation testing scrambling group labels; **ST8**).

It will be important for future studies to further refine the possible developmental interactions with biological sex, as current work already highlights it importance for mental health[60,101]. The GAMLSS approach could be adapted to study sex-by-development interactions. As noted in the Main Text, it important to stress that there are no known clinical, cognitive or behavioural implications for differences in brain size related to sex. It is also important to note the limitations of binary classification and conflation of sex and gender present in almost all available imaging data[102].

### Reference database details: demographics and processing pipelines

This section provides an overview of primary datasets currently included in the aggregated reference dataset.

#### 17. Demographics of reference database

**Fig. S17.1** shows we have curated data from across the world, with the majority of studies coming from Europe and North America. **Fig. S17.2** shows that we have strong representation across the lifespan with 500 or more subjects in each of the Kang- defined developmental windows[59] from 15 post-conception weeks (pcw) onwards. Interestingly, in addition to early development, the age-range between 30 and 40 years is particularly under-represented in currently available data.

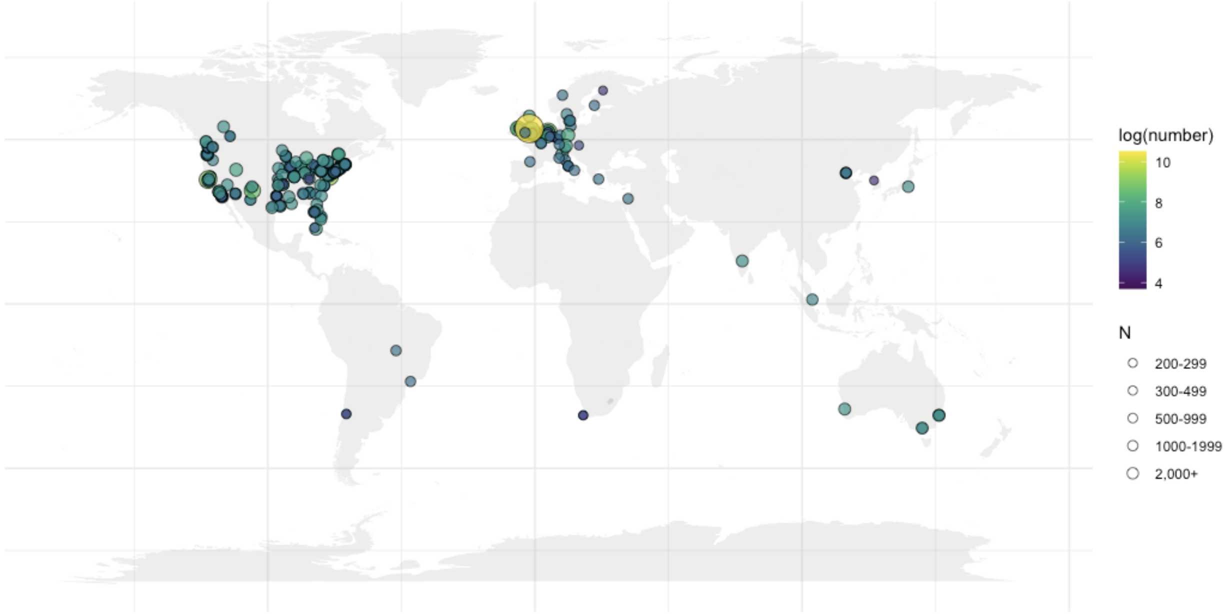


***Fig. S17.1. Locations of primary datasets included in the aggregated, reference database for the Lifespan Brain Chart project.***

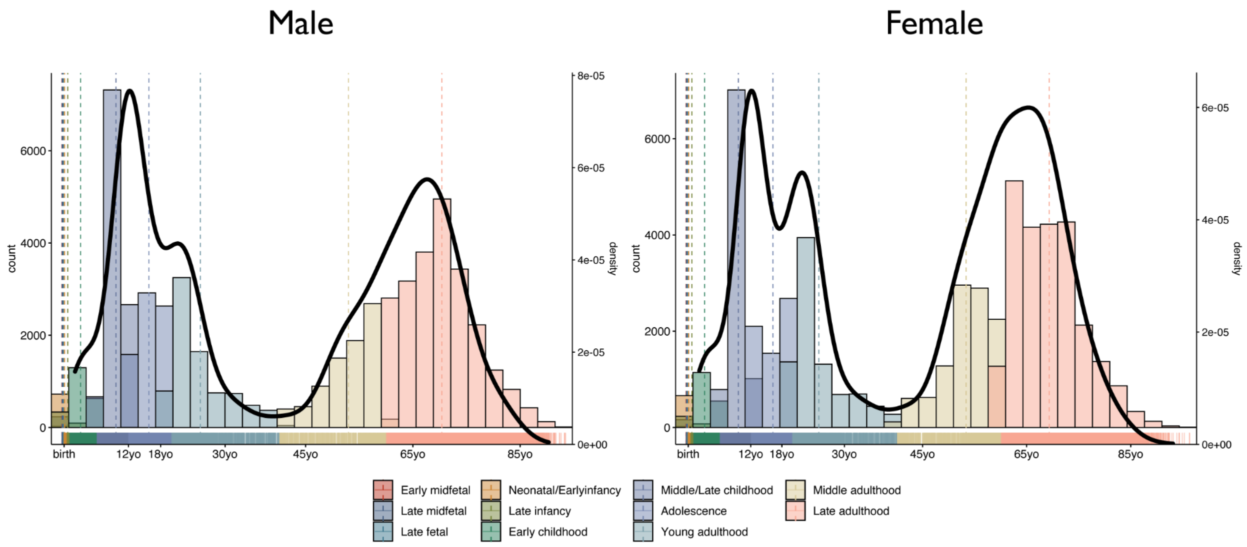


***Fig. S17.2. Histograms and probability density plots of age of participants in the reference database, stratified by sex.***

#### 18. Data processing

If T1- and T2/FLAIR-weighted raw data were available, as they were for approximately 95% of scans), these data were processed on the same server at the University of Cambridge with FreeSurfer 6.0.1[32] using the combined T1-T2 recon-all pipeline for improved grey-white matter boundary estimation. If only raw T1-weighted data were available, and subjects were aged over 2 years, data were processed with FreeSurfer 6.0.1 using the standard recon-all pipeline. If subjects were aged 0–2 years, data were processed with Infant FreeSurfer v1[103]. Briefly, the first processing stage of recon-all includes: non-uniformity correction, projection to Talairach space, intensity normalisation, skull-stripping, automatic tissue and subcortical segmentation. Subsequently, surface interpolation, tessellation and registration are done at the second and third stages of the recon-all pipeline. **ST1.1** lists the number of subjects per site per processing pipeline alongside their respective MRI acquisition and quality control protocols. We noticed that Infant FreeSurfer estimated total subcortical grey matter volume (sGMV) differently from other pipelines included in this dataset, while other cerebrum tissue volumes were estimated consistently across pipelines. We therefore excluded scans processed with Infant FreeSurfer from growth curve estimation for subcortical GMV. All four cerebrum tissue volumes were extracted from the aseg.stats files output by the recon-all process: 'Total cortical gray matter volume' for GMV; 'Total cortical/cerebral (FreeSurfer version dependent) white matter volume' for WMV; ‘Subcortical gray matter volume’ for sGMV (inclusive of thalamus, caudate nucleus, putamen, pallidum, hippocampus, amygdala, and nucleus accumbens area; https://freesurfer.net/fswiki/SubcorticalSegmentation); and the difference between 'BrainSegVol" and 'BrainSegVolNotVent' for Ventricular volume. Regional volume was estimated for each of 34 bilaterally averaged cortical regions defined by the Desikan-Killiany[57] parcellation template following the final stages of the recon-all pipeline and using the hemisphere-specific aparc.stats files generated by FreeSurfer.

#### 19. Primary dataset descriptions

##### 3R-BRAIN - Brain Consortium for Reproducibility, Replicability and Reliability

The 3R-BRAIN is built by the Chinese Academy of Sciences (CAS) and aims to recruit a total of 200 adults (20-35 years), with each participant visiting three ultra-high field (two 3T and one 7T) MRI scanners located at the CAS Institute of Psychology (GE-MR750) and the CAS Institute of Biophysics (SMS-Prisma and SMS-Terra). Each participant received 5 repeated scans including two-week test-retest data at GE-MR750 and SMS-Prisma, respectively, as well as a single scan at SMS-Terra. This represents an open resource for reproducible, replicable and reliable brain research and imaging cognitive neuroscience [104]. In the present study all the first scans are included and all raw structural T1-weighted scans were processed with FreeSurfer 6.0.1.

##### ABCD - Adolescent Brain and Cognitive Development

The ABCD Study is a landmark, longitudinal study of brain development and child health[50]. Investigators at 21 sites around the country will measure brain maturation in the context of social, emotional and cognitive development, as well as a variety of health and environmental outcomes. Minimally processed T1 and T2 weighted imaging for 10,588 individuals was downloaded through the NIMH Data Archive (NDAR) and processed using FreeSurfer 6.0.1 using the combined T1-T2 processing pipeline when both modalities were available (and conventional T1 when no T2 was available). Individuals were included in the reference model as healthy controls (CN) based on the parental response to the ABCD screening and risk questionnaire (https://nda.nih.gov/data_structure.html?short_name=abcd_screen01) indicating the individual had never been diagnosed with a mental health disorder.

##### ABIDE - Autism Brain Imaging Data Exchange

The Autism Brain Imaging Data Exchange (ABIDE) initiative has aggregated functional and structural brain imaging data collected from laboratories around the world to accelerate our understanding of the neural bases of ASD[105,106]. All T1-weighted structural data from this collection was processed using recon-all as implemented in FreeSurfer 6.0.1.

##### ABVIB - Aging Brain: Vasculature, Ischemia, and Behavior

This study is based on a prospective, longitudinal cohort study started in 2008 known as the Aging Brain: Vasculature, Ischemia, and Behavior Study (ABVIB; https://ida.loni.usc.edu/login.jsp?project = ABVIB)[107]. The primary goal of ABVIB was to assess the contributions of cardiovascular risk factors (laboratory studies) and cerebrovascular disease (carotid intima media thickness and retinal vessels) to brain structure and function, alone or in combination with AD. Measures of brain structure and function included serial MRI and neuropsychological testing. Exclusion criteria were: age younger than 55 years, non-Englishspeaking, cortical strokes, severe illnesses other than cardiovascular or dementia, and use of medications that affect cognition. A total of 280 participants completed the neuropsychological assessments as well as the brain imaging. For the purpose of this study, here we only included data from those participants who had valid brain data[107].

#### ACE and IBIS

The ACE and IBIS cohorts included in the reference database were aggregated from several NDAR projects: Longitudinal MRI Study of Infants at Risk for Autism (19), Biomarkers of Developmental Trajectories and Treatment in ASD (2026) and A Longitudinal MRI Study of Infants at Risk for Autism (ACE 2; 2027). Structural T1 weighted imaging was processed with Infant FreeSurfer[103] for individuals younger than 36 months, while individuals older than 36 months were processed with FreeSurfer 6.0.1.

#### ADHD200

The ADHD-200 Sample is a grassroots initiative, dedicated to accelerating the scientific community's understanding of the neural basis of ADHD through the implementation of open data-sharing and discovery-based science. It includes the unrestricted public release of 776 anatomical datasets aggregated across 8 independent imaging sites, 491 of which were obtained from typically developing individuals and 285 in children and adolescents with ADHD (ages: 7–21 years old). Accompanying phenotypic information includes: diagnostic status, dimensional ADHD symptom measures, age, sex, intelligence quotient (IQ) and lifetime medication status. Preliminary quality control assessments (usable vs. questionable) based upon visual timeseries inspection are included for all resting state fMRI scans. T1-weighted structural data was processed using recon-all as implemented in FreeSurfer 6.0.1.

#### ADNI - Alzheimer's Disease Neuroimaging Initiative

Data used in the preparation of this article were obtained from the Alzheimer’s Disease Neuroimaging Initiative (ADNI) database (adni.loni.usc.edu). The primary goal of ADNI has been to test whether serial magnetic resonance imaging (MRI), positron emission tomography (PET), other biological markers, and clinical and neuropsychological assessment can be combined to measure the progression of MCIand early AD. For up-to-date information, see www.adni-info.org. MRI data is one component of the comprehensive data set collected in ADNI participants. ADNI began in 2004 and to date 3 different phases of ADNI have been undertaken. The MR protocol evolved over these 3 phases. The MRI protocol for ADNI1 (2004–2009) focused on consistent longitudinal structural imaging on 1.5T scanners using T1- and dual echo T2-weighted sequences. One-fourth of ADNI 1 subjects were also scanned using essentially the same protocol on 3T scanners. In ADNI-GO/ADNI2 (2010-2016), imaging was performed at 3T with T1-weighted imaging parameters similar to ADNI1. In place of the dual echo T2-weighted image from ADNI1, 2D FLAIR and T2*-weighted imaging was added at all sites. Both fully sampled and accelerated T1-weighted images were acquired in each imaging session. ADNI 3 imaging is being done exclusively on 3T scanners. Nearly all of the imaging sequences from ADNI2 have been updated for inclusion in ADNI 3. Each of the ADNI 2advanced imaging sequences is now included in the basic ADNI 3 protocol with a few site-wise exceptions related to sequence license issues. For the present study all structural T1-weighted scans were processed with FreeSurfer 6.0.1.

#### AIBL - Australian Imaging. Biomarkers and. Lifestyle Flagship Study of Ageing

The Australian Imaging, Biomarker & Lifestyle Flagship Study of Ageing (AIBL) is a study to discover which biomarkers, cognitive characteristics, and health and lifestyle factors determine subsequent development of symptomatic AD. The AIBL MRI and PiB images were acquired with ADNI protocols. Data was collected by the AIBL study group. AIBL study methodology has been reported previously[108]. T1-weighted structural data was processed with FreeSurfer 6.0.1.

#### AOBA

The subjects were Japanese volunteers recruited by the Aoba Brain Imaging Research Center, Sendai, Japan. All were normal and right-handed. The group consisted of 772 men (age range: 16–79 years) and 775 women (age range: 18–79 years). Data from subjects who had a history or symptoms of a central nervous system disease of any kind or brain injury were excluded from the database. Each subject was interviewed by medical doctors of the Institute of Development, Aging and Cancer (IDAC), Tohoku University. For the present study raw structural T1-weighted scans were processed with FreeSurfer 6.0.1.

#### AOMIC ID1000, PIOP1 & PIOP2 - Amsterdam Open MRI Collection

The Amsterdam Open MRI Collection (AOMIC) consists of three datasets with multimodal (3T) MRI data including structural (T1-weighted), diffusion-weighted, and (resting-state and task-based) functional BOLD MRI data, as well as detailed demographics and psychometric variables from a large set of healthy participants (N=928, N=226, and N=216). Data from all three datasets were scanned on the same Philips 3T scanner (Philips, Best, the Netherlands), but underwent several upgrades in between the three studies. The ID1000 dataset was scanned on the 'Intera' version, after which the scanner was upgraded to the 'Achieva' version (converting a part of the signal acquisition pathway from analog to digital) on which the PIOP1 dataset was scanned. After finishing the PIOP1 study, the scanner was upgraded to the 'Achieva dStream'' version (with even earlier digitisation of the MR signal resulting in less noise interference), on which the PIOP2 study was scanned. All studies were scanned with a 32-channel head coil (though the head coil was upgraded at the same time as the dStream upgrade). Full dataset description and methods are found in Snoek et al.[109] All T1-weighted structural data was processed using FreeSurfer 6.0.1.

#### ARWIBO - Alzheimer's disease Repository Without Borders

Data used in the preparation of this article were obtained from the Alzheimer's Disease Repository Without Borders (ARWiBo) (www.arwibo.it). The primary aim of ARWiBo is to publish all clinical, neuropsychological, EEG, neuroimaging, and biological data of patients with neurodegenerative diseases and CN individuals collected in over 10 years by a number of researchers of IRCCS Fatebenefratelli, Brescia, Italy[110,111]. The overall goal of ARWiBo is to contribute, through synergy with neuGRID (https://neugrid2.eu), to global data sharing and analysis in order to develop effective therapies, prevention methods and a cure for AD and other neurodegenerative diseases. All structural T1-weighted scans were processed with FreeSurfer 6.0.1.

#### ASRB - Australian Schizophrenia Research Bank

The Australian Schizophrenia Research Bank (ASRB) is a comprehensive biobank of clinical, neuroimaging and genetic data acquired in individuals with schizophrenia and healthy comparison individuals. Certain data comprising the biobank can be accessed subject to approval of the ASRB Access Committee. Participants were recruited from five sites in Australia, with all sites implementing the same recruitment procedures and MRI acquisition protocols. Exclusion criteria included any neurological disorder, history of brain trauma followed by a long period of amnesia (>24 h), intellectual disability (full-scale IQ<70), current drug or alcohol dependence, as well as electroconvulsive therapy in the past 6 months. Patients had a confirmed diagnosis of schizophrenia or schizoaffective disorder, based on DSM-IV criteria. Structural and diffusion-weighted MRI scans of brain anatomy were acquired using Siemens Avanto MRI scanners located in Melbourne, Sydney, Brisbane, Perth and Newcastle. The same acquisition sequence was used at all sites. An individual traveled to all five sites and was scanned at each site to quantify gross inter-site differences. A Siemens MRI phantom was also scanned at each site to enable inter-site calibration. This calibration was done prior to MRI acquisition to help minimize potential inter-site variability. Structural T1-weighted images were acquired using an optimized MPRAGE sequence (voxel resolution: 1 $mm^3$ isotropic, TR: 1980 ms, TE: 4.3 ms). Participants showing gross artefacts, cerebellar cropping and/or significant head motion were excluded, following protocols established as part of a prior study in this cohort. For the current study, the T1-weighted scans were processed using FreeSurfer 6.0.1. The reconstructed cortical surfaces were not manually edited. Approval to contribute ASRB data to this study was granted by the ASRB Access Committee on December 17, 2020. The ASRB is funded by a National Health and Medical Research Council (NHMRC) Enabling Grant (386500; Carr V, Schall U, Scott R, Jablensky A, Mowry B, Michie P, Catts S, Henskens F, Pantelis C, Loughland C), and the Pratt Foundation, Ramsay Health Care, the Viertel Charitable Foundation, and the Schizophrenia Research Institute, using an infrastructure grant from the NSW Ministry of Health.

#### BCP - Baby Human Connectome Project

The Baby Connectome Project (BCP) is a four-year study of children from birth through five years of age, intended to provide a better understanding of how the brain develops from infancy through early childhood and the factors that contribute to healthy brain development. Data is collected at two sites, UNC and UMN, using a Siemens 3T Prisma (32-channel coil) at both sites. The imaging modalities are structural, diffusion, and functional (resting state). For the present study all structural T1-weighted scans were processed with a customised version of FreeSurfer 6.0.1 and has undergone quality control. For more information on the BCP see Howell et al.[112]

#### BGSP - Brain Genomics Superstruct Project

The Brain Genomics Superstruct Project Open Access Data Release[113,114] comprises a carefully vetted collection of neuroimaging, behaviour, cognitive, and personality data for over 1,500 human participants. Each neuroimaging data set includes one high-resolution MRI acquisition and one or more resting-state functional MRI acquisitions. Each functional acquisition is accompanied by a fully-automated quality assessment and pre-computed brain morphometrics. For the present study raw structural T1-weighted scans were processed with FreeSurfer 6.0.1.

#### BHRCS - Brazilian High Risk Cohort Study for Mental Conditions

The Brazilian High Risk Cohort Study for Childhood Psychiatric Disorders (BHRC)[115] is a large community school-based study that is following 2,511 children from Brazil since 2010. Psychological, genetic, and neuroimaging data were obtained with the aim to investigate typical and atypical trajectories of psychopathology and cognition over development. Investigators first assessed childhood symptoms and family history of psychiatric disorders in a screening interview, collecting information from 9,937 index children at 57 schools in the cities of São Paulo and Porto Alegre, as well as from 45,394 family members. In the second stage, a random subsample (intended to be representative of the community, N=957) and a high-risk subsample (children at increased risk for mental disorders, based on family risk and childhood symptoms, N=1,554) were selected for further evaluation. In addition, 750 children were invited to take part in a neuroimaging study and to provide blood samples for the assessment of peripheral blood biomarkers. Participants were re-scanned in the same scanner for all follow-up assessments. The sample has two completed follow-up waves at 3 and 6 years. Recreational activities were performed for desensitisation on the day of scanning. All three waves were collected in the same scanners for each participant. For the present study raw structural T1-weighted scans were processed with FreeSurfer 6.0.1.

#### BioDep - Biomarkers of Depression

BioDep is an observational, multi-site (KCL, Cambridge & Oxford), case-control study to investigate depression. Depressed cases screened positive for current depressive symptoms on the Structured Clinical Interview for DSM-5 (SCID)[116] and had a total score >13 on the Hamilton Rating Scale for Depression[117]. CN individuals screened negative for past or current depressive disorder on the SCID. Complete data was collected for 143 eligible participants categorised into three groups: CN (N=53), depressed cases with CRP <3mg/L (loCRP MDD, N=55), and depressed cases with CRP >3 mg/L (hiCRP MDD, N=35). All groups were matched for mean age, sex and handedness. For the present study raw structural T1-weighted scans were processed with FreeSurfer 6.0.1.

#### BSNIP - Bipolar & Schizophrenia Consortium for Parsing Intermediate Phenotypes

BSNIP (Bipolar & Schizophrenia Consortium for Parsing Intermediate Phenotypes) is a multi-centre dataset available through NDAR (ID: 2274) comprising individuals with psychosis, BD, their 1st degree relatives as well as non-psychiatric CN individuals. For the present study raw structural T1-weighted scans were processed with FreeSurfer 6.0.1.

#### Calgary

The Calgary Preschool MRI Dataset was obtained from the Developmental Neuroimaging Lab at the University of Calgary[118]. The dataset consists of multi-modal imaging of 244 individuals in early childhood (2–8 years) and was processed using FreeSurfer 6.0.1.

#### CALM - Centre for Attention Learning and Memory

The children with problems of attention, learning and memory (CALM) cohort [119] consists of 800 children with problems in attention, learning and memory, as identified by a health or educational professional, and 200 typically-developing children recruited from the same schools as those with difficulties. A subset of this cohort underwent a voluntary brain scan at the MRC CBU (www.mrc-cbu.cam.ac.uk). Ethical approval was obtained from the Cambridgeshire Research Ethics Committee and participants or their carers gave written informed consent. MRI measures were collected in a one-hour session conducted on the same site as the CALM clinic on a 3T Siemens Prisma with a 32-channel quadrature head coil. Prior to scanning all children were acquainted with the scanning procedure in a mock scanner. Subsequently a high-resolution 3D T1-weighted structural image was acquired using a Magnetisation Prepared Rapid Gradient Echo (MPRAGE). For the present study raw structural scans were processed with FreeSurfer 6.0.1 using the combined T1 and T2 pipeline where both scans were available and T1 only if T2-weighted images were absent.

#### Cam-CAN - Cambridge Centre for Aging and Neuroscience

The present study included 648 participants from the Cambridge Centre for Aging and Neuroscience (Cam-CAN, www.cam-can.org)[120,121]. These participants were cognitively healthy adults (age range, 18–88) recruited from the local community. Ethical approval was obtained from the Cambridgeshire Research Ethics Committee and participants gave written informed consent. The MRI data come from the same MRC CBU scanner and sequences as the CALM sample above. This sample has previously been processed using FreeSurfer 5.3 and quality controlled[122,123].

#### CAM-FT - Cambridge foetal testosterone

CAM-FT is the Cambridge foetal testosterone cohort [124,125] comprised of 68 individuals recruited from a longitudinal study of the effects of FT on cognitive, behavioural, and brain development [126] to undergo neuroimaging. All imaging for the FT cohort took place at the Wolfson Brain Imaging Centre at Addenbrooke's Hospital (Cambridge, UK) on a Tim Trio 3T magnet (Siemens Medical Solutions). For the present study all raw structural T1-weighted scans were processed with FreeSurfer 6.0.1.

#### CCNP-devCCNP - Chinese Color Nest Project devCCNP

The Chinese Color Nest Project (CCNP) is a a twenty-year project, with a long-term goal to create neurobiologically sound developmental curves for the brain to characterize phenomenological changes associated with the onset of varying forms of mental health and learning disorders, as well as to predict the developmental status (i.e., age-expected values) of an individual brain's structure or function[127]. It consists of three phases (see more details at http://deepneuro.bnu.edu.cn/?p=163): developing CCNP (devCCNP), maturing CCNP (matCCNP) and aging CCNP (ageCCNP). The devCCNP was named as 'Growing Up in China', targeting longitudinal data from 480 typically developing kids. As an initial sample the devCCNP design has been tested at Southwest University, which includes 3 waves of neuroimaging data from 192 developing children (6–18 years) across five years (2013–2017)[128]. More details of these samples can be found in Dong et al.,[129,130]. In the present study all time-points from this cohort are included and all raw structural T1-weighted scans were processed with FreeSurfer 6.0.1.

#### Cuban Human Brain Mapping Project (CHBMP)

The Cuban Human Brain Mapping Project (CHBMP) repository is an open multimodal neuroimaging and cognitive dataset from 282 young and middle age healthy participants (31.9 ± 9.3 years, age range 18–68 years)[131]. This dataset was acquired from 2004 to 2008 as a subset of a larger stratified random sample of 2,019 participants from La Lisa municipality in La Habana, Cuba. Magnetic resonance imaging (MRI) was performed on a 1.5T scanner (MAGNETOM Symphony Siemens Erlangen Germany). All data was subsequently processed using FreeSurfer 6.0.1.

#### CHILD - Cambridge Human Imaging and Longitudinal Development

Cambridge Human Imaging and Longitudinal Development (CHILD) Study will follow the development of babies who have an autistic mother or an autistic older sibling from pregnancy until 36 months old in order to better understand early brain and behaviour differences in babies who have an increased genetic likelihood of being autistic. The current cohort includes one pre- (~33 pcw) and one post-natal (~52 pcw) 1.5T T1-weighted MRI scan. Scans were completed using a GE Optima MR450w 1.5T scanner at the Evelyn Perinatal Imaging Centre, Addenbrookes. Total scan time for each participant was approximately 30 minutes. At the postnatal stage, scans were completed during natural sleep, without sedation. Preprocessing was not done using standard FreeSurfer pipelines; instead the following procedure was followed. Firstly, the orientation of the foetal scans were determined. This was achieved through manually drawing an oil capsule next to left temporal lobe using an image viewer which does not employ orientation codes (MRIcro) and then, using an image viewer which does employ orientation codes (itksnap), manually resetting the orientation code so that the oil capsule appears on the left. The image origin was then reset to the ACPC fibre bundle. To perform skull stripping brain ROIs were manually drawn and multiplied with original head images. A study specific template was then created from a random sample of participants with corresponding foetal and infant scans using the 'buildtemplateparallel.sh' script from the Advanced Normalisation Toolkit. To move STA31 anatomical ROIs[132] into participant space the STA31 template was co-registered to the study specific template, the study specific template itself being co-registered to each participant. After concatenation of the resulting transformations and re-slicing of ROIs, the number of voxels under each warped STA31 ROI was counted and multiplied by appropriate millimeter dimensions resulting in a volume estimate.

COBRE - Center for Biomedical Research Excellence

The Cobre dataset[133] was downloaded from the SchizConnect database (http://schizconnect.org), where it had been obtained from the COllaborative Informatics and Neuroimaging Suite Data Exchange tool (COINS; http://coins.mrn.org/dx). In this dataset, a diagnosis of schizophrenia was made using the Structured Clinical Interview for DSM Disorders (SCID; Diagnostic and Statistical Manual of Mental Disorders, DSM-IV). Exclusion criteria included confirmed or suspected pregnancy, any history of neurological disorders and a history of mental retardation. The data was acquired using a 3T Siemens scanner. For more details on the included sample and processing see[134]. For the present study all structural T1-weighted scans were processed with FreeSurfer 6.0.1.

CONTE

Participants were part of the University of North Carolina (UNC) Early Brain Development Study, approved by the institutional review boards of UNC and Duke University [135]. Mothers of singletons and twins were recruited at the prenatal diagnostic clinics of UNC Hospitals and Duke University Medical Center, as well as by local advertising. Informed consent was obtained on enrollment from each mother and from a parent at each postnatal imaging visit. Exclusion at enrollment included major maternal medical or psychiatric illness, substance use during pregnancy, or abnormalities on prenatal ultrasound. All magnetic resonance images were acquired at UNC using either a Siemens Allegra head-only 3T scanner or a Siemens TIM Trio 3T scanner, which replaced the Allegra in 2011 (Siemens Medical System, Inc., Erlangen, Germany). Infants were scanned during natural sleep after being fitted with earplugs and secured using a vacuum-fixed immobilisation device after birth and at ages 1 and 2 years. At 4 and 6 years, children were scanned awake watching a movie after being trained in a mock scanner. T1- and T2-weighted images were rated for motion artefacts on a scale of 1 to 4; images with a rating of 4 were excluded if artefacts were present in more than a few slices. Neonatal global tissue volumes were determined using an atlas-based expectation-maximisation segmentation algorithm based on both T1- and T2-weighted images specifically adapted to the neonate brain[136]. Tissues were automatically segmented into GM, WM, and CSF; and cortical tissue volumes were derived from a 28-region parcellation of the cerebrum achieved by nonlinear warping of a parcellation atlas template as previously described[137]. Data were processed using CIVET, see [135] for full processing details.

#### Cornell

The Cornell dataset is based on a multi-site study of MDD that has been previously described in detail[138]. All psychiatric diagnoses were based on structured clinical interviews (MINI or SCID) conducted by a trained clinician. High-resolution T1-weighted anatomical scans (MP–RAGE or SPGR) were obtained with specific scanning parameters variable by site. Most sites used a TR of ~2 s, in-plane resolution of ~3.5 mm, and obtained 150–180 volumes in ~5–6 min. Detailed scanning parameters for each site have previously been reported[138].

#### CTAAC - Cape Town Adolescent Antiretroviral Cohort

The purpose of the Cape Town Adolescent Antiretroviral Cohort (CTAAC) is to investigate chronic disease processes in perinatally HIV-infected South African adolescents. The focus is on four key domains: the impact of chronic HIV infection on development; the neuropsychiatric manifestations of HIV in adolescence; the development of chronic lung disease; and early markers of cardiovascular dysfunction. Throughout, the emphasis of this research is on understanding the interactions between chronic disease processes across organ systems. The proposal will enroll 520 perinatally-infected children ages 9-14 years established on antiretroviral therapy. We will follow these children with regular measures (including measures of physical and psychological development, clinical well-being, lung function, cardiovascular status, and emergent risk behaviours) at 6-monthly intervals over 36 months. A CN group of 80 HIV-negative controls will be matched on age, gender and socioeconomic status, in order to collect normative data on key parameters.

#### cVEDA - Consortium on Vulnerability to Externalizing Disorders and Addictions

The full characteristics of the cVEDA cohort are described elsewhere[139]: In short cVEDA is a longitudinal planned cohort study of 10,000 individuals between 6 and 23 years of age, of all genders, representing five geographically, ethnically, and socio-culturally distinct regions in India. Structural (T1, T2, DTI) and functional (resting state fMRI) MRI brain scans have been performed on approximately 15% of the individuals. In addition, 250 T1w scans from individuals aged between 6 and 60y were obtained from the India Brain Template project[140]. For the present study raw structural T1-weighted scans were processed with FreeSurfer 6.0.1.

#### dHCP - Developing Human Connectome Project

The Developing Human Connectome Project (dHCP) is an Open Science project funded by the European Research Council to provide a large dataset of functional and structural brain images from 20 to 44 weeks of gestational age (GA). Neuroimaging was acquired in a single scan session for each infant at the Evelina Newborn Imaging Centre, Evelina London Children's Hospital, using a 3T Philips Achieva system (Philips Medical Systems, Best, The Netherlands). All infants were scanned without sedation in a scanner environment optimized for safe and comfortable neonatal imaging, including a dedicated transport system, positioning device and a customized 32-channel receive coil, with a custom-made acoustic hood [87]. MR-compatible ear putty and earmuffs were used to provide additional acoustic noise attenuation. Infants were fed, swaddled and comfortably positioned in a vacuum jacket prior to scanning to promote natural sleep. All scans were supervised by a neonatal nurse and/or paediatrician who monitored heart rate, oxygen saturation and temperature throughout the scan. Using data from the 2nd DHCP release, tissue segmentation and cortical surface extraction was performed using the MCRIBS pipeline for neonatal cortical parcellation[141] utilizing DHCP tools DrawEM[142] and Deformable[143], respectively. Cortical surfaces were parcellated using the following sequence of steps: inflation and spherical mapping[144], registration to a template[144] and Bayesian labelling[145]. Structural volumes were obtained from segmented tissue images corrected by the location of the surfaces.

#### DCHS - Drakenstein Child Health Study

The Drakenstein Child Health Study as included in the present study consists of two related cohorts:

##### DCHS Infants

This is a nested sub-study that included infants enrolled in a larger population-based birth cohort study, the Drakenstein Child Health Study (DCHS). This DCHS is located in South Africa, in a low to middle-income community of approximately 200,000 people in which there is limited migration. Mothers were recruited at 20–24 weeks gestation, written informed consent obtained, and background data collected for the DCHS. In this nested sub-study, data from two to four week old infants underwent brain magnetic resonance imaging. They were wrapped, fed and then imaged in quiet, natural (unsedated) sleep. Earplugs and mini-muffs were used for double ear protection; a pulse oximeter was used to monitor pulse and oxygenation, and a qualified neonatal nurse or pediatrician was present with the infant in the scanner room for the duration of the imaging session. Multimodal neuroimaging assessment was done at the Cape Universities Body Imaging Centre (CUBIC). The imaging modalities performed included: (1) structural MRI with T1-weighting and T2-weighting to examine cortical and subcortical volumes; (2) diffusion tensor imaging for white matter microstructure; magnetic resonance spectroscopy; and (4) resting state functional MRI for regional connectivity[146].

##### DCHS Mothers

In addition to the recruitment of neonates as described above the DCHS cohort also included mothers. In brief, participants were recruited from two primary health care clinics (TC Newman clinic and Mbekweni clinic) in the Paarl area of the Western Cape[147]. Inclusion criteria for the DCHS included: women over the age of 18 years, who were between 20 and 28 weeks pregnant, who presented to one of two health care clinics for antenatal care (TC Newman and Mbekweni clinics), and had no intention of moving out of the area within the following year, and were able to give written consent[148]. Imaging for this cohort took place at the Cape University Body Imaging Centre (CUBIC) at Groote Schuur Hospital at the University of Cape Town (3T Magnetom Skyra (Siemens).

#### DLBS - Dallas Lifespan Brain Study

The Dallas Lifespan Brain Study (DLBS) is an open imaging dataset aiming to provide understanding of the antecedents of preservation and decline of cognitive function at different stages of the adult lifespan, with a particular interest in the early stages of a healthy brain's march towards AD. It includes 350 healthy adults, aged 20-89 thoroughly characterized in terms of cognition, brain structure and brain function across the adult lifespan. Participants received a structural MRI with DTI, three task-based functional MRI scans, and a resting state scan on a Philips 3T scanner. All participants were scanned on a single 3T Philips Achieva scanner equipped with an 8-channel head coil. For the present study raw structural T1-weighted scans were processed with FreeSurfer 6.0.1.

#### EDSD - European DTI Study on Dementia

The EDSD data include 471 Diffusion Tensor Imaging (DTI) and 471 structural MRI scans (MPRAGE) from patients with AD, MCI, and Healthy Elderly subjects. EDSD is a cross-sectional multicenter study. As of March 2016, the EDSD study sample consists of 139 AD patients, 160 MCI patients and 194 HC. Dementia patients were diagnosed with clinically probable AD according to the NINCDS-ADRCA criteria[149] and were required to be free of any other significant neurological, psychiatric, or medical conditions. Patients with MCI were diagnosed according to the Petersen criteria, exhibiting subjective and objective cognitive impairment (exceeding 1.5 standard deviations in the Consortium to Establish a Registry of Alzheimer's Disease [CERAD] testing battery, controlled for age and education) and being free of dementia[150]. Some MCI patients (N=19) exhibited past or current psychiatric symptoms such as depressive or anxiety symptoms, which were recorded in the clinical data. Cerebrospinal fluid (CSF) information on biomarkers of amyloid and tau pathology is available for 76 MCI subjects, allowing a classification according to the revised diagnostic criteria[151]. CN were required to be free of cognitive complaints and to have performed according to the age and education adjusted norms in all subtests of the CERAD testing battery [152]. For the present study all structural T1-weighted scans were processed with FreeSurfer 6.0.1.

#### EMBARC - Establishing Moderators and Biosignatures Of Antidepressant Response for Clinical Care

The Establishing Moderators and Biosignatures Of Antidepressant Response for Clinical Care (EMBARC) Study is a comparative effectiveness trial of three mechanistically distinct treatments for Major Depressive Disorder (MDD) (citalopram, bupropion, and cognitive behavioural therapy) in which investigators will assess a comprehensive array of carefully selected clinical (i.e., anxious depression, early life trauma) and biological (i.e., genetic, neuroimaging, serum, epigenetic) moderators and mediators of outcome. For the present study raw structural T1-weighted scans were processed with FreeSurfer 6.0.1.

#### Female ASD

The dataset termed 'Female ASD' corresponds to the NDAR dataset 'Multimodal Developmental Neurogenetics of Females with ASD' under ID 2011. It includes a sex-balanced cohort of individuals with and without ASD scanned at George Washington University. For the present study raw structural T1-weighted scans were processed with FreeSurfer 6.0.1.

#### FinnBrain

Participants were mother–infant dyads recruited from the FinnBrain Birth Cohort Study (www.finnbrain.fi)[153], which is a population-based cohort that included participants from Southwestern Finland (Turku region and Åland islands). The studies were conducted according to the Declaration of Helsinki and were reviewed and approved by the Ethics Committee of the Hospital District of Southwest Finland (ETMK:31/180/2011). Infant T1-weighted data were successfully processed with Infant FreeSurfer (N=106) (for total cortical grey matter and white matter), using majority vote segmentation for the subcortex [154]. Toddler data were obtained with a silent T1-weighted PETRA sequence (N=11), and we averaged two images before processing with FreeSurfer 6.0. For T1-weighted (MPRAGE) images from 4-year-olds (N=31) and 5-year-olds (N=121) we obtained GMV, WMV and CSF measures with FreeSurfer 6.0 and used FSL FIRST (FSL v6.0) for segmenting subcortical volumes (based on validation against manual segmentations of the subcortical structures[155]).

#### Frankfurt

The Frankfurt dataset is an openly available multi-modal imaging dataset linked to Genc et al. [156] made available through OSF. It includes a T1-weighted high-resolution anatomical image data preprocessed with FreeSurfer 5.3 included in the public release.

#### GOBS - Genetics of Brain Structure and Function study

Since 2006, the Genetics of Brain Structure and Function study (GOBS), has recruited randomly ascertained extended pedigrees of Mexican American descent living in San Antonio (McKay et al. 2014). High-resolution structural MRI scans were acquired on a Siemens 3T TIM Trio at UTHSCSA. The sample included 1,443 individuals (836 female) with mean age of 40.7 years (SD=15.5, range=18–85). Only a subset of individuals with minimal relatedness (1/8th) were used for the normative models in this study.

#### GOSICH - Great Ormond Street Institute for Child Health

GOSICH is a neurotypical control cohort from within the larger MELD project[1] scanned at Great Ormond Street Institute for Child Health. All participants were scanned on a 3T whole body MRI system (Magnetom Prisma, Siemens Medical Systems), using a 20-channel receive head coil and

[1] https://www.protocols.io/view/meld-protocol-1-patient-and-control-inclusion-in-t-ne2dbge

body coil for transmission and 80mT/m magnetic field gradients. For the present study raw structural T1-weighted scans were processed with FreeSurfer 5.3.

#### GUSTO - Growing Up in Singapore Towards Healthy Outcomes

The Growing Up in Singapore Towards Healthy Outcomes (GUSTO)[157] cohort consists of pregnant Asian women attending the first trimester antenatal ultrasound scan clinic at the National University Hospital and KK Women's and Children's Hospital in Singapore. The parents were Singapore citizens or permanent residents of Chinese, Malay or Indian ethnic background. Birth outcomes and pregnancy measures were obtained from hospital records. Socioeconomic status (household income) was extracted from survey questionnaires conducted as a part of a scheduled appointment during pregnancy. The GUSTO study was approved by the National Healthcare Group Domain Specific Review Board and the Sing Health Centralized Institutional Review Board, and all participating mothers provided informed consent. Children were scanned at different ages throughout their development. The present study includes longitudinal data (3 time points) from children between the age of 4 and 8 (with scanning completed around years 4, 6 and 8). For the present study, structural T1-weighted MPRAGE scans were processed with FreeSurfer 6.0.1.

#### HABS - Harvard Aging Brain Study

The Harvard Aging Brain Study (HABS NIH-P01AG036694)[158] is a longitudinal observational study designed to further our understanding of differentiating 'normal' aging from preclinical AD. Longitudinal data collection in HABS is ongoing and now in its eleventh year. Inclusion criteria included: 50 years of age or older (minimum age for the lifespan MRI project at enrollment was 62); a score of 0 on the Clinical Dementia Rating Scale; a score of greater than 25 on the Mini-Mental State Examination; scores above age and education-adjusted cutoffs on the 30-Minute Delayed Recall of the Logical Memory Story A[159] (ADNI based cut-offs, http://www.adni-info.org/); and a score of less than 11 on the Geriatric Depression Scale. Exclusion criteria included: history of alcoholism, drug abuse, head trauma, or current serious medical/psychiatric illness. The criteria utilized for inclusion/exclusion ensures the HABS cohort consists of a diverse group of cognitively normal, healthy older individuals at study enrollment. As of March 2020, 15% of the initial cohort had progressed to MCI, and we expect additional individuals to progress over the next five years. For the present study raw structural T1-weighted scans were processed with FreeSurfer 6.0.1.

#### Harvard foetal

Harvard foetal cohort of in-utero foetal MRIs combines (i) foetal MRIs of pregnant mothers that were prospectively recruited as control subjects for previous research projects and (ii) clinical foetal MRIs that were performed to screen for foetal brain abnormalities but were clinically interpreted as normal by two board-certified radiologists, experienced in foetal MRI. Inclusion criteria for enrollment were as follows: no serious maternal medical conditions during pregnancy, between 15 and 37 GWs; fetuses recruited prospectively as controls in other research studies; and fetuses with MRI that were read as normal by radiologists and pediatric neuroradiologists. Exclusion criteria were as follows: multiple gestation pregnancies, dysmorphic features on US examination, brain malformations or brain lesions identified on MRI or US, other identified organ anomalies, known chromosomal abnormalities, and known congenital infections. The gestational age of fetuses was estimated based on the available clinical data (crown-rump length and/or the first day of the last menstrual period) as recommended by the American College of Obstetricians and Gynecologists and Committee on Obstetric Practice, 2017. For the present study, the raw structural T2-weighted scans were processed using in-house-built pipelines that involved inter-slice motion correction for super-resolution volume reconstruction followed by brain extraction and tissue segmentation[160–162].

#### HBN - Healthy Brain Network

The Healthy Brain Network (HBN)[163], is an ongoing initiative focused on creating and sharing a biobank of data from 10,000 New York area participants (ages 5–21) organised by the Child Mind Institute. The HBN Biobank houses data about psychiatric, behavioural, cognitive, and lifestyle phenotypes, as well as multimodal brain imaging (resting and naturalistic viewing fMRI, diffusion MRI, morphometric MRI), electroencephalography, eye-tracking, voice and video recordings, genetics and actigraphy. In the present project pre-processed FreeSurfer output was used alongside the primary diagnostic labels from all data available up to and including release 7.

#### Human Connectome Project

Publically available data from the human connectome project (HCP; http://www.humanconnectome.org/), comprised MRI data from 1,113 individuals (606 female) from 457 unique families (including 170 dizygotic twins, 286 monozygotic twins, 576 non-twin siblings, and 25 non-sibling familial relations) with mean age 28.8 years (SD=3.7, range=22–37). As previously described in detail[83,164], T1-weighted and T2-weighted structural images were acquired on a 3T Siemens Skyra employing a 32-channel head coil.

#### Human Connectome Project Aging and Development

The Human Connectome Project Aging[165] and Development studies, HCPa and HCPd respectively are extensions of the aforementioned HCP study, specifically aimed at extending the coverage of HCP to a fuller lifespan. They follow a comparable acquisition and recruitment protocol to HCP[166], albeit with some key adaptations aimed to tackle the challenges of scanning younger and older populations compared to HCP. Key differences include a slightly larger T1w voxel size (0.8mm) to allows some additional SNR margins, use of volumetric navigators for prospective motion correction, only one acquisition per modality to reduce scanning time, multi-echo acquisition for T1 (TE=1.8, 3.6, 5.4 and 7.2ms), slower TR (800ms) to allows maintenance of full Fourier k-space acquisition necessitated by the increased number of echoes. For the present project, minimally processed data was obtained directly from HCP and was run through FreeSurfer 6.0.1.

#### iADNI - Italian Alzheimer's Disease Neuroimaging Initiative

I-ADNI is a cross sectional study and consists of 262 patients with subjective memory impairment, mild cognitive impairment, AD dementia and frontotemporal dementia enrolled in 7 Italian centers. Few cognitively healthy elderly CN individuals were also included. This study has provided standardisation of MRI acquisition and imaging marker collection across different Italian clinical units and equipment. This is a mandatory step to the implementation of imaging biomarkers in clinical routine for early and differential diagnosis. MRI site qualification and MP-RAGE quality assessment was applied following the ADNI-1 procedures. For the present study all structural T1-weighted scans were processed with FreeSurfer 6.0.1.

#### ICBM

The International Consortium for Brain Mapping aims to collect comprehensive neuroimaging and genetics data from 7000 individuals in an effort to generate a probabilistic human brain atlas[167,168]. Current data are available through the University of Southern California's Laboratory of Neuroimaging (LONI; http://loni.usc.edu/about_loni). For the present project all T1-weighted anatomical images were downloaded from the LONI repository and processed using FreeSurfer 6.0.1.

#### IMAGEN

IMAGEN is a European research project examining how biological, psychological, and environmental factors during adolescence may influence brain development and mental health[169]. It includes longitudinal data of up to 3 time-points for an early adolescent sample from across Europe. All T1-weighted structural data was processed using the standard recon-all pipeline included in FreeSurfer 6.0.1.

#### IMAP - Multi-modal Neuroimaging in Alzheimer's Disease

Two hundred and fifteen cognitively unimpaired individuals from the IMAP study (multimodal neuroimaging of early Alzheimer's disease; Caen, France; PIs: Gaël Chételat (scientific) & Vincent de La Sayette (MD); sponsor: Caen University Hospital) [170], aged between 19 and 85 (88 participants < 40 years old, 56 between 40 and 60 years old, and 71 participants > 60 years old), were included in this study. Participants were recruited from the general population through advertisement or word of mouth. They had no history or clinical evidence of major neurological or psychiatric disorder and performed in the normal range in all neuropsychological tests (including tests of episodic memory, working memory, language skills, executive functions, and visuospatial abilities). A regional review board has approved the use of human participants for this study, and consent forms from all participants were obtained. All neuropsychological, MRI, and PET assessments were performed in close temporal proximity (within 3 months). MRI scans were acquired at the Cyceron Center (Caen, France) on a Philips (Eindhoven, The Netherlands) Achieva 3T scanner using a 3D fast-field echo sequence. All T1-weighted structural data was processed using the standard recon-all pipeline included in FreeSurfer 6.0.1.

#### IXI

The IXI data set consists of a variety of MR images[171] from nearly 600 normal, healthy subjects with their respective demographic information that are freely available for download (https://brain-development.org/ixi-dataset/). Only the T1-weighted images were used in the present project. MRI data were acquired in three different scanners, two of which were 1.5T and one was 3T. The 3T scanner was a Philips Intera and the T1-weighted acquisition consisted of a standard T1 with the following parameters TR=9.6, Echo Time=4.6, FoV=208 x 208, Flip Angle=8.0. The 1.5T scanner was a Philips Gyroscan Intera using a T1 weighted (TR=9.6, Echo Time=4.6, Flip Angle=8.0) acquisition. All scans were processed with FreeSurfer 6.0.1.

#### KNE96 - Korean normal elderly brain template study

The KNE96 dataset includes 96 (M/F=48/48) right-handed, cognitively normal (CDR=0) Koreans aged 60 year or older (M=69.5±6.2 years, F=70.1±7.0 years) selected from the participants of the Korean Longitudinal Study on Cognitive Aging and Dementia (KLO-SCAD). Three-dimensional (3D) .

#### LA5c - UCLA Consortium for Neuropsychiatric Phenomics LA5c Study

The LA5c dataset comprises data on OpenNeuro from the UCLA Consortium for Neuropsychiatric Phenomics LA5c Study [172] which includes 272 subjects with and without psychiatric diagnoses. For the present study all structural T1-weighted scans were processed with FreeSurfer 6.0.1.

#### LATAM

The cohort referred to as LATAM in the present dataset was collected as part of the Latin American Network for the Study of Early Psychosis (ANDES; www.cyted.or/ede/NDES), a consortium of research groups from six Latin American countries: Argentina, Bolivia, Brazil, Chile, Colombia and Mexico[173]. The data was specifically collected at the Pontificia Universidad Católica de Chile and included a T1 weighted acquisition from a Philips Ingenia 3T MRI scanner with a 16-channel coil[174].

#### LIFE - Leipzig Research Centre for Civilization Diseases Study

The Leipzig Research Centre for Civilization Diseases Study (LIFE Study) is a population-based study from Leipzig, Germany, with the objective to investigate the development of major modern diseases[175]. Overall, 10,000 participants were randomly drawn from the local population, of whom 2,667 underwent MRI and detailed screening. With age-associated diseases such as mild and major neurocognitive disorder being one of the main focuses of this study, most participants were adults older than 60 years of age.

#### MCIC

The Mental Illness and Neuroscience Discovery Institute (MIND) Institute, now the Mind Research Network (MRN, www.mrn.org) formed the MIND Clinical Imaging Consortium (MCIC) in 2003 to conduct a multi-institutional, cross-sectional study of patients with schizophrenia and demographically matched, by sex and age, healthy controls to identify quantitative neuroimaging biomarkers for this devastating disease[176]. Standardisation of acquisition across sites was previously evaluated in a separate calibration and validation study[177]. Structural T1 weighted imaging data from this consortium was made available through the Collaborative Informatics Neuroimaging Suite (COINS). For the present study all data was processed with FreeSurfer 6.0.1.

#### MCSA (Mayo Clinic Olmsted Study of Aging)

The objectives of the Mayo Clinic Study of Aging were to determine in the population of Olmsted County, Minn., (1) the prevalence of MCI; (2) the incidence of MCI; (3) conversion rates from MCI to dementia or AD; (4) risk factors for MCI; and (5) risk factors for the progression from MCI to dementia or AD[178]. Details on the recruitment procedure and study design can be found in Roberts et al.[178] For the current project the first time-point scans of the first 1,000 subjects were shared. All images were processed using FreeSurfer software version 6.0.1.

#### MRi-Share

The i-Share (for internet-based Student Health Research enterprise; www.i-share.fr) cohort project was conceived to investigate the impact of learning and social changes associated with higher education on maturational changes in the brain, and how it interacts with the personal traits, physical and mental health status to influence immediate as well as later-life events. An important sub-component of the i-Share study, which was called 'MRi-Share', is a multi-modal brain magnetic resonance imaging (MRI) database collected in a subset of i-Share participants [179]. The specific motivations behind MRi-Share were to 1) characterize late-maturational changes of post-adolescence brain; 2) investigate the impact of higher education on late maturational processes of the brain; 3) study the associations between brain phenotypes and neuropsychiatric conditions prevalent in young adults, such as migraine, depression and anxiety disorders, and substance abuse; and 4) establish the early occurrence of imaging biomarkers of late-life disorders, such as white matter hyperintensities (WMH) and enlarged perivascular space (ePVS)'. The MRI acquisition protocol for the MRi-Share database was designed to closely emulate that of the UKB MR brain imaging study, in terms of both modalities and scanning parameters for each. All neuroimaging data were acquired on the same Siemens 3T Prisma scanner with a 64-channels head coil (gradients: 80 mT/m - 200 T/m/s), in the 2-year period between November 2015 and November 2017. All T1-weighted images were processed using FreeSurfer software version 6.0.1.

#### Narratives

The "Narratives" collection[180] aggregates auditory story-listening fMRI datasets acquired over the course of roughly seven years (2011–2018). Stimuli comprised 28 naturalistic spoken stories ranging from ~3 to ~56 minutes for a total of ~5 hours of unique audio stimuli. The collection includes 345 unique subjects participating in over 750 functional scans with accompanying anatomical data. For the present study all structural T1-weighted scans were processed with FreeSurfer 6.0.1.

#### NeuroScience and Psychiatry Network

The Neuroscience and Psychiatry Network (NSPN) study comprises a primary cohort of 2402 healthy young people, recruited from schools, colleges, NHS primary care services and direct advertisements in north London and Cambridgeshire. Participants were stratified into five age groups (14–15, 16–17, 18–19, 20–21 and 22–25 years) and each stratum was evenly balanced for sex and ethnicity. Primary participants completed demographic, medical, childhood trauma and mental health questionnaires by post. The secondary cohort sub-sampled approximately 60 individuals from each stratum in the primary cohort, maintaining the sex and ethnicity balance. Secondary participants completed MRI scanning as part of a whole-day assessment at one of two sites (Cambridge and London, UK), on at least two occasions. Cohort retention for the MRI follow-up was 74%[181]. The present study included 295 individuals for which baseline T1 images were available. These data were preprocessed using FreeSurfer 5.3 and manually quality controlled as detailed previously[182,183].

#### NHGRI

The NHGRI dataset refers to NDAR project 2936 and consists of imaging data shared by the NIMHP-IRP Data Science and Sharing Team. All T1-weighted images were processed using FreeSurfer 6.0.1.

#### NIH

The National Institute of Health (NIH) sample [44] was part of a study of normal brain development where participants submitted a blood sample, underwent a comprehensive neuropsychological evaluation, and a T1-weighted structural brain scan every two years over a period of up to 12 years. All T1-weighted images were processed using FreeSurfer 6.0.1.

#### NIHPD Infant and Adult

The NIH study of pediatric development (NIHPD) is a multisite, combined cross-sectional and longitudinal study of normal, healthy developing children (representative of US Census 2000 statistics for gender, family income, race/ethnicity) from early childhood through young adulthood.

A complete list of the sites and procedures can be found at: https://www.nitrc.org/docman/view.php/98/288/MRI_Manual_Nov06.pdf.

#### NKI - Nathan Kline Institute Rockland Sample

The Nathan Kline Institute (NKI) Rockland Sample is an ongoing initiative to generate a deeply phenotype and community ascertained lifespan sample with advanced neuroimaging and genetics[184]. For the present study we downloaded multi-modal imaging data from the 100-functional connectomes project for 532 quality controlled T1 images and these were processed using FreeSurfer 5.3.

#### NTB_Yale

The Turk-Browne lab dataset (NTB_Yale) comprises anatomical scans from the initial three cohorts of data collection in an ongoing awake infant fMRI project. Participants were scanned at one of three sites on a 3T MRI: the Scully Center for the Neuroscience of Mind and Behavior at Princeton University (Siemens Skyra), the Magnetic Resonance Research Center (MRRC) at Yale University (Siemens Prisma), and the Brain Imaging Center at Yale University (Siemens Prisma). Participants in the Princeton cohort were recruited through flyers and word-of-mouth, and in the Yale cohorts from maternity ward visits at the Yale-New Haven Hospital. Informed consent was obtained from a parent or guardian according to a protocol approved by the institutional IRB. The sample included healthy participants between 3 and 36 months of age. Most of the scans were collected while the infant was watching a movie, though on rare occasions they had fallen asleep. See reference[185] for further details about data acquisition for the first two cohorts.

#### OASIS3 - Open Access Series of Imaging Studies

The Open Access Series of Imaging Studies (OASIS) is a multimodal collection of data focused on the effects of healthy aging and AD that is freely available to the scientific community. OASIS-3 incorporates data from 1,098 participants covering the adult life span aged 42 to 95, including cognitively normal individuals and individuals with early-stage AD dementia. OASIS-3 includes participants enrolled into several ongoing studies through the Charles F. and Joanne Knight Alzheimer Disease Research Center (Knight ADRC) at Washington University in St. Louis spanning over 15 years and several research studies - Memory and Aging Project, Adult Children Study[186], and Healthy Aging and Senile Dementia. Each study targets varying cohorts but includes similar assessments and visit intervals. The following cohorts were recruited: (1) Individuals who were generally healthy, were cognitively normal (CDR=0), and had a family history of AD, defined as being a biologic child of at least one parent with a reported history of AD dementia with onset age 80 years; (2) individuals who were generally healthy, were cognitively normal (CDR=0), and had no family history of AD for either biological parent and lived at least to age 70 years; and (3) healthy individuals 65 and older, both those who were cognitively normal (CDR 0) and this with very mild-mild symptomatic AD (CDR 0.5 and 1). Exclusion criteria included medical conditions that precluded longitudinal participation (e.g., end-stage renal disease requiring dialysis) or medical contraindications for the study arms (e.g., pacemaker for MRI, anticoagulant use for lumbar puncture). Participants were recruited from the community via flyers, word of mouth, and community engagements. Participants from all cohorts agreed to submit an initial blood sample for genetic testing, complete regular cognitive testing, and neuroimaging and lumbar punctures approximately every 2-3 years. Each participant was enrolled along with a collateral source, someone who knew the participants well (e.g., spouse or adult child) and could report whether the participant's current cognitive and functional performance was or was not at previously attained levels. Participants enrolled in studies at the Knight ADRC Clinical Core were referred to the Knight ADRC Research Imaging (KARI) Program for magnetic resonance imaging (MRI) and positron emission tomography (PET) scans. All participants were required to have a CDR ≤1 at the time of most recent Clinical Core assessment. Participants completed screening for general health information to assess any contraindications to PET or MR imaging. Participants were excluded for the following health reasons: women who were pregnant or breastfeeding; implanted medical devices such as pacemakers and drug pump; history or risk of metal in the eye; and history of claustrophobia. Eligible participants signed informed consent for one of the KARI imaging studies that included MR only, PET only, or MR and PET scans. To the best effort of investigators, participants underwent scan sessions within six months of Clinical Core visits. Across the years of scanning, gaps in funding, funding for additional sub-studies, or participant related delays, have caused variations in visit timelines resulting in extended or decreased intervals.

OHSU

The Oregon Health & Science University (OHSU) study (PIs: Damien Fair, Joel Nigg) included individuals with ADHD, ASD and typically-developing CN individuals as previously described[187–191]. Participants were recruited via mailings to commercial mailing lists and public advertisements. Families participated in a multi-gated procedure that included an initial phone screen. Parents of children who remained eligible upon completion of the initial phone screen were invited to complete the ADHD Rating Scale, Conner's Rating Scale, 3rd edition, and the Strengths and

Difficulties Questionnaire, and an in-person semi-structured diagnostic interview (Kiddie Schedule for Affective Disorders and Schizophrenia, KSADS) while the child completed IQ screening and brief academic achievement testing. A best estimate DSM-IV diagnosis was established by a multidisciplinary diagnostic team. Blind to one another's ratings and to the subsequent cognitive test scores, they formed a diagnostic opinion based on all available information. Their agreement rate was excellent (ADHD diagnosis kappa =.88). Disagreements were conferenced and consensus reached. Cases where consensus was not readily achieved were excluded. ASD youth were recruited in a more targeted fashion. Children who met criteria for DSM-IV diagnosis of either Autistic Disorder, Asperger's Disorder, or Pervasive Developmental Disorder Not Otherwise Specified were recruited through the University's Autism Clinic in the Child Development and Rehabilitation Center, community support groups and outreach, and targeted mailings based on charted ICD-9 codes. After passing initial screening criteria, participants in the ASD group were administered the Autism Diagnostic Observation Schedule and parents completed the Autism Diagnostic Interview–Revised. Parents also completed the Social Responsiveness Scale, 2$^{nd}$ edition (SRS-2), Children's Communication Checklist, and a detailed developmental history questionnaire. Parents had the option to provide any existing documentation regarding a previous ASD diagnosis including IEP's, chart records, and psychological evaluations. A multi-disciplinary diagnostic team that included three licensed clinicians then utilized all of the aforementioned materials to determine a consensus diagnosis. Children in all three diagnostic groups were excluded if they: were prescribed long-acting psychotropic medications; had neurological impairment, seizure history, head injury with loss of consciousness, other major medical conditions, or substance abuse; had a prior diagnosis of intellectual disability, or psychosis; were currently experiencing a major depressive episode; or had estimated IQ <70. Other comorbidities were assessed by the multi-disciplinary teams, but were not exclusionary in any of the diagnostic groups except that children in the ADHD and Control samples were excluded if they had a parent-reported history of ASD diagnosis or if the multidisciplinary team identified a diagnosis of ASD. Children with ASD or ADHD taking stimulant medications were included in the study but were required to be off medication for 24 (for short-acting preparations) to 48 hours (for long-acting preparations) prior to testing or MRI.

#### OpenPain

The OpenPain (https://www.openpain.org) project (PI : A. Vania Apkarian, Northwestern University) is supported by the National Institute of Neurological Disorders and Stroke (NINDS) and National Institute of Drug Abuse (NIDA). It is an aggregation of several subcohorts described below. For the present study all structural T1-weighted scans from these cohorts were processed with FreeSurfer 6.0.1.

##### Placebo 1 (PL1)

This study was conducted in the setting of a clinical randomized controlled trial specifically designed for assessing the placebo response (registered at https://www.clinicaltrials.gov/ct2/show/NCT02013427). The study consisted of 6 visits spread over ~8 weeks, including a baseline monitoring/screening period and two treatment periods, each followed by a washout period. The overall protocol included four scanning sessions collected before and after each treatment period.

##### Subacute longitudinal study (SA1 & SA2):

Subjects with SBP were recruited who reported a single intense episode of back pain lasting 4-16 weeks and no prior back pain for at least 1 year, performed brain scans as soon as possible (mean ± SEM pain duration from injury at visit 1: 9.14 ± 0.48 weeks) and followed their pain and mood parameters, as well as brain activity, over three additional visits for the next year (visit 2: 7.15 ± 0.26 weeks; visit 3: 29.20 ± 0.63 weeks; visit 4: 54.36 ± 2.14 weeks; from visit 1).

##### Placebo predict Tetreault (PLOA)

Data includes a discovery group used to identify and localize brain functional differences between placebo responders and nonresponders. Additionally, a validation study is included, which involved a double-blinded trial in which patients received placebo or duloxetine for 3 mo.For all patients, brain scans were collected prior to treatment.

##### Brain network change Mano (RS)

All the scans were performed on a 3T Magnetom Trio with TIM system (Siemens, Erlangen, Germany) equipped with echo planar imaging (EPI) capability and a standard 12-channel phased array head coil either at Addenbrooke's hospital (Cambridge, UK) or CiNet (Osaka, Japan). Participants remained supine and wore MR-compatible headphones with their heads immobilised with cushioned supports during scanning.

##### Accumbens Chronic Pain Signature (SAB)

Subacute back pain patients were followed longitudinally. The Data were collected at Yale University. The study recruited 40 SBP patients (16 females, average age ± SEM: 31.7 ± 1.7 y), 28 CLBP patients (17 females, 32.2 ± 2.0 y), and 30 healthy controls (14 females, 31.1 ± 2.0 y). CLBP patients were studied at one time point only. The SBP patients and healthy controls were followed up longitudinally for a median duration of ~1 y. Of the 40 SBP patients, 35 (87.5%) presented for follow-up (32.5 ± 1.9 y, 14 females) and completed questionnaires, 26 (65%) consented to scan, and 5 were lost to follow-up. Of the 30 healthy controls, 16 (53.3%) presented for follow-up (age=31.6 ± 2.5 y, 7 females) 14 (46.7%) consented to scan, 11 (36.7) were not yet due for follow-up, and 5 (16.7%) were lost to follow-up. The median duration at follow-up was 59.4 wk. SBP patients were dichotomized into recovered back-pain patients (SBPr, N=19) if their back-pain intensity dropped ≥30% on the VAS relative to the pain at entry into the study or into persistent back-pain patients (SBPp, N=16) otherwise. This study was approved by the Yale University Institutional Review Board. All participants gave informed consent for inclusion in our study.

#### Oslo

The sample was drawn from a longitudinal research project run by Center for Lifespan Changes in Brain and Cognition and was an extension of a previously published protocol[192]. The study was approved by the Regional Ethical Committee of South Norway (REK-Sør), and written informed consent was obtained from all participants prior to the examinations. Volunteers were primarily recruited by advertisements in newspapers and social media. Participants were required to be right-handed native Norwegian speakers, feel healthy, not use medicines known to affect central nervous system (CNS) functioning, including psychoactive drugs, not be under psychiatric treatment, be free from worries regarding their memory abilities, and not have injury or diseases known to affect CNS function, including neurological or psychiatric illness, serious head injury, or history of stroke. All MR scans were subjected to a radiological evaluation by a specialist in neuroradiology, and the participants were required to be deemed free of significant injuries or conditions. Data were processed using FreeSurfer 6.0.1.

#### Oulu

The Oulu dataset refers to a dataset shared as part of the International Neuroimaging Datasharing Initiative (INDI)[193] and includes 103 subjects (37M, 66F) between the ages of 20 and 23. All structural T1-weighted images from this dataset were processed with FreeSurfer 6.0.1 to obtain tissue segmented volumes.

#### Penn-CHOP Developmental Connectome (PCDC)

PCDC (www.brainmrimap.org) is a cohort on typical brain development including infancy (0-2years) at Children's Hospital of Philadelphia (CHOP) for establishing next-generation developmental brain MRI atlases (quantitative UPenn-CHOP brain atlases). Infant MRI data of PCDC was used. All participants were scanned on a 3T Siemens Prisma scanner, using a 32-channel receive head coil. In addition to a T1 MPRAGE, high-resolution 3D T2-weighted structural images were acquired using Sampling Perfection with Application-optimized Contrasts using different flip angle Evolutions (SPACE) with the following parameters: TR=3200ms, TE=564 ms, 208 slices, sagittal acquisition, FOV=256×240mm, and voxel size=0.8mm isotropic.

#### PING - Pediatric Imaging, Neurocognition, and Genetics

The PING Data Resource is the product of a multi-site project involving developmental researchers across the United States including UC San Diego the University of Hawaii UC Los Angeles Children's Hospital of Los Angeles of the University of Southern California UC Davis Kennedy Krieger Institute of Johns Hopkins University Sackler Institute of Cornell University University of Massachusetts Massachusetts General Hospital at Harvard University and Yale University. The Data Resource includes neurodevelopmental histories, information about developing mental and emotional functions, multimodal brain imaging data, and genotypes for well over 1,000 children and adolescents between the ages of 3 and 20. The current data was obtained from the NIMH Data Archive (NDAR ID: 2607) and all T1 weighted structural imaging was run through FreeSurfer 6.0.1.

#### Pixar

Pixar[194] is an OpenNeuro dataset comprising 155 children who watched Disney Pixar's 'Partly Cloudy' while lying in the scanner. There was no task; participants were simply instructed to lie still and watch the movie. The movie began after 10s of rest (black screen; TRs 1-5). The first 10s of the movie are the opening credits (disney castle, pixar logo; TRs 6-10). For the present study all structural T1-weighted scans were processed with FreeSurfer 6.0.1.

#### PNC - Philadelphia Neuroimaging Cohort

Recruitment and study protocols for the PNC have been described in detail previously[195–197]. Briefly, 9,498 individuals 8 to 21 years old were recruited from the Children's Hospital of Philadelphia care network. Study procedures were approved by the institutional review boards of both the University of Pennsylvania and the Children's Hospital of Philadelphia. All MRI scans were acquired on a single 3T Siemens TIM Trio scanner with 32-channel head coil as described previously[197].

#### POND - Province of Ontario Neurodevelopmental Disorders

T1-weighted images were obtained from the Province of Ontario Neurodevelopmental Disorders (POND) study[2], which was approved by each of the participating research ethics boards and conducted in accordance with its guidelines. Informed written consent was obtained from all participants and/or their parents. Participants were recruited via the Province of Ontario Neurodevelopmental Disorders Network, across five Centers in Ontario, Canada (Holland Bloorview Kids Rehabilitation Hospital, Toronto; The Hospital for Sick Children, Toronto; McMaster Children's Hospital, Hamilton; Queen's University, and Lawson Health Research Institute, London). Controls were recruited through advertising in public transit, in hospitals, and on social media. Inclusion criteria were age <18 years, and a clinical diagnosis of ADHD, ASD, or OCD. Controls had no developmental diagnosis, and no first-degree family history of these disorders[198]. The majority of scans (74%) were done on a 3T Siemens Trio TIM; a hardware upgrade to the Siemens Prisma scanner took place in June, 2015 (this affected 26% of the ADHD sample, 24% of the ASD sample, 62% of controls, and 7% of the OCD sample). All images were processed with FreeSurfer 6.0.1.

#### PPMI - Parkinson's Progression Markers Initiative

The Parkinson's Progression Markers Initiative (PPMI)[199] is an observational study to better define and measure Parkinson's disease to speed therapeutic development. PPMI makes its data set and biorepository—the most robust in Parkinson's to date—available to academia and industry to accelerate breakthroughs. PPMI has gathered longitudinal data from more than 1,400

[2] https://offordcentre.com/studies/pond-network-imaging-sub-study/

individuals at 33 clinical sites in 11 countries. These data are shared through the University of Southern California's Neuroimaging Laboratories database (https://ida.loni.usc.edu/). T1-weighted images were downloaded from the LONI database and processed using FreeSurfer 6.0.1

#### PREVENT-AD - PRe-symptomatic EValuation of Experimental or Novel Treatments for AD

In 2010 investigators at McGill University and the Douglas Mental Health University Institute Research Centre created a Centre for **St**udies **o**n **P**revention of **A**lzheimer's **D**isease (StoP-AD Centre). The Centre's prime objective was to pursue innovative studies of pre-symptomatic AD, with efforts to provide relatively enriched samples for prevention trials requiring individuals at-risk of developing the disease. To this end, the StoP-AD Centre developed an observational cohort for **PR**e-symptomatic **EV**aluation of **E**xperimental or **N**ovel **T**reatments for **AD** (PREVENT-AD)[200]. To increase the probability that participants would harbor the earliest changes associated with pre-symptomatic AD, entry criteria required intact cognition and a parental or multiple-sibling family history of AD. Participants had to be 60 years of age or older, with an exception that individuals between 55–59 years old were eligible if their own age was within 15 years of symptom onset of their youngest-affected first-degree relative. All participants were scanned longitudinally on a Siemens TIM Trio 3T MRI scanner at the Brain Imaging Centre of the Douglas Mental Health University Institute using a Siemens standard 12 or 32-channel coil (Siemens Medical Solutions, Erlangen, Germany) using the same acquisition protocol as ADNI. For the present study all structural T1-weighted scans were processed with FreeSurfer 6.0.1.

#### PSYSCAN Maastricht

The Maastricht GROUP dataset comes from an MRI study in Maastricht, the Netherlands, led by the GROUP consortium. Patients were identified by screening caseloads of representative clinicians for inclusion criteria in selected representative geographic areas of the Netherlands and Belgium. All patients satisfied DSM-IV diagnostic criteria for schizophrenia or other nonaffective psychotic disorders. The data was acquired using a 3T Siemens Magnetom Allegra head scanner. For more information, see [134,201]. For the present study all structural T1-weighted scans were processed with FreeSurfer 6.0.1. Note that this dataset forms part of legacy data collated as part of the PSYSCAN project.

#### PSYSCAN Dublin

The Dublin dataset was acquired and scanned in the Trinity College Institute of Neuroscience as part of a Science Foundation Ireland-funded neuroimaging genetics study—a structural and functional MRI investigation of genetics, cognition and emotion in schizophrenia. Patients were recruited through local clinical services whilst healthy control subjects reported no history of psychiatric disease. All patients satisfied DSM-IV diagnostic criteria for schizophrenia or other nonaffective psychotic disorders. Both groups were recruited in the same geographical area through local advertisement and exclusion criteria for both groups included confirmed or suspected pregnancy, any history of neurological disorders or intellectual disability and substance misuse in the preceding 3 months. The data was acquired using a 3T Philips Intera Achieva scanner. For more information and details on the included sample have been described previously[134]. For the present study all structural T1-weighted scans were processed with FreeSurfer 6.0.1. Note that this dataset forms part of legacy data collated as part of the PSYSCAN project.

#### RDB

This dataset includes 494 subjects from the Robert Debré Hospital (RDB) in Paris. It spans an age range with most subjects being young children and adolescents (age range 1.5–55.2, average 10.83, median 8). It includes both individuals diagnosed with ASD (N=325) and undiagnosed control subjects (N=169). The majority of subjects were male (127 female, 367 male). Accompanying phenotypic information include sex, age at scan and diagnosis of the participants, as well as a standardized evaluation of their cognitive abilities using a variety of tests depending on chronological age, productive language skills and functioning level of subjects. All T1-weighted imaging data were processed using recon-all as implemented in FreeSurfer 6.0.1. The MRI and volume data were visually quality controlled.

#### SALD - Southwest University Adult Lifespan Dataset

The data generated in the Southwest University Adult Lifespan Dataset (SALD) comprises a large cross-sectional sample (N=494; age range=19–80) undergoing a multi-modal (sMRI, rs-fMRI, and behavioural) [202]. The goals of the SALD are to give researchers the opportunity to map the structural and functional changes the human brain undergoes throughout adulthood and to replicate previous findings. The data were collected at the Center for Brain Imaging, Southwest University. For the present study all structural T1-weighted scans were processed with FreeSurfer 6.0.1.

#### SCZIowa

The SCZIowa cohort refers to the Phenomenology and Classification of Schizophrenia (Iowa Longitudinal Study)[67,203] listed in NDAR (2125; description below taken from NDAR). This study follows first-episode patients, some of whom have been followed for as long as 15 years. The study emphasizes understanding the phenomenology of schizophrenia by examining the lifetime trajectory of the illness and its long-term outcome. Since the disorder is characterized by a prolonged lifetime course, longitudinal study of a large group of informative patients is one of the most powerful strategies for examining measures that will illuminate its mechanisms or refine the definition of its phenotype. This study examines 4 domains of variables: Symptoms, psychosocial function, brain morphology as measured by morphometric magnetic resonance imaging, and cognition as measured by both standard neuropsychological tests and a group of experimental tests. We have found that, although symptoms stabilize relatively quickly after initial onset in the majority of patients, the other domains tend to worsen throughout the first decade after onset. This suggests that the disorder may have a worse prognosis, despite adequate treatment, than originally anticipated. We also will divide the patients into groups based on levels of recovery and determine the predictors of recovery group. In order to understand the long-term outcome of schizophrenia at the clinical, neural, and cognitive levels, it is important that we continue to study this large group of patients longitudinally on into the second decade of the illness so that we can examine the interrelationships between these four domains and determine whether the downward trends persist or stabilize. For the present study all structural T1-weighted scans were processed with FreeSurfer 6.0.1.

##### SLIM - Southwest University Longitudinal Imaging Multimodal Dataset

The SLIM dataset includes brain and behavioural data across a long-term retest-duration within three and a half years. MRI scans provided a set of structural, diffusion and resting-state functional MRI images, along with rich samples of behavioural assessments addressed including cognitive and emotional information[204]. A total of 167 healthy undergraduate students from the local community of Southwest University in China participated in this study as a part of our ongoing project investigating the associations among brain imaging, mental health, and creativity. For the present study all structural T1-weighted scans were processed with FreeSurfer 6.0.1.

##### STRIvE - Stress in Eating

The Stress in Eating (STRIvE) dataset was acquired from the University of Cambridge as a part of a multimodal neuroimaging study, which sought to examine the neurobiological correlates of disordered eating. The study was approved by the Cambridge East NHS Research Ethics Committee (HRA Ref. 17/EE/0304). Eighty-five women (age range 18–34 years) were recruited to three groups: those acutely ill with DSM-*5* anorexia nervosa (binge-eating/purging subtype; AN-BP), DSM-*5* bulimia nervosa (BN) and women with no personal history of mental illness. Participants underwent scanning on two consecutive days on a 3T Siemens SkyraFit MR scanner (Erlangen, Germany) at the Wolfson Brain Imaging Center at Addenbrooke's hospital in Cambridge, UK. Anatomical scans were co-registered with a linear transformation (AFNI program *3dAllineate*) and averaged across days via *3dMean.* The averaged structural image was then processed with FreeSurfer 6.0.1.

##### SYS Adults & Adolescents

The Saguenay Youth Study (SYS)[3] is a two-generational study of adolescents and their parents (N=1,029 adolescents and 962 parents) aimed at investigating the aetiology, early stages and trans-generational trajectories of common cardiometabolic and brain diseases[205,206]. High-resolution anatomical T1 images are acquired using the following parameters: 3D RF-spoiled

[3] https://academic.oup.com/ije/article/46/2/e19/2617159?login=true#112555109

gradient echo scan with 140–160 slices, 1mm isotropic resolution, TR=25 ms, TE=5 ms, flip angle=30° [207]. Scans were processed using FreeSurfer 5.3.

#### TEBC - Theirworld Edinburgh Birth Cohort

Theirworld Edinburgh Birth Cohort (TEBC) is a prospective longitudinal cohort study[208]. We plan to recruit 300 infants born at <33 weeks of gestational age (GA) and 100 CN infants born after 37 weeks of GA. Multiple domains are assessed: maternal and infant clinical and demographic information; placental histology; immunoregulatory and trophic proteins in umbilical cord and neonatal blood; brain macrostructure and microstructure from structural and diffusion MRI (dMRI); DNA methylation; hypothalamic-pituitary-adrenal axis activity; social cognition, attention and processing speed from eye tracking during infancy and childhood; neurodevelopment; gut and respiratory microbiota; susceptibility to viral infections; and participant experience. A Siemens MAGNETOM Prisma 3T MRI clinical scanner (Siemens Healthcare, Erlangen, Germany) and 16-channel phased-array paediatric head receive coil are used to acquire three-dimensional (3D) T1-weighted magnetisation-prepared rapid acquisition with gradient echo (MPRAGE) structural volume scan (acquired voxel size=1 mm isotropic); a 3D T2-weighted sampling perfection with application-optimized contrasts by using flip angle evolution (SPACE) structural scan (voxel size=1mm isotropic); and a multishell axial dMRI scan with optimal angular coverage. A subset has axial 3D susceptibility-weighted imaging and axial 2D fluid-attenuated inversion-recovery BLADE imaging, and magnetisation transfer saturation imaging is acquired for evaluation of tissue myelin content, consisting of three sagittal 3D multiecho spoiled gradient echo scans, 2mm isotropic acquired resolution, magnetisation-transfer, proton density-weighted and T1w acquisitions. For this project, tissue volumes were derived from structural data that were preprocessed using the developing Human Connectome Project (dHCP) minimal structural processing pipeline for neonatal data[88].

#### TOPSY

TOPSY refers to a high-resolution 7T MRI/spectroscopy study of untreated first episode psychosis (NCT02882204)[209]. Data were acquired on a Siemens MAGNETOM 7.0T MRI (Erlangen, Germany) using an 8-channel transmit/32-channel receive, head-only, radiofrequency coil at the Centre for Metabolic Mapping at Western University in London, Ontario. Images were first corrected for gradient nonlinearities using spherical harmonic coefficients provided by the manufacturer, and implemented in a BIDS app using spline interpolation and modulation by the determinant Jacobian of the warp (gradcorrect, https://github.com/khanlab/gradcorrect). In the current aggregated dataset a mixed sample of cognitively normal volunteers, individuals with schizophrenia and schizoaffective disorder for which the tissue segmentations were manually quality controlled were included.

#### UKB - UK Biobank

The UK BioBank (UKB) provides a unique, large and comprehensive dataset that includes both extensive phenotypic information as well as neuroimaging and genetics[38,210,211]. Structural minimally processed[4] T1- and T2-FLAIR weighted data was obtained from UK BioBank (application 20904) and further preprocessed with FreeSurfer 6.0.1[145] using the T2-FLAIR weighted image to improve pial surface reconstruction. Recon-all reconstruction included bias field correction, registration to stereotaxic space, intensity normalisation, skull-stripping, and white matter segmentation. A triangular surface tessellation fitted a deformable mesh model onto the white matter volume, providing grey-white and pial surfaces with >160,000 corresponding vertices registered to fsaverage standard space. When no T2-FLAIR image was available FreeSurfer reconstruction was done using the T1-weighted image only. Individuals were included in the reference dataset as healthy controls (CN) based on the response recorded in data-field 20544 (https://biobank.ndph.ox.ac.uk/showcase/field.cgi?id=20544) of the UKB mental health questionnaire, including only individuals who had never had mental health problem diagnosed by a mental health professional.

#### VETSA - Vietnam Era Twin Study of Aging

As outlined previously[43], VETSA is a longitudinal multi-modal (behavior, cognition, genetics, neuroimaging) study, comprising over 1,200 adult male twin pairs recruited from the Vietnam Era Twin Registry. All participants were in some branch of the United States military at some point between 1965 and 1975, with most not participating in combat or deployed in Vietnam. Data from ‘Wave 1’, which was conducted between 2003 and 2007, were included in the present study. Structural MRIs on 545 twins were collected on a Siemens 1.5T scanner, and were processed with FreeSurfer 6.0.1.

#### VITA - Vienna Transdanube Aging study

VITA is a population based cohort-study of all 75-years old inhabitants of a geographically defined area of Vienna. VITA is composed of 606 subjects followed longitudinally for 4 years. Recruitment took place between May, 2000 and October, 2002. The primary focus of the VITA work group was to establish a prospective age cohort for evaluation of prognostic criteria for the development of AD. All subjects derive from the prospective Vienna Trans-Danube Aging (VITA) study that targeted all 1,750 inhabitants of the age of 75 in the starting year of 2000 in two districts of Vienna and included irregular follow-ups until death, irrespective of clinical symptoms or diagnoses. All subjects featured in this analysis underwent one MRI measurement at the age of 75–76 years. Thereby, a 1T unit (Siemens Impact Expert; Siemens Medical Systems, Inc., South Iselin, NJ) and a circular polarized skull coil were used. For the present study all structural T1-weighted scans were processed with FreeSurfer 6.0.1.

[4] https://biobank.ctsu.ox.ac.uk/crystal/crystal/docs/brain_mri.pdf

### 20. Replication/validation datasets

We used several external datasets throughout the sensitivity analyses, all of which are described in more detail below.

##### 10k-in-a-day

The 10k-in-a-day dataset originated from a large collaborative international workshop where participants worked on 'connectomising' their own dataset for joined analysis[212]. The subsequent data has been made publicly available with minimal demographic information and age-binned age windows. Together, a total of 15,947 MRI datasets were processed using standard FreeSurfer pipelines and approximately 8000+ were made publicly available. While the lack of more detailed demographic data (e.g., age in years) and the likelihood that data may overlap with existing cohorts included in the present sample precluded us from using the full dataset into the original modelling. It did provide us with the opportunity to validate the generalisability of our derived data against a potentially noisy OoS dataset. We used the mean age of each 5-year age bracket, extracted the total grey matter volume from the dataset and compared these against the summed trajectories of GMV and sGMV generated by our model.

##### Hsu et al. (Ultrasound) - estimated

Using the software digitizeit (https://www.digitizeit.xyz/) we extracted total brain volume from Hsu et al.[213] who assessed the total volume and the blood flow index of the foetal brain in normal pregnancies using 3D ultrasound (Voluson 730). The study included 126 fetuses, ranging from 15 to 38 weeks of gestation. These total volumes were compared against the summed model trajectories from all 4 phenotypes.

##### Chang et al. (Ultrasound) - estimated

Analogous to Hsu et al.[213], this study evaluated foetal brain volume using 3D ultrasound. Chang and colleagues[214] measured total brain volume in 203 singleton fetuses ranged between 20 and 40 weeks of gestation. These total brain volumes were extracted using digitizeit (https://www.digitizeit.xyz/) and compared against the summed model trajectories from all 4 phenotypes.

##### Roelfsema et al. (Ultrasound) - estimated

Roelfsema et al.[215] used serial 3D sonography to measure foetal brain volume in 68 normal singleton pregnancies at 18 to 34 weeks of gestation. These 3D measurements were internally validated against foetal brain volume estimates from two-dimensional (2D) sonography measurement of head circumference and published postmortem foetal brain weights. We extracted these reported total brain volumes using digitizeit (https://www.digitizeit.xyz/) and compared them against the summed model trajectories from all 4 phenotypes.

#### Brain Weight

Postmortem brain weight estimates were available from the authors of two studies containing multiple historical data sources[41,216]. Additional postmortem brain weight data was aggregated from two recent large-scale genomic and transcriptomic initiatives GTEx[217] and PsychEncode [218]. The former was accessed through the GTEx access portal (https://www.gtexportal.org/home/) and dbGap (application https://www.ncbi.nlm.nih.gov/projects/gap/cgi-bin/study.cgi?study_id=phs000424.v8.p2), the latter through the PsychENCODE website (http://www.psychencode.org) and a Synapse application (https://www.synapse.org/#!Synapse:syn4921369/wiki/235539). These post-mortem brain weights were compared against the total cerebrum volume estimates obtained from the summed individual trajectories as a further validation of the trajectory slope.

#### International prenatal (HC)

The INTERGROWTH-21st consortium is a population-based project that assessed foetal growth and newborn size in eight geographically defined urban populations[219]. The project provided growth curve standards for head circumference during prenatal development. The head circumference values at each percentile (3rd, 5th, 10th, 50th, 90th, 95th, 97th) from 14–40 weeks gestation was averaged across males and females, and compared with total cerebrum volume estimates.

#### WHO postnatal (HC)

The WHO international standards on postnatal growth are an openly available resource for growth charts for height, weight, and head circumference[26]. These charts were used as input into digitizeit (https://www.digitizeit.xyz/) to extract the averaged percentile estimates across males and females, and were analyzed as above for the INTERGROWTH-21st cohort.

### 21. A note on data sharing

The complete dataset aggregated for the purposes of this study contains primary datasets that differ quite widely in terms of their "openness," i.e., their availability for secondary use without restrictions or special efforts by the primary study team. Primary studies ranged from fully open and downloadable datasets in the public domain to more restricted datasets that could only be used for specific purposes, under specific agreements, or after special efforts had been made to provide QC'd data in shareable form. There can be several reasons why data aren't always and immediately shared openly and/or without the active involvement of the researchers who collected the data[220]. In our experience within the context of this project, the various factors operating to prevent complete openness can be organised roughly into four categories:

- No informed consent was obtained for the open sharing of data at the time of collection[221] (or the informed consent does not extend to other uses in general).
- Data protection regulations, either at national or institutional levels, prevent the sharing of more detailed data such as essential demographics.
- The funding agency mandated or encouraged explicit involvement of researchers who collected primary study data in secondary studies where data was shared.
- Primary studies are still ongoing and data cannot be shared openly until the primary study objectives and/or milestones have been achieved.

There are also several reasons for not sharing data openly that cut across these categories such as general concerns about privacy or confidentiality of participants (which may be expressed by researchers, funders or governance bodies), as well as issues of data ownership (which are actively evolving as a result of changing legislation in some jurisdictions, e.g., General Data Protection Regulations [GDPR] in the European Union since 2016).

For these reasons, in practice, data is often shared under individually tailored and specific data usage or material transfer agreements. In the absence of a unified standard academic agreement this means that there is considerable variability in the terms under which data is or can be shared. For the present project, we sometimes had to make the difficult decision not to include potentially relevant datasets because abiding by the terms of the proposed sharing agreements would not have satisfied journal criteria for authorship and/or would have created an unbalanced acknowledgement of individual authors' contributions.

The benefits of truly open data are very clear from a scientific perspective. More open datasets would increase the number and diversity of researchers who are able to conduct secondary or meta-analytic studies without the need to negotiate multiple individual usage agreements. The present project would not have been possible without the availability of several exemplary open datasets[105,118,163,171,194,222–227], which were particularly valuable at the outset of this project, by facilitating pilot studies of brain charting methods. However, journal authorship criteria meant that we could not include members of some of the most open consortia as co-authors because their data were readily available to us without any significant additional contribution meriting authorship. We note that this situation potentially disincentives open science, since the people who do most to make their data openly available could be least likely to merit co-authorship of secondary studies. We therefore consider it is important for all stakeholders (funders, journals, investigators) to continue to think about how open human brain science can be properly recognised and rewarded. Here we have explicitly referenced and acknowledged our debt to the several open MRI datasets without which this study would not have been possible, because and although it has not always been appropriate to list the principal architects of these datasets as co-authors of this paper.

### 22. Affiliations of authors


[1] Autism Research Centre, Department of Psychiatry, University of Cambridge, Cambridge, CB2 0SZ, UK.
[2] Brain Mapping Unit, Department of Psychiatry, University of Cambridge, Cambridge, CB2 0SZ, UK
[3] Department of Psychiatry, University of Pennsylvania, Philadelphia, PA 19104
[4] Department of Child and Adolescent Psychiatry and Behavioral Science, The Children's Hospital of Philadelphia, Philadelphia, PA 19104
[5] Department of Psychiatry, University of Cambridge, Cambridge, CB2 0SZ, UK
[6] MRC Biostatistics Unit, University of Cambridge, Cambridge, England
[7] Lifespan Informatics & Neuroimaging Center, University of Pennsylvania, Philadelphia, PA 19104
[8] Department of Psychology, Yale University, New Haven, CT, USA
[9] Developmental Imaging, Murdoch Children's Research Institute, Melbourne, Victoria, Australia
[10] Department of Medicine, Monash University, Melbourne, Victoria, Australia
[11] UCL Great Ormond Street Institute for Child Health, 30 Guilford St, Holborn, London WC1N 1EH
[12] Weill Cornell Institute of Geriatric Psychiatry, Department of Psychiatry, Weill Cornell Medicine
[13] Department of Pediatrics University of Toronto
[14] Holland Bloorview Kids Rehabilitation Hospital, Toronto, Canada
[15] The Clinical Hospital of Chengdu Brain Science Institute, MOE Key Lab for NeuroInformation, University of Electronic Science and Technology of China, No. 2006, Xiyuan Ave., West Hi-Tech Zone, Chengdu, 611731, China
[16] University of Pinar del Río “Hermanos Saiz Montes de Oca”, Cuba
[17] MRC Cognition and Brain Sciences Unit, University of Cambridge, Cambridge UK
[18] Department of Psychology, School of Philosophy, Psychology and Language Sciences, University of Edinburgh, Edinburgh, United Kingdom
[19] University College London, Mental Health Neuroscience Research Department, Division of Psychiatry, London UK
[20] Department of Paediatrics, University of Melbourne, Melbourne, Victoria, Australia
[21] Cambridge Lifetime Asperger Syndrome Service (CLASS), Cambridgeshire and Peterborough NHS Foundation Trust, Cambridge, United Kingdom
[22] Centre for Addiction Medicine, National Institute of Mental Health and Neurosciences, Bengaluru, India 560029
[23] Department of Neurology, Max Planck Institute for Human Cognitive and Brain Sciences, Leipzig, 04103, Germany
[24] Department of Neuropsychiatry, Seoul National University Bundang Hospital, Seongnam, Korea
[25] Department of Human Genetics, South Texas Diabetes and Obesity Institute, University of Texas Rio Grande Valley
[26] MRC Centre for Reproductive Health, University of Edinburgh, UK
[27] foetal and Neonatal Institute, Division of Neonatology, Children's Hospital Los Angeles, Department of Pediatrics, Keck School of Medicine, University of Southern California, Los Angeles, California USA
[28] McGill Centre for Integrative Neuroscience, Ludmer Centre for Neuroinformatics and Mental Health, Montreal Neurological Institute
[29] McGill University
[30] Department of Brain Sciences, Imperial College London, London UK & Care Research & Technology Centre, UK Dementia Research Institute
[31] Tri-institutional Center for Translational Research in Neuroimaging and Data Science, Georgia State University, Georgia Institute of Technology, and Emory University, Atlanta, GA, USA

[32] Computational Brain Anatomy (CoBrA) Laboratory, Cerebral Imaging Centre, Douglas Mental Health University Institute
[33] Penn Statistics in Imaging and Visualization Center, Department of Biostatistics, Epidemiology, and Informatics, Perelman School of Medicine, University of Pennsylvania, Philadelphia, PA, USA
[34] Normandie Univ, UNICAEN, INSERM, U1237, PhIND "Physiopathology and Imaging of Neurological Disorders", Institut Blood and Brain @ Caen-Normandie, Cyceron, 14000 Caen, France
[35] Singapore Institute for Clinical Sciences, Agency for Science, Technology and Research, Singapore
[36] Department of Obstetrics and Gynaecology, Yong Loo Lin School of Medicine, National University of Singapore, Singapore
[37] Centre for Medical Image Computing (CMIC), University College London
[38] Dementia Research Centre (DRC), University College London
[39] Department of Psychiatry, Trinity College, Dublin, Ireland
[40] Cerebral Imaging Centre, Douglas Mental Health University Institute, Verdun, Canada
[41] Undergraduate program in Neuroscience, McGill University, Montreal, Canada
[42] Department of Neuroscience, University of California, San Diego, San Diego, CA 92093, USA
[43] Autism Center of Excellence, University of California, San Diego, San Diego, CA 92037, USA
[44] Institute of Neurodegenerative Disorders, CNRS UMR5293, University of Bordeaux
[45] Melbourne Neuropsychiatry Centre, University of Melbourne, Melbourne, Australia
[46] The Hospital for Sick Children, Toronto, Canada
[47] Department of Psychiatry, School of Medicine, Pontificia Universidad Católica de Chile, Diagonal Paraguay 362, Santiago 8330077, Chile
[48] Department of Psychosis Studies, Institute of Psychiatry, Psychology and Neuroscience, King's College London, De Crespigny Park, London SE5 8AF, UK
[49] Child and Adolescent Psychiatry Department, Robert Debré University Hospital, AP-HP, F-75019, Paris France
[50] Human Genetics and Cognitive Functions , Institut Pasteur, F-75015, Paris France
[51] Social, Genetic and Developmental Psychiatry Centre, Institute of Psychiatry, Psychology & Neuroscience, King's College London, London, United Kingdom
[52] Cerebral Imaging Centre, Douglas Mental Health University Institute, Montreal, QC, Canada
[53] Department of Psychiatry, McGill University, Montreal, QC, Canada
[54] Department of Psychiatry, Brigham and Women’s Hospital, Harvard Medical School, Boston, Massachusetts, United States
[55] Max Planck UCL Centre for Computational Psychiatry and Ageing Research, University College London, London, UK.
[56] Wellcome Centre for Human Neuroimaging, University College London, London, UK
[57] University of Cape Town, South Africa, Cape Town, South Africa
[58] Center for Neuroimaging, Cognition & Genomics (NICOG), School of Psychology, National University of Ireland Galway, Galway, Ireland
[59] Weil Family Brain and Mind Research Institute, Department of Psychiatry, Weill Cornell Medicine
[60] Centre for the Developing Brain, King's College London, London, UK
[61] Evelina London Children's Hospital
[62] Institute of Child Development, Department of Pediatrics, Masonic Institute for the Developing Brain, University of Minnesota, Minneapolis, MN, United States
[63] Department of Psychiatry, Center for Behavior Genetics of Aging, University of California, San Diego, La Jolla, CA
[64] Desert-Pacific Mental Illness Research Education and Clinical Center, VA San Diego Healthcare, San Diego, CA, USA
[65] Department of Psychiatry, University of California San Diego, Los Angeles, CA, USA
[66] Department of Psychiatry, University of Cambridge, and Wellcome Trust MRC Institute of Metabolic Science, Cambridge Biomedical Campus, Cambridge, United Kingdom
[67] Cambridgeshire and Peterborough NHS Foundation Trust
[68] Department of Clinical, Educational and Health Psychology, University College London, London, UK
[69] Department of Psychiatry, Center for Behavior Genetics of Aging, University of California, San Diego, La Jolla, CA 92093
[70] Cuban Center for Neuroscience, La Habana, Cuba
[71] Computational Radiology Laboratory, Boston Children’s Hospital, Boston, MA 02115

[72] Department of Child and Adolescent Psychiatry, University of California, San Diego, San Diego, CA 92093, USA
[73] Department of Psychiatry, University of California San Diego, San Diego, CA, USA
[74] Department of Psychiatry, University of North Carolina, Chapel Hill, NC, USA
[75] Department of Psychiatry, Boston Children's Hospital and Harvard Medical School, Boston, MA 02115
[76] Harvard Medical School, Boston, MA 02115
[77] Division of Newborn Medicine and Neuroradiology, foetal Neonatal Neuroimaging and Developmental Science Center, Boston Children's Hospital, Harvard Medical School, Boston, MA 02115, USA
[78] Department of Paediatrics and Child Health, Red Cross War Memorial Children's Hospital, SA-MRC Unit on Child & Adolescent Health, University of Cape Town, South Africa
[79] Neuroscience Institute, University of Cape Town, Cape Town, South Africa
[80] Lifespan Brain Institute, The Children's Hospital of Philadelphia, Philadelphia, PA 19104
[81] Lifespan Brain Institute, The Children's Hospital of Philadelphia, Philadelphia, PA 19105
[82] Mouse Imaging Centre, Toronto, Canada
[83] Clinical Memory Research Unit, Department of Clinical Sciences Malmö, Lund University, Malmö, Sweden
[84] Memory Clinic, Skåne University Hospital, Malmö, Sweden
[85] Department of Neurology, Icahn School of Medicine at Mount Sinai, New York, NY 10029, USA
[86] Athinoula A. Martinos Center for Biomedical Imaging, Department of Radiology, Massachusetts General Hospital, Harvard Medical School, Boston, MA 02129, USA
[87] Department of Psychiatry and Psychotherapy, Charite University Hospital Berlin, Berlin, Germany
[88] Department of Psychiatry, University of Cambridge, Cambridge, UK
[89] Department of Neuropsychology, Max Planck Institute for Human Cognitive and Brain Sciences, Leipzig, Germany
[90] Université de Paris, Paris, France
[91] Department of Psychiatry, University of Cape Town, Cape Town, South Africa.
[92] Departments of Psychiatry and Integrative Medicine, NIMHANS, Bengaluru, India
[93] Departments of Psychology and Psychiatry, Yale University, New Haven, CT, USA
[94] Radiology Research, Children's Hospital of Philadelphia, Philadelphia, United States
[95] The Department of Radiology, Perelman School of Medicine, University of Pennsylvania, Philadelphia, United States
[96] Division of Newborn Medicine, foetal Neonatal Neuroimaging and Developmental Science Center, Boston Children's Hospital, Harvard Medical School, Boston, MA 02115, USA
[97] Boston Children's Hospital, Boston, MA 02115
[98] Department of Psychiatry and Mental Health, Clinical Neuroscience Institute, University of Cape Town
[99] Department of Radiology, Mayo Clinic, Rochester, MN 55905, USA
[100] Department of Psychiatry, Universidade Federal de São Paulo
[101] National Institute of Developmental Psychiatry, CNPq
[102] Institute of Science and Technology for Brain-Inspired Intelligence, Fudan University, Shanghai, 200433, China
[103] Key Laboratory of Computational Neuroscience and BrainInspired Intelligence (Fudan University), Ministry of Education, Shanghai, China
[104] Centre for Population Neuroscience and Precision Medicine (PONS), Institute of Psychiatry, Psychology and Neuroscience, SGDP Centre, King's College London, London SE5 8AF, UK
[105] Harvard Aging Brain Study, Department of Neurology, Massachusetts General Hospital, Boston, MA 02114
[106] Center for Alzheimer Research and Treatment, Department of Neurology, Brigham and Women's Hospital, Boston, MA 02115
[107] Department of Radiology, Massachusetts General Hospital, Boston, MA
[108] Cambridgeshire and Peterborough NHS Foundation Trust, Huntingdon, United Kingdom
[109] Department of Neurology, Mayo Clinic, Rochester, MN, USA
[110] Department of Radiology, Mayo Clinic, Rochester, MN, USA
[111] Department of Psychiatry Division of Neurosciences, University Medical Center, Utrecht, The Netherlands
[112] Department of Psychiatry, Icahn School of Medicine, Mount Sinai, New York, USA
[113] Department of Clinical Medicine, Department of Psychiatry and Turku Brain and Mind Center, FinnBrain Birth Cohort Study, University of Turku and Turku University Hospital, Turku, Finland
[114] Centre for Population Health Research, Turku University Hospital and University of Turku, Turku, Finland
[115] Institute of Development, Aging and Cancer, Tohoku University, Seiryocho, Aobaku, Sendai 980-8575, Japan

[116] Queen's University, Departments of Psychology and Psychiatry, Centre for Neuroscience Studies, Kingston, Ontario, Canada
[117] Institute of Neuroscience and Physiology, University of Gothenburg, Gothenburg, Sweden
[118] Department of Brain and Cognitive Sciences, Seoul National University College of Natural Sciences, Seoul, Republic of Korea
[119] Department of Neuropsychiatry, Seoul National University Bundang Hospital, Seongnam, Republic of Korea
[120] Department of Psychiatry, Seoul National University College of Medicine, Seoul, Republic of Korea
[121] Department of Brain and Cognitive Science, Seoul National University College of Natural Sciences
[122] Section on Developmental Neurogenomics, Human Genetics Branch, National Institute of Mental Health, Bethesda, MD, USA
[123] Department of Brain & Cognitive Sciences, Seoul National University College of Natural Sciences, Seoul, Korea
[124] Department of Medical Biophysics, University of Toronto, Toronto, ON, Canada
[125] Mouse Imaging Centre, The Hospital for Sick Children, Toronto, ON, Canada
[126] Wellcome Centre for Integrative Neuroimaging, FMRIB, Nuffield Department of Clinical Neuroscience, University of Oxford, Oxford, UK
[127] Montreal Neurological Institute, McGill University, Montreal, Canada
[128] The Clinical Hospital of Chengdu Brain Science Institute, University of Electronic Science and Technology of China, Chengdu 611731, China
[129] Department of Psychiatry and Brain and Mind Research Institute, Weill Cornell Medicine
[130] Laboratory for Autism and Neurodevelopmental Disorders, Center for Neuroscience and Cognitive Systems @UniTn, Istituto Italiano di Tecnologia, Rovereto, Italy
[131] School of Biomedical Engineering & Brain and Mind Centre, The University of Sydney, Sydney, NSW, Australia
[132] Department of Psychology, University of Texas, Austin, Texas 78712, USA
[133] Department of Psychiatry and Neuropsychology, School of Mental Health and Neuroscience, EURON, Maastricht University Medical Centre, PO Box 616, 6200 MD, Maastricht, the Netherlands; Institute for Mental Health Care Eindhoven (GGzE), Eindhoven, the Netherlands.
[134] McConnell Brain Imaging Centre, Montreal Neurological Institute, McGill University, Montreal, QC H3A 2B4, Canada
[135] Bordeaux University Hospital
[136] Professor, Department of Psychosis Studies, Institute of Psychiatry, Psychology and Neuroscience, King's College London, UK
[137] Ludmer Centre for Neuroinformatics and Mental Health, Douglas Mental Health University Institute, McGill University, Montreal, Quebec, Canada; Singapore Institute for Clinical Sciences, Singapore
[138] Department of Computer Science and Technology, University of Cambridge, Cambridge CB3 0FD, United Kingdom
[139] Department of Psychiatry, University of Cambridge, Cambridge CB2 0SZ, United Kingdom
[140] The Alan Turing Institute, London, NW1 2DB
[141] Department of Psychology, School of Business, National College of Ireland, Dublin, Ireland
[142] School of Psychology & Center for Neuroimaging and Cognitive Genomics, National University of Ireland Galway, Galway, Ireland
[143] Department of Psychiatry, Trinity College Dublin, Dublin, Ireland
[144] Department of Psychiatry, School of Medicine, Oregon Health and Science University, Portland, United States
[145] Center for Sleep and Cognition, Yong Loo Lin School of Medicine, National University of Singapore, Singapore
[146] Department of Pediatrics, Washington University in St. Louis, St. Louis, Missouri, United States
[147] Alzheimer Center Amsterdam, Department of Neurology, Amsterdam Neuroscience, Vrije Universiteit Amsterdam, Amsterdam UMC, Amsterdam, The Netherlands
[148] Lund University, Clinical Memory Research Unit, Lund, Sweden
[149] Robarts Research Institute & The Brain and Mind Institute, University of Western Ontario,London,Ontario,Canada.
[150] Department of Psychiatry, Federal University of Sao Poalo (UNIFESP)
[151] National Institute of Developmental Psychiatry for Children and Adolescents (INPD), Brazil.
[152] Melbourne Neuropsychiatry Centre, Department of Psychiatry, The University of Melbourne and Melbourne Health, Carlton South, Victoria, Australia
[153] Melbourne School of Engineering, The University of Melbourne, Parkville, Victoria, Australia
[154] Florey Institute of Neuroscience and Mental Health, Parkville, VIC, Australia

[155] Department of Psychiatry, Schulich School of Medicine and Dentistry, Western University, London, ON, Canada
[156] Department of Psychiatry, Faculty of Medicine and Centre Hospitalier Universitaire Sainte-Justine, University of Montreal, Montreal, Quebec, Canada
[157] Departments of Psychiatry and Psychology, University of Toronto, Toronto, ON, Canada.
[158] Departments of Physiology and Nutritional Sciences, University of Toronto, Toronto, Canada
[159] Cuban Neuroscience Center, Havana, Cuba
[160] Department of Psychiatry, Faculty of Medicine, McGill University, Montreal, Qc, H3A 1Y2, Canada
[161] Douglas Mental Health University Institute, Montreal, Qc, H4H 1R3, Canada.
[162] Autism Center of Excellence, Department of Neurosciences, University of California, San Diego La Jolla, CA, USA
[163] School of Psychology, Southwest University, Chongqing 400715, P.R. China
[164] Department of Biomedical Engineering, The N.1 Institute for Health, National University of Singapore.
[165] Department of Clinical Neurosciences, University of Cambridge, Cambridge UK
[166] Department of Neurology, Harvard Medical School
[167] Department of Neurology, Boston Children's Hospital, Boston, MA 02115
[168] Instituto de Biomedicina de Sevilla (IBiS) HUVR/CSIC/Universidad de Sevilla, Dpto. de Fisiología Médica y Biofísica, Spain
[169] Department of Psychology, Neuroscience Institute, University of Chicago
[170] Department of Paediatrics and Wellcome-MRC Cambridge Stem Cell Institute, University of Cambridge, Hills Road, Cambridge, UK
[171] Department of Psychiatry, Universidade Federal do Rio Grande do Sul (UFRGS)
[172] National Institute of Developmental Psychiatry (INPD)
[173] Otto Hahn Group Cognitive Neurogenetics, Max Planck Institute for Human Cognitive and Brain Sciences, Leipzig, Germany
[174] Institute of Neuroscience and Medicine (INM-7: Brain and Behaviour), Research Centre Juelich, Juelich, Germany
[175] Athinoula A. Martinos Center for Biomedical Imaging, Department of Radiology, Massachusetts General Hospital, Charlestown, MA 02129, USA
[176] Centre for Population Neuroscience and Stratified Medicine (PONS), SGDP Centre, IoPPN, KCL, UK
[177] PONS-Centre, Dept of Psychiatry and Psychotherapy, Campus Charite Mitte, Humboldt University, Berlin, Germany
[178] PONS Centre, Institute for Science and Technology of Brain-inspired Intelligence (ISTBI), Fudan University, Shanghai, China
[179] Wallenberg Centre for Molecular and Translational Medicine, University of Gothenburg, Gothenburg, Sweden
[180] Department of Psychiatry and Neurochemistry, University of Gothenburg, Sweden
[181] Dementia Research Centre, Queen's Square Institute of Neurology, University College London, UK
[182] Center for Biomedical Image Computing and Analytics, Department of Radiology, Perelman School of Medicine, University of Pennsylvania, Philadelphia, PA, USA
[183] Departments of Neurology, Pediatrics, and Radiology, Washington University School of Medicine, St. Louis, United States
[184] SA MRC Unit on Risk & Resilience in Mental Disorders, Dept of Psychiatry and Neuroscience Institute, University of Cape Town, Cape Town, South Africa
[185] Division of Psychiatry, Centre for Clinical Brain Sciences, University of Edinburgh, UK
[186] Department of Neuroscience, Institut Pasteur, Paris, France
[187] Center for Research and Interdisciplinarity (CRI), Université Paris Descartes, Paris, France
[188] Department of Psychology, University of Cambridge, Cambridge, UK
[189] Wu Tsai Institute, Yale University, New Haven, CT, USA
[190] Department of Clinical Medicine, Department of Psychiatry, FinnBrain Birth Cohort Study, University of Turku, Turku, Finland
[191] Department of Clinical Medicine, University of Turku, Turku Finland
[192] Turku Collegium for Science, Medicine and Technology, University of Turku, Turku, Finland
[193] Univ. Bordeaux, Inserm, Bordeaux Population Health Research Center, U1219, CHU Bordeaux, F-33000 Bordeaux, France
[194] Department of Anesthesia, Faculty of Medicine, McGill University, Montreal, Qc, H3A 1G1, Canada
[195] Faculty of Dentistry, McGill University, Montreal, Qc, H3A 1G1, Canada
[196] Alan Edwards Centre for Research on Pain (AECRP), McGill University, Montreal, Qc, H3A 1G1, Canada

[197] Joint China-Cuba Lab,University of Electronic Science and Technology, Chengdu China/Cuban Center for Neuroscience, La Habana, Cuba
[198] University of Electronic Science and Technology of China/Cuban Center for Neuroscience
[199] Institute for Neuroscience and Medicine 7, Forschungszentrum Juelich; Max Planck Institute for Human Cognitive and Brain Sciences
[200] Department of Psychiatry & Neurosychology, Maastricht University, Maastricht, The Netherlands
[201] Department of Biostatistics, Vanderbilt University, Nashville, Tennessee, USA
[202] Department of Biostatistics, Vanderbilt University Medical Center, Nashville, Tennessee, USA
[203] Division of Newborn Medicine, foetal Neonatal Neuroimaging and Developmental Science Center, Department of Pediatrics, Boston Children's Hospital, Boston, MA 02115
[204] McConnell Brain Imaging Center, Montreal Neurological Institute, McGill University, Montreal, Quebec, Canada
[205] Clinic for Cognitive Neurology, University of Leipzig Medical Center, Leipzig, 04103, Germany
[206] The Alan Turing Institute, London NW1 2DB, UK
[207] Wellcome Centre for Human Neuroimaging, Institute of Neurology, University College London, WC1N 3AR
[208] State Key Laboratory of Cognitive Neuroscience and Learning, Beijing Normal University, Beijing 100875, China
[209] Developmental Population Neuroscience Research Center, IDG/McGovern Institute for Brain Research, Beijing Normal University, Beijing 100875, China
[210] National Basic Science Data Center, Beijing 100190, China
[211] Research Center for Lifespan Development of Brain and Mind, Institute of Psychology, Chinese Academy of Sciences, Beijing 100101, China
[212] Division of Clinical Geriatrics, Center for Alzheimer Research, Department of Neurobiology, Care Sciences and Society, Karolinska Institutet, Stockholm, Sweden
[213] Faculty of Medicine, CRC 1052 'Obesity Mechanisms', University of Leipzig, Leipzig, 04103, Germany
[214] Department of Electrical and Computer Engineering, National University of Singapore, Singapore
[215] Centre for Sleep & Cognition and Centre for Translational MR Research, Yong Loo Lin School of Medicine, National University of Singapore, Singapore
[216] N.1 Institute for Health & Institute for Digital Medicine, National University of Singapore, Singapore
[217] Integrative Sciences and Engineering Programme (ISEP), National University of Singapore, Singapore
[218] foetal Neonatal Neuroimaging and Developmental Science Center, Division of Newborn Medicine, Boston Children's Hospital, Harvard Medical School, Boston, MA 02115, USA
[219] Melbourne Neuropsychiatry Centre, University of Melbourne, Melbourne, Australia; Department of Biomedical Engineering, University of Melbourne, Melbourne, Australia.
[220] SAMRC Unit on Child & Adolescent Health, University of Cape Town, South Africa
[221] Center for Translational Magnetic Resonance Research, Yong Loo Lin School of Medicine, National University of Singapore, Singapore
[222] Wellcome Trust-MRC Institute of Metabolic Science,  University of Cambridge, Cambridge, CB2 0SZ
[223] Cambridgeshire and Peterborough Foundation Trust, Cambridge, CB21 5EF
[224] National Institute of Mental Health (NIMH), National Institutes of Health (NIH), Bethesda, Maryland, USA
[225] Department of Psychiatry, Escola Paulista de Medicina, São Paulo, Brazil.
[226] Key Laboratory of Brain and Education, School of Education Science, Nanning Normal University, Nanning 530001, China
* Data used in this article were obtained from the brain consortium for reproducibility, reliability and replicability (3R-BRAIN).
** Data used in the preparation of this article was obtained from the Australian Imaging Biomarkers and Lifestyle flagship study of ageing (AIBL) funded by the Commonwealth Scientific and Industrial Research Organisation (CSIRO) which was made available at the ADNI database (www.loni.usc.edu/ADNI). The AIBL researchers contributed data but did not participate in analysis or writing of this report. AIBL researchers are listed at www.aibl.csiro.au.
*** Data used in preparation of this article were obtained from the Alzheimer's Disease Neuroimaging Initiative (ADNI) database (adni.loni.usc.edu). As such, the investigators within the ADNI contributed to the design and implementation of ADNI and/or provided data but did not participate in analysis or writing of this report. A complete listing of ADNI investigators can be found at: http://adni.loni.usc.edu/wp-content/uploads/how_to_apply/ADNI_Acknowledgement_List.pdf
**** A complete listing of ARWiBo researchers can be found in the Supplementary Materials

***** Data used in this article were collected and processed by the UMN Baby Connectome Project (BCP) consortium and the Masonic Institute for the Developing Brain (MIDB) Informatics group. The groups are comprised of the following individuals: Amanda Ruetger, Audrey Houghton, Ben Lynch, Thomas Pengo, Jim Wilgenbusch, Tim Hendrickson, Trevor Day, Sooyeon Sung, Kathy Snider, Lucille Moore, Alice Graham, Damien Fair, Eric Feczko and Jed Elison
****** The Centre for Attention Learning and Memory (CALM) research clinic is based at and supported by funding from the Medical Research Council Cognition and Brain Sciences Unit, University of Cambridge, as well as funding from the EU Horizon 2020 Personalised Medicine “LifeBrain” project (H2020-SC1-2016-2017, Topic SC1-PM-04-2016). The lead investigators are Duncan Astle, Kate Baker, Susan E. Gathercole, Joni Holmes, Rogier A. Kievit, and Tom Manly. Data collection was assisted by a team of researchers and PhD students that includes Danyla Akarca, Joe Bathelt,Giacomo Bignardi, Sarah Bishop, Erica Botanic, Lara Bridge, Diandra Bkric, Annie Bryant, Sally Butterfield, Elizabeth Byrne, Gemma Crickmore, Edwin Dalmaijer; Fánchea Daly, Tina Emery, Grace Franckel, Laura Forde, Delia Fuhrmann, Andrew Gadie, Sara Gharooni, Jacalyn Guy, Erin Hawkins, Agniezska Jaroslawska, Sara Joeghan, Amy Johnson, Jonathan Jones, Elise Ng-Cordell, Sinéad O’Brien, Cliodhna O’Leary, Joseph Rennie, Ivan Simpson-Kent, Roma Siugzdaite, Tess Smith, Stepheni Uh, Francesca Woolgar, Mengya Zhang, and Natalia Zdorovtsova. We thank the many professionals working in children’s services in the southeast and east of England for their support and to the children and their families for giving up their time to visit the clinic, and the radiographer for facilitating pediatric scanning.
******* The Cambridge Centre for Ageing and Neuroscience (Cam-CAN) research was supported by the Biotechnology and Biological Sciences Research Council (BB/H008217/1), as well as funding from the Medical Research Council Cognition and Brain Sciences Unit, and the EU Horizon 2020 Personalised Medicine “LifeBrain” project (H2020-SC1-2016-2017, Topic SC1-PM-04-2016). We thank the Cam-CAN respondents and their primary care teams in Cambridge for their participation in this study. Further information about the Cam-CAN corporate authorship membership can be found at http://www.cam-can.org/index.php?content=corpauth#12.
******** Data used in this article were obtained from the developmental component 'Growing Up in China' of Chinese Color Nest Project (http://deepneuro.bnu.edu.cn/?p=163).
********* Data was downloaded from the COllaborative Informatics and Neuroimaging Suite Data Exchange tool (COINS; http://coins.mrn.org/dx) and data collection was performed at the Mind Research Network, and funded by a Center of Biomedical Research Excellence (COBRE) grant 5P20RR021938/P20GM103472 from the NIH to Dr. Vince Calhoun.
********** The ENIGMA Developmental Brain Age working group principally consists of Drs. James Cole, Niall Bourke, Heather Whalley, David Glahn, Laura Han, Francesca Biondo, Katherine Karlsgodt, Carrie Bearden, Jakob Seidlitz, Richard Bethlehem, Eileen Xu, Marieke Bos, Sam Mathia, Sophia Frangou, Miruna Carmen Barbu, Yoonho Chung, and Aaron Alexander-Bloch
*********** Data used in the preparation of this article were obtained from the Harvard Aging Brain Study (HABS - P01AG036694; https://habs.mgh.harvard.edu). The HABS study was launched in 2010, funded by the National Institute on Aging. and is led by principal investigators Reisa A. Sperling MD and Keith A. Johnson MD at Massachusetts General Hospital/Harvard Medical School in Boston, MA.
************ Data used in the preparation of this article were obtained from the IMAGEN consortium.
************* Data used in this article were obtained from the Korean Longitudinal Study on Cognitive Aging and Dementia (KLOSCAD) (https://recode.re.kr).
************** A full list of NSPN consortium members can be found at: https://www.nspn.org.uk/nspn-team/
*************** The POND network is a Canadian translational network in neurodevelopmental disorders, primarily funded by the Ontario Brain Institute.

### 23. Acknowledgements

RAIB is supported by a British Academy Postdoctoral fellowship and Autism Research Trust. JS is supported by NIMH T32 MH019112-29 and K08MH120564. SRW was funded by UKRI Medical Research Council MC_UU_00002/2 and was supported by the NIHR Cambridge Biomedical Research Centre (BRC-1215-20014). ETB is supported by an NIHR Senior Investigator award. AFA-B is supported by NIMH K08MH120564. Data were curated and analysed using a computational facility funded by an MRC research infrastructure award (MR/M009041/1) and supported by the NIHR Cambridge Biomedical Research Centre. The views expressed are those of the authors and not necessarily those of the NIH, NHS, the NIHR or the Department of Health and Social Care. E.A and POND collection is funded by the Ontario Brain Institute;D.E.A. was supported by MRC Programme Grant MC-A0606-5PQ41;M.A. is supported by NIH 1R01MH112904-01;G.B. was supported by an NHMRC Investigator Grant (#1194497) and the Royal Children's Hospital Foundation.;cVEDA is jointly funded by the Indian Council for Medical Research (ICMR/MRC/3/M/2015-NCD-I) and the Newton Grant from the Medical Research Council(MR/N000390/1), United Kingdom.;The Genetics of Brain Structure and Function project was funnded by the NIH (MH078143, MH083824, and MH078111);JBB was supported by Brain Canada (243030), the Fonds de recherche du Québec (FRQ) Healthy Brains Healthy Liles (HBHL) FRQ/Canada-Cuba-China Axis (246117), the Canada First Research Excellence Fund (CFREF)/HBHL BigBrain Analytics and Learning Laboratory (HIBALL), and Helmholtz (252428) ;IMAP study (PI (scientific): G Chetelat; PI (MD) V de La Sayette)) was funded by Programme Hospitalier de Recherche Clinique (PHRCN 2011-A01493-38 and PHRCN 2012 12-006-0347) and Agence Nationale de la Recherche (LONGVIE 2007). Dr Chetelat's research including IMAP was also funded by Institut National de la Santé et de la Recherche Médicale (Inserm), Fondation Plan Alzheimer (Alzheimer Plan 2008-2012); Région Basse-Normandie; Association France Alzheimer et maladies apparentées, Fondation Vaincre Alzheimer.;E.C. is supported by grants: NIMH P50-MH081755, NIMH R01-MH036840, NIMH R01-MH110558, NIMH U01-MH108898, NIDCD R01-DC016385;The Developing Human Connectome Project was supported by the European Research Council under the European Union Seventh Framework Programme (FP/2007-2013)/ERC Grant Agreement No. 319456;The BCP was supported by grants: U01MH110274 & R01MH104324;PCF is supported by the Wellcome Trust (Reference No. Reference No. 206368/Z/17/Z) and by the Bernard Wolfe health Neuroscience Fund;C.E.F. is supported by grants: R01s AG050595, AG022381, AG037985; & P01 AG055367; The content is the sole responsibility of the authors and does not necessarily represent official views of the NIA,

NIH, or VA. The U.S. Department of Veterans Affairs, Department of Defense; National Personnel Records Center, National Archives and Records Administration; National Opinion Research Center; National Research Council, National Academy of Sciences; and the Institute for Survey Research, Temple University provided invaluable assistance in the creation of the VET Registry. The Cooperative Studies Program of the U.S. Department of Veterans Affairs provided financial support for development and maintenance of the Vietnam Era Twin Registry. We would also like to acknowledge the continued cooperation and participation of the members of the VET Registry and their families.;A.G. is supported by grants: NIH R01 EB018988, and R01 NS106030;The Genetics of Brain Structure and Function project was funded by the NIH (MH078143, MH083824, and MH078111);RNH is supported by MRC Programme Grant SUAG/046 G101400;"The CHILD study was funded by the Autism Research Trust. RH received funding from the Innovative Medicines Initiative 2 Joint Undertaking (JU) under grant agreement No 777394. The JU receives support from the European Union's;Horizon 2020 research and innovation programme and EFPIA and AUTISM SPEAKS, Autistica, SFARI.; ";FinnBrain was funded by Jane and Aatos Erkko Foundation, Signe and Ane Gyllenberg Foundation;LK was funded by the Brain and Behavior Research Foundation, NARSAD YI Grant #1956 and the Academy of Finland Profi 5 #325292;E.A.K. is funded by the Masonic Foundation of Ontario, the National Institute of Mental Health, and the Ontario Brain Institute;SK was financed by grants from the Swedish state under the agreement between the Swedish government and the county councils, the ALF-agreement ( ALFGBG-81392, ALF GBG-771071). The Alzheimerfonden (AF-842471, AF-737641, AF-939825 ). The Swedish Research Council (2019-02075);The KNE96 study was supported by a grant of the Korean Health Technology R&D Project, Ministry for Health, Welfare, and Family Affairs, Republic of Korea (grant number I09C1379(A092077);W.S.K. was supported by NIA grants R01s AG050595, AG022381, AG037985, and P01 AG055367. ;The POND study was supported by the Ontario Brain Institute (grant number IDS-I 1-02). This organisation did not play a role in thedesign of the study, the collection, analysis and interpretation ofthe data, and in writing the manuscript.;BHRCS was supported with grants from the National Institute of Development Psychiatric for Children and Adolescent (INPD). Grant: Fapesp 2014/50917-0 - CNPq 465550/2014-2;CP was supported by a National Health and Medical Research Council (NHMRC) Senior Principal Research Fellowship (1105825), an NHMRC L3 Investigator Grant (1196508) and NHMRC Program Grant (ID: 1150083). ;K.P. is supported by grants: NIMH R01-MH080134, NIMH R01-MH104446;This research/project is supported by the National Science Foundation (NSF:2010778) and National Research Foundation, Singapore under its AI Singapore Programme (AISG Award No: AISG-GC-2019-002). Additional funding is provided by the

Singapore Ministry of Education (Academic research fund Tier 1; NUHSRO/2017/052/T1-SRP-Partnership/01), NUS Institute of Data Science;ADNI: NIH funding. Dataused in preparation of this article were obtained from the Alzheimer's Disease Neuroimaging Initiative (ADNI) database (adni.loni.usc.edu).As such, the investigators within the ADNI contributed to the design and implementation of ADNI and/or provided data but did not participate in analysis or writing of this report. A complete listing of ADNI investigators can be found at: http://adni.loni.usc.edu/wp-content/uploads/how_to_apply/ADNI_Acknowledgement_List.pdf.
NACC: The NACC database is funded by NIA/NIH Grant U01 AG016976. NACC data are contributed by the NIA-funded ADCs: P30 AG019610 (PI Eric Reiman, MD), P30 AG013846 (PI Neil Kowall, MD), P50 AG008702 (PI Scott Small, MD), P50 AG025688 (PI Allan Levey, MD, PhD), P50 AG047266 (PI Todd Golde, MD, PhD), P30 AG010133 (PI Andrew Saykin, PsyD), P50 AG005146 (PI Marilyn Albert, PhD), P50 AG005134 (PI Bradley Hyman, MD, PhD), P50 AG016574 (PI Ronald Petersen, MD, PhD), P50 AG005138 (PI Mary Sano, PhD), P30 AG008051 (PI Thomas Wisniewski, MD), P30 AG013854 (PI Robert Vassar, PhD), P30 AG008017 (PI Jeffrey Kaye, MD), P30 AG010161 (PI David Bennett, MD), P50 AG047366 (PI Victor Henderson, MD, MS), P30 AG010129 (PI Charles DeCarli, MD), P50 AG016573 (PI Frank LaFerla, PhD), P50 AG005131 (PI James Brewer, MD, PhD), P50 AG023501 (PI Bruce Miller, MD), P30 AG035982 (PI Russell Swerdlow, MD), P30 AG028383 (PI Linda Van Eldik, PhD), P30 AG053760 (PI Henry Paulson, MD, PhD), P30 AG010124 (PI John Trojanowski, MD, PhD), P50 AG005133 (PI Oscar Lopez, MD), P50 AG005142 (PI Helena Chui, MD), P30 AG012300 (PI Roger Rosenberg, MD), P30 AG049638 (PI Suzanne Craft, PhD), P50 AG005136 (PI Thomas Grabowski, MD), P50 AG033514 (PI Sanjay Asthana, MD, FRCP), P50 AG005681 (PI John Morris, MD), P50 AG047270 (PI Stephen Strittmatter, MD, PhD).;Supported the NIH including the NINDS K23NS101120 (C.K.R.), the NHLBI K23HL141602 (C.M.O), NIBIB R01EB013248 (S.K.W.), R01EB018988 and R01NS106030 (A.G.), and a NHLBI Pediatric Heart Network Scholar Award (C.K.R.); the American Academy of Neurology Clinical Research Training Fellowship (C.K.R.); the Brain and Behavior Research Foundation NARSAD Young Investigator (C.K.R.) and Distinguished Investigator (S.K.W.) Awards; the McKnight Foundation Technological Innovations in Neuroscience Award (A.G.); Office of Faculty Development at Boston Children's Hospital Career Development Awards (A.G., C.K.R.); and the Mend A Heart Foundation (C.M.O.). ;RRG was supported by the Guarantors of Brain, Cancer Research UK Cambridge Centre ;LR was supported by the Bernard Wolfe Health Neuroscience Fellowship;M.D.R. is supported by Bill & Melinda Gates Foundation INV-015711;NIHR Cambridge BRC;T.D.S. is supported by grants: NIH R01MH112847, R01MH120482, & R01MH113550 ;M.S. is supported by the Knut and Alice

Wallenberg Foundation (Wallenber Centre for Molecular and Translational Medicine), the Swedish Research Council and the Swedish State under the ALF-agreement.;RTS is supported by R01MH112847.;STRADL study was supported and funded by the Wellcome Trust Strategic Award “Stratifying Resilience and Depression Longitudinally” (Reference 104036/Z/14/Z). Processing of the dataset used the resources provided by the Edinburgh Compute and Data Facility (ECDF) (http://www.ecdf.ed.ac.uk/).;K.A.T. was funded by the Guarantors of Brain (G101149).;The preparation and initiation of the i-Share project was funded by the program ‘Invest for future’ (reference ANR-10-COHO-05). The i-Share Project is currently supported by an unrestricted grant of the Nouvelle-Aquitaine Regional Council (Conseil Régional Nouvelle-Aquitaine) (grant N° 4370420) and by the Bordeaux ‘Initiatives d’excellence’ (IdEx) program of the University of Bordeaux (ANR-10-IDEX-03-02). It has also received grants from the Nouvelle-Aquitaine Regional Health Agency (Agence Régionale de Santé Nouvelle-Aquitaine, grant N°6066R-8), Public Health France (Santé Publique France, grant N°19DPPP023-0), , and The National Institute against cancer INCa (grant N°INCa_11502).;SNV was supported by NIH R01MH123563;K.S.W. was supported by the Wellcome Trust (215901/Z/19/Z);S.K.W. was supported by NIH R01EB013248;S.R.W was funded by UKRI Medical Research Council MC_UU_00002/2 and was supported by the NIHR Cambridge Biomedical Research Centre (BRC-1215-20014). The views expressed are those of the author(s) and not necessarily those of the NIHR or the Department of Health and Social Care;A.V.W. was supported by grants from the German Research Foundation (WI 3342/3-1; 209933838-02);B.T.T.Y. is funded by the Singapore National Research Foundation (NRF) Fellowship (Class of 2017), the NUS Yong Loo Lin School of Medicine (NUHSRO/2020/124/TMR/LOA), the Singapore National Medical Research Council (NMRC) LCG (OFLCG19May-0035) and the USA NIH (R01MH120080).;The DCHS was funded by the Bill & Melinda Gates Foundation (OPP1017641). HJZ is funded by the SA-MRC;J.H.Z. is funded by National Medical Research Council, Singaore and the National University of Singapore Yong Loo Lin School of Medicine (NUHSRO/2020/124/TMR/LOA);X.N.Z is funded by the Startup Funds for Leading Talents at Beijing Normal University, the National Natural Science Foundation of China (81220108014), the National Basic Research (973) Program (2015CB351702), the National Basic Science Data Center `Chinese Data-sharing Warehouse for In-vivo Imaging Brain' Program (NBSDC-DB-15), the Beijing Municipal Science and Tech Commission (Z161100002616023, Z181100001518003), the China - Netherlands CAS-NWO Programme (153111KYSB20160020), Guangxi BaGui Scholarship (201621), the Key Realm R&D Program of Guangdong Province (2019B030335001), and the Major Project of National Social Science Foundation of China (20&ZD296); LIFE is funded by means of the European Union, by the

European Regional Development Fund (ERDF) and by funds of the Free State of Saxony within the framework of the excellence initiative. A.V.W. was supported by grants from the German Research Foundation (WI 3342/3-1; 209933838-02).
